# Sequence- and chemical specificity define the functional landscape of intrinsically disordered regions

**DOI:** 10.1101/2022.02.10.480018

**Authors:** Iris Langstein-Skora, Andrea Schmid, Frauke Huth, Drin Shabani, Lorenz Spechtenhauser, Mariia Likhodeeva, Franziska Kunert, Felix J. Metzner, Ryan J. Emenecker, Mary O.G. Richardson, Wasim Aftab, Maximilian J. Götz, Sarah K. Payer, Niccoló Pietrantoni, Valentina Sjeničić, Sakthi K. Ravichandran, Till Bartke, Karl-Peter Hopfner, Ulrich Gerland, Philipp Korber, Alex S. Holehouse

## Abstract

Intrinsically disordered protein regions (IDRs) pervasively engage in essential molecular functions, yet they are often poorly conserved as assessed by sequence alignment. To understand the seeming paradox of how sequence variability is compatible with persistent function, we examined the functional determinants for a poorly conserved but essential IDR. We show that IDR function depends on two distinct but related properties: sequence- and chemical specificity. While sequence-specificity works via linear binding motifs, chemical specificity reflects the sequence-encoded chemistry of multivalent interactions through amino acids across an IDR. Unexpectedly, a binding motif that is essential in the wild-type IDR can be removed if compensatory changes to the sequence chemistry are made, highlighting the orthogonality and interoperability of both properties and providing a much deeper sequence space compatible with function. Our results provide a general framework to understand the functional constraints on IDR sequence evolution.

## INTRODUCTION

Intrinsically disordered proteins and protein regions (collectively referred to as IDRs) play important and often essential roles in many biological processes across all three kingdoms of life (*1*). IDRs frequently engage in molecular interactions, and their inherent structural plasticity allows them to bind through a variety of mechanisms (*1–3*). As such, understanding how IDR sequences determine the modes of molecular recognition is key to mapping from sequence to function.

Despite their importance for function, IDRs are often poorly conserved as assessed by alignment-based methods (*4–7*). The notable exceptions to this are short linear motifs (SLiMs); 5-15 amino acid modules which often contain multiple conserved positions in a consensus motif and engage in sequence-specific binding (*8–10*). The modular nature of many SLiMs is demonstrated by their ability to mediate molecular recognition when inserted into otherwise neutral contexts (*9*, *11*). In addition to molecular recognition driven by SLiMs, IDRs that lack SLiMs can engage in distributed multivalent interactions driven by sequence-encoded chemistry (*1*, *3*, *12–15*). These interactions can mediate both stoichiometric protein-protein interactions or drive the formation of biomolecular condensates (*16–20*). For these distributed multivalent interactions, the precise sequence order is often unimportant, yet the amino acid composition and patterning are critical (*16*, *17*, *21*). These two modes of interaction are generally considered to drive orthogonal types of molecular recognition – that is, loss of a SLiM cannot be compensated for by changing the degree of multivalency outside of the SLiM (*16*, *22*). This orthogonality can also be considered a reflection of the degree of order in the bound state (*23*). With this in mind, substantial sequence variation appears to be tolerated in many IDRs as long as conserved positions in SLiMs are retained and/or the amino acid composition and patterning (i.e., bulk sequence properties) are maintained (*5*, *24–26*).

Here, using *S. cerevisiae* as a model organism, we sought to investigate the interplay between these two modes of interaction in the context of conservation and function in an essential IDR. Our results suggest that rather than existing as two distinct modes of interaction, IDR-mediated molecular recognition should be considered on a two-dimensional landscape, whereby SLiMs cooperate with and can even be fully replaced by distributed multivalent interactions. Accordingly, apparently modular sequence motifs may impart function not by acting as *bona fide* SLiMs, but simply by altering the overall sequence chemistry. Our results imply that sequence context and the presence/absence of SLiMs can compensate for and buffer one another, providing a key missing piece in our understanding of the sequence constraints on IDR evolution.

## RESULTS

### Proteome-wide analyses reveal weak alignment-based conservation for yeast IDRs

IDRs often undergo more substantial sequence variation than folded domains (*4*, *6*). We therefore anticipated that *S. cerevisiae* proteins may possess conserved folded regions alongside less well-conserved IDRs (**Fig. 1A**). To assess this, we performed a systematic analysis of sequence conservation and predicted disorder across the *S. cerevisiae* proteome (see *Methods*) (*27*, *28*). This analysis revealed pervasive disorder and confirmed an expected anti-correlation between conservation and disorder (**Fig. 1B**, **fig. S3-6**). Interestingly, essential proteins in *S. cerevisiae* are – on average – as disordered as non-essential proteins (**Fig. 1C**) (*29*). We, therefore, wondered if seemingly poorly conserved IDRs from essential proteins could be functionally conserved despite large variations in sequence.

**Fig 1:**
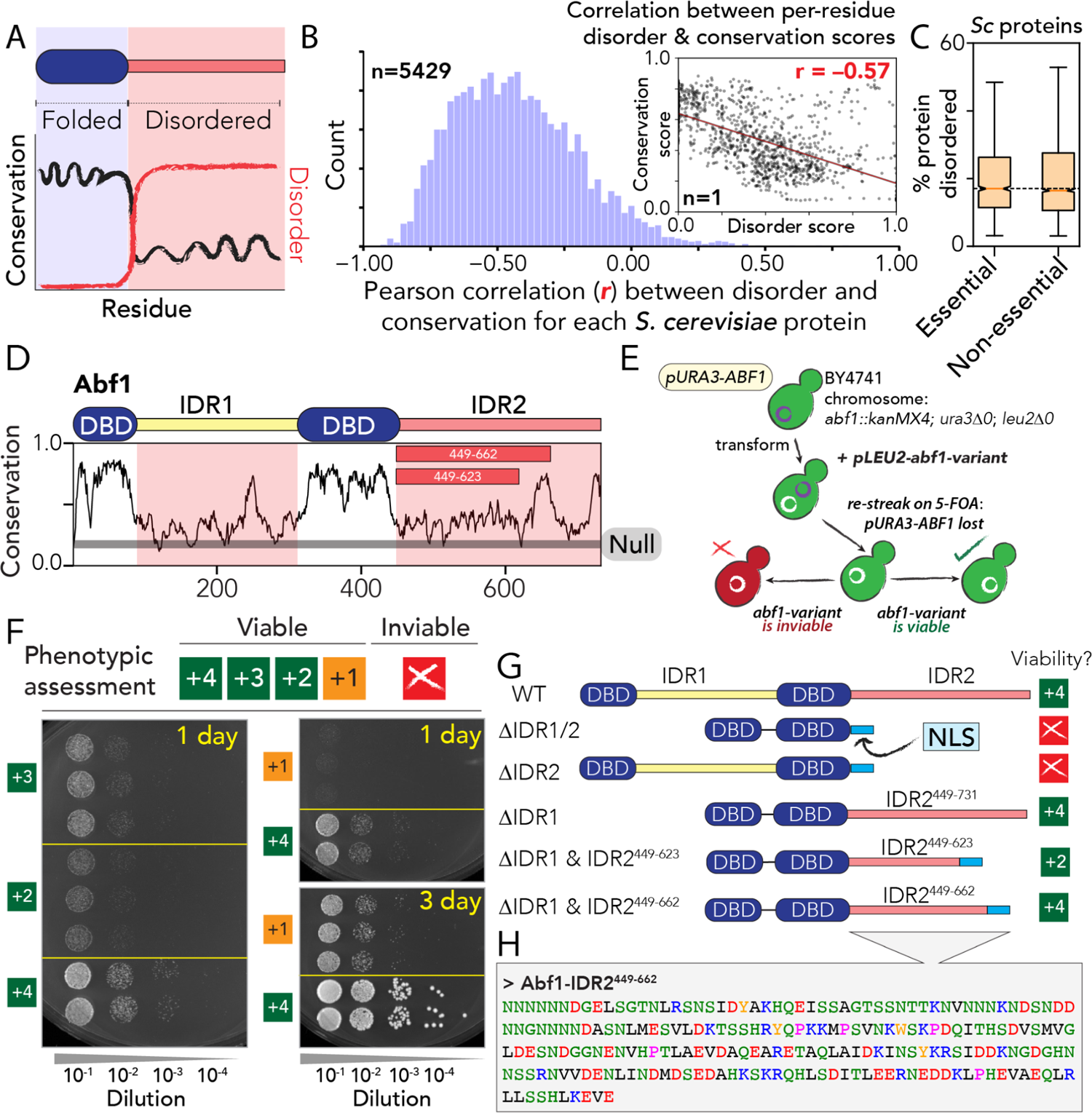
IDRs are poorly conserved as assessed by linear alignment, and Abf1 is an essential protein with a poorly conserved yet essential IDR. (A) Schematic showing conservation and disorder across a hypothetical protein. (B) Histogram of per-protein correlations (*r*) between per-residue conservation and disorder scores. Inset shows an example of a single protein, with each marker representing one amino acid in this protein. The histogram reports on *r* values for the entire yeast proteome. (C) Percentage of the sequence defined as IDRs for essential *vs.* non-essential *S. cerevisiae* proteins. (D) Sequence analysis of Abf1 with per-residue conservation is shown. The horizontal grey line corresponds to the conservation score expected for a randomly shuffled sequence (Null). (E) Schematic of plasmid shuffle assay. Variants measured are expressed from a plasmid. (F) Spot dilution assays for semi-quantitative assessment of sequence-dependent growth rate, scoring each viable construct between +4 and +1 (fig. S2). (G) Domain diagram for Abf1 truncation mutants with their viability shown in the right-hand side column. NLS: SV40 nuclear localization signal (H) Abf1-IDR2^449-662^ amino acid sequence, the focus of this study.

### Abf1 is an essential chromatin-binding protein with an essential IDR

We sought to identify a model protein to explore the conservation of function in the absence of sequence conservation. Because long IDRs are abundant in the context of chromatin-associated proteins (**fig. S7**) (*30*, *31*) we turned to the class of yeast general regulatory factors (GRFs). GRFs consist of sequence-specific DNA binding domains (DBDs) and long IDRs (**fig. S8**) (*32*), are abundant, and are mostly essential for viability. They share features with mammalian “architectural factors” (e.g., CTCF) (*33*, *34*), but with binding sites mostly in promoter regions where they function as barriers/boundaries/insulators in organizing or regulating chromatin in various contexts (*35–43*). Although often subsumed under the category of “transcription factors” due to their strong effects on transcription, they are distinct from classical transcription factors in that they mostly do not harbor transactivation domains (*44*). In contrast to transcription factor IDRs, and despite their numerous genetic and physical interactions (*45*, *46*), little is known about how GRF IDRs function besides the general notion that they interact with a range of partners and modulate genomic processes by opening and partitioning chromatin (*35*).

Abf1 is an essential *S. cerevisiae* GRF (*47*) that modulates nucleosome positioning and, thereby, chromatin accessibility at promoters (*36*, *38*, *40*). Rather than directly driving transcription, it functions as an insulator and mediates unidirectional transcription from potentially bidirectional promoters (*35*, *48*, *49*), as well as participates in roadblock termination of RNA polymerase (*50–52*). For our purpose, Abf1 has an ideal prototypical domain architecture, with a bipartite DNA binding domain and two poorly conserved IDRs (IDR1 and IDR2) (**Fig. 1D**). While the DBDs mediate sequence-specific DNA recognition, the IDRs are considered malleable protein-protein interaction modules, based on extensive genetic interaction experiments, although direct physical interactions are still mostly ill-defined (*45*, *53–56*).

Our interest in Abf1 is buoyed by prior observations – reproduced by us (**fig. S1, S2**) – that the full-length *K. lactis* orthologous Abf1 can complement function in *S. cerevisiae* (*57*). This suggests that despite 150+ million years of evolutionary divergence, Abf1 likely performs similar functions across fungi. With this in mind, we can say that full-length Abf1 appears functionally conserved (i.e., distant orthologs are sufficient for function). However, whether disordered regions excised from proteins offer the same functional conservation is often assumed but rarely tested. Finally, we have strong reason to believe the IDRs in Abf1 function in the context of protein-protein interactions, in no small part because Abf1 has a large number of physical interactors identified through various experiments (28 physical interactors in BioGrid (*45*)) yet possesses only a bipartite DNA binding domain and two IDRs. The absence of canonical protein-protein interaction domains strongly implicates the IDRs as the main protein interaction modules.

We first established which Abf1 regions were essential for function. Previous studies on Abf1 focused on the DBD, although truncation experiments identified apparently essential C-terminal sequences (CS1/2) in IDR2 (*58*, *59*). We extended this truncation approach through a classical plasmid shuffling assay (*60*). In this assay, the chromosomal *ABF1* gene was deleted and a wild-type *ABF1* gene was provided on a plasmid with a *URA3* marker (**Fig. 1E**). Transformation with a plasmid bearing an *abf1* mutant gene and selection on 5-FOA plates for loss of the wild-type *ABF1* gene plasmid (5-FOA is toxic in the presence of *URA3*) assessed if the *abf1* mutant gene supported viability (see *Methods*, **fig. S1** and **Table S1-S8**).

As Abf1 is an essential protein, viability in this assay was a clear readout for the essential function of the respective Abf1 mutant protein. For all Abf1 constructs that supported viability, we compared growth rates in a semi-quantitative way, ranging from +4 (wildtype-like) to +1 (very slow growth), by spotting serial dilutions (**Fig. 1F, fig. S2**). Constructs that we class as inviable are unambiguously inviable. Importantly, for the mutant inviable constructs, co-expressed wildtype and Flag-tagged mutant to perform a ChIP assays (see *Methods*, **fig. S9**) to verify that these mutant proteins (i) do not confer gain-of-function toxicity, (ii) are expressed, (iii) enter the nucleus, and (iv) bind specifically to Abf1 DNA binding sites. This allowed us to ascribe inviable constructs in terms of losing essential IDR2 function, as opposed to spurious mislocalization, impaired DNA binding, or aberrant protein degradation.

Our molecular dissection revealed that IDR2 but not IDR1 is essential (**Fig. 1G**). The maximal C-terminal IDR2 truncation, IDR2^449-662^ (**Fig. 1H**) that was reported to be viable in the presence of IDR1 (*59*) was also viable at wild-type growth rate in the absence of IDR1 (**Fig. 1G**). We used IDR2^449-662^ as our “wildtype” IDR reference sequence and refer to it as IDR2 for the remainder of this work. IDR2 is poorly conserved across all orthologs (**Fig. 1D**), yet essential for viability, at least in *S. cerevisiae*. The previously identified CS1/2 region that corresponds to a small island of conservation (peak in **Fig. 1D** at residue ∼650) was previously shown to be a nuclear localization sequence (NLS) (*56*). We confirmed that this region was not strictly necessary if a heterologous SV40 NLS was included (**Fig. 1F**, bottom). We included the SV40 NLS in all *abf1* mutant constructs to avoid scoring effects on nuclear localization.

### Abf1-IDR2 is poorly conserved by alignment but modestly conserved by chemistry

Based on prior work, we anticipated that conservation in IDRs could be considered in terms of two distinct metrics: compositional conservation and linear sequence conservation (*5*, *16*, *18*, *61*). Using this framework, we can identify proteins that are well conserved in terms of linear sequence (and hence composition), well conserved in terms of composition alone, or poorly conserved by both (**Fig. 2A**). A comprehensive analysis of conservation across *S. cerevisiae* IDRs and folded domains confirmed that many IDRs, despite being poorly conserved in terms of linear sequence, are well-conserved in terms of composition (**Fig. 2B, fig. S10-S11)**. Importantly, this analysis revealed that IDR2 is more conserved in terms of charged and hydrophobic residues than most IDRs with a similar degree of sequence conservation (**Fig. 2B, fig. S10**). In contrast, polar residue composition appeared less well-conserved.

**Fig 2:**
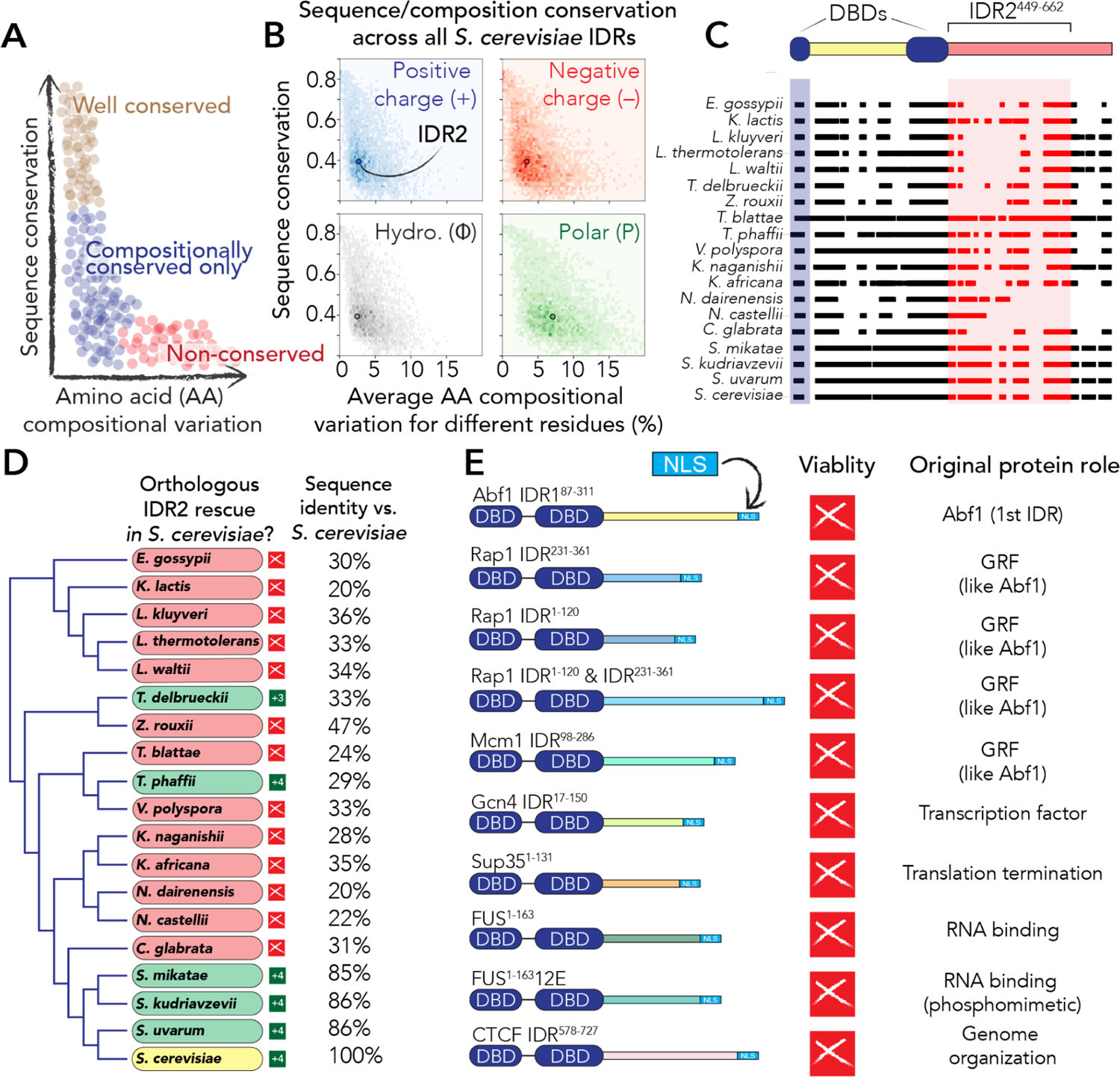
IDR2 shows relatively conserved composition, yet most orthologous sequences cannot rescue function. **(A)** Schematic of compositional and sequence conservation. **(B)** An analysis of all *S. cerevisiae* IDRs reveals Abf1-IDR2 (colored dot) is relatively conserved in charge and hydrophobicity composition but less so for polar composition. See also **fig. S10, S11**. **(C)** Schematic of orthologous IDR2s shown across full sequence alignment. **(D)** Phylogenetic tree (for the whole organism) vs. IDR2 rescue ability. Most IDR2 orthologs do not support viability in *S. cerevisiae*. Sequence identity vs. *S. cerevisiae* IDR2 is shown. See also fig. S12. **(E)** Alternative IDRs tested with their original function noted to the right. See also fig. S12.

### IDR2s from orthologous Abf1 proteins largely fail to rescue function in S. cerevisiae

We assumed that the conservation of composition seen for IDR2 would explain functional conservation, and anticipated that orthologous IDRs with similar compositions would support viability in *S. cerevisiae*. To test this, we took eighteen Abf1 orthologs, identified their IDRs corresponding to *S. cerevisiae* IDR2^449-662^ from the full-length proteins (**Fig. 2C**), and replaced the *S. cerevisiae* IDR2^449-662^ in our test plasmid with each of these orthologous IDR2s. We then tested the resulting chimeric constructs in our plasmid shuffle assay (**Fig. 2D**). Unexpectedly, outside of the *sensu stricto S. cerevisiae* complex (the bottom four species in **Fig. 2D**), only two of the fifteen orthologous IDR2s were viable (**Fig. 2D**), with no obvious relationship between sequence composition, sequence identity, or sequence length and function (**Fig. 2D, fig. S12**). To our surprise, our expectation that compositionally conserved IDRs would be functionally conserved proven incorrect.

We next wondered whether IDRs from proteins with similar functions might confer viability. We tested several candidates with similar IDR amino acid compositions, including IDRs from Abf1 (IDR1), other GRFs (Rap1, Mcm1), a yeast transactivator (Gcn4), and a human insulator (CTCF) (**Fig. 2E, fig. S12, S13**). We also tested unrelated but compositionally similar low-complexity IDRs from the human RNA binding protein FUS and the yeast translation termination factor Sup35 (**Fig. 2E, fig. S12, S13**). These IDRs failed to confer viability, suggesting that Abf1’s IDR2 provides specific molecular recognition.

Although most orthologous IDR2s could not rescue viability, we unexpectedly identified several IDRs from other *S. cerevisiae* proteins that conferred viability in place of IDR2 (**Fig. 3A**). These included compositionally similar IDRs from the yeast transactivators Gal4 and Pho4, and from the GRF Reb1 (**fig. S14**). Gal4, Pho4, and Reb1 are DNA-binding proteins that can trigger chromatin opening *in vivo* (*36–41*, *62–64*). These results illustrate that perhaps paradoxically, while many orthologous IDRs are inviable (**Fig. 2D**), there exist IDRs that differ substantially in length and linear sequence that confer wildtype-like growth.

**Fig 3:**
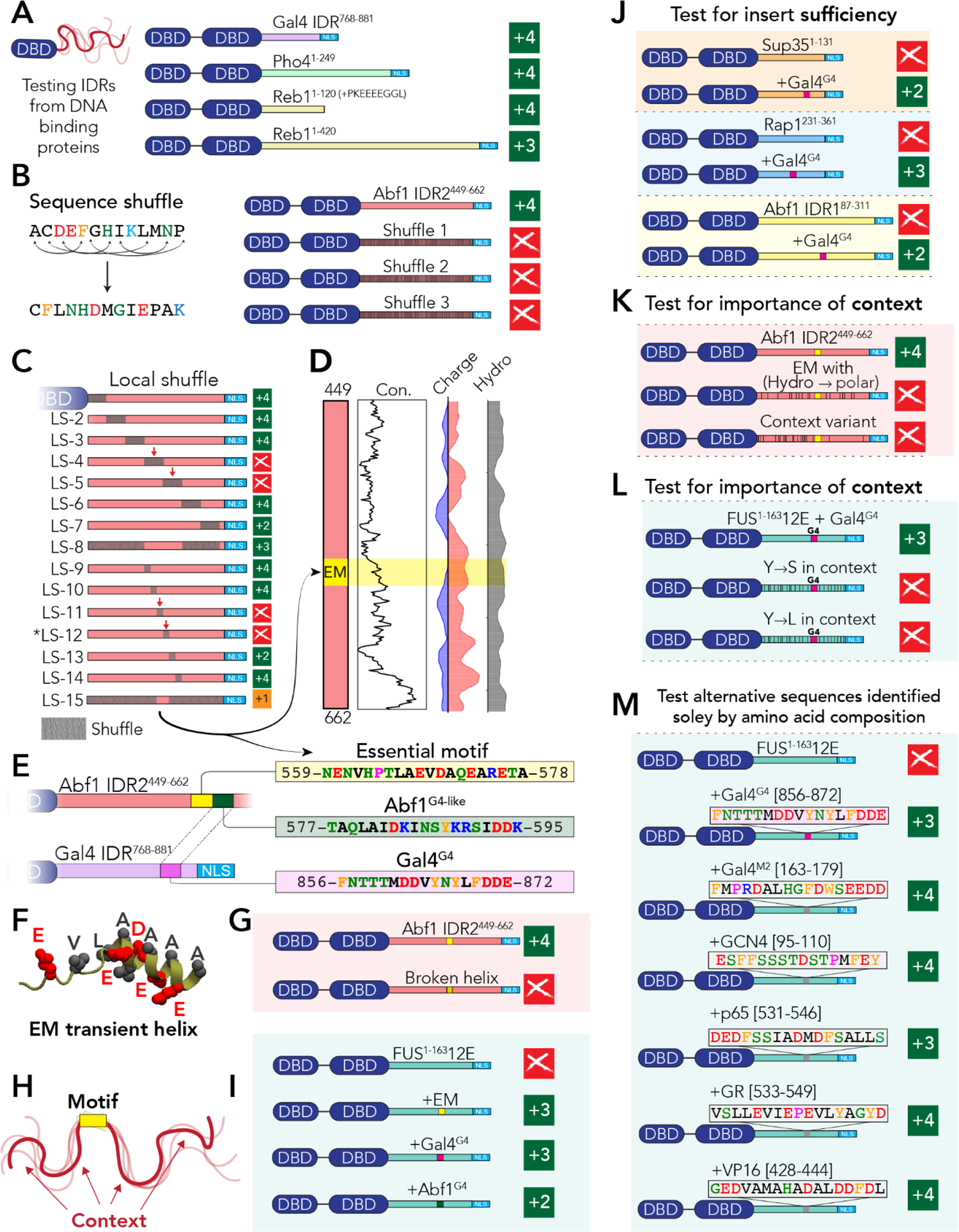
Sequence motifs play a key role in IDR2 function. **(A)** While most orthologs are inviable, several entirely unrelated IDRs can confer viability with wildtype-like growth. **(B)** Global shuffles are inviable, demonstrating that IDR2 composition alone is insufficient for viability. **(C)** Sequential sequence shuffling pin-points an essential motif (EM) in the center of IDR2. *NB* LS12 is also referred to as EM shuffle in Fig. 6. **(D)** The essential motif is not conserved across orthologs or enriched for charge or hydrophobicity compared to the rest of the sequence. **(E)** Three putative motifs are shown in the context of their IDRs. Abf1^G4-like^ and Gal4^G4^ were identified by sequence alignment of IDR2 and Gal4^768-881^ (**fig. S15**). **(F)** Structural bioinformatics implicates the essential motif in forming a transient helix (**fig. S16**). **(G)** Helix-disrupting mutations (glycine substitutions for other polar residues) abrogate viability. **(H)** Schematic showing motif and context. **(I)** Insertion of each of the three putative motifs into a non-functional IDR (FUS^1-163^12E) rescues that IDR to be functional. **(J)** Gal4^G4^ rescues viability in several unrelated non-functional IDR contexts. **(K)** Mutations to the IDR2 context that preserve the essential motif but reduce hydrophobicity in the context are non-viable. **(L)** Mutations to the FUS^1-163^12E context that preserve Gal4^G4^ but reduce hydrophobicity in the context are non-viable. **(M)** Compositionally matched subsequences taken from a range of transcription factors also provide viability inserted into FUS^1-163^12E.

### Amino acid composition in IDR2 is insufficient to explain function

Considering Gal4, Pho4, and Reb1 can mediate chromatin remodeling (one of Abf1’s functions), we wondered if a common <u>S</u>hort <u>Li</u>near <u>M</u>otif (SLiM) for recruiting the requisite machinery may be shared across these IDRs. Given SLiMs depend on their specific linear sequence (*8*), we reasoned that shuffling a SLiM would disrupt its function. As an initial test, we designed three globally shuffled variants of IDR2 with a subset of evenly distributed positions held fixed (**Fig. 3B**). All global shuffles were inviable, demonstrating that IDR2-like composition is insufficient for viability, implicating linear sequence-specific regions that must be essential. Taken together, our evolutionary comparisons and global shuffles suggest that wild-type Abf1 sequence composition is insufficient for function.

### IDR2 possesses an essential short linear motif

Our global shuffle variants imply IDR2 contains a SLiM. To identify this putative motif, we developed an unbiased approach termed sequential sequence shuffling (**Fig. 3C**). IDR2 was subdivided into non-overlapping 30-residue windows, and the sequence in each window was locally shuffled. This revealed two central windows that were intolerant to shuffling, which we confirmed by shuffling everything except the central 60-residue subsequence (**Fig. 3C**). We then repeated the procedure using 10-residue windows within the 60-residue subsequence. We identified a 20-residue subsequence (the essential motif, EM) that could not be shuffled and was essential for IDR2 function (**Fig. 3C**). This region is unremarkable with respect to other sequence properties and not conserved across orthologs (**Fig. 3D**), yet is unambiguously essential for IDR2 function.

Might similar motifs exist in the other functional IDRs identified in **Fig. 3A**? A global sequence alignment between IDR2 and Gal4^768-881^ was relatively poor, with only one sub-region showing alignment (**fig. S15**). Despite its low identity, this alignment revealed two remotely homologous regions, which we named Abf1^G4^ and Gal4^G4^ (**Fig. 3E**). Intriguingly, while viable, the shuffle construct in which this Gal4^G4^ region is shuffled does show a noticeable growth defect (**Fig. 3C**, LS13). We therefore sought to understand how these motifs may work and if they are sufficient for viability.

What might underlie the molecular basis for EM function? Structural bioinformatics predicts this region to form a transient helix (**Fig. 3F**), a feature frequently associated with IDR-mediated interactions (**fig. S16**) (*22*, *65*). To establish whether helicity may play a key role, we made a small number of helix-disrupting point mutations within the EM, abrogating viability (**Fig. 3G**), confirming the importance of this region for function and implicating the ability to form a helix as a possible molecular determinant of binding.

### Motif context is a critical determinant of viability

Given modular SLiMs should confer function when inserted into a neutral context, we next tested if the essential motif met this definition. Here, ‘context’ refers to the IDR sequence surrounding the SLiM (**Fig. 3H**). As our neutral context, we selected the phosphomimetic variant of the low-complexity IDR from the human RNA binding protein FUS (FUS^1-163^12E) (*66*) (**fig. S17**). FUS^1-163^12E is a compositionally uniform low-complexity disordered region that lacks secondary structure or known binding motifs (*66*, *67*). However, FUS^1-163^12E contains uniformly spaced hydrophobic (aromatic) and acidic residues, making it an ideal neutral IDR with similar sequence properties to IDR2. While FUS^1-163^12E alone was inviable (**Fig. 2E**), insertion of the essential motif into the FUS^1-163^12E context “rescued” the FUS^1-163^12E sequence to make it viable (**Fig. 3I**). Importantly, we obtained similar rescue phenotypes when inserting Gal4^G4^ and Abf1^G4^ into the FUS^1-163^12E context (**Fig. 3I**). Taken together, we interpreted these results to mean we had identified three distinct motifs that mediated essential interaction with one or more Abf1 partners.

Next, we tested whether our motif-insertion experiments (**Fig. 3I**) depended on the specific sequence context of the FUS^1-163^12E mutant. We inserted the Gal4^G4^ in three more otherwise inviable IDR contexts (Sup35^1-131^, Rap1^231-361^, Abf1-IDR1^87-311^) (**Fig. 3J**). In these three cases, the 17-residue Gal4^G4^ conferred viability, confirming the general ability of Gal4^G4^ to rescue function. Curiously, we noticed that while all three constructs were viable, we observed distinct patterns in growth rate; Abf1-IDR1 context and Sup35 context both conferred slower growth than the Rap1 and FUS^1-163^12E contexts. This observation made us curious: could we modulate viability without altering the motif but simply by altering the context?

To test the importance of context, we revisited our bioinformatic analysis (**Fig. 2B**), which identified the conservation of charge and hydrophobicity. We created two mutants of IDR2 that preserved the essential motif but altered the composition of the context, reducing hydrophobicity in two different ways. In both cases, these variants were inviable (**Fig. 3K**). This starkly contrasts with large-scale shuffle mutants that repositioned almost every residue in the sequence yet preserved composition and were viable (**Fig. 3C**). This demonstrates that sequence context can play a critical role in determining IDR2 function.

Next, we designed variants of FUS^1-163^12E + Gal4^G4^ where the contextual aromatic residues were converted to serine or leucine (**Fig. 3L**). Aromatic residues are critical for FUS-dependent IDR interaction in other systems, where they function via distributed multivalent interactions (*17*, *67*). Despite the presence of the Gal4^G4^, these variants were inviable (**Fig. 3L**). Furthermore, if Gal4^G4^ was inserted into a glutamine-rich IDR from the yeast transcriptional co-repressor Ssn6, this variant was also inviable (**fig. S1**). These results further illustrate how IDR context can also be essential to determine function.

To confirm that Abf1^G4^ and Gal4^G4^ contained *bona fide* SLiMs, we reasoned that an essential control experiment would be to take unrelated but length-matched subsequences with Abf1^G4^/Gal4^G4^-like amino acid compositions and demonstrate that these were inviable. We identified five subsequences compositionally similar to Gal4^G4^ from yeast and non-yeast transcription factor IDRs (**fig. S18**). If IDR2 function is SLiM-dependent, then these non-alignable subsequences from other species should be inviable, given that - other than composition - they were randomly selected. To our surprise, all six of these 17-residue subsequences were viable in the FUS^1-163^12E context, giving mostly wildtype-like or near-wildtype-like growth (**Fig. 3M)**.

### IDR-mediated interactions are driven by sequence- and chemical-specificity

This result prompted us to step back and reconsider our data. Conventionally speaking, the ability to insert a short (<20 residue) sequence into a non-functional IDR context and confer function is interpreted as a simple and unambiguous demonstration of a *bona fide* modular motif (*e.g.,* a SLiM). Given that sequence-specific motifs are often defined by three characteristics (an inability to tolerate shuffling, sensitivity to point mutations, and autonomous modular activity), the essential motif is an actual motif (**Fig. 3C**,**G****,I**). However, the finding that a collection of compositionally similar but unrelated subsequences were also functional implied we *either* had a remarkable ability to identify *de novo* motifs *or* that something more general was at play.

Our results thus far identified two determinants of function: (i) the presence of a motif (Fig. **3G,I****,J**), and (ii) the presence of a sequence context that we interpret to mediate distributed multivalent interactions (*16*, *17*, *68*) as hydrophobic residues were critical (**Fig. 3K,L**). These two binding modes can be considered in terms of sequence-specificity (the dependence on a precise amino acid sequence) and chemical specificity (the dependence on sequence-encoded complementary chemistry without the requirement for an exact amino acid order). Generally, these two modes of interaction are discussed separately. Motifs are considered in specific stoichiometric interactions, while distributed multivalent binding is mainly associated with biomolecular condensates (*9*, *16*, *18*, *22*). However, our results prompted us to wonder if these two interaction modes might instead exist on a combined two-dimensional landscape (**Fig. 4A)**.

**Fig 4:**
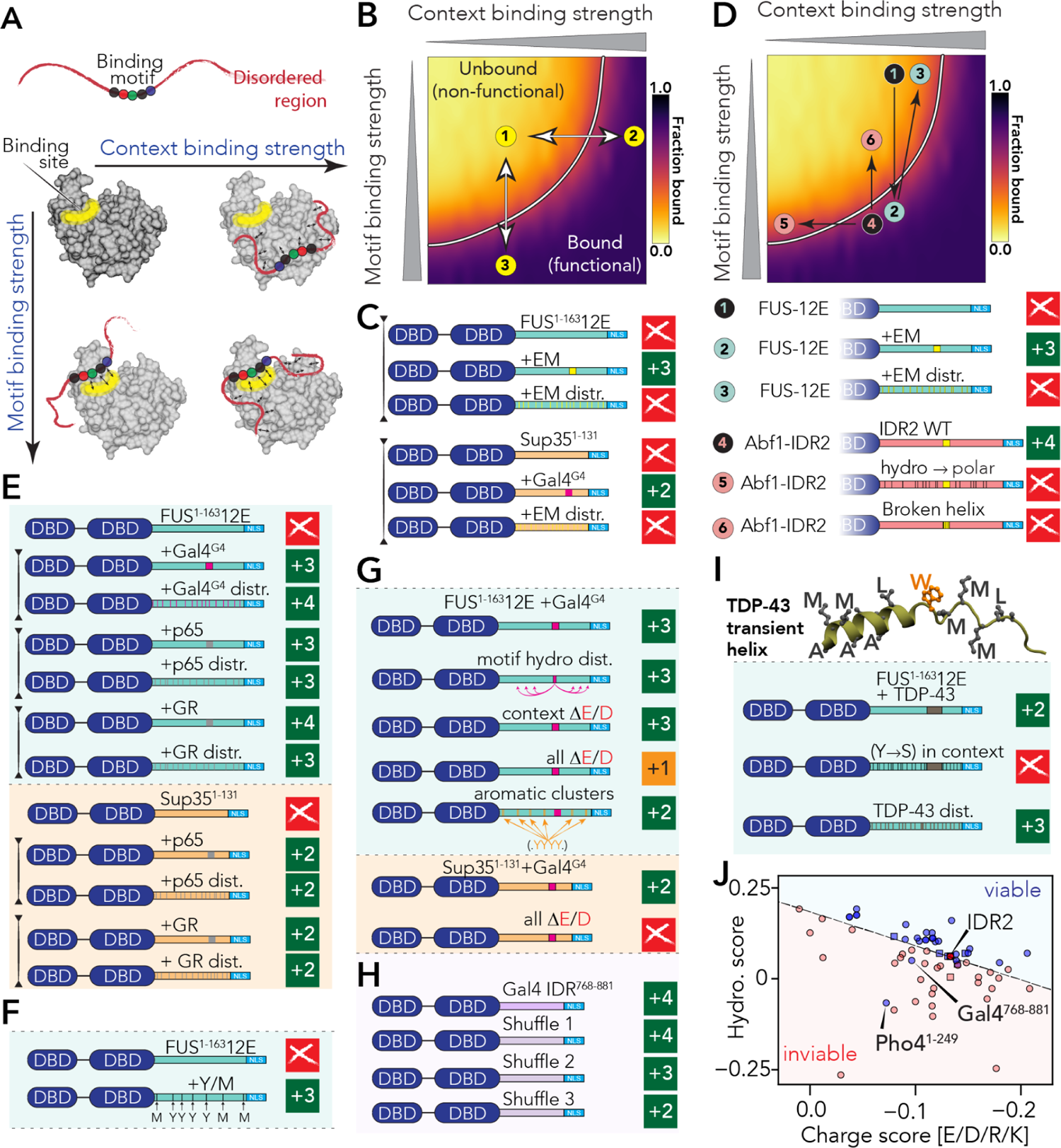
IDR2-mediated interactions can be understood via a two-dimensional binding landscape. **(A)** IDR-mediated interactions can be understood in terms of motif binding and context binding. **(B)** Motif and context binding can be projected onto a simple two-dimensional binding surface. **(C)** The essential motif is a true motif, in that it cannot be distributed across the IDR sequence. **(D)** Previous designs can be interpreted through the two-dimensional binding landscape. **(E)** Variants with distributed motifs identified by composition are functional in both FUS^1-163^12E and Sup35 backgrounds, with minimal change in growth phenotype. **(F)** Rational design of a motif-free FUS^1-163^12E variant. **(G)** Sufficient acidity and aromatic patterning can tune growth phenotypes. **(H)** Global shuffles of the Gal4^768-881^ construct reveal a variety of functional consequences. **(I)** The TDP-43 helical region can impart function upon the FUS^1-163^12E background, yet this effect is not dependent on the presence of a local motif. **(J)** Viable and inviable sequences can be classified based on charge and binding scores and parameters based on the weighted sequence composition. Circles are sequences that lack an essential motif, while squares are sequences that have an essential motif.

To guide our intuition, we performed simple coarse-grained simulations using the PIMMS simulation engine to quantify 1:1 binding for an IDR with its partner as a function of motif and context strength (**Fig. 4B,D**, **fig. S19**). These simulations reveal a two-dimensional landscape whereby a bound state can be achieved via many combinations of motif and context-binding strengths. Sequence changes can alter the context (**Fig. 4B**, from 1→2), the motif (**Fig. 4B**, 1→3), or both. Accordingly, we sought to test this conceptual framework via further rational sequence design.

Motifs are - by definition - sequence-specific, i.e., they depend on their contiguous linear amino acid sequence. It should therefore not be possible to distribute the amino acids of a motif across the IDR context and maintain function. In keeping with this, a variant with the essential motif residues distributed across the FUS^1-163^12E context was inviable (as was the redistribution of the essential motif in the Sup35 background) (**Fig. 4C**). This variant can be interpreted as simultaneously disrupting the motif but also (modestly) enhancing the context through the hydrophobic residues that are redistributed (**Fig. 4D**, 1→2→3). In contrast, variants that disrupt the context chemistry (**Fig. 4D**, 4 →5) or the motif alone (**Fig. 4D**, 4→6) effectively walk along orthogonal axes on our binding landscape. This conceptual framework allows us to re-interpret our additional mutants in a new light.

### Identification of motifs requires motif shuffling or re-distribution as a control

Our binding landscape model raised an intriguing possibility: What if Gal^G4^ and the other sequences identified in **Fig. 3K** were not *bona fide* motifs but instead altered the IDR context, albeit very locally, without being an actual motif? To test this, we asked if variants where the amino acids of these sequences were distributed were viable. In all cases and over multiple distinct contexts, we discovered that these distribution variants were viable, confirming our hypothesis (**Fig. 4E**). Importantly, the growth phenotype for the redistributed “motif” was the same or similar to the original “motif” construct.

### Rational design reveals the complex topology of chemical specificity

The functional yet motif-free sequences identified in **Fig. 4E** prompted us to pursue a strategy of rational design based on chemical principles. Given removing hydrophobic residues from contexts abolished viability (**Fig. 3K,L**) and given Gal4^G4^ and the other motifs in **Fig. 4E** must function by modulating the context, we reasoned this likely occurs through an increased number of hydrophobic residues. Accordingly, we asked if a FUS^1-163^12E variant with additional evenly distributed hydrophobic residues (+4 tyrosine (aromatic), +3 methionine (aliphatic), as in Gal4^G4^) would be viable. Indeed, even though this design was wholly artificial and even though wild-type IDR2 requires a *bona fide* motif, this design was viable with near wildtype-like growth (**Fig. 4F**). This result illustrates how an IDR with an essential SLiM can be rescued using a rationally-designed IDR that is lacking a SLiM if instead compensatory chemical groups are presented.

We sought to further explore the chemical determinants of function through rational IDR design. Taking the FUS^1-163^12E+Gal4^G4^ construct, re-distributing hydrophobic residues from the Gal^G4^ across the context had no impact on growth (**Fig. 4G**). Our bioinformatic analysis had previously implicated acidic residues as also being conserved (**Fig. 2B**). To explore this further, we removed all acidic residues from the context in the FUS^1-163^12E+Gal4^G4^ sequence, which had no impact on growth compared to the FUS^1-163^12E+Gal4^G4^ (**Fig. 4G**). However, removing every acidic residue in the sequence (context and motif) yielded a viable but extremely slow-growing phenotype (**Fig. 4G**). Intriguingly, the Sup35+Gal4^G4^ construct showed slower growth at baseline than the FUS^1-163^12E+Gal4^G4^ construct, arguing that Sup35 provided a weaker context than FUS^1-163^12E. Indeed, in the Sup35 context, removing all acidic residues led to an inviable construct (**Fig. 4G**). These results further emphasize the complex interplay between different types of sequence chemistry.

Recent work has implicated the patterning of aromatic residues in tuning intermolecular interactions in various contexts (*16*, *69*, *70*). To investigate the impact of aromatic patterning, we designed a variant (FUS^1-163^12E aromatic clusters) in which aromatic residues were clustered together (**fig. S20**). This variant was also viable, suggesting evenly-spaced aromatic residues are not a prerequisite for function in IDR2 (**Fig. 4G**). This variant showed a reduction in growth compared to the FUS^1-163^12E+Gal4G4, demonstrating that subtle changes in chemical context can be tuned *in vivo* by altering the composition and patterning of IDR sequences.

We originally identified Gal4^768-881^ as showing wildtype-like growth (**Fig. 3A**). While the Gal4^G4^ sequence is not a *bona fide* motif, we wondered if a motif existed within Gal4^768-881^. To test this, we generated three global shuffles. While each of these was viable (refuting the idea that a motif exists), each shuffle showed a different growth phenotype, ranging from wildtype (+4) to relatively slow-growing (+2) (**Fig. 4H**). This result again demonstrates that local sequence chemistry (beyond just composition) can play a crucial role in dictating function.

Finally, given the essential motif appeared to function as a hydrophobic helix, we wondered if we could rationally design an IDR with a hydrophobic helix. Excising a hydrophobic helix-forming region from the C-terminal IDR of the RNA binding protein TDP-43, this 25-residue segment did confer viability when inserted into the FUS^1-163^12E background in a background-dependent manner (**Fig. 4I**). However, redistribution of the helix residues was also viable indicating the importance of performing motif-redistribution controls to establish whether an inserted subsequence is a *bona fide* motif or not.

In sum, we found we can score each of our variants based on a hydrophobicity score (enhanced by aliphatic and aromatic residues, reduced by proline) and a charge score (reflecting the net charge of the sequence) (**Fig. 4J**). In this empirical chemical space, viable and inviable constructs predominantly fall on either side of a dividing line, with Pho4^1-249^ a notable exception (**fig. S21**). To test whether Pho4^1-249^ possesses a *bona fide* motif, we tested a global shuffle of this sequence, which – satisfyingly – is non-viable (**fig. S1**). While this does not unambiguously establish that Pho4 possesses a motif, it implies a sequence-specific dependence for a function that is not captured by composition alone, justifying the outlying behavior of Pho4^1-249^ in **Fig. 4J**.

While IDR-mediated interactions have often been viewed through the lens of sequence-specific motif binding, here we uncovered numerous examples in which wildtype-like function – at least in the context of our assay – is preserved in the absence of a motif. Moreover, we have examples in which the essential motif is preserved, yet function is lost. Taken together, we interpret these results to mean chemically specific interactions are sufficient and necessary. In contrast, sequence-specific interactions are, in fact, essential only in a small window of chemical contexts. More generally, our results imply that the conservation of chemically specific interactions can occur despite massive changes in the exact amino acid sequence. We therefore sought to explore this idea more systematically.

### Conservation of chemical specificity is invisible to alignment-based methods

If IDR function can be preserved via chemically specific interactions, we wondered how the conservation of chemical specificity would appear in a multiple sequence alignment. To investigate this, we leveraged a recently developed computational approach to predict IDR-mediated chemically specific interactions from sequence (*71*). This approach allows us to take a disordered region and a partner protein and predict which regions and residues in the disordered region facilitate attractive and repulsive interactions (**Fig. 5A**). We reasoned we could “evolve” IDR2 under the constraints of interacting with a partner via chemically specific interactions and investigate the degree of sequence conservation that emerged.

**Fig 5:**
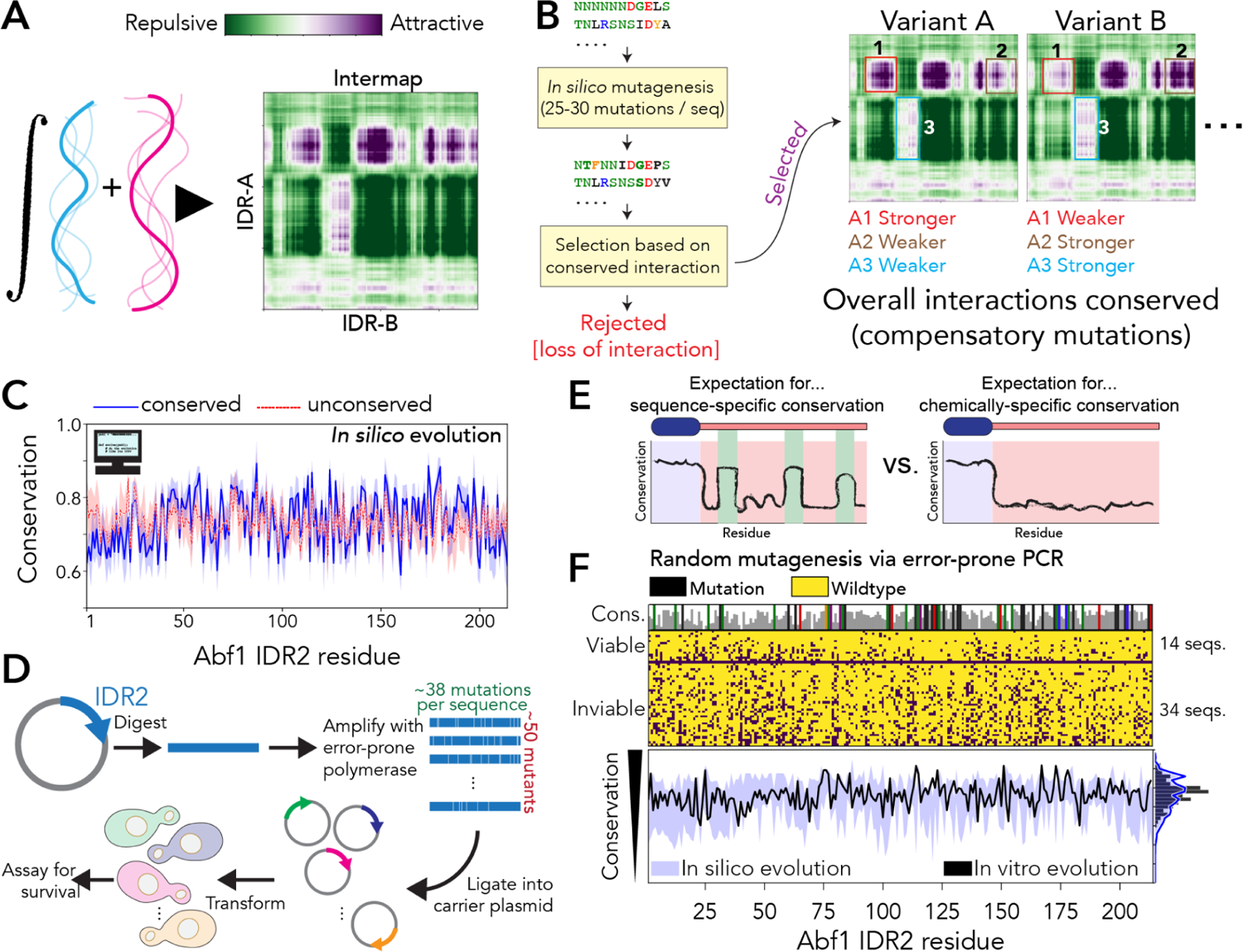
Chemical specificity imparts cryptic conservation that is challenging to identify from multiple sequence alignments. **(A)** Chemical specificity between two IDRs can be predicted directly from sequence using FINCHES. This computational approach repurposes physical chemistry developed for molecular simulations as a mean-field energy function to identify putative intermolecular attractive or repulsive interactions. (B) *In silico* evolution scheme. Mutations are generated by converting the protein sequence to nucleic acid and stochastically mutating nucleotides using standard transition/transversion rates and converting back to protein space. Selection is then performed by computing the predicted intermolecular interaction map (intermap) between the mutant IDR2 and the partner protein, requiring the overall conservation of attractive interactions. **(C)** Comparison of per-residue conservation for sets of orthologs evolved under selection (blue) vs. no selection (dashed red). These two profiles are indistinguishable from one another. **(D)** Schematic for error-prone PCR protocol used for *in vitro* evolution. (**E**) Expected conservation patterns for sequence-(*left*) vs. chemical (*right*) specificity. If sequence-specific conservation is in play, we anticipate seeing local peaks of conservation reflecting those regions that are highly conserved. **(F)** Results from *in vitro* and *in silico* evolution experiments. *Top*: *In vitro* evolution identified 14 viable and 34 inviable constructs. Viable constructs have mutations distributed throughout the sequence, including in the essential motif, without any conserved peaks. Residues that are never mutated in viable constructs are colored according to their amino acid chemistry (top bar plot). *Bottom:* Comparison of per-residue conservation from *in vitro* (black) and *in silico* (blue) evolution, where *in silico* experiments show the distribution that emerges when sets of fourteen viable mutants are generated many different times with the same number of average mutations.

While the *bona fide* IDR2 partners essential for function remain unknown (and are the source of ongoing investigation), Abf1 has previously been isolated as a stable trimeric complex with Rad7 and Rad16 (*54*, *72*). Moreover, the N-terminal IDR of Rad7 (Rad7^NTD^) possesses several distinct regions predicted to interact with IDR2 (**fig. S22**). We, therefore, performed *in silico* evolution of IDR2 under selection for chemically-specific interactions with the Rad7^NTD^ (**Fig. 5B**). However, we emphasize our conclusions here do not depend on the specific partner (**fig. S23**). As a control, we also evolved IDR2 without selection, in both cases ensuring the same number of average mutations was acquired for the two sets of “orthologs” (∼28 mutations). In the limit of small sets of orthologs (20-30 sequences), we discovered we cannot distinguish conserved vs. unconserved sets of artificial orthologs via alignment-based methods, *despite* the conserved variants being under strong selective pressure (**Fig. 5C**). Taken together, these results suggest multiple sequence alignments may be largely blind to conservation of chemical specificity. Next, we sought to test this model experimentally.

We next performed random mutagenesis to investigate the signatures of conservation in IDR2 (**Fig. 5D**). We generated a large numbers of length-matched randomly mutagenized versions of IDR2, tested them *in vivo*, and then analyzed the viable sequences to ask what features or regions were conserved. If specific residues/regions were highly conserved, this would strongly imply conservation of sequence specificity (**Fig. 5E**, top). Alternatively, if viable constructs appeared randomly mutagenized with no clear conservation pattern, this would be inconsistent with the conservation of sequence specificity and consistent with (although – we recognize – not unambiguously demonstrative of) the conservation of chemical specificity (**Fig. 5E** bottom). This approach yielded 48 variants with a distribution of point mutations, yielding 14 viable and 34 inviable constructs (**Fig. 5F**, top). Not only were no regions or residues statistically enriched for conservation as assessed by alignment, but a pool of mutationally-matched sequences evolved under conservation of chemical specificity was statistically indistinguishable from our data (**Fig. 5F** bottom). While caveats remain (see *Discussion*), our results demonstrate that IDRs can conserve chemical specificity despite appearing unconserved based on alignment-based analysis.

Finally, we wondered if the conservation of chemical specificity could be identified informatically across the yeast proteome. To assess this, we identified IDRs across the yeast proteome that were poorly conserved for multiple sequence-based conservation scores yet highly conserved to chemical specificity across various chemical interactions (see *Methods*). This analysis revealed ∼400 IDRs from proteins involved in essential cellular processes, including transcriptional regulation, DNA repair, and cell signaling (**Table S9**). In short, we conclude that the conservation of chemical specificity is widespread across the yeast proteome.

### IDR mutants lead to idiosyncratic changes in Afb1-responder sites

Our work thus far has focussed on Abf1 as a model system for examining sequence-to-function relationships in IDRs. However, we wondered what the molecular basis for Abf1-associated dysfunction in different IDR mutants might be. Abf1 has a well-described role in nucleosome organization as it participates in the generation of nucleosome-free regions (NFRs) over its binding sites flanked by arrays of regularly spaced nucleosomes (*41*, *73*, *74*). To investigate the functional consequences of a subset of Abf1 constructs, we asked how the replacement of wild-type Abf1 IDR2 with other IDRs affected nucleosome organization.

As Abf1 is essential for viability, wild-type Abf1 cannot simply be replaced by a non-viable variant. To circumvent this and avoid secondary adaptations to viable Abf1 variants, we turned to an *ad hoc* depletion approach with an Abf1 anchor away system (**Fig. 6A**) (*49*, *75*, *76*). FRB-tagged wild-type Abf1 is removed from the nucleus during 75 minutes of incubation in the presence of rapamycin, while either an untagged wild-type Abf1, no Abf1, or an Abf1 variant is retained in the nucleus. *Ad hoc* depletion of Abf1 recapitulates prior work; we observe partial filling up of the NFR and disorganization of the flanking nucleosomal arrays (shifts of peaks towards the NFR and decreased peak-to-trough ratios) at some (responder) but not other (non-responder) Abf1 sites (**fig. S24A,B**) (*76*). Moreover, if responder sites are in promoters, the corresponding genes are downregulated upon Abf1 ablation (*76*), an effect we also reproduced (**Fig. 6B**, no Abf1). These results give us confidence we have a working assay to interrogate Abf1 function *in vivo*.

**Fig. 6.**
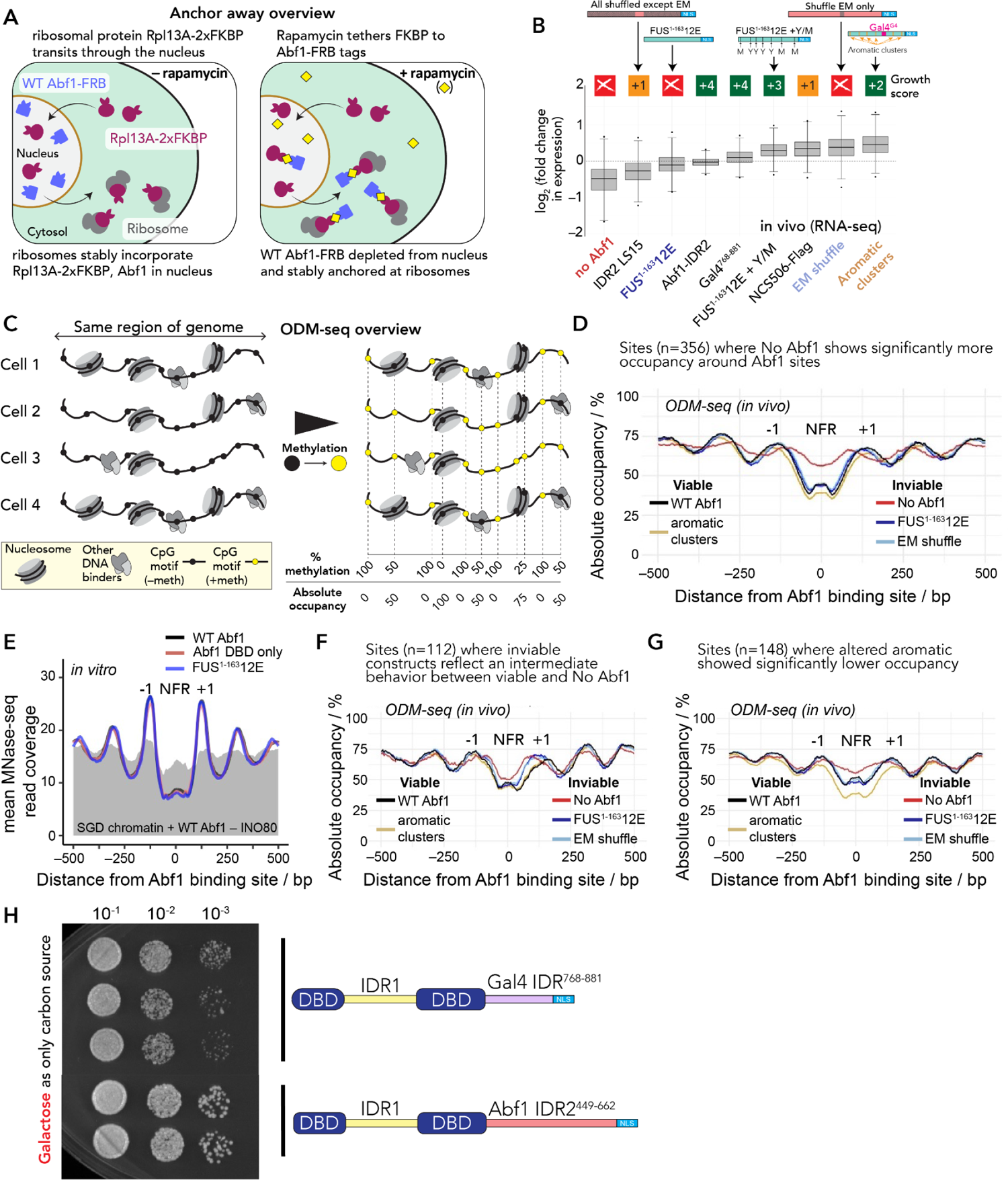
Variant IDRs have diverse effects on Abf1 function. **(A)** Anchor away enables *ad hoc* depletion of wild-type Abf1 from the nucleus. The wildtype copy of Abf1 is genetically fused to FRB while the ribosomal protein Rpl13A – which is constantly evicted from the nucleus during ribosome assembly – is fused to the double FKBP12 tag (2x FKBP). After adding rapamycin, FRB and FKBP bind, and nuclear Abf1 is sequestered into ribosomes, leading to *ad hoc* depletion. **(B)** Different Abf1 constructs enhance or suppress RNA levels for genes (n = 279) linked to Abf1 responder sites (fig. S24, see *Methods*). Change in total RNA levels in Abf1 anchor away strains with the indicated Abf1 constructs left in the nucleus relative to a strain with WT Abf1 left in the nucleus. n = 3 for WT Abf1, FUS^1-163^12E + Y/M, no Abf1, n = 2 for all other constructs. Note that EM shuffle was defined as Local Shuffle 12 (LS12) in **Fig. 3C**. NCS506 is a random mutagenesis construct (**Fig. 5**) with just 13 point mutations. **(C)** Schematic of ODM-seq chromatin mapping. (**D)** Examples for Abf1 sites (n = 356) aligned on positions where the no Abf1 construct showed significantly more occupancy in a 141 bp window centered on the Abf1 site. In these examples, variants had a minimal effect on chromatin structure. In the absence of Abf1, the nucleosome-free region (NFR) around Abf1 sites becomes occupied while the flanking nucleosomes (−1, +1) and up-/downstream nucleosome arrays become disordered, as seen by decreased peak-to-trough ratios and shifts of occupancy peaks towards the NFR. **(E)** *In vitro* genome-wide nucleosome positioning assay (MNase-seq) at 1025 Abf1 sites (ODM-seq responder sites for No Abf1 strain, **fig. S29**) reveals Abf1 IDRs are not required for robust nucleosome positioning. Lines compare different purified Abf1 variants in combination with ATP and the chromatin remodeler INO80. The grey shaded area reflects the same experiment in the absence of INO80. **(F)** Abf1 sites (n = 112) where both inviable Abf1 constructs (FUS^1-163^12E and EM shuffle) have similar effects on nucleosome organization, while the viable construct (Altered Aromatic Patterning) behaved like WT. **(G)** Examples for Abf1 sites (n = 148) where viable construct (Altered Aromatic Patterning) specifically affected nucleosome organization while both inviable constructs FUS^1-163^12E and EM shuffle did not. **(H)** The Gal4 transactivation domain in the Gal4 IDR^768-881^ construct allowed growth even under fully GAL-inducing conditions with galactose as the sole carbon source. Spot dilution assay after two days of incubation. All strains were incubated on the same plate.

A naïve expectation may be that the degree of downregulation of these responder genes, which were defined by effects on nucleosome organization (MNase-seq assay) in the absence of Abf1 (*76*), would reflect the degree of viability (growth rate score, **Fig. 1F**, **Table S7**) of Abf1 variants. However, this was not the case. Different IDRs fused to the Abf1 DBD led to idiosyncratic transcription responses (**Fig. 6B**). In hindsight, such varied outcomes may even be expected if the range of our tested IDRs mediated more or less specific interactions with various partner proteins that exert diverse effects depending on the Abf1 sites, i.e., where in the genome these IDRs are recruited via the Abf1 DBD. In sum, different IDRs need not necessarily lead to a “loss of function” in terms of an inability to recruit native Abf1 partners but also a “gain of function” in recruiting other nuclear components.

### Abf1 nucleosome barrier function does not depend on the IDR

While experimentally tractable, the transcriptional consequences of Abf1 variants are likely a secondary effect for the determinants of viability. We therefore next investigated if and how Abf1 variants influence chromatin organization at known Abf1 responder sites. Nucleosome occupancy measured by MNase-seq notoriously depends on limited MNase digestion degrees, making normalization difficult and introducing considerable technical noise on top of possible biological variation. Further, MNase-seq only scores the nucleosomal but not the non-nucleosomal fraction of the genome. In contrast, reliable absolute DNA occupancy in chromatin can be measured via next-generation DNA methylation footprinting and Nanopore sequencing (“ODM-seq” or “Fiber-seq,” **Fig. 6C** (*77*, *78*)). ODM-seq scores both occupied and non-occupied DNA (**Fig. 6C**), employs saturation of methylation (**fig. S25, S26, S38**), and yields low variation among replicates (**fig. S27**).

We generated such next-generation chromatin mapping data for the anchor away strains with WT Abf1, no Abf1, the inviable constructs (FUS^1-163^12E [**Fig. 3G**] and Local Shuffle 12 (LS12) [**Fig. 3C**], which we refer to as EM shuffle), or the viable constructs with compromised growth (aromatic clusters (**Fig. 4G**)) left in the nucleus. To identify Abf1 responder sites, we performed an unbiased search of all 2126 potential Abf1 binding sites across the genome, as predicted by position weight matrices (PWMs; (*79*, *80*)), and then identified locations in which a significant change in absolute occupancy around the Abf1 site or in the proximal/distal flanks of the neighboring nucleosomes was observed (**fig. S28,** *Methods*). Using the “No Abf1” construct to establish a baseline, this analysis identified many responder sites (**fig. S29**). We then investigated different subsets of Abf1-responsive sites to investigate the impact of IDR variation.

We first wondered if non-viable Abf1 variants would phenocopy the no Abf1 scenario. To our surprise, this was not the case in general. We identified many Abf1 sites where the lack of Abf1

– but not Abf1 variants – affected nucleosome organization (**Fig. 6D**, **fig. S30**). This implies that
– at least at some sites – the mere presence of a factor with the cognate DNA binding domain could serve as a nucleosome positioning barrier. To test this unexpected result directly, we performed *in vitro* genome-wide chromatin reconstitution assays with purified factors. Indeed, neither the lack of an IDR (ΔIDR1/2, i.e., Abf1 DBD only construct) nor an IDR that did not support viability in vivo (FUS^1-163^12E) compromised the generation of phased arrays with regularly spaced nucleosomes at Abf1 sites by an ATP-dependent chromatin remodeler like yeast INO80 (**Fig. 6E, fig. S36, S37**). These results establish the unexpected result that – at least for Abf1 – nucleosome barrier function does not depend on the IDRs.

### Abf1 nucleosome phasing function is strongly influenced by the IDR

Despite this result, our IDR variants did compromise nucleosome organization at Abf1 sites *in vivo*. In particular, both non-viable constructs (FUS^1-163^12E & EM shuffle) led to similar impairments with flanking nucleosome(s) moving towards the NFR, as seen in the absence of Abf1, at many Abf1 sites (**Fig. 6F, fig. S30**). The fact that the NFR was maintained nonetheless demonstrates that mere DNA binding at the Abf1 sites was sufficient to exclude nucleosomes *in vivo*, yet nucleosome phasing required the “right” IDR. This suggests that the essential function of Abf1 goes beyond just generating NFRs but is involved in organizing flanking nucleosomes, too. Quite surprisingly, the viable Abf1 variant (aromatic clusters) changed nucleosome organizations at some Abf1 sites in particular ways, e.g., even decreasing occupancy at the NFR and flanking nucleosomes, different from the lack of Abf1 and even where both inviable Abf1 constructs had not much of an effect (**Fig. 6G, fig. S30**). While such effects were compatible with viability, they may explain the reduced growth rate for this construct. Collectively, our ODM-seq chromatin data with Abf1 variant constructs strongly suggested a pivotal role for IDR2 of wild-type Abf1 in chromatin organization as it was impaired in more or less severe ways by alternative IDRs fused to Abf1 DBD.

### Abf1 can accommodate a strong and transcriptionally-active activation domain

Unlike canonical transcription factors, Abf1 lacks a transactivation domain (*44*) and participates in the insulation of transcription regulation (*48*, *81*). It was therefore unclear *a priori* if a GRF like Abf1 could tolerate bearing a strong transactivation domain. Our work here gives a clear answer, as we identified several viable Abf1 constructs with well-known transactivation domains, especially the Gal4 IDR^768-881^ construct with the notoriously strong Gal4 transactivation domain (**Fig. 3A**). Nonetheless, the Gal4 activation domain, including the short Gal4^G4^ activation domain sequence present in several of our constructs, may be repressed by Gal80 in our glucose-containing media (*82*). We confirmed that the Gal4 IDR^768-881^ construct still supported viability with galactose as the sole carbon source, meaning the GAL regulon was fully induced (**Fig. 6H**). Based on this result, we conclude that the essential GRF function of Abf1 is compatible with the presence of active and strong transactivation potential.

## DISCUSSION

IDR-mediated interactions have generally been viewed through the lens of either sequence-specific binding motifs (*e.g.,* SLiMs) or distributed multivalent interactions. These interaction modes are determined by sequence-specificity and chemical specificity, respectively, and elegant work from many groups established the functional importance of both (*15*, *19*, *22*, *83*, *84*). Here, we uncover the surprising result that – at least in Abf1 – these two modes can be compensatory for one another. A weak context could be compensated by introducing a motif (**Fig. 3**) and, more surprisingly, the absence of a motif could be compensated by the gain of context strength (**Fig. 4**). Importantly, we show that testing motif shuffling/distribution is essential to identify true motifs/SLiMs.

Our results can be rationally interpreted via a two-dimensional binding landscape (**Fig. 4B**). In this model, the determinants of function reflect how IDR2 engages in intermolecular interactions via some interoperable combination of motif-dependent and context-dependent binding. Based on our molecular understanding, we were able to rationally design new IDRs that were functional, although dramatically different and wholly unrelated to the wild-type sequence (**Fig. 4**). We note that biophysical hints for a model in which sequence- and chemical specificity co-operate to drive IDR-mediated binding have been found in numerous *in vitro* reconstituted systems (*3*, *19*, *20*, *85–87*). Finally, our work offers insight into plausible rules for evolutionary conservation in IDRs. While our *in silico* and *in vitro* evolution experiments cannot definitively prove one model over another, they are at least consistent with a model in which the conservation of chemical specificity shapes sequence variation in IDRs. Moreover, proteome-wide analyses find many examples in which this type of conservation suggests conserved intermolecular interactions despite large changes in amino acid sequence.

Over the last decade, there has been a growing appreciation for the intersection of sequence- and chemical specificity in the context of IDR function. The chemical determinants of IDR function have been explored in various contexts (*5*, *44*, *61*, *70*, *88–97*), yet how these properties influence the evolutionary landscape of disordered regions remains less clear. Our work suggests compensatory changes facilitated by chemical specificity may “buffer” the loss of SLiMs. Further, our *in silico* and *in vitro* evolution experiments imply that the conservation of chemical specificity may be almost impossible to infer from alignment-based analyses. This further motivates the need for novel routes to interpret conservation in disordered regions (*5*, *24*, *25*, *91*, *98–100*).

Recent work has invoked intracellular phase transitions to explain molecular principles underlying chromatin organization (*101–103*). While FUS^1-163^12E is highly soluble alone, it is plausible that it may phase-separate with an appropriate partner (*66*, *67*). We, therefore, wondered if our results could be interpreted through a phase transition model. Rational designs that either altered aromatic clustering (**Fig. 4G**) or substantially varied sequence length while maintaining a fixed composition (**fig. S1, S2, S31**) are viable. Given the strong dependence that phase behavior has on both aromatic residue clustering and the effective valence (determined in part by the sequence length), these results are inconsistent with a phase separation model underlying Abf1’s function (*16*, *70*, *104*, *105*) (**fig. S31)**. As such, while we cannot rule out the role of phase separation in this system, we find no strong evidence supporting it.

A key feature of our work is the use of rationally designed sequences to investigate the link between sequence and function. To help other groups investigate sequence and chemical specificity in their own systems, we have – in parallel – developed a set of computational methods for analyzing and designing IDRs with specific chemical and biophysical properties (*71*, *99*, *106*). Important for the generality of our work, our tools enable the prediction of chemical specificity directly from sequence, setting up clear and testable hypotheses in situations where an IDR’s partner is known. Emerging work from many groups has highlighted the importance of chemical specificity in various contexts (*19*, *23*, *107–112*). We propose that the systematic investigation of this phenomenon will require hypothesis-driven experiments facilitated by rationally designed IDRs with desired chemical or biophysical properties, approaches enabled by our methods.

Functions based on molecular interactions generally involve some degree of specificity, both towards a partner of interest as well as against off-target inhibitory interactions. Specificity is typically considered in terms of shape complementarity and chemical compatibility, two features afforded by folded domains and, to a lesser extent, SLiMs (*113*, *114*). Here, we find that even for an IDR that depends on a *bona fide* motif, alternative sequences with rationally interpretable chemical features can uphold function without any motif. We speculate that this chemical specificity offers an evolutionarily plastic mode of molecular recognition (**fig. S32**) (*18*, *19*, *61*).

The idiosyncratic functional consequences of Abf1 IDR variants on chromatin organization and gene expression likely reflect IDR-dependent changes in Abf1’s interactome. Therefore, an important feature absent from our current study is the interactome mapping for wildtype and Abf1 variants. Identifying the interaction partners is ongoing and will provide insight into the molecular basis for chemical specificity and Abf1 function. An additional layer of complexity reflects that IDR-mediated chemical specificity can alter genomic binding preferences for transcription factors (*110*, *115–118*). While this remains to be tested, we note that the Abf1 IDR2 has a relatively high fraction of negatively charged residues, yet charge-depleted variants are tolerated (**Fig. 4G**). This contrasts the well-studied yeast transcription factor Msn2, where negatively charged residues were essential for proper genomic targeting (*110*). This may indicate a different IDR grammar underlying GRF function versus transcription factor binding site selection.

Our ODM-seq chromatin mapping data for Abf1 variants *in vivo* demonstrated that the function of GRFs like Abf1 does not just entail nucleosome exclusion via binding to specific sites to generate NFRs. While NFR maintenance was grossly impaired upon Abf1 depletion (**Fig. 6D**, **fig. S24**, **S30**), as described (*76*), even inviable Abf1 constructs could maintain NFRs (**Fig. 6D,F,G**, **fig. S30**). Instead, the combination of NFRs and proper organization of nucleosomes – both at Abf1 sites – appears to constitute the function. This parallels recent findings in the context of replication (*119*) and 3D genome organization (*120*), where in both cases, NFRs alone were insufficient for function, yet the combination of NFRs and flanking phased arrays of regularly spaced nucleosomes was key.

In light of the increased evolutionary leeway offered by the interoperability of chemical and sequence-specificity, the limited functional conservation in IDR2 across orthologs despite the conservation of amino acid composition and sequence features may seem surprising (**Fig. 2, fig. S33**). To verify that full-length Abf1 performs analogous functions in other species, we confirmed that full-length Abf1 from *K. lactis* is viable in *S. cerevisiae* (**fig. S1**) (*57*). With this in mind, we emphasize that our study is focussed on IDR sub-regions orthologous to IDR2^449-662^, and not orthologous full-length proteins. Therefore, there are several (non-mutually exclusive) explanations for the inviability of orthologs. Firstly, interaction networks co-evolve by coupled evolutionary changes, such that SLiMs in orthologs may be incompatible with partners found in *S. cerevisiae*. Secondly, functionally important features can relocate across an IDR-containing protein. In this model, the specific location of a binding motif in the protein may be relatively unimportant, such that motifs can be lost from one region and emerge in another. An intriguing prediction from this model is that we should *expect* motifs to rapidly appear and disappear from a given IDR, a prediction supported and compatible with previous work on *ex nihilo* motif evolution (*121*).

While the essential motif appears poorly conserved, we still expect evolutionarily conserved SLiMs to be important and ubiquitous across proteomes. In line with this expectation, we identified thousands of short and conserved hydrophobic subsequences within IDRs, with almost twice as many conserved hydrophobic subsequences in essential proteins as non-essential ones (**fig. S34, S35, tables S10-S14**). We also identified many IDRs with low sequence conservation yet extremely high conservation of chemical specificity (**Table S9**). As a corollary, we wondered if other non-conserved regions of transient structure may be found in the yeast proteome and, upon analysis, identified 963 short (<40-residue) subregions in IDRs with predicted transient structure (**Table S17**). This set of *de novo* predictions includes the previously identified Pho4 activation domain (Pho4^69-94^) and four separate subregions in the N-terminal IDR of Reb1, both of which conferred viability (**Fig. 3A**). Moreover, a global shuffle of Pho4^1-249^ is non-viable, supporting the idea that there may exist a *bona fide* motif within Pho4 (**fig. S1**). While many such regions may be inert, others may offer specific binding interfaces, as is the case in Abf1, or specific helical regions identified in transactivation domains (*94*, *122*). In summary, this analysis offers specific, testable predictions for putative functional modules in IDRs at a proteome-wide scale, and we encourage others to use these predictions.

Finally, we focused on function measured by viability in *S. cerevisiae* growing under low-challenge laboratory conditions. This specific growth niche likely does not assess all facets of Abf1 function. The importance of other Abf1 regions and features in alternative growth conditions remains unassessed. How might alternative motifs or sequence features contribute to IDR function in Abf1? Ongoing work implies large-scale IDR-dependent remodeling of transcription during glucose starvation, and IDRs have been proposed to function as intrinsic sensors of intracellular state (*123*). If chemical specificity is defined by IDR chemistry, mechanisms to tune this chemistry (either via post-translational modifications, changes in protonation state, or changes in sidechain solvation properties) offer an attractive route for modulating specificity.

## Acknowledgments

We thank Rohit V. Pappu for his immediate willingness to collaborate once approached by P.K. and for allowing A.S.H. to work independently on this project while completing his postdoctoral work. We thank Rahul Das for his help in jump-starting this project and Benoit Kornmann for helping M.O.G.R. handle his SATAY data at an early stage of the project. We thank Wim de Jonge and Frank Holstege for sharing their Abf1 anchor away strain and their Abf1 site annotations, as well as numerous discussions and advice. We are grateful to Tobias Straub for advice on bioinformatics analyses and to Slawomir Kubik and David Shore for sharing their class I to V responder Abf1 site annotations. We thank Shahar Sukenik, David Moses, Broder Schmidt, and Stephen Plassmeyer for their critical comments and feedback on the manuscript. We thank Amy Keating and the members of the Keating lab for helpful discussion. We are grateful to Axel Imhof, Christoph Kurat, and Tamas Schauer for their input as thesis advisory committee members for I.L.-S. and especially Axel Imhof for his suggestion to use the ChIP assay as a control. We thank Stefan Krebs, Alexander Graf, and Helmut Blum (LaFuGa, Gene Center, LMU Munich) for Illumina and Oxford Nanopore sequencing and data handling and Mam Hataichanok (The Histone Source, Colorado State University) for purified recombinant histones. We acknowledge funding by the German Research Foundation (grants KO 2945/3-1 and within SFB1064) to P.K., and to ASH from the Longer Life Foundation (an RGA/Washington University Collaboration), the MOLSSI (via a Seed Fellowship), the Human Frontier Science Program (grant RGP0015/2022), and a CAREER award from the United States National Science Foundation (2338129).

## Data availability

Sequencing data are deposited at NCBI Gene Expression Omnibus (GEO; https://www.ncbi.nlm.nih.gov/geo/) under accession number GSE (to be released on publication). Source codes are deposited at https://github.com/gerland-group/ and https://github.com/holehouse-lab/supportingdata/tree/master/2024/Langstein-Skora_2024. GOOSE is available at https://github.com/idptools/goose/, and documentation for GOOSE is available at https://goose.readthedocs.io/. FINCHES is available at https://github.com/idptools/finches/.

## Author contributions

Conceptualization: PK, ASH

Methodology: PK, ILS, ASH, AS, RJE, MOGR, FH, DS, LS, FK, TB, ML, FJM, NP, VS, SKR

Investigation: PK, ILS, ASH, AS, RJE, MJG, SKP, MOGR, FH, DS, SKR

Visualization: ILS, PK, ASH, FH, DS, WA Funding acquisition: PK, ASH, UG, KPH

Project administration: PK, ASH, UG, KPH

Supervision: PK, ASH, UG, KPH, TB

Writing – original draft: ILS, PK, ASH

Writing – review & editing: ILS, PK, ASH, RJE

## Supplementary Materials

### Materials and Methods

Supplementary Text

Figs. S1 to S42

Tables S1 to S17 References (1–56)

