## Supplementary material for "Sequence- and chemical specificity define the functional landscape of intrinsically disordered regions": Fig S2

Unless noted otherwise, the following images show YPDA plates that were incubated for ca. 1 day at 30°C. If one of the independent clones for each construct did not show the same growth rate as the others, it is labeled as „outlier“ and we categorized the growth rate according to the two similarily growing clones. All rating is relative to the growth of the strain with wild type (WT) Abf1 on the same plate.

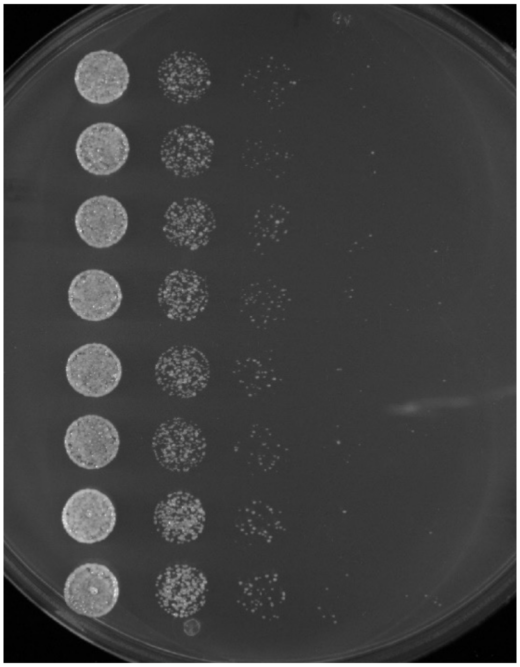

++++ FUS$^{1-163}$12E + GR

++++ FUS$^{1-163}$12E + Gal4 M2

++++ Abf1 WT

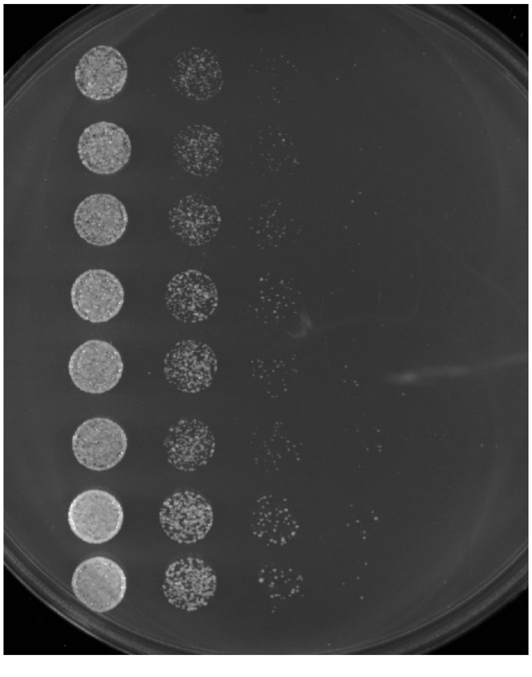

++++ VP16

++++ FUS$^{1-163}$12E + GCN4

++++ Abf1 WT

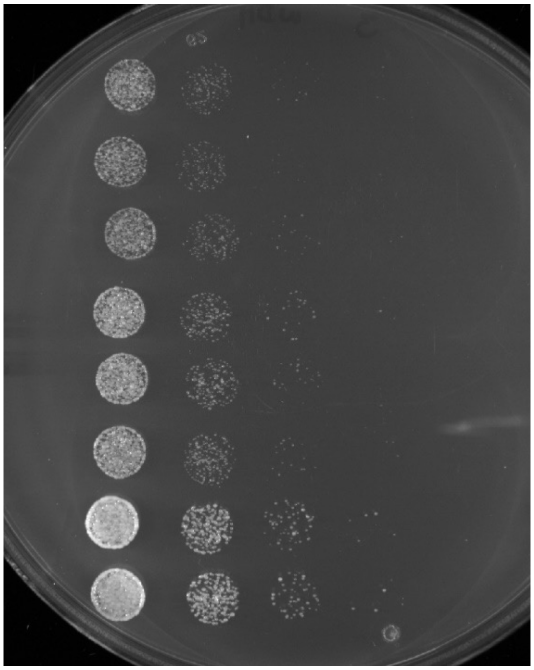

++ | Abf1 IDR1$^{87\text{-}311}$ + Gal4$^{G4}$ motif

+++ | Rap1$^{231\text{-}361}$ + Gal4$^{G4}$ motif

++++ | Abf1 WT

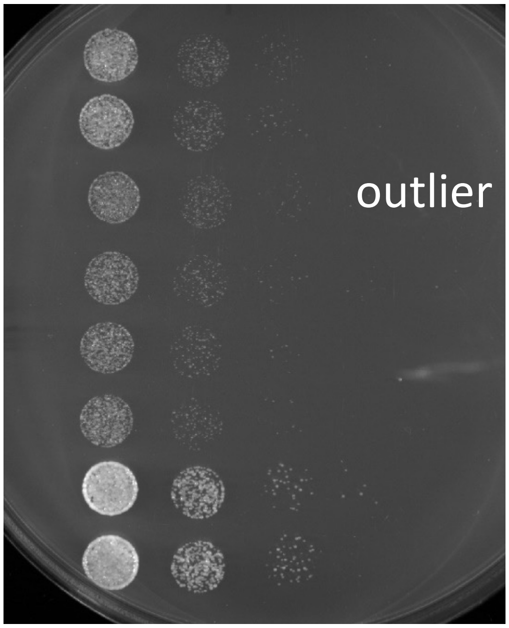


+++ | Local Shuffle 8

++ | Sup35$^{1\text{-}131}$ + Gal4$^{G4}$ motif

++++ | Abf1 WT

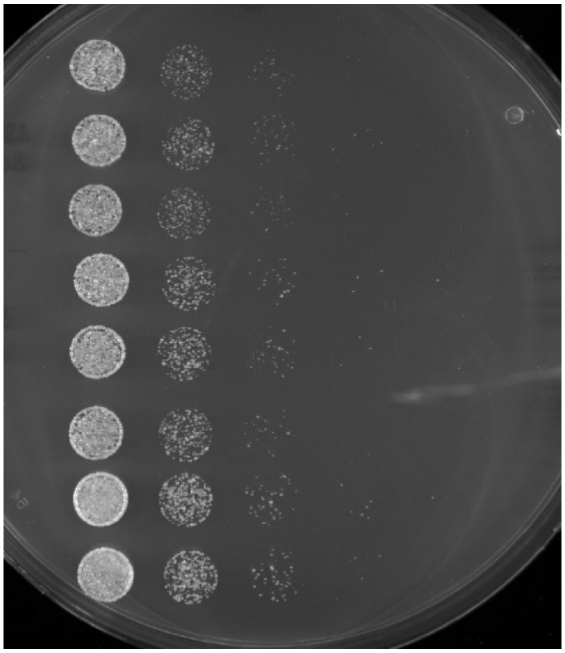

++++ FUS$^{1-163}$12E + Gal4$^{G4}$ distributed

++++ T. phaffii

++++ Abf1 WT

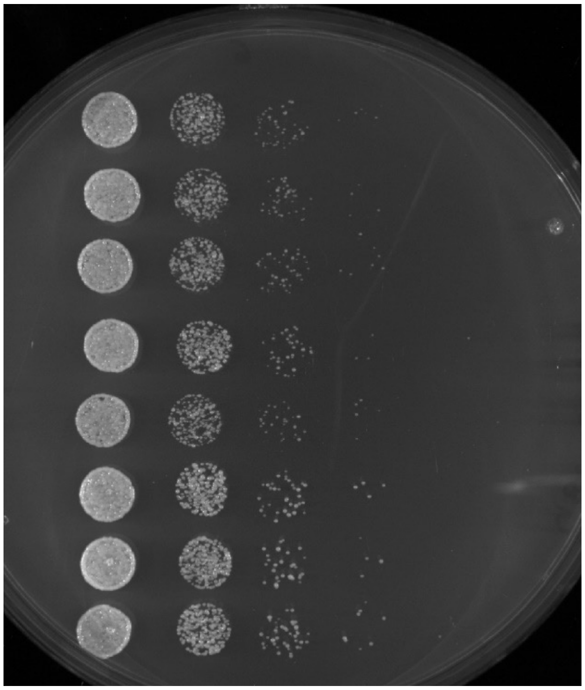

++++ Local Shuffle 2

++++ Local Shuffle 3

++++ Abf1 WT

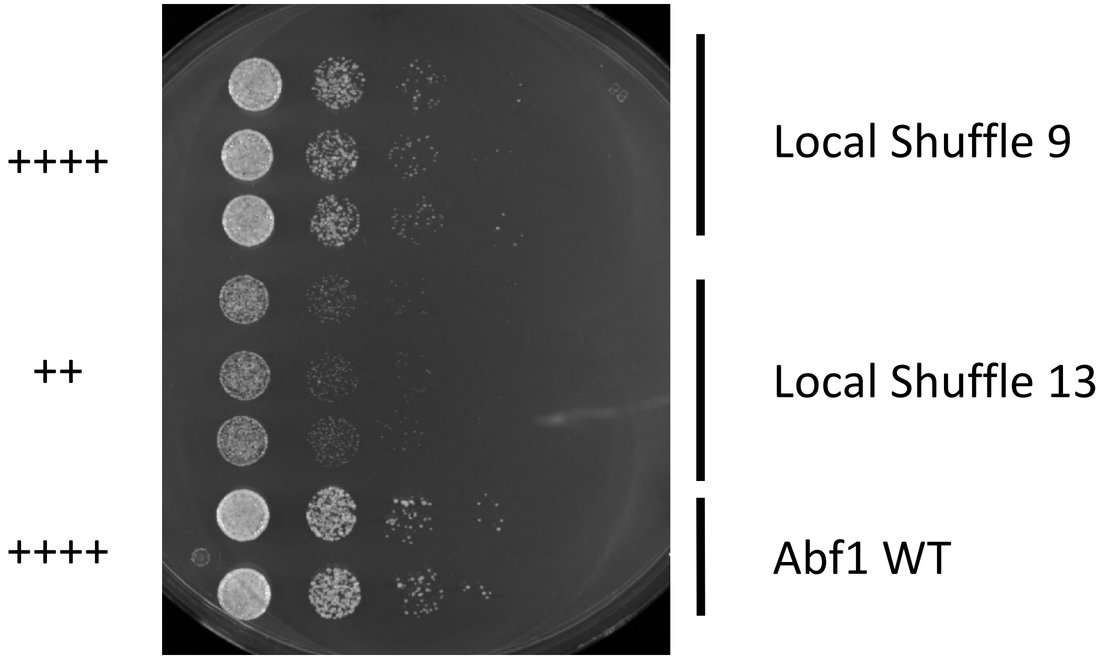
++++
++
++++
Local Shuffle 9
Local Shuffle 13
Abf1 WT

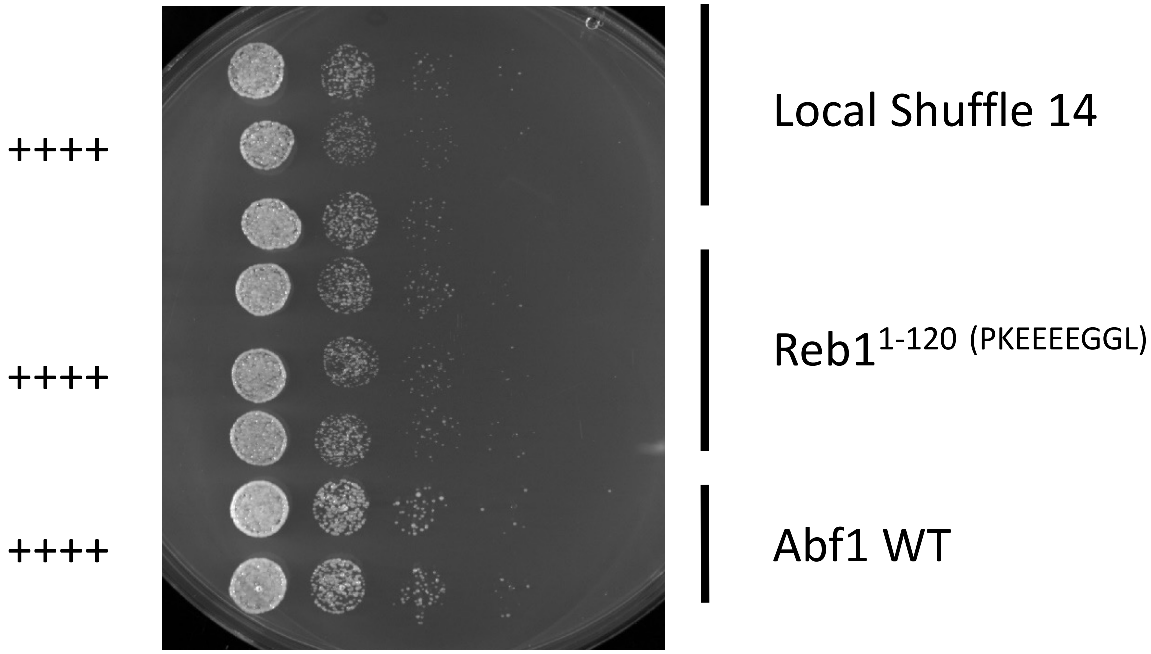
++++
++++
++++
Local Shuffle 14
Reb1$^{1-120\ (PKEEEEGGL)}$
Abf1 WT

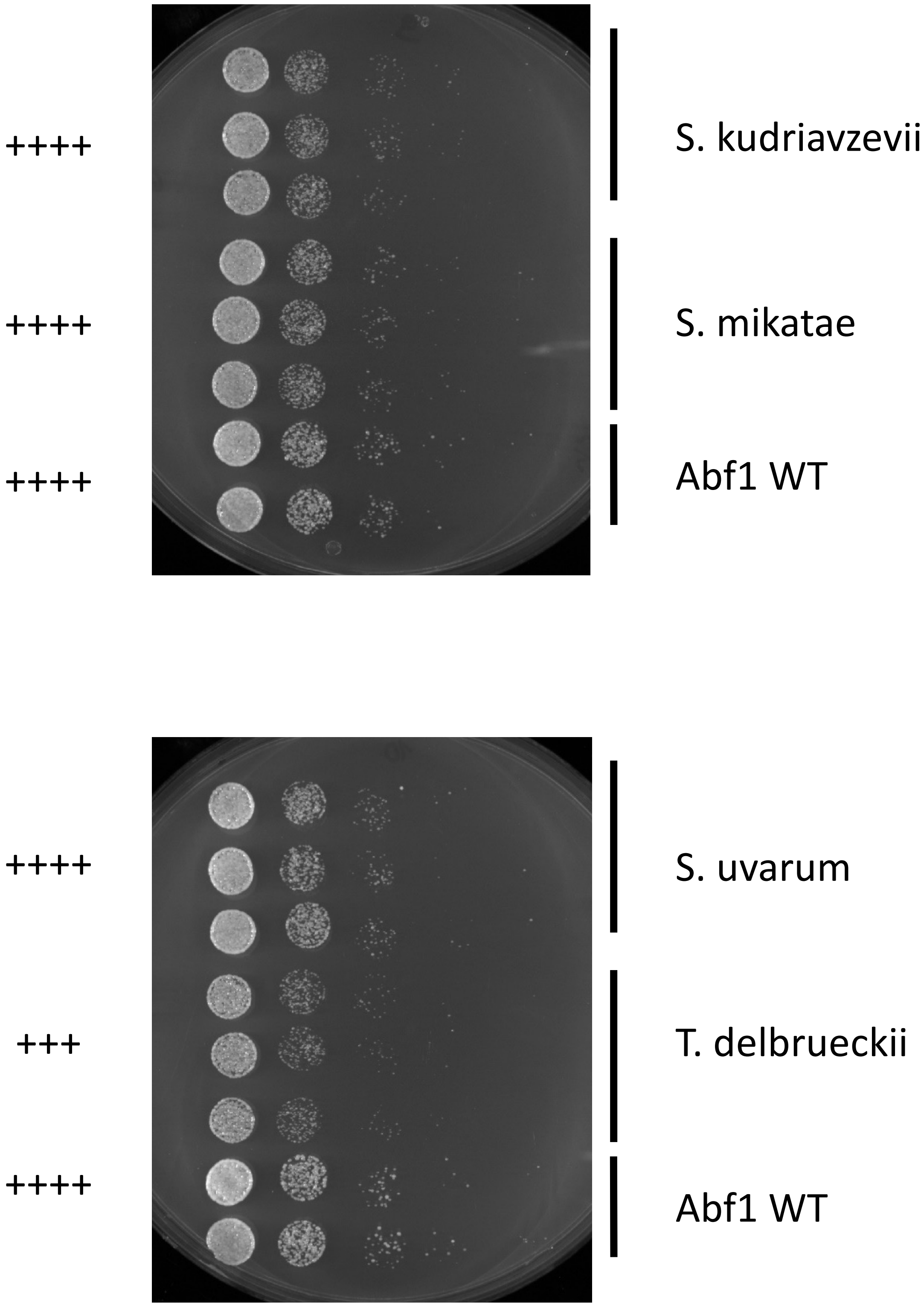

++++
S. kudriavzevii
++++
S. mikatae
++++
Abf1 WT
++++
S. uvarum
+++
T. delbrueckii
++++
Abf1 WT

+++

++

++++

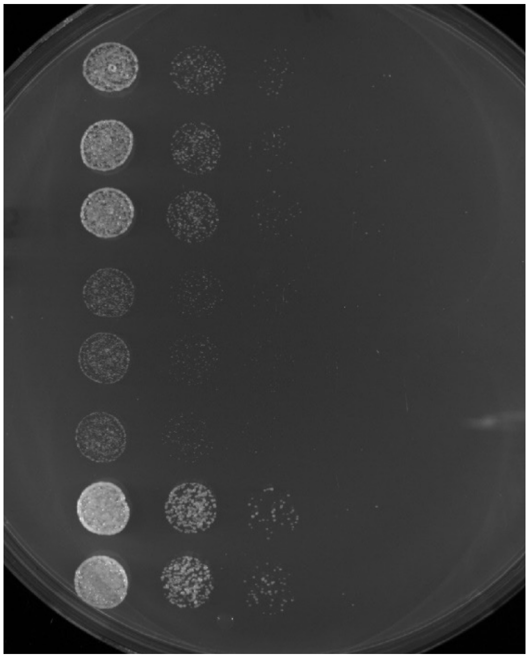

FUS12E + p65

FUS12E altered aromatic patterning

Abf1 WT

+++

++

++++

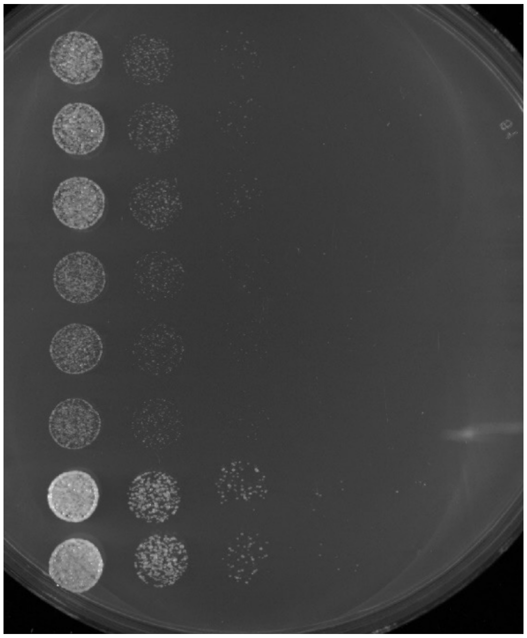

FUS12E + p65 distr.

Sup35 + p65

Abf1 WT

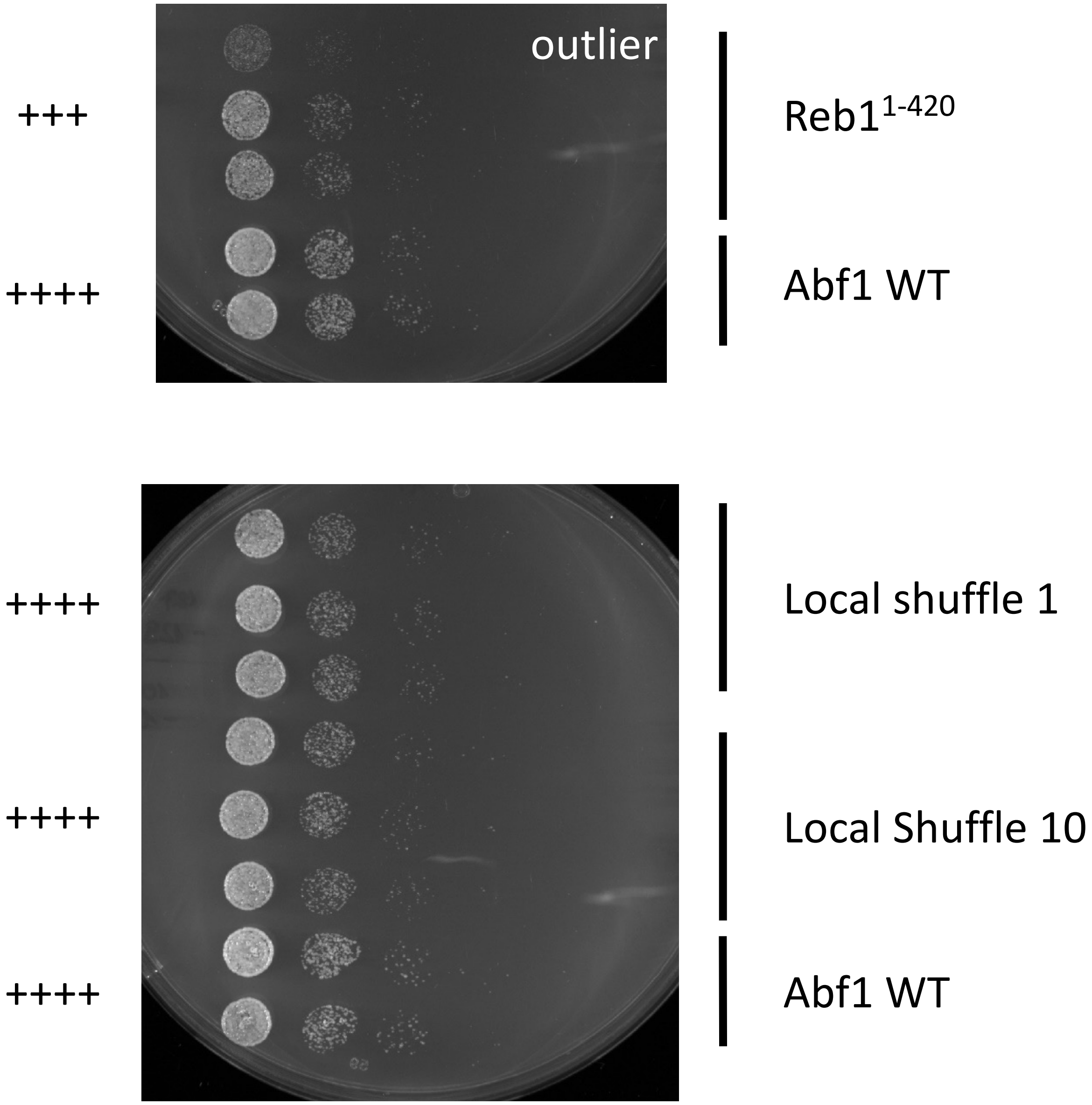
outlier
+++
Reb1^1-420
++++
Abf1 WT
++++
Local shuffle 1
++++
Local Shuffle 10
++++
Abf1 WT

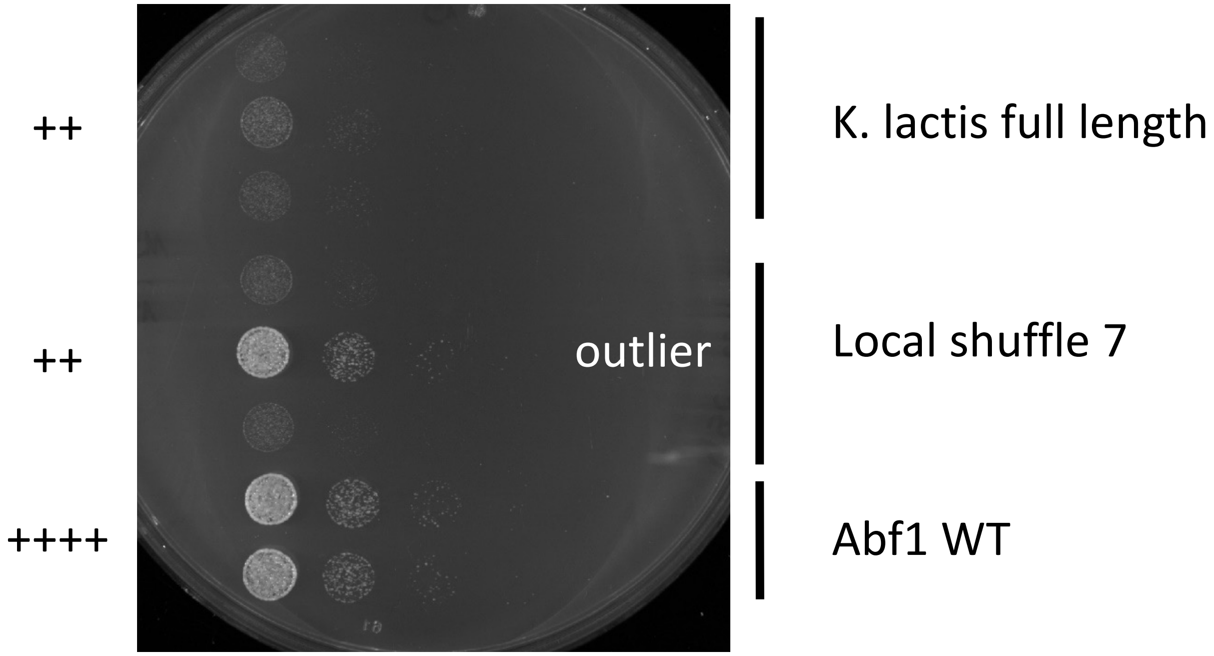
++
K. lactis full length
++
outlier
Local shuffle 7
++++
Abf1 WT

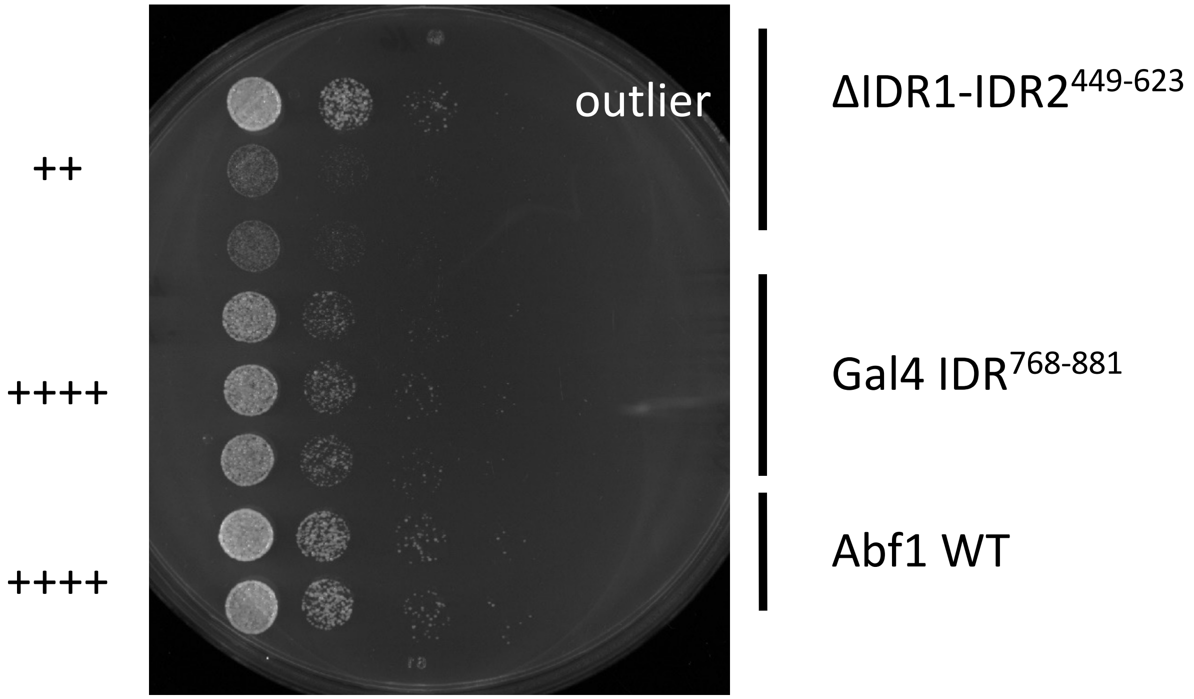
outlier
ΔIDR1-IDR2$^{449-623}$
++
++++
Gal4 IDR$^{768-881}$
++++
Abf1 WT

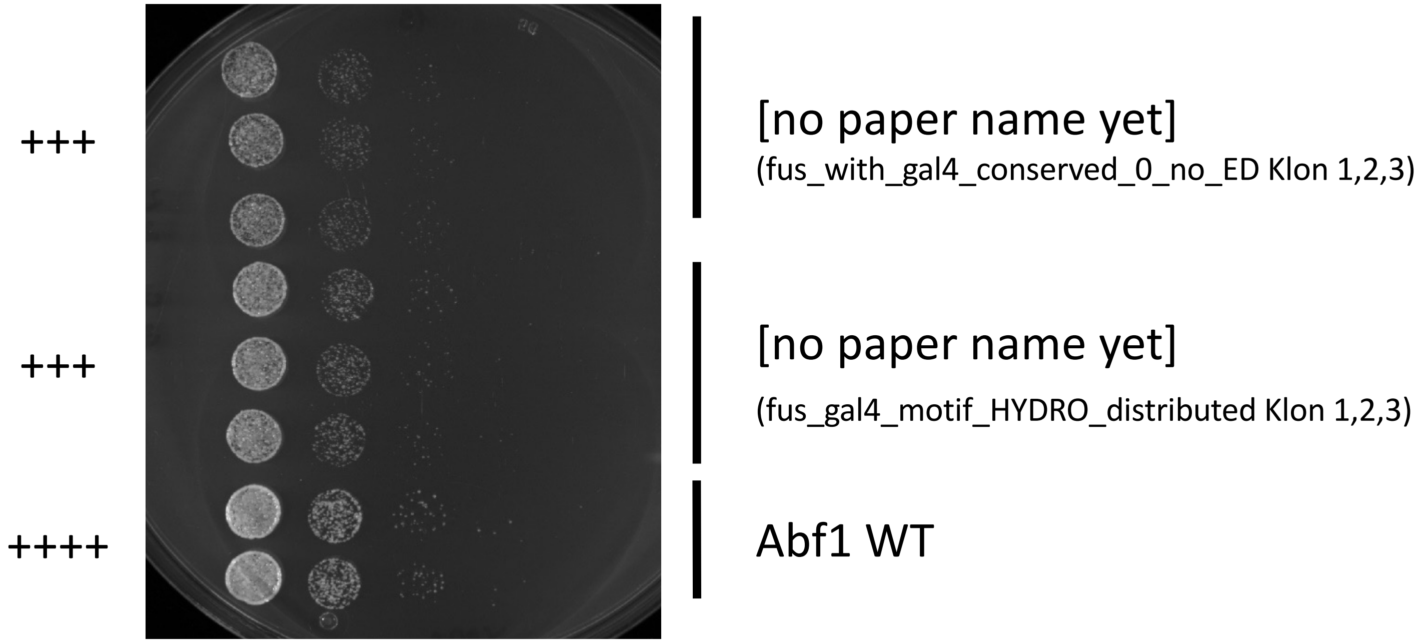
+++
+++
++++
[no paper name yet]
(fus_with_gal4_conserved_0_no_ED Klon 1,2,3)
[no paper name yet]
(fus_gal4_motif_HYDRO_distributed Klon 1,2,3)
Abf1 WT

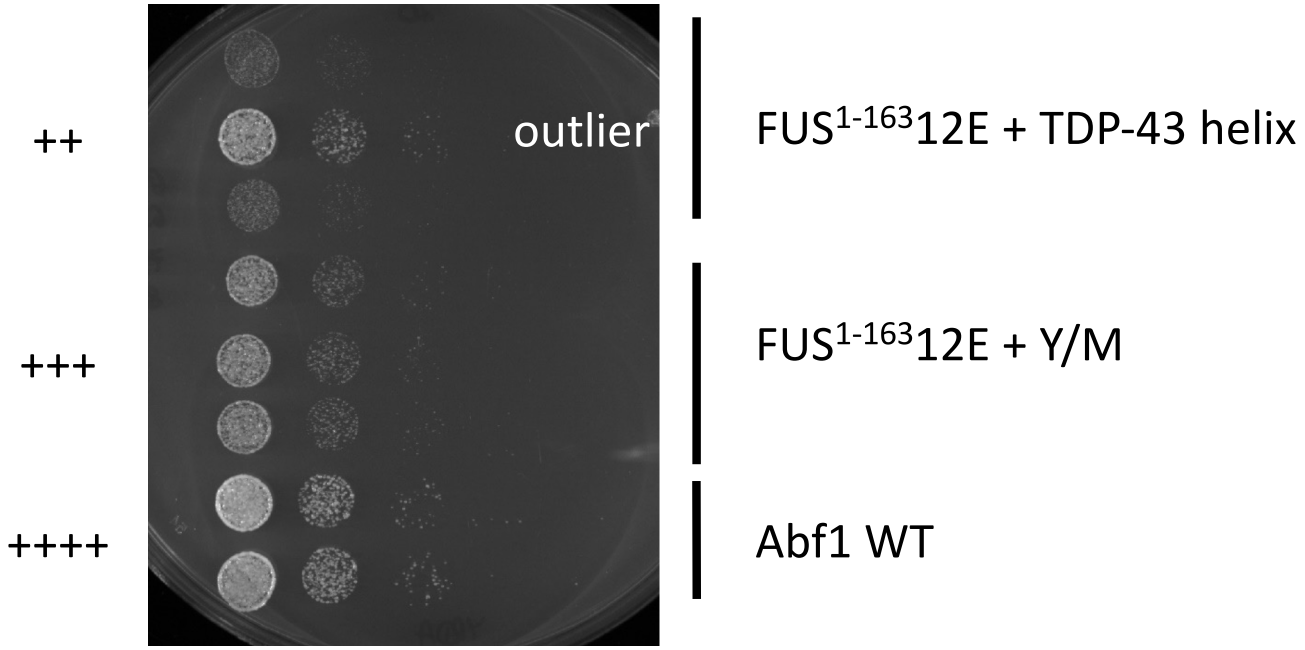
++
+++
++++
outlier
FUS$^{1-163}$12E + TDP-43 helix
FUS$^{1-163}$12E + Y/M
Abf1 WT

+++

$FUS^{1\text{-}163}$12E + GR distr.

++

$Sup35^{1\text{-}131}$ + GR

++++

Abf1 WT

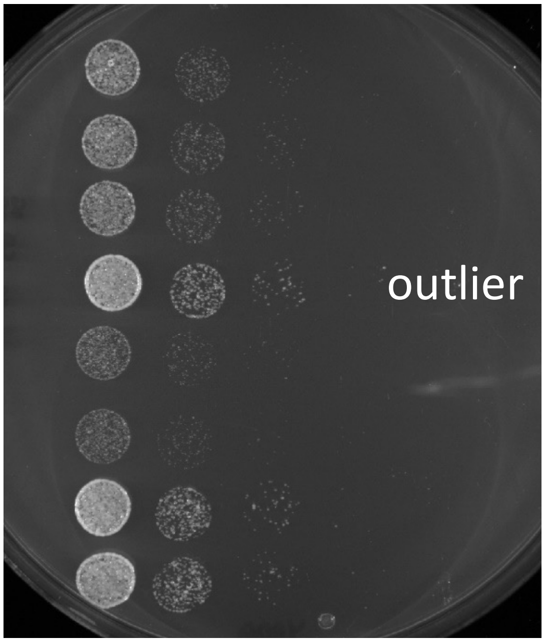


++

$Sup35^{1\text{-}131}$ + GR distr.

++++

Local shuffle 6

++++

Abf1 WT

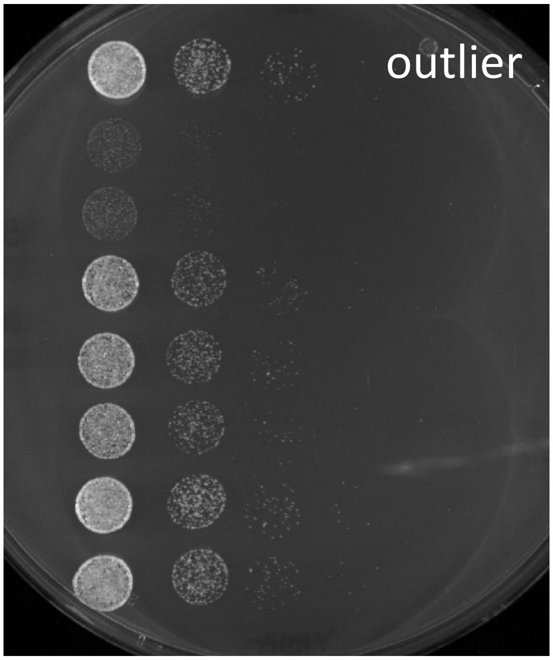

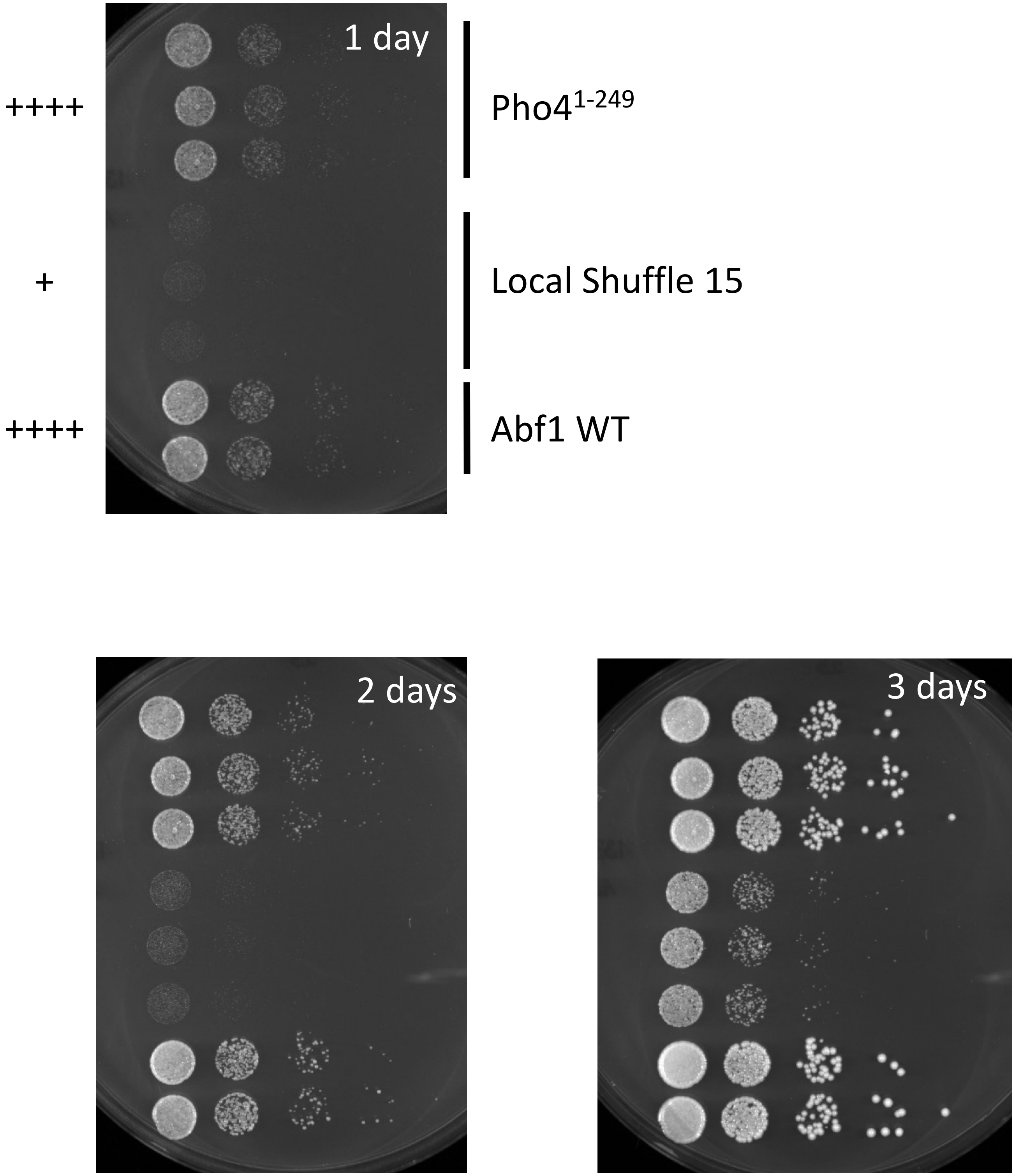

1 day
++++
Pho4$^{1-249}$
+
Local Shuffle 15
++++
Abf1 WT
2 days
3 days

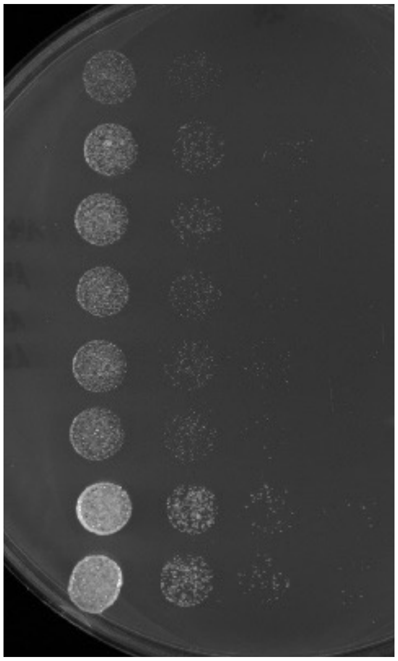

+++ Altered valence 2

+++ [No paper name yet]
(FUS12E_TDP_dist. Klon 1,2,3)

++++ Abf1 WT

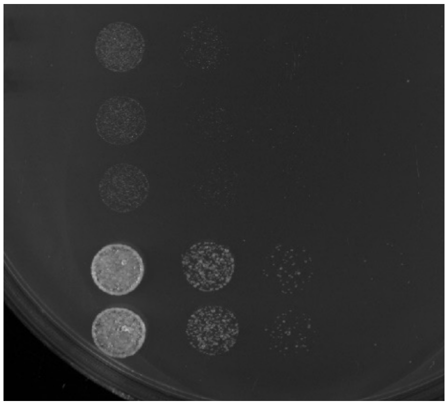

++ FUS12E + Abf1$^{G4}$ motif

++++ Abf1 WT

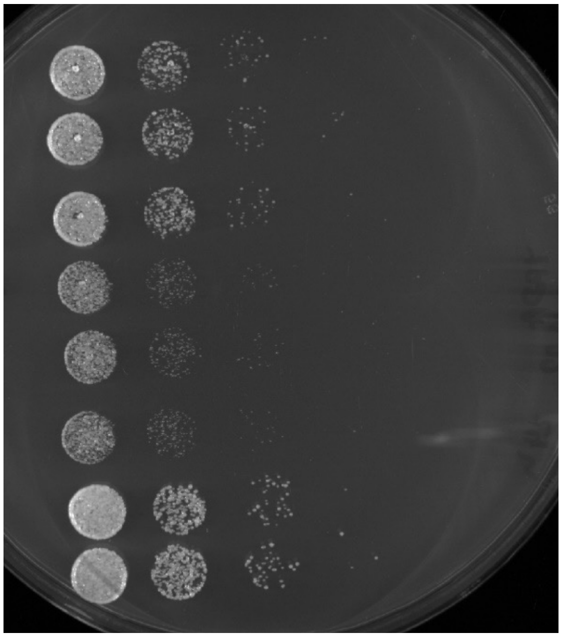

++++

[No paper name yet]
(Gal4_global_shuffle_1 Klon 1,2,3)

+++

[No paper name yet]
(Gal4_global_shuffle_2 Klon 1,4,5)

++++

Abf1 WT

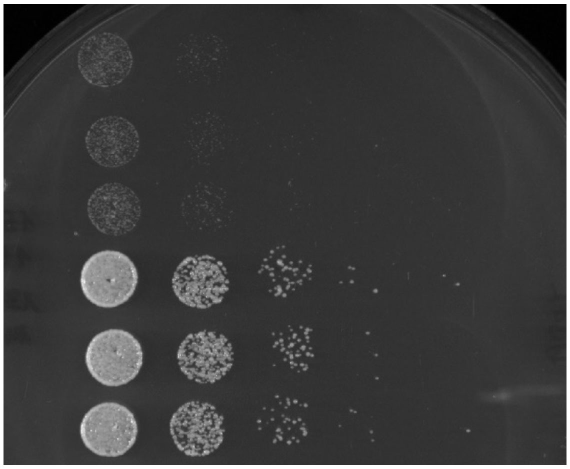

++

[No paper name yet]
(Gal4_global_shuffle_3 Klon 1,2,3)

++++

Abf1 WT

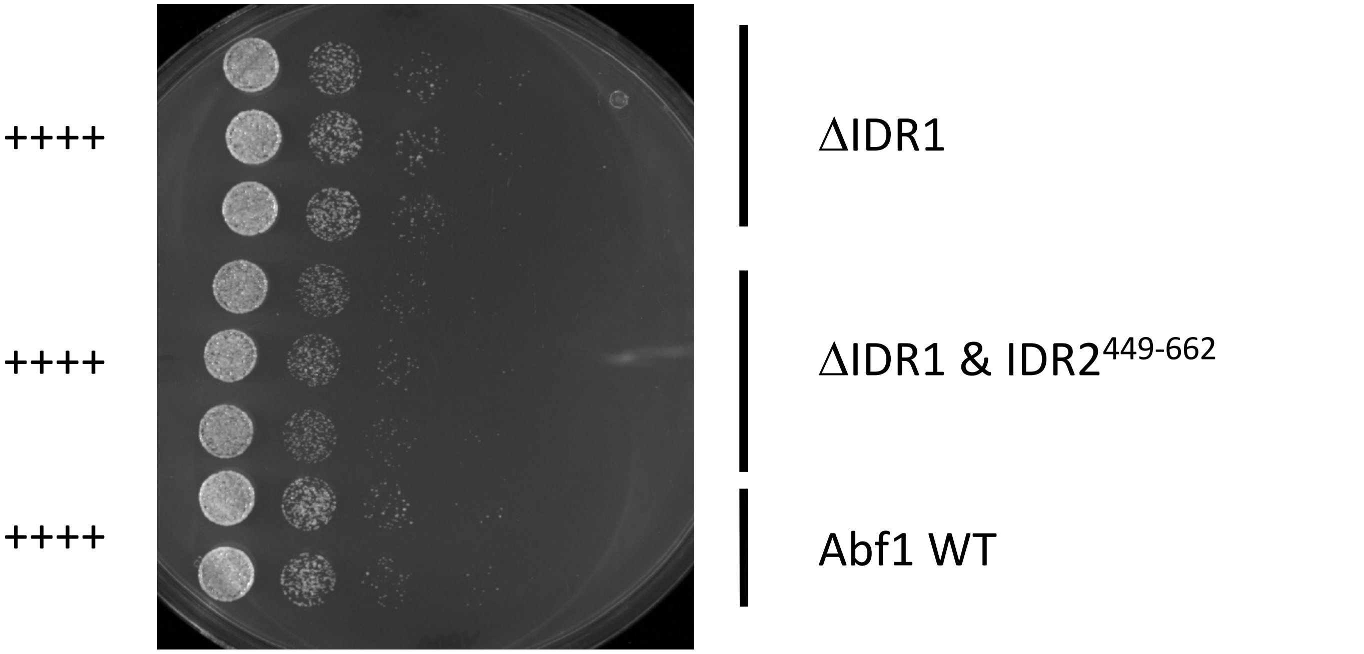
++++
ΔIDR1
++++
ΔIDR1 & IDR2$^{449-662}$
++++
Abf1 WT

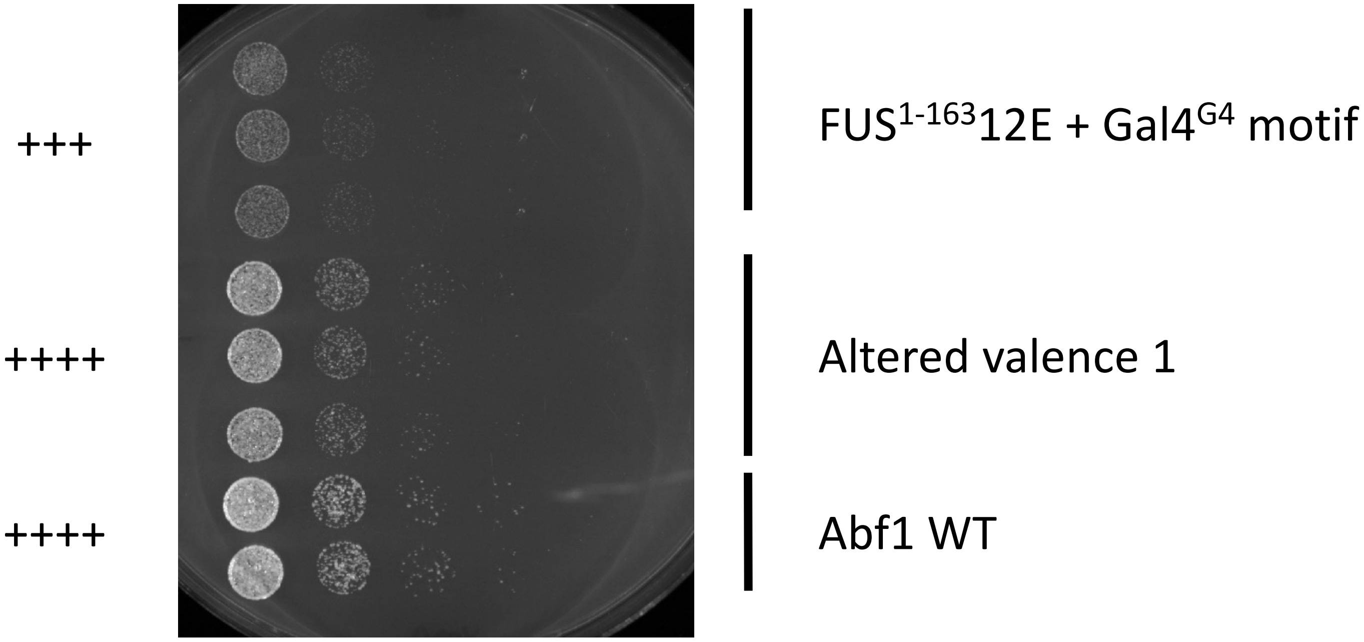
+++
FUS$^{1-163}$12E + Gal4$^{G4}$ motif
++++
Altered valence 1
++++
Abf1 WT

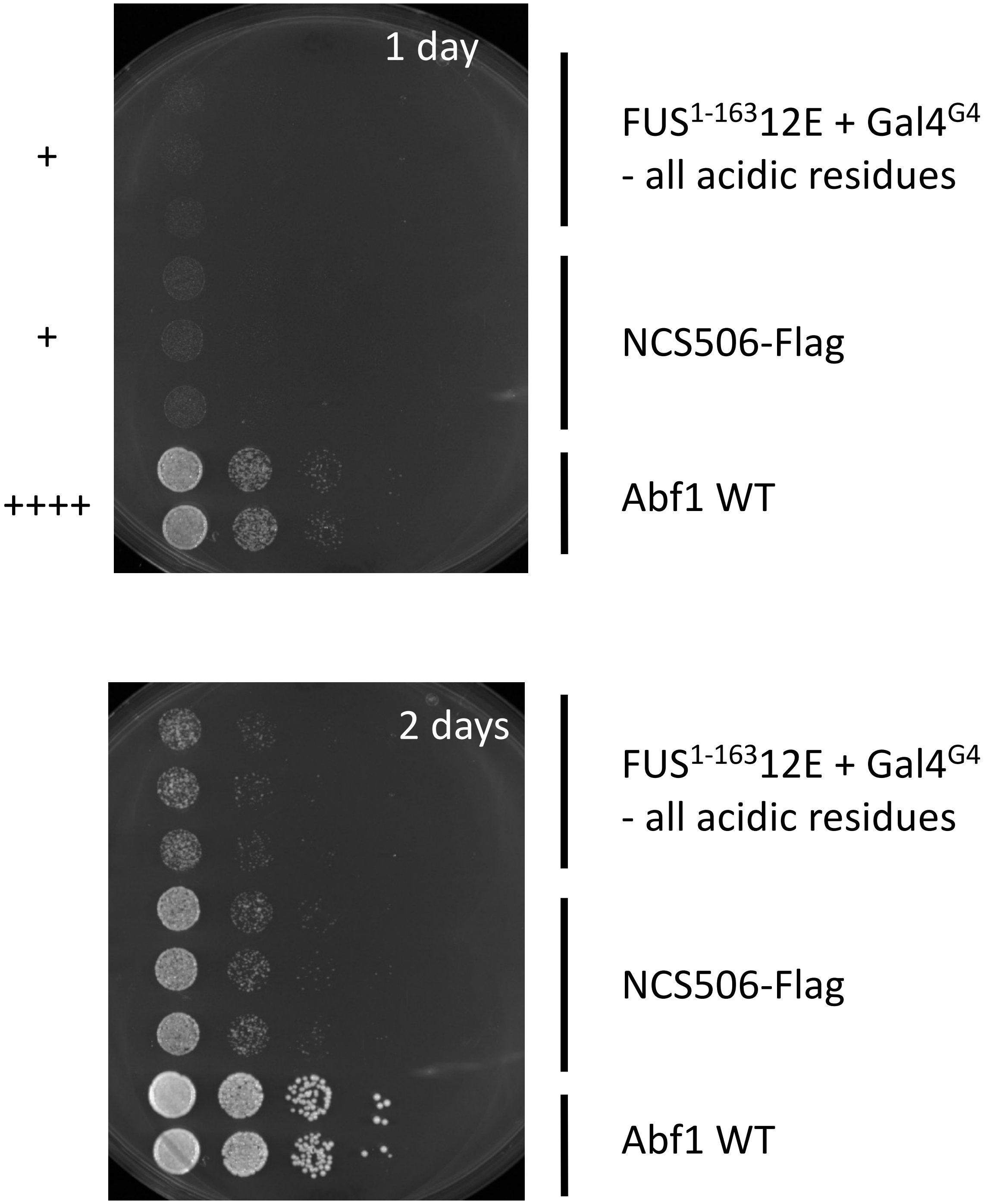
1 day
+
FUS$^{1-163}$12E + Gal4$^{G4}$
- all acidic residues
+
NCS506-Flag
++++
Abf1 WT
2 days
FUS$^{1-163}$12E + Gal4$^{G4}$
- all acidic residues
NCS506-Flag
Abf1 WT

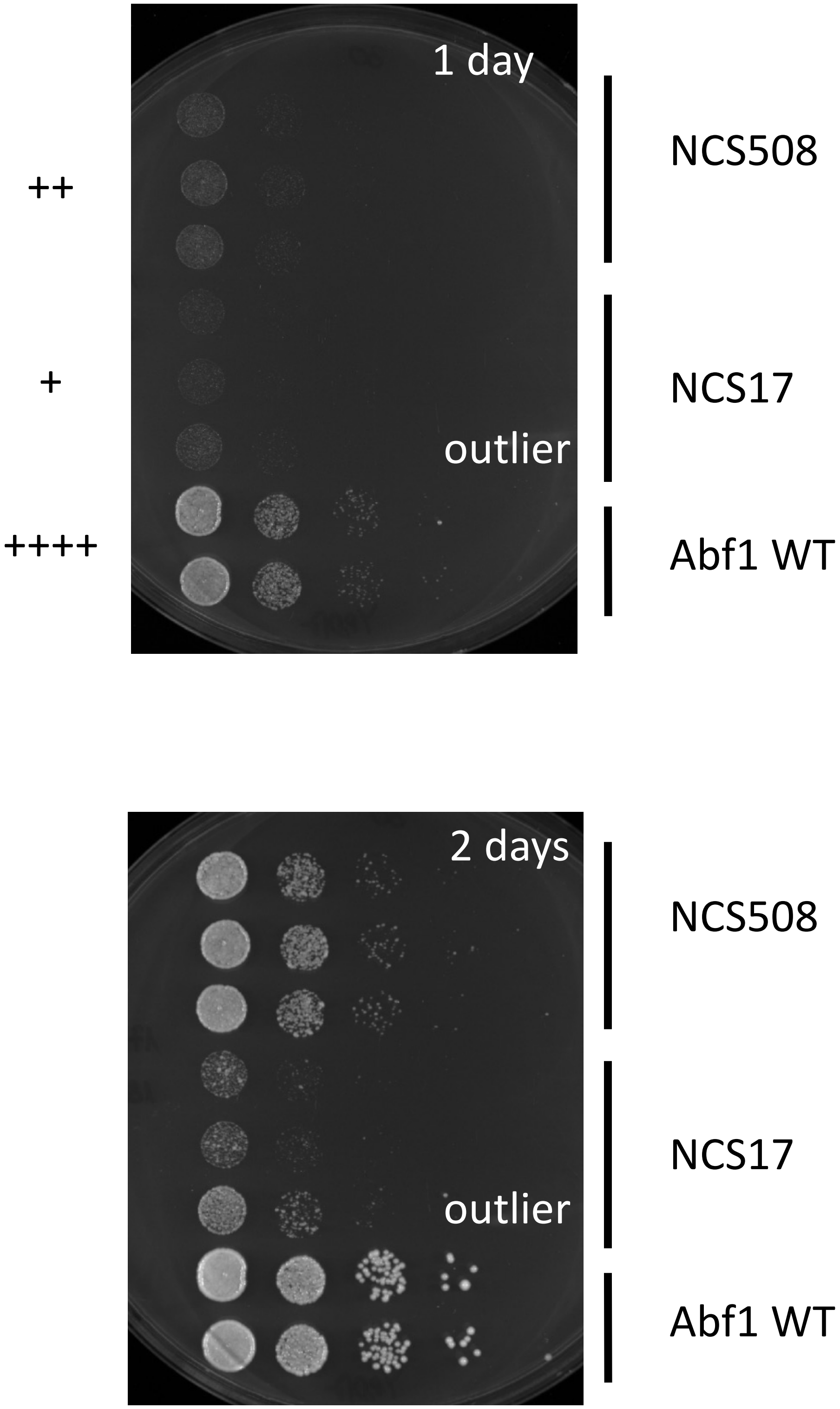
1 day
++
+
++++
outlier
NCS508
NCS17
Abf1 WT
2 days
outlier
NCS508
NCS17
Abf1 WT

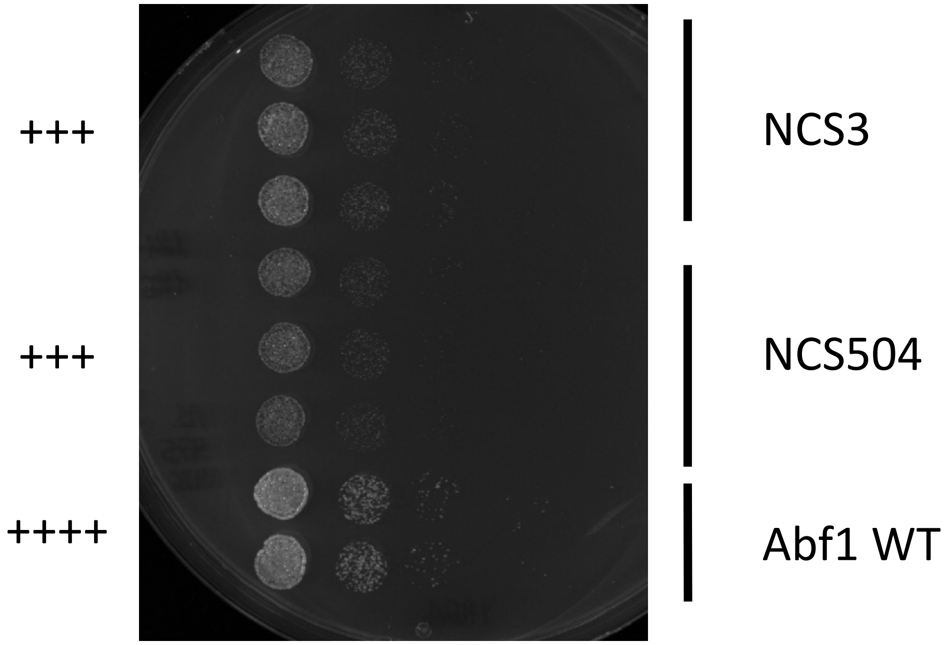
+++
NCS3
+++
NCS504
++++
Abf1 WT

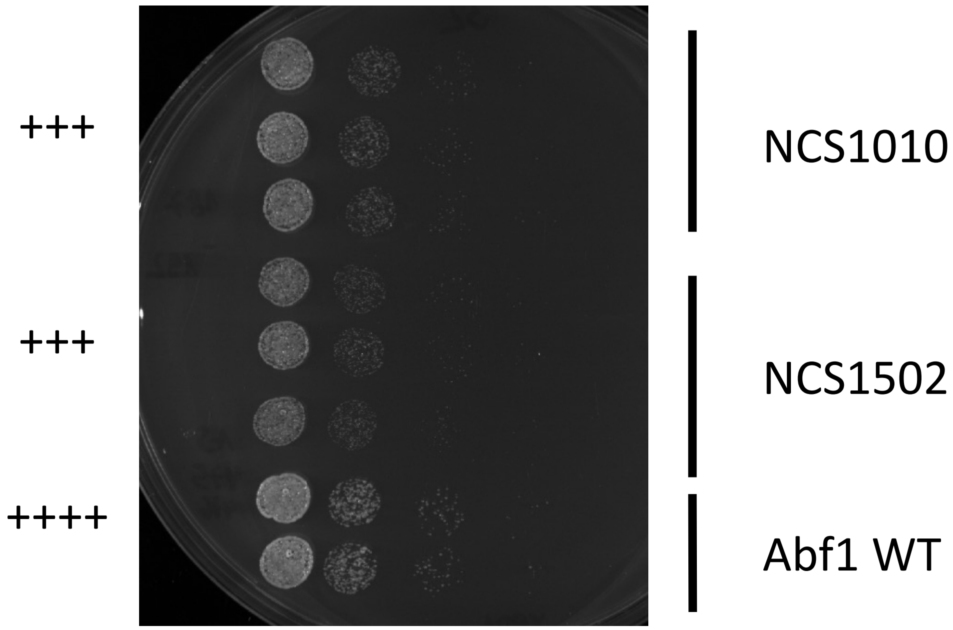
+++
NCS1010
+++
NCS1502
++++
Abf1 WT

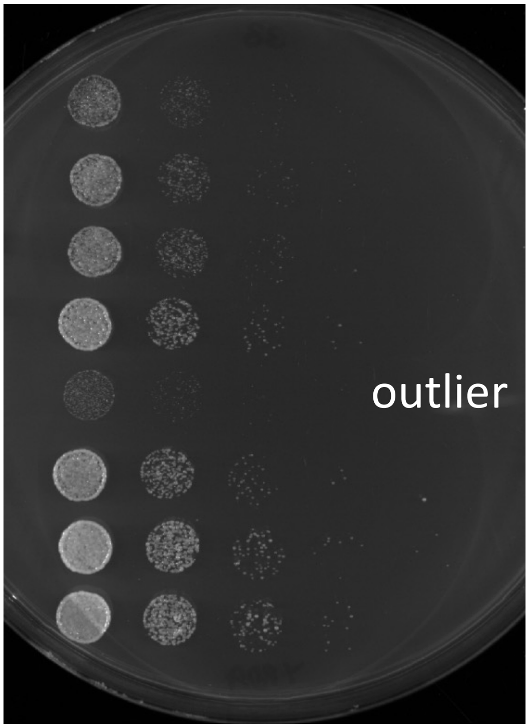


+++ NCS1508

++++ NCS509

++++ Abf1 WT

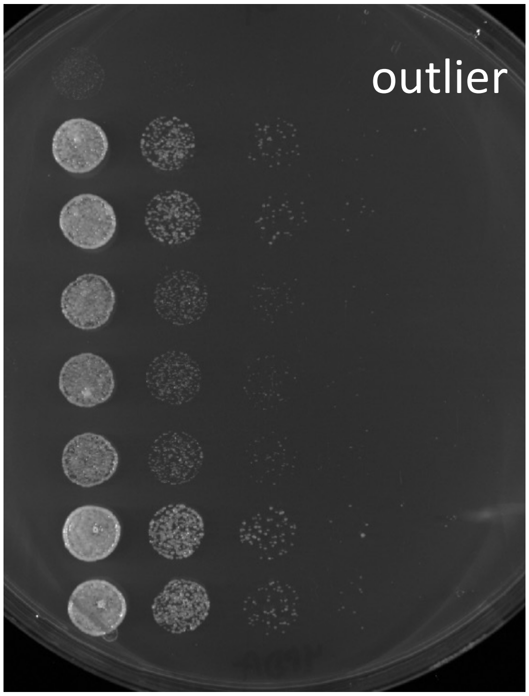


++++ NCS510

+++ NCS21

++++ Abf1 WT

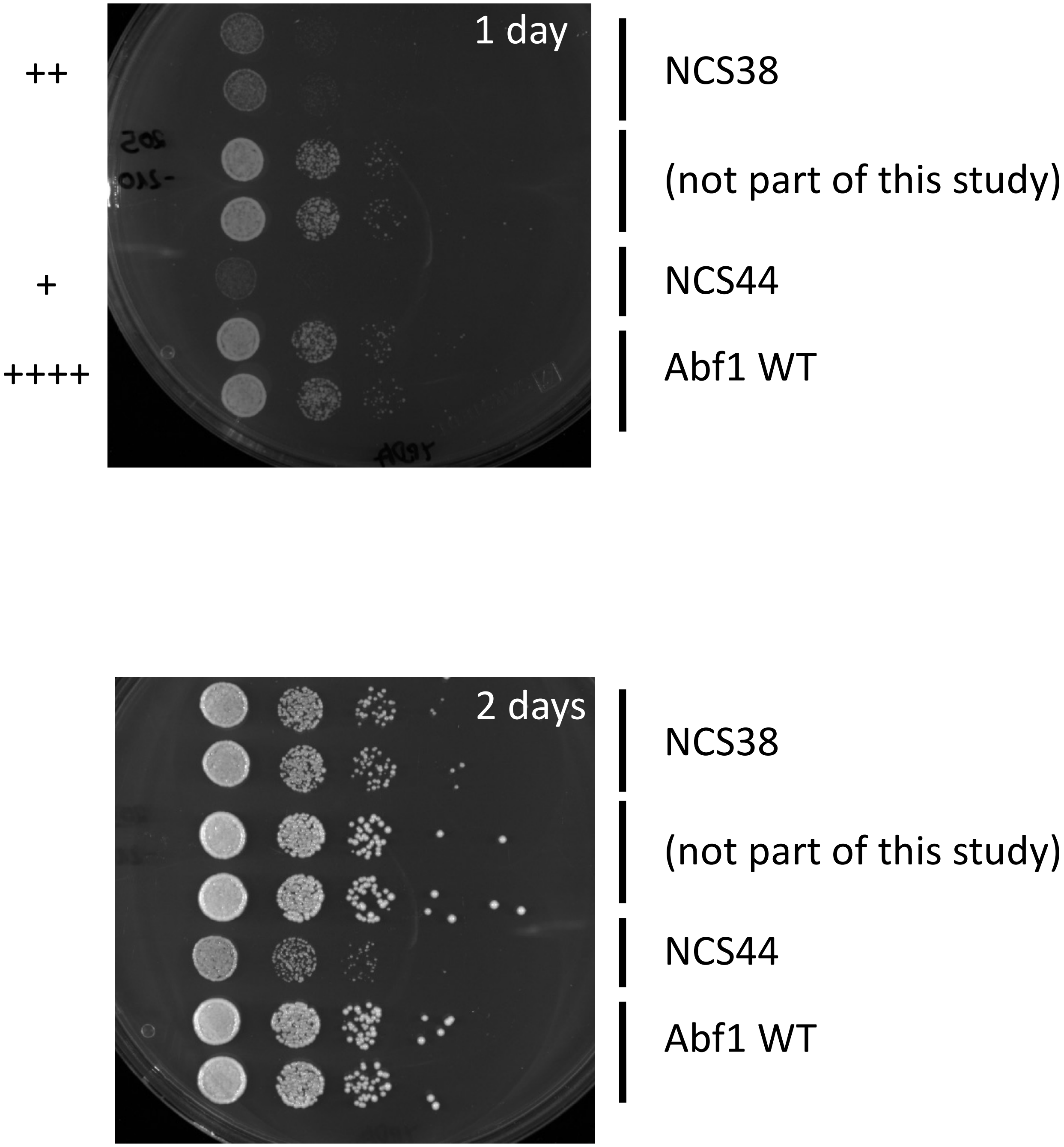
1 day
++
NCS38
(not part of this study)
+
NCS44
++++
Abf1 WT
2 days
NCS38
(not part of this study)
NCS44
Abf1 WT

++

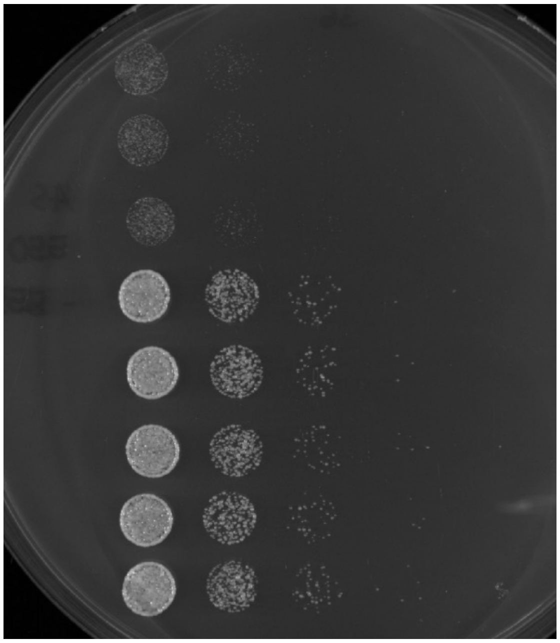

Sup35$^{1\text{-}131}$ + p65 distributed

(not part of this study)

++++

Abf1 WT

+++

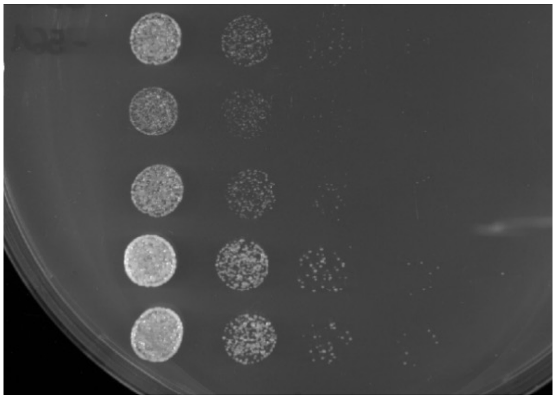

FUS$^{1\text{-}163}$12E + EM

++++

Abf1 WT

+

++++

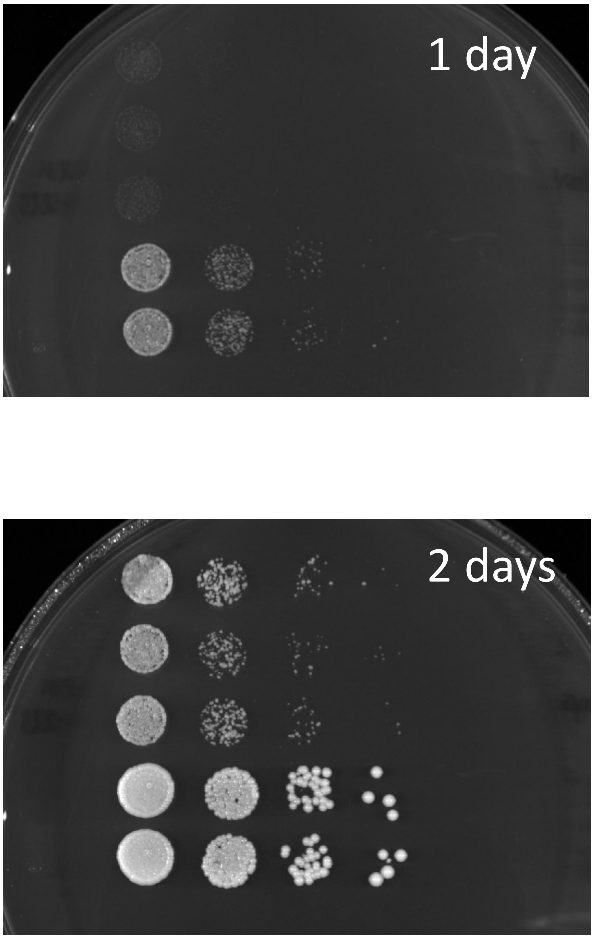


[no paper name yet]
NCS21 hydro2polar to wt

Abf1 WT
