## Supplementary Information for "Sequence- and chemical specificity define the functional landscape of intrinsically disordered regions"



Version 2.1 (October 2024)

#### Table of contents:

This PDF file includes:

Materials and Methods
Supplementary Text
Figs. S3 – S42
Captions for Tables S1 to S17

Fig. S1 is found in a separate PDF (fig_s1.pdf)
Fig. S2 is found in a separate PDF (fig_s2.pdf)
Tables S1 to S17 are found in the supplementary tables excel file, with one table per tab.

### Materials and Methods

##### Plasmids and yeast strains for 5-FOA plasmid shuffling assay and growth rate scoring by spot assay

Plasmid pRS416-Abf1 contained the wild type *ABF1* gene (a PCR fragment from 690 bp upstream and 170 bp downstream of the coding region generated with primers *pRS416_Abf1_for* and *pRS416_Abf1_rev* (**Table S2**) and genomic DNA of BY4741 as template) inserted into the EcoRI and BamHI sites of plasmid pRS416 (*1*). Plasmid pRS315-Abf1 contained the equivalent insert as plasmid pRS416-Abf1 (PCR fragment amplified with primers *pRS315_Abf1_for* and *pRS315_Abf1_rev* (**Table S2**) and genomic DNA of BY4741 as template) but inserted via BamHI and HindIII in plasmid pRS315 (*1*).

Rationally designed *abf1* mutant constructs were generated via Gibson cloning (*2*, *3*) using the PCR primers and templates in **Table S3** for insert and backbone fragments. The templates for inserts were mainly ordered by gene synthesis (ThermoFisher, **Table S3**) or amplified from genomic DNA of strain BY4741. All constructs in pRS315 contained an SV40 nuclear localization signal (amino acid sequence: PKKKRKV) at the C-terminus. All rationally designed constructs in pRS315 except for ΔIDR1 contained the 3xFlag tag introduced via Gibson cloning using primers and template as specified in **Tables S2 and S3**.

For PCR, Phusion polymerase (New England Biolabs) and thermo cycling conditions were used according to the manufacturer's recommendations. A pool of inserts for randomly mutagenized *abf1* constructs was generated by error-prone PCR (Jena Bioscience dNTP-Mutagenesis Kit #PP-101) with primers *N2-IDR2-N*dC* and *N3-IDR2-N*dC* and assembled via Gibson cloning into a backbone PCR fragment generated with primers *N1-IDR2-N*dC* and *N4-IDR2-N*dC* (**Table S4**).

For both mutagenized insert and plasmid backbone PCR, pRS315-ΔIDR1&IDR2$^{449\text{-}662}$ was used as a template. After transformation of the Gibson assembly reaction into *E. coli* strain DH5α and DNA isolation (Macherey-Nagel NucleoSpin Plasmid NoLid REF 740499.250) from single bacterial colonies, insert DNA sequences were confirmed by Sanger sequencing with primers listed in **Table S5**.

Parent diploid strain with heterozygous deletion of the *ABF1* gene was Y24962 (BY4743; MATa/MATα; *his3Δ1/his3Δ1*; *leu2Δ0/leu2Δ0*; *LYS2/lys2Δ0*; *met15Δ0/MET15*; *ura3Δ0/ura3Δ0*; *YKL112w/YKL112w::kanMX4*) obtained from EUROSCARF (www.euroscarf.de). Y24962 was transformed with plasmid pRS416-Abf1 and sporulated in liquid culture (1% potassium acetate, 5 µg/ml histidine, 25 µg/ml leucine). Sporulation and tetrad dissection were according to (*4*). Spores with both the *abf1::kanMX* deletion allele and plasmid pRS416-Abf1 were selected after tetrad dissection (MSM400, Singer Instruments) on YNB –uracil plates and subsequent growth on YPDA + kanamycin (0.2 mg/ml) selection medium.

Three strains originating from independent such spores were kept for further use: 7-A2 (MATa), 7-C5 (MATα), and 7-D4 (MATa). Their mating type was determined by colony PCR with primers as published (*5*). These strains were transformed with plasmids (pRS315-"construct name") bearing *abf1* mutant constructs detailed in **Table S3** and selected on YNB – uracil – leucine medium. The insert region of every plasmid batch was checked by Sanger sequencing prior to transformation into yeast. For each construct, the number of independent single colonies after transformation (= number of replicates, always ≥ 3) given in **Table S1** was re-streaked for single colonies on YNB – uracil – leucine selection medium, and single colonies thereof streaked out as patches on the same medium. From these patches, strains were streaked again as patches on 5-FOA –leucine plates (0.67% YNB, 0.2% SC–ura –leu drop-out mix, 2% glucose, 50 µg/mL uracil, 0.1% 5-FOA (Toronto Research Chemicals, #F595000 or Diagnostic Chemicals Limited, #1555)) and incubated for 2-3 days at 30°C. Viability on 5-FOA plates was visually scored by comparison to known viable or inviable strains on the same plate. Inviable strains were distinguished from viable strains because they showed a smeary appearance and, at most tiny/sparse single colonies. In contrast, viable strains grew as one more or less dense patch. (**fig. S1**, **Table S1**). Constructs viable on 5-FOA plates were tested for uracil auxotrophy on YNB –uracil medium, and three strains from independent single colonies that were viable on 5-FOA and inviable on –uracil medium were stored as glycerol culture. Three independent single colonies of strains with constructs scored inviable on 5-FOA plates were stored from the patches on YNB –uracil – leucine medium.

The three strains originating from the three independent single colonies for each viable Abf1 construct were scored for growth rate by spotting serial dilutions in water (1:10, 1:100, 1:1000, 1:10,000) of overnight cultures adjusted with YPDA media to the same $OD_{600}$ of 0.5 (Genesys 20 photometer, ThermoFisher) on YPDA plates and incubation at 30°C for 1-3 days (**fig. 2**). Growth rates were scored visually into four broad categories (+1 to +4) in comparison to the rate of two independent clones of strain 7-A2 bearing plasmid pRS315-Abf1 spotted on the same plates and equivalent to growth rate score +4 (**fig. S2**, **Table S7**). The correct sequence of the plasmid insert was checked for at least one clone of each viable construct by Sanger sequencing of a colony PCR product from a colony on the spot dilution assay YPDA plates using the sequencing primers given in **Table S5**.

Three independent single colonies of strains with constructs scored inviable as patches on 5-FOA plates were stored as glycerol cultures from the patches on YNB –uracil – leucine medium. Starting from these glycerol cultures, all rational mutagenesis constructs and the random mutagenesis construct NCS506-Flag were also tested for viability by spot serial dilutions as above but on 5-FOA –leucine plates. In most cases, this immediately confirmed the inviability of the constructs. Nonetheless, if a strain showed notable growth in this spot dilution assay, the plasmid insert was checked by Sanger sequencing of colony PCR products from colonies on the 5-FOA –leucine plates. In all but two cases, colonies of clones scored as inviable by the patch assay that showed growth in the spot dilution assay on 5-FOA –leucine plates carried inserts with full-length or IDR2 sequences of WT Abf1. We speculate this is driven by recombination between plasmids pRS416-Abf1 and pRS315-[abf1 mutant construct] during over night incubation in YNB –leu –ura medium before spotting the dilutions. Only the constructs NCS506-Flag and FUS1-16312E + Gal4G4 –all acidic residues showed very slow growth, visible after prolonged incubation, in this spot dilution assay and still carried the proper construct plasmid. Clones of these two constructs were also included in the growth rate scoring togther with the clones of the other viable constructs and received the viability score +1 (**Table S7**).

**Chromatin immunoprecipitation and quantitative real-time PCR**

ChIP assays were performed for all inviable constructs bearing a Flag-tag and some viable constructs as indicated by the respective ChIP values in **Table S1**. Corresponding yeast strains were grown to logarithmic phase in –uracil – leucine selection medium (inviable constructs or viable constructs together with plasmid pRS416-Abf1) or in –leucine selection medium (viable constructs) at 30°C. One hundred $OD_{600}$ units (Zeiss PMQ II photometer) were crosslinked for 15 min at room temperature with 1% final concentration of formaldehyde while shaking slowly, and crosslinking was quenched by 250 mM final concentration of glycine for 10 min at room temperature while shaking slowly. Cells were harvested by centrifugation for 5 min at 4,000 rpm (Heraeus Cryofuge 6000i) and 4°C. Cell pellets were washed with ice-cold ST buffer (10 mM Tris-HCl pH 7.5, 150 mM NaCl) and washed twice in FA lysis buffer (50 mM HEPES-KOH pH 7.5, 150 mM NaCl, 1 mM EDTA, 1% Triton X-100, 0.1% sodium deoxycholate, 0.1% SDS). Washed pellets were flash-frozen in liquid nitrogen and stored at -80°C. For cell lysis, pellets of 50 $OD_{600}$ units were resuspended in 250 µl FA lysis buffer supplemented with protease inhibitors (1x cOmplete EDTA-free protease inhibitor cocktail (Roche), 1 mM PMSF). Zirconia beads (350 µl BioSpec Products 0.5 mm glass beads, 11079105) were added, and cells were lysed in a bead beater (Percellys 24, Bertin Technologies). Lysis efficiency was controlled by comparing $OD_{600}$ values before and after bead beating and was 75-90%. Beads were removed by centrifugation, and cell lysates were adjusted to 1.2 ml with FA lysis buffer and chromatin sheared by sonication at 4.5°C (Bioruptor, Diagenode, 30 cycles, 30 sec on/off, high intensity) and fragment lengths were usually around 1 kb as assessed in a Bioanalyzer after DNA purification (high sensitivity DNA chip). If this fragmentation range was not met, preparation of the sample was repeated starting from a new culture. The chromatin solution was precleared by incubation with 25 µl protein G beads for 1-2 hrs at 4°C with soft rolling. For immunoprecipitation, 500 µl of precleared chromatin solution was incubated with or without (no antibody control) 2.5 µl anti-Flag M2 antibody (Sigma) overnight at 4°C with soft rolling and for four further hours with 25 µl of pre-blocked (10% BSA) protein G beads. Beads were washed twice with high salt FA lysis buffer (0.5 instead of 0.15 M NaCl), twice with ChIP Wash buffer (10 mM Tris-HCl pH 8.0, 250 mM LiCl, 0.5% NP-40, 0.5% sodium deoxycholate; 1 mM EDTA) and once with TE buffer (10 mM Tris-HCl, pH 8.0; 1 mM EDTA). Elution was done twice for 15 min at 65°C while shaking in 100 µl ChIP Elution buffer (50 mM Tris-HCl pH 7.5, 10 mM EDTA, 1 % SDS). Eluates were pooled and crosslinking reversed by incubation overnight at 65°C while shaking. 50 µl of chromatin sample for input DNA was taken off after sonication, adjusted to 200 µl with ChIP Elution buffer, and treated in parallel for DNA purification. Immunoprecipitated or input chromatin was treated with RNaseA (0.5 mg/ml) for 30 min at 37°C, then with proteinase K (0.5 mg/ml) for 2 hrs at 37°C while shaking. DNA was purified with AmPure beads (1.8 vol), according to the manufacturer's instructions.

Input DNA (1 µl, 1:100 dilution in water), immunoprecipitated DNA (1 µl) and mock precipitated (no antibody) DNA (1 µl) were analyzed in triple replicates by quantitative real-time PCR (qPCR; Light Cycler 480 II (Roche)) in 10 µl total volume using the primers given in **Table S6** at a final concentration of 3 µM and the 1x Fast Sybr Green Master Mix (Applied Biosystems). ChIP values were calculated from qPCR CT values as an average of PCR triplicate averages for each of the three amplicons with Abf1 site divided by the average of PCR triplicate averages for each of the three amplicons without Abf1 site. For unknown reasons, some replicates of inviable constructs failed to score in the ChIP assay (= ChIP value close to one; **Table S1**). Nonetheless, if at least one replicate was positive (= ChIP value > 10), this confirmed that the respective construct was expressed, imported into the nucleus, and could bind specifically to Abf1 sites. ChIP values for inviable versus viable constructs were on average lower **fig. S9**. This was likely due to the competition between the untagged wild type Abf1 and the Flag-tagged inviable construct for binding to Abf1 sites. This competition was unavoidable for inviable constructs but usually not the case for viable constructs as these were tested in the absence of the untagged wild type Abf1. To confirm this, we tested two viable constructs in the presence of untagged wild type Abf1 to mimic the competition between wildtype Abf1 and the Flag-tagged construct (marked in **fig. S9**) and found that they indeed showed ChIP values that matched the expected values from the inviable constructs (p = 0.58, Mann Whitney U test) in the range of the viable constructs.

**Abf1 anchor away strains and growth conditions for RNA-seq and ODM-seq analyses**

The Abf1 anchor away strain ABF1-aa-V5 clone A9 was obtained from Wim de Jonge and Frank Holstege(*6*). The strain was transformed with the same pRS315-based plasmids encoding wild type Abf1, select Abf1 constructs, or the empty pRS315 plasmid (no Abf1 control), as was used for the 5-FOA plasmid shuffle assays. At least three independent clones were isolated and stored as glycerol cultures after transformation and used for biologically independent replicate experiments. Anchor away strains were grown to log phase (ca. $OD_{600}$ of 0.4, Genesys 20, ThermoFisher) in YNB –leucine media, incubated for 75 min with 8 mg/ml rapamycin (Santa Cruz) and crosslinked for 5 min (for ODM-seq) with 1% formaldehyde at room temperature. Formaldehyde crosslinking was quenched for 15 min with 250 mM glycine at room temperature.

**RNA-seq**

Total RNA was prepared after Abf1 anchor away induction with the RNeasy kit (Qiagen) according to the manufacturer's protocol. Briefly, 4 $OD_{600}$ units (Genesys 20, ThermoFisher) of cells were washed with PBS (50 mM phosphate buffer pH 7, 150 mM NaCl) and flash frozen in liquid nitrogen. The cell wall was broken up by incubation with beta-mercaptoethanol and Zirconia bead-beating (6x 3 min 30 Hz with 3 min breaks, Precellys, Bertin). The cell lysate was cleared by centrifugation, mixed with ethanol, and bound to a column provided in the kit. We also employed the optional DNaseI digestion on the column. RNA was washed on the resin, eluted, and digested using DNase I (Qiagen) in solution. After ethanol precipitation, the total RNA preparation was converted into sequencing libraries with the NEBNext Ultra II directional RNA library prep kit for Illumina.

**Cloning, expression, purification, and characterization of M.SssI DNA methyltransferase**

*DNA methyltransferase cloning*

The DNA sequence coding for the M. SssI methyltransferase, where the TGA stop codons coding for tryptophan in the original organism were replaced by TGG codons, was PCR amplified from plasmid pCMV-FLAG-4azf-M.SssI (gift from Kevin Ford, Kings College London) and cloned into the NcoI and SalI sites of the pBAD24 plasmid (87399, ATCC) to achieve tight transcriptional control by the *E. coli* arabinose-regulated *araBAD* promoter. An in-frame C-terminal 6xHis-tag was introduced via the 3'-primer during the PCR. The resulting expression plasmid is called pBAD24-M.SssI-His6, and was transformed (selection on LBamp plates with 0.2% glucose) into *E. coli* One Shot™ TOP10 (ThermoFisher) yielding the expression strain for M.SssI expression.

A single colony streaked out on LBamp plates + 0.2% glucose was used to inoculate a 100 ml overnight culture in LBamp + 0.2% glucose at 37°C and 130 rpm. Six times 1 l LBamp media (without glucose) were inoculated with 10 ml each of the overnight culture and incubated at 37°C and 130 rpm until $OD_{600}$ reached 0.5 (Genesys 20, ThermoFisher). Expression was induced by the addition of 0.0002% arabinose for 4 h at 37°C and 130 rpm. Cells were harvested by centrifugation (4000 rpm, 30 min, 4°C, Avanti JXN-26, JLA 8.1000 rotor, Beckman-Coulter), resuspended, and combined in 75 ml Lysis Buffer (50 mM Tris-HCl pH 8, 300 mM NaCl). Aliquots of 35 ml were snap-frozen in liquid nitrogen and stored at -70°C.

*DNA methyltransferase purification*

Purification started with one frozen cell pellet aliquot that was thawed at room temperature, topped up to 50 ml with Lysis Buffer, and supplemented with 1 mg/ml chicken egg white lysozyme (28260, Serva) and protease inhibitors (0.7 µg/ml pepstatinA, 1 µg/ml leupeptin, 1 µg/ml aprotinin, 0.2 mM PMSF), incubated on ice or 30 min and then sonicated (Branson sonifier 250D, 6 cycles of 10 s burst and 10 s break at 50% peak power). Lysed cells were centrifuged for 1 h at 4°C and 20,000 g (Optima XPN-80, Ti-45 rotor, Beckman-Coulter). During centrifugation, 5 ml slurry of Ni-NTA agarose (745400.25, Macherery-Nagel) was washed twice with 20 ml Lysis Buffer (1 min, 1000 rpm, 4°C Eppendorf 5810R centrifugation for collecting the agarose) and resuspended in 30 ml Lysis Buffer, distributed into three 10 ml aliquots and Ni-NTA beads were collected by centrifugation and the supernatant discarded. After ultracentrifugation, the lysed cells' supernatant was evenly distributed onto the pre-washed

Ni-NTA beads and incubated for 1.5 h at 4°C on a rotating wheel. Beads were collected by centrifugation (2 min, 1000 rpm, 4°C, Eppendorf 5810R), the supernatant discarded, and the beads resuspended in 20 ml cold Lysis Buffer with protease inhibitors and transferred to a gravity flow column (Econo-Pac, 7321010, Bio-Rad) at room temperature. The column was then washed by gravity flow with cold 100 ml Lysis Buffer, cold 20 ml Wash Buffer (50 mM Tris-HCl pH 8, 250 mM NaCl, 20 mM imidazole-HCl pH 6.6). After the addition of 2 ml cold Elution Buffer (50 mM Tris-HCl pH8, 250 mM NaCl, 300 mM imidazole-HCl pH 6.6), the column was closed and lightly vortexed and incubated on ice for 10 min with light vortexing every 2 min. One 2 ml-fraction was collected on ice by gravity flow, and three 1 ml-fractions after the addition of 3 ml Elution Buffer. Fractions of the steps during the purification were analyzed by SDS-PAGE (**fig. S39A**).

Fractions from the column enriched for the 42 kDa M.SssI methyltransferase were pooled and directly loaded in the cold room onto a HiLoad 16/600 Superdex 200 PG size exclusion column (Cytiva) equilibrated with Size Exclusion Buffer (20 mM Tris-HCl pH 8, 500 mM NaCl, 0.1 mM EDTA, 0.1 mM DTT). Elution was with Size Exclusion Buffer at 0.5 ml/min flow rate, and 2 ml-fractions were collected in the cold. Fractions were analyzed by SDS-PAGE (**fig. S39B**) and fractions enriched for M.SssI combined and dialyzed against Dialysis Buffer (10 mM Tris-HCl pH 8, 0.2 mM EDTA, 1 mM DTT, 50% glycerol), which also concentrates the protein solution. The dialyzed and concentrated M.SssI solution was analyzed again by SDS-PAGE (**fig. S39C**), and the methyltransferase activity was quantified by restriction enzyme activity assay. Aliquots corresponding to ca. 440 units were snap frozen in liquid nitrogen and stored at -80°C.

*Quantification of CpG-specific DNA methyltransferase activity:*
Five hundred ng of plasmid pUC19-ftz (containing a fragment of the Drosophila ftz locus from sequence TAGTTTCCTAATGAT to GCCGAAGATGATGCT cloned into pUC19 multiple cloning site) were incubated in 122 µl final volume with increasing amounts of purified M.SssI solution or commercially aquired M.SssI (M0226, NEB) in NEB buffer 2 (NEB) supplemented with 160 µM SAM for 1 h at 25 °C. The reaction was stopped by the addition of 1/10 volume 10x Stop Buffer (50 mM Tris-HCl, pH 7.5, 4% SDS, 100 mM EDTA) and incubated with 5% proteinase K. After incubation for 1 h to overnight at 37°C, 1/5 volume of 5 M $NaClO_4$ was added to the solution, then an equal volume of phenol (77607, Sigma), vortexed, an equal volume of chloroform-isoamylalcohol (24:1), vortexed, and phases separated by centrifugation (Hettich Microliter). The DNA was recovered from the aqueous phase by ethanol precipitation and resuspended in 37 µl of rCutSmart Buffer (NEB). The DNA was incubated with 30 units each of BamHI-HF and PvuI-HF (NEB) for 1 hr at 37°C. The complete reaction was loaded onto a 1% agarose gel in 1xTAE buffer and electrophoresed for 1.5 h at 100 V. The DNA in the gel was stained with 5 µl/100ml Midori Green solution (Nippon Genetics Europe) for a few minutes and directly imaged (Chemidoc, BioRad) (**fig. S40**). BamHI linearized the 5769 bp pUC-ftz plasmid regardless of CpG methylation. PvuI cut in a CpG methylation-sensitive way and yielded 2544, 2329, and 896 bp fragments in combination with the BamHI cut if cleavage was complete, but only the 5769 linearized plasmid band if complete CpG methylation blocked PvuI digestion. Comparison of the degree of blocked PvuI digestion between reactions with purified and

commercial M.SssI (NEB) allowed us to quantify the specific activity of the purified M.SssI according to the unit definition by NEB.

The M.SssI preparation was checked for contaminating nucleases by incubation with 500 ng supercoiled pUC19-ftz plasmid in either rCutSmart or NEB Buffer 2 (NEB) for 2 h or overnight at 25°C. The reaction was electrophoresed in 1% agarose 1x TAE gel, stained, and imaged. The persistence of the supercoiled form demonstrated the lack of contaminating nucleases (**fig. S40B**).

**Absolute nucleosome occupancy mapping by ODM-seq**
Chromatin was prepared from formaldehyde-crosslinked yeast cultures as described (*7*). Chromatin corresponding to 0.1 g wet cell pellet was washed in cold Kladde Buffer (20 mM HEPES-NaOH pH 7.5, 70 mM NaCl, 0.25 mM EDTA pH 8.0, 0.5 mM EGTA pH 8.0, 0.5% (v/v) glycerol), resuspended in 870 µl of Kladde Buffer and incubated with 4000 units M.SssI methyltransferase at 25°C. After 90 min, 4000 units M.SssI were added, together with plasmid pUC19-ftz which was included as spike-in control. After a total of 180 min, one 430 µl aliquot was removed, and plasmid pFMP233 (*8*) was added to the remaining aliquot as a second spike-in contro. The reaction was further incubated for 60 min (240 min total) at 25°C. A similar degree of CpG methylation after 180 vs. 240 min (**fig. S25**) in combination with ongoing CpG methylation as probed by methylation of the spike-in plasmids (**fig. S26**) indicated saturated CpG methylation. After DNA methylation, the reaction was stopped by the additon of 1/10 volume 10x Stop Buffer. The resulting DNA was purified by proteinase K digestion, phenolization, ethanol precipitation, and then resuspended in TE Buffer. Nanopore sequencing libraries were prepared and sequenced on a Promethion by the Laboratory for Functional Genomics (LaFuGa), Gene Center, LMU Munich.

**Cloning, expression, and purification of WT Abf1, and constructs FUS$^{1\text{-}163}$12E and IDR1/2**
Expression plasmids for WT Abf1 and Abf1 variant constructs were constructed in pPROEX as in Krietenstein et al. (*9*), but starting from the respective pRS315-plasmids with the respective Abf1 constructs and using the primers for backbone and insert PCR, respectively, which were combined by Gibson cloning:

| | |
|---|---|
| pPROEX_V_rev | AATTTGTCCATggGATCCATGG |
| Abf1_I_for | GATCCCATGGACAAATTAGTCGTGAAT |
| Abf1_I_rev | ACCGCATGCCTCGAGCTActtatcgtcatcgtctttg |
| pPROEX_V_for | CTCGAGGCATGCGGTACC |

WT Abf1 and variant proteins were expressed using BL21 star (DE3 pLysS) strain (ThermoFisher). 100 mL LBamp media were inoculated with a single colony and incubated overnight at 130 rpm, 37 °C. 20 ml of this preculture were used to inoculate 1 l of LBamp and incubated at 130 rpm, 37°C until $OD_{600}$ was 0.5-0.6 (Genesys 20, ThemoFisher). IPTG was added to a final concentration 1 mM for 4 h at 37 °C, 130 rpm. Cells were collected by centrifugation (4000 rpm, 20 min, 4 °C, Avanti JXN-26, JLA 8.1000 rotor, Beckman-Coulter), combined a 50 ml Greiner tube in 30 ml Lysis Buffer with protease inhibitors and centrifuged (1 min, 1000 rpm, 4°C Eppendorf 5810R). The pellet was flash-frozen and stored at -70°C.

For purification in the cold room, the cell pellet was thawed on ice, resuspended in 40 ml Lysis Buffer with mg/ml chicken egg white lysozyme, and incubated for 30 min on ice. Cell lysis was completed by sonication on ice (Branson sonifier 250D, 6 cycles of 10 s burst and 10 s break at 50% peak power). The cell lysate was centrifuged at 45,000 rpm for 1 h at 4°C (Optima XPN-80, Ti-45 rotor, Beckman-Coulter), and the supernatant collected and transferred to a 50 ml Greiner tube where it was mixed with pre-washed Ni-NTA resin (see above) equivalent to 2 ml slurry. This mixture was incubated for 1.5 h at 4°C on a rotating wheel, then washed with 100 ml Lysis Buffer and with 20 ml Wash buffer and eluted with 5-10 ml Elution Buffer. The eluate was dialyzed overnight at 4°C against 1-2 l Dialysis Buffer II (20 mM HEPES-NaOH pH 7.5, 70 mM NaCl, 0.1% Tween, 40% glycerol, 1 mM DTT) and concentrated if necessary (Amicon Ultra -4 Ultracells 10K or 30K, Millipore). The dialyzed and maybe concentrated sample, for WT Abf1 and $FUS^{1-163}$12E variants, was loaded onto a 1ml HiTrap Q FF anion exchange chromatography column (Cytiva) equilibrate in Low Salt Buffer (50 mM Tris-HCl pH 8.0, 80 mM NaCl, 10% glycerol, 1 mM DTT, 1 mM EDTA) and washed to baseline with Low Salt Buffer. Elution was by linear gradient from 0% to 100% High Salt Buffer (50 mM Tris-HCl pH 8.0, 350 mM NaCl, 10% Glycerol, 1 mM DTT, 1 mM EDTA) over 20 column volumes. The dialyzed and maybe concentrated IDR1 IDR2 construct was loaded onto a 1 ml Hi Trap Heparin HP column (Cytiva) equilibrated with Heparin buffer A (25 mM Tris-HCl pH 7.6, 10% glycerol, 1 mM DTT, 50 mM NaCl, 1 mM EDTA), washed to baseline with Heparin buffer A and eluted by linear gradient from 0% to 100% of Heparin buffer B (25 mM Tris-HCl pH 7.6, 10% glycerol, 1 mM DTT, 1 M NaCl, 1 mM EDTA) over 30 column volues.

The purified proteins (**fig. S41**) were dialyzed overnight against Storage Buffer (20 mM HEPES-NaOH pH 7.5, 350 mM NaCl, 0.1% Tween, 40% glycerol, 1 mM DTT), aliquotized, flash frozen in liquid nitrogen and stored at -80°C.

**Genome-wide chromatin reconstitution and remodeling assay**

Genome-wide chromatin reconstitution, remodeling assay, MNase-seq assay, and Illumina sequencing were done as described in (*10*), with the exception that (a) Illumina sequencing was in paired-end mode and (b) for the samples shown in **Fig. 6E**, recombinant human octamers were used, reconstituted from H2A/H2B dimers and H3/H4 tetramers according to Kunert et al. (*11*). Recombinant human histones were obtained from the Histone Source (Colorado State University). To verify the robustness of this assay, we repeated the experiment (as seen in **Fig. 6E**) using Drosophila embryo histone octamers (**fig. S37**), yielding comparable results. WT and variant Abf1 proteins were used at 30 nM in **Fig. 6E** and 20 nM in **fig. S37**.

**Data processing for RNA-seq, ODM-seq and MNase-seq assays**

*RNA-seq*: Raw sequence reads were preprocessed with a Snakemake pipeline that involved several steps, including quality control, alignment of reads to the *S. cerevisiae* SacCer3 (R64-1-1 build) genome using the STAR aligner (*12*), and quantification of gene expression via RSEM(*13*). Rule-based directives in Snakemake made it easier to perform key preprocessing steps, ensuring effective management of computational resources and data dependencies. The processed data were imported into R Studio, where a summarized experiment object was created to encapsulate gene expression metrics alongside sample metadata. This structured data object made robust data manipulation and subset selection in R easier. Prior to creating plots, the transcripts per million (TPM) data were log2 transformed to normalize expression levels, improving the statistical reliability of comparisons. The ggplot2 (*14*, *15*) package was utilized to generate boxplots that illustrate the differences in gene expression between the WT Abf1 and the variant conditions.

*Occupancy measurement via DNA methylation and high-throughput sequencing* (*ODM-seq):* Both natural and modified bases were called for Oxford Nanopore sequencing reads, and reads were filtered and demultiplexed with Dorado (Oxford Nanopore Technologies (ONT) Basecalling Software, (C) Oxford Nanopore Technologies plc. Version 7.2.13+fba8e8925, client-server API version 16.0.0). The resulting reads were mapped to the S. cerevisiae genome (SacCer3) with minimap2 (version 2.17). Methylation information was added to the bam files using modkit repair, and methylation probabilities were extracted from bam files using modkit pileup (ONT, version 0.1.13). The modkit output corresponds to the averaged methylation probabilities over all reads per genomic coordinate per sample. The genome average methylation was calculated for each sample by averaging the modkit output over all coordinates (**fig. S38**). As slight batch effects were visible on this level of genome average methylation, we introduced a correction factor. The genome average methylation of each of the four individual samples (180 and 240 min of replicates 1 and 2) for each Abf1 variant (WT Abf1, no Abf1, FUS$^{1\text{-}163}$12E, Local Shuffle 12, Altered Aromatic Patterning) was averaged. The resulting averages were divided by the average of the four WT Abf1 samples, and this quotient was used as a correction factor for each individual sample. These corrected modkit outputs were plotted in composite plots (ggplot2) relative to alignment points. Genomic coordinates were converted to new coordinates relative to the alignment points, which received the coordinate zero in each case (data.table). Averages per sample were calculated for every new coordinate, and plots were smoothed by plotting the rolling mean over a 50 bp window at the center base pair of each window. In vivo +1 nucleosome and transcription start site (TSS) annotations were taken from Oberbeckmann et al. (*16*).

The annotation of responder Abf1 sites was either taken from Kubik et al.(*17*) or derived from our ODM-seq data in the following way. Responder Abf1 sites were determined by starting from 2126 Abf1 sites predicted by position weight matrices (PWMs) obtained from either Pachkov et al. or MacIsaac et al. (*18*, *19*). If predicted binding sites were within 20 bp of each other, only the site with the lower genomic coordinate was used. As an alignment point, we used the center base pair (rounded up for even number of bp) of each PWM. We defined five windows for calling occupancy differences of Abf1 variants versus WT Abf1, either at the Abf1 sites or in the proximal or distal flank of the +1 and -1 nucleosomes (**fig. S28**): one ± 70 bp window is centered at the Abf1 site (0 bp viewpoint), or two ± 40 bp windows are centered at ± 100 bp in both up- and down-stream direction from the Abf1 site (100 bp viewpoints) or at ± 180 bp (180 bp viewpoints) (**fig. S26**). For each individual sample (180 and 240 min of replicates 1 and 2 for each Abf1 variant), we calculated the average methylation probability within every of the five windows based on the corrected modkit output. The averages of the five windows for four individual samples for no Abf1, $FUS^{1\text{-}163}$12E, Local Shuffle 12, and altered aromatic patterning were compared per window to the respective window averages for four WT Abf1 samples with a two-sided Student's t-test (stats). Abf1 binding sites were called responder sites per window if $p \leq 0.01$ was in that window (**Table S8**). A subgroup of the Abf1 responder sites was called if the sites were within 80-180 bp of a TSS. For composite plots of ODM-seq data at such defined Abf1 responder sites, the data for 0 viewpoint windows were oriented such that the nearest TSS, if present within 1000 bp, was to the right of the Abf1 sites or the PWM was set to the top strand. For the mirrored ± 100 bp and ± 180 bp viewpoint windows, the window with the smaller p-value was oriented to the right. Composite plots were only done if responder site counts exceeded 40. For comparison of our called Abf1 responder sites with responder sites defined by Kubik et al. (*17*), sites were considered the same if their genomic coordinates were within 50 bp of each other.

*NB*: we recovered many more responder sites than previously identified, and also some sites identified by Kubik *et al.* were not recovered in our analysis (**fig. S29**). The larger number of responder sites may be explained because we did not limit our search to Abf1 sites in promoters that were confirmed by ChIP data, but included in an unbiased way all 2126 sites predicted by position weight matrices (PWMs; (*18*, *19*)). Conversely, the responder sites missed by us relative to the responder sites called by Kubik *et al.* may be due to our stringent significance threshold of $p \leqslant 0.01$.

*Selection of sites in Fig. 6*. For analysis in Figure 6, subsets of Abf1 responder sites were selected for ODM-seq composite plot construction. These sites were selected based on the following criterion:

**Panel B:** Genes (n = 279) linked to Abf1 responder sites, where responder sites were identified usingMNase-seq of WT vs. no Abf1 or as Class I and II as defined by Kubik et al. (*17*). See also fig. S24.

**Panel D**: Sites (n = 356) defined as all Abf1 sites where the "no Abf1" strain showed significantly more occupancy in a 141 bp window centered on the Abf1 site.

**Panel E**: Sites (n = 1025) in the yeast plasmid library where nucleosome organization was affected *in vivo* (as assessed by ODM-seq) in the "no Abf1" anchor away strain.

**Panel F**: Sites (n = 112) where the $FUS^{1\text{-}163}$12E strain showed significantly more occupancy in at least one 81 bp window centered at ± 100 bp distance to the Abf1 site.

**Panel G**: Sites (n = 148)  swhere the "Altered Aromatic Patterning" strain showed significantly higher occupancy in a 141 bp window centered on the Abf1 site.

Sites for RNA-seq MNase-seq of WT vs. no Abf1, Class I and II) of Kubik et al., 2018 Mol Cell, fig. S24

*MNase-seq*: Illumina sequencing data were mapped to *S. cerevisiae* SacCer3 (R64-1-1 build) genome using Bowtie2(*20*), excluding reads with multiple matches. The bam files were converted to bed files and then to RDS files of ranges (start to end), which were imported into R Studio. For comparison samples with different total read numbers, we applied subsampling. To mainly include DNA fragments generated by MNase digest of canonical nucleosome core particles, we selected only 120-170 bp reads. For visualization of the MNase-seq patterns, we plotted 50 bp extended dyads. Regions of > 200 bp contiguous nulls (no coverage) were omitted among all compared samples for a particular graph.

As a final note, we also showed that INO80 cannot generate nucleosomal arrays phased to Abf1 sites without Abf1 (**fig. S36**).

**Proteome-wide analysis**

Motivated by previous work, the set of aligned syntenic genes from twenty different yeast species was obtained from the Yeast Genome Order Browser as described previously (*21*–*25*). This dataset contains 5430 protein sequences. Orthologous sets of sequences were then aligned using Clustal omega to generate alignments for every *S. cerevisiae* protein (*26*). Per-residue sequence conservation was calculated using the normalized Jensen-Shannon (JS) divergence based on the BLOSUM62 matrix, as first introduced by Capra and Singh (*27*–*29*). This analysis yields a per-residue conservation score where values range between 0.019 and 0.948 across the yeast proteome (**Fig. 1**).

Disorder scores and predicted pLDDT scores for analyses were calculated using metapredict (V1), where defined IDRs were identified using the IDR delineation mode (*30*). The predicted pLDDT value reflects a prediction of the per-residue pLDDT score (predicted Local Distance Difference Test), a metric introduced by DeepMind in the context of structural evaluation for AlphaFold2. Metapredict implements a deep learning model to predict the pLDDT score from sequence alone.

Almost all disorder predictions in this work used metapredict V1, instead of the more accurate metapredict V2 or V3. This was a deliberate decision driven by the underlying training data used for metapredict V2 and V3. Metapredict V1 was trained in a manner that avoids convolving evolutionary information inferred from AlphaFold, whereas V2 and V3 combine AlphaFold-derived information with a set of additional predictions, meaning it (indirectly) has evolutionary information encoded given AlphaFold's ability to confidently predict structure is in part determined by the availability of large sets of alignable homologous sequences. Using the V1 predictions allows us to legitimately compare disorder and conservation without inadvertently capturing trends encoded in the underlying bidirectional recurrent neural networks. The one exception for this is examining IDRs that are conserved in terms of chemical specificity but not sequence specificity (**Fig. 5**). For this analysis, we used metapredict V3 first to identify IDRs across the yeast proteome, and then followed up using these IDRs. We made this decision given the accuracy metapredict V3 has for correctly identifying large contiguous IDRs. At the time of writing, metapredict V3 was not published but was publicly available in a branch of metapredict.

pLDDT-defined structured domains are set using a predicted pLDDT threshold of 65 or greater; gaps between contiguous stretches of predicted pLDDT values over 65 that are three residues or smaller are filled in. A minimum region size of five residues is set for convenience.

**Sequence analysis**

All sequence analysis was performed using SHEPHARD (https://shephard.readthedocs.io/). The complete set of sequences, conservation scores, disorder scores, ppLDDT scores, and IDR domain boundaries are provided as SHEPHARD-compatible data files (structured, tab-separated data files). IDR sequence analysis (compositional analysis, hydrophobicity, predicted disorder, and predicted pLDDT scores) was performed using localCIDER and metapredict (*30*–*32*).

**Sequential sequence shuffling**

IDR designs were generated using GOOSE, a Python package for the rational design of IDRs and IDR variants. Documentation for GOOSE is available at https://goose.readthedocs.io/ and the open-source code is available at https://github.com/idptools/goose.

**Identification of conserved subsequences in the yeast proteome**

Conserved subsequences were identified via the following procedure. First, hydrophobic (I/L/V/M/Y/F/W) residues in an IDR with a conservation score above 0.65 or higher were identified. These residues are in the 85$^{th}$ (or higher) percentile for conservation across all disordered regions (dashed lines, **fig. S4**). For each conserved hydrophobic residue (initiator methionine residues are excluded) in an IDR, we identified the residues +/– 6 of that central conserved IDR (truncating at the N- and C-termini as necessary). We then calculated the median disorder for this subregion of (up to) 13 amino acids. Finally, having done this for an entire protein, we extract contiguous subregions where the median disorder score is above some threshold. A lower disorder threshold finds more regions but runs the risk of identifying residues at the boundaries between disordered and folded domains, which may, in fact be folded. A higher disorder threshold finds fewer regions but benefits from higher confidence that a region identified is found in a bona fide IDR. With the most permissive disorder threshold, we identified 5173 conserved subregions. With the least permissive disorder threshold, we identified 884 subregions. All subsequences, including their position and parent protein, are provided in **Tables S7-S11**.

**Identification of subsequences with predicted transient structure**

Using a deep-learning model trained on ~360,000 protein structures from AlphaFold2, we used the metapredict package to predict per-residue predicted structure across the yeast proteome (*30*). Cross-referencing those scores against predicted disordered regions and excluding subsequences directly adjacent to folded domains, we finally filtered for contiguous subsequences with a predicted pLDDT score above 40, where the subsequence was up to 40 residues long. The complete set of these subsequences, the average disorder score, and the average conservation is provided in **Table S17**.

**Identification of IDRs with compositional conservation**

Compositional conservation is calculated by taking a set of orthologous aligned IDRs and computing the mean Euclidean distance between fractional amino acid composition across all the IDRs. Compositional conservation is only calculated for IDRs of length ten residues or longer (without alignment gaps) and for sequences where ten or more orthologous IDRs are identified. The complete set of IDRs with median and mean sequence conservation and compositional conservation is provided in **Table S16**.

**Hydrophobicity and charge scores**
Hydrophobicity and charge scores (**Fig. 4J**) are calculated as compositional weighted metrics.

The hydrophobicity score (*B*) for a sequence is calculated with the following expression;

$$B = \sum_{i}^{\beta} f_i \beta_i \qquad (1)$$

Where for a given protein *B* is the binding score, *β* is the set of residues used to calculate the binding score (Y, W, F, M, V, I, L, A, P), $\beta_i$ is the binding score coefficient associated with residue identity *i* and $f_i$ is the fraction of the sequence made up of residue *i*.

The charge scores (*C*) are calculated in an analogous way;

$$C = \sum_{i}^{\theta} f_i \theta_i \qquad (2)$$

Where the residues in the set θ are R, K, E, and D, $\theta_i$ is the charge score coefficient, and those coefficients are listed in **Table S16**.

**AlphaFold2 model**
The AlphaFold2 model was taken directly from the EBI AlphaFold2 protein structure prediction database (https://alphafold.ebi.ac.uk/entry/P14164) (*33*, *34*).

**Gene Ontology Analysis**
The gene ontology analysis described in **fig. S7** and for proteins with IDRs that are conserved in terms of chemical specificity but not sequence was performed using PANTHER (*35*).

**All-atom simulations**
All-atom simulations were performed with the CAMPARI Monte Carlo engine and ABSINTH implicit solvent model (*36*, *37*). The combination of ABSINTH and CAMPARI has been effectively used to recapitulate IDR ensembles in a variety of systems (*38–41*). Simulations were performed on a peptide with the sequence “AEVDAQEARETAQLAIDKINSYKRSIDD”. Twenty independent simulations were run, where each simulation was run for 6,000,000 equilibration steps followed by 14,000,000 production steps. Conformations were saved every 20,000 frames, during the production stage, such that the final conformational ensemble is made up of 15,000 conformations.

Simulations were run at 320 K with 10 mM NaCl, using ABSINTH 3.2 OPLS with the ion parameters of Mao and Pappu, as previously done (*38*, *40*, *42*).

**Coarse-grained simulations**

Coarse-grained simulations were performed using the PIMMS simulation engine (https://github.com/holehouse-lab/pimms) to generate the two-dimensional binding landscapes. A fifty-bead flexible polymer is defined, in which the central six residues are designated as a motif while the flanking residues on either side are defined as the context. A second fifty-bead polymer that collapses due to attractive intramolecular interactions defines a binding partner. Motif:globule interactions (M) and context:globule (C) interactions were systematically titrated to generate a two-dimensional sweep of parameter space (see **fig. S19**). The fraction bound is calculated for each simulation run at a unique combination of M and C interaction strengths. Binding here is defined as the two molecules in direct interaction and does not distinguish between motif-based or context-based binding.

Monte Carlo simulations were run on a cubic lattice of 30 x 30 x 30 sites. The system is evolved via a combination of chain repetition and rigid-body translation moves. Each simulation is run for a total of on the order of 30 x $10^6$ individual Monte Carlo moves.

***In silico evolution***

In silico evolution (**Fig. 6**) was performed by iteratively acquiring a set of mutations and then evaluating the fitness of the sequence with respect to some selection function. In our case, mutations were introduced as single point mutations only (no insertions or deletions) and fitness was evaluated in terms of evaluating attractive interactions with a specific binding partner.

Chemical specificity was calculated using FINCHES using the Mpipi-GG forcefield (*43–45*). *In silico* evolution used a fitness function whereby the summed attractive interactions across the predicted intermolecular interaction map (intermap) were computed and mutants “survived” if the summed attractive interactions remained approximately equal to or more attractive than the wildtype value. Mutations were introduced by sequentially making a series mutations before evaluating fitness, enabling compensatory mutations to appear.

Mutations were introduced through an iterative mutational strategy. First, the amino acid sequence was converted into nucleotide sequence. Next, the underlying nucleotide sequence was stochastically mutated based on expected transition/transversion rates for nucleobases found in coding sequences. This is done by randomly selecting a nucleotide and having it undergo a transition or a transversion event. We typically performed 30-40 events per nucleotide sequence to mutagenize the underlying nucleic acid sequence to a consistent number of amino acid changes. Having mutagenized the underling nucleic acid sequence, this sequence is converted back into protein space. If a premature stop codon was introduced, the sequence is discarded, otherwise that sequence is saved as one of a set of possible mutants. This procedure is repeated many times to generate an initial library of variants. Finally, each variant is assessed with respect to the underlying fitness function, and only those variants that are sufficiently “fit” are kept. For null models (evolution absent selection), the same procedure is carried out, except we do not apply a fitness selection filter at the end. The aquisition of all mutations before evaluation by the fitness function facilitates the aquisition of compensatory mutations. It also more accurately mirrors the logistical steps taken for the *in vitro* error-prone PCR experiment.

The described protocol for *in silico* evolution was done to create two libraries of 350 sequences; in one library (real-lib), every sequence was evaluated and kept based on the fitness function, while in the other (null-lib), sequences were randomly mutageneized without constraints on which sequences were kept or discarded. We ensure that the number of mutations observed in sequences in the real-lib and null-lib are always matched, and are typically around 28-31 mutations per sequence (i.e. 12-15% of the residues). However, it is worth noting that the results do not change if we increase or decrease the number of mutations.

To perform statistical comparisons, we performed a bootstraping-like analysis to investigate per-residue conservation for sequences generated for our two libraries (**Fig. 5C**). This involved randomly sampling twenty sequences (without replacement) from either real-lib or null-lib, and for those twenty sequences calculated the per-residue conservation score. This was then repeated ten times to build a distribution of per-residue conservation scores expected for sequences generated under selection vs. sequence generated from the null model. For each residue position, the mean and standard deviation for those distributions are shown in **Fig. 5C**.

We initially compared conservation of IDR2 vs. the N-terminal IDR of Rad7 as our hypothetical target protein. To establish the apparent lack of conservation through alignment-based methods is a general phenomenon and not a fluke, we repeated this analysis using eight additional IDRs from proteins Abf1 has been proposed to interact with. Putative interactors were taken as physical interactors from BioGRD(*46*), although we note that in many cases the interactions were identified in the presence of DNA, such that we cannot distinguish between a scenario in which Abf1 interacts directly with the putative partner, or that both proteins are interacting with the same DNA molecule. Rad16 and Rad7 are notable exceptions where protein-only complexes have been identified (*47*, *48*). Regardless, in silico evolution for the eight additional potential partners leads us to the same conclusion (**Fig. S23**).

For comparing *in silico* evolution vs. the results from error prone PCR experiments (**Fig. 5E**), a similar strategy was taken as described above, except we matched the number of mutations to the average number of mutations seen experimentally (~28 mutations) and we subsampled groups of 14 sequences to mirror the number of viable sequences identified experimentally.

As a final note, our *in silico* evolution is far from a faithful reproduction of the putative dynamics that shape mutations in codoing sequences of proteins. We consider this a toy system to ask a stylized question - if the ONLY determinants of proteins sequence changes were constrained in this way, could conservation be obviously detected by multiple sequence alignments. In reality, partners are co-evolving, post-transcriptional restraints (i.e. RNA secondary structure, processing, and degradation) will also shape protein coding sequences, and insertions and deletions will contribute additional noise. We consider our simplified model be an overly-stringent and deliberately “simple” model for selection - if signatures of conservatoin are not easily detectable here, then the ability to do so in even noisier scenarios seems even less likely.

**Conservation of chemical specificity**

To assess IDRs in which chemical specificity was conserved, we first identified those IDRs with a low degree of average sequence conservation. We used an average sequence conservation value of 0.35. To assess chemical specificity, we computed average IDR interactions with a set of 36 of chemically orthogonal dipeptides identified previously (*45*). For each set of orthologous IDRs, we calculated the variance in mean-field interaction scores with each of those 36 peptides. We then identified those IDRs where – despite very low sequence conservation – the variance among one or more dipeptide interactions was in the bottom 15%. The set of proteins, IDR sequences, and the dipeptides with which they were conserved with respect to are included in **Table S9**.

This analysis focussed explicitly on large IDRs, and is a relatively unsophisticated way to systematically identify signatures of conservation with respect to chemical specificity. This is an area of active and ongoing investigation.

**Flory-Huggins phase diagrams**

To generate hypothetical Flory-Huggins phase diagrams, we solved the Flory-Huggins free energy of mixing, as described previously (*49–51*). This allows us to calculate volume fraction (ϕ) and temperature phase diagrams for different chain lengths (**fig. S31C**). Reduced temperature was calculated by dividing the binodal temperature values by the critical temperature ($T_c$). To convert volume fraction into molar concentration, we operate under the assumption that

$$\varphi = c\left(\frac{v_m N_a}{M_m}\right) \qquad (3)$$

Where ϕ is the volume fraction, $c$ is the mass concentration (in mg/ml), $v_m$ is the average volume occupied by a single amino acid (140 $Å^2$), $N_a$ is Avagadro's number, and $M_m$ is the average mass of an amino acid (110 g/mol). Given three of the four terms here are known, this expression can be re-written as

$$c = \varphi\rho_0 \qquad (3)$$

Where $\rho_0$ is 1310.16 mg/ml. As such, to convert volume fraction to mass concentration (in units of mg/ml) involves multiplying the volume fraction by 1310.16.

To convert from mass concentration into molar concentration involves dividing mass concentration by molar mass of the molecule, which for the short and long synthetic repetitive FUS sequences are 7,208.9 g/mol and 17,422.1 g/mol, respectively. The precise sequences are listed in **Table S1**.

**Fig. S1. Viability of Abf1 constructs assessed by 5-FOA plasmid shuffling assay.** Strains were restreaked from patches from YNB – ura – leu plates to 5-FOA – leu plates. Only constructs discussed in the paper are labeled in plate schemes, where names of viable strains are in blue, names of inviable ones in black. Viability on 5-FOA plates was visually scored by comparison to known viable or inviable strains on the same plate. Inviable strains were distinguished from viable strains because they showed a smeary appearance and at most tiny/sparse single colonies, whereas viable strains grew as one more or less dense patch. *See separate PDF data file.*

**Fig S2. Growth phenotypes by dilution plating for Abf1 constructs.** Unless noted otherwise, the following images show YPDA plates that were incubated for ca. 1 day at 30°C. If one of the independent clones for each construct did not show the same growth rate as the others, it is labeled as “outlier”, and we categorized the growth rate according to the two similarily growing clones. All rating is relative to the growth of the strain with wild type (WT) Abf1 on the same plate. *See separate PDF data file.*

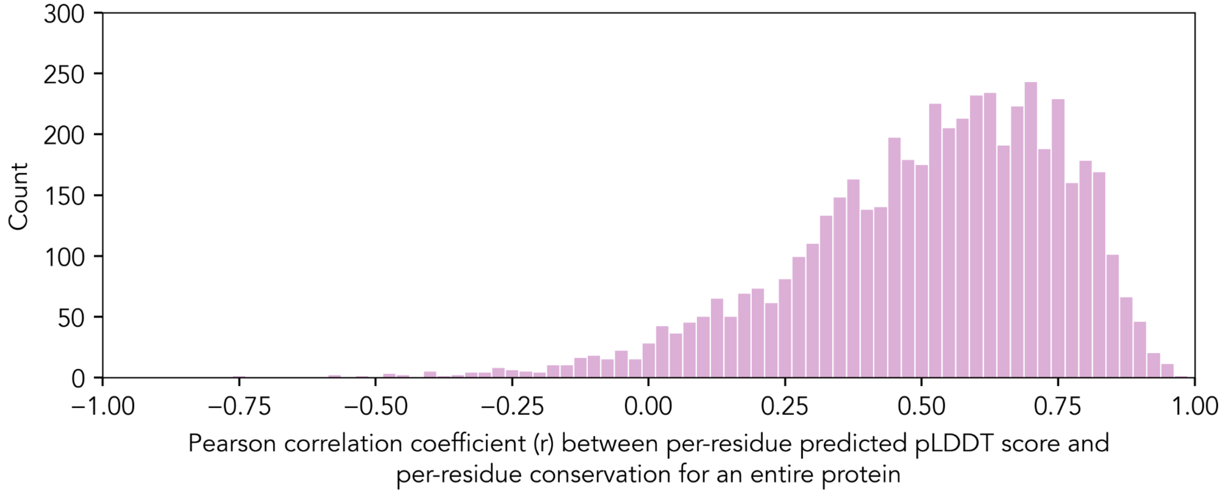


**Fig. S3. Distribution of correlation between per-residue predicted structure and per-residue conservation across the *S. cerevisiae* proteome.** Per-residue predicted structure is calculated using the predicted pLDDT, a metric that reports on the likelihood that AlphaFold2 would be able to correctly predict a structure for the residue in a specific sequence context. This prediction is implemented in metapredict (*30*) and correlated (r) with per-residue conservation.

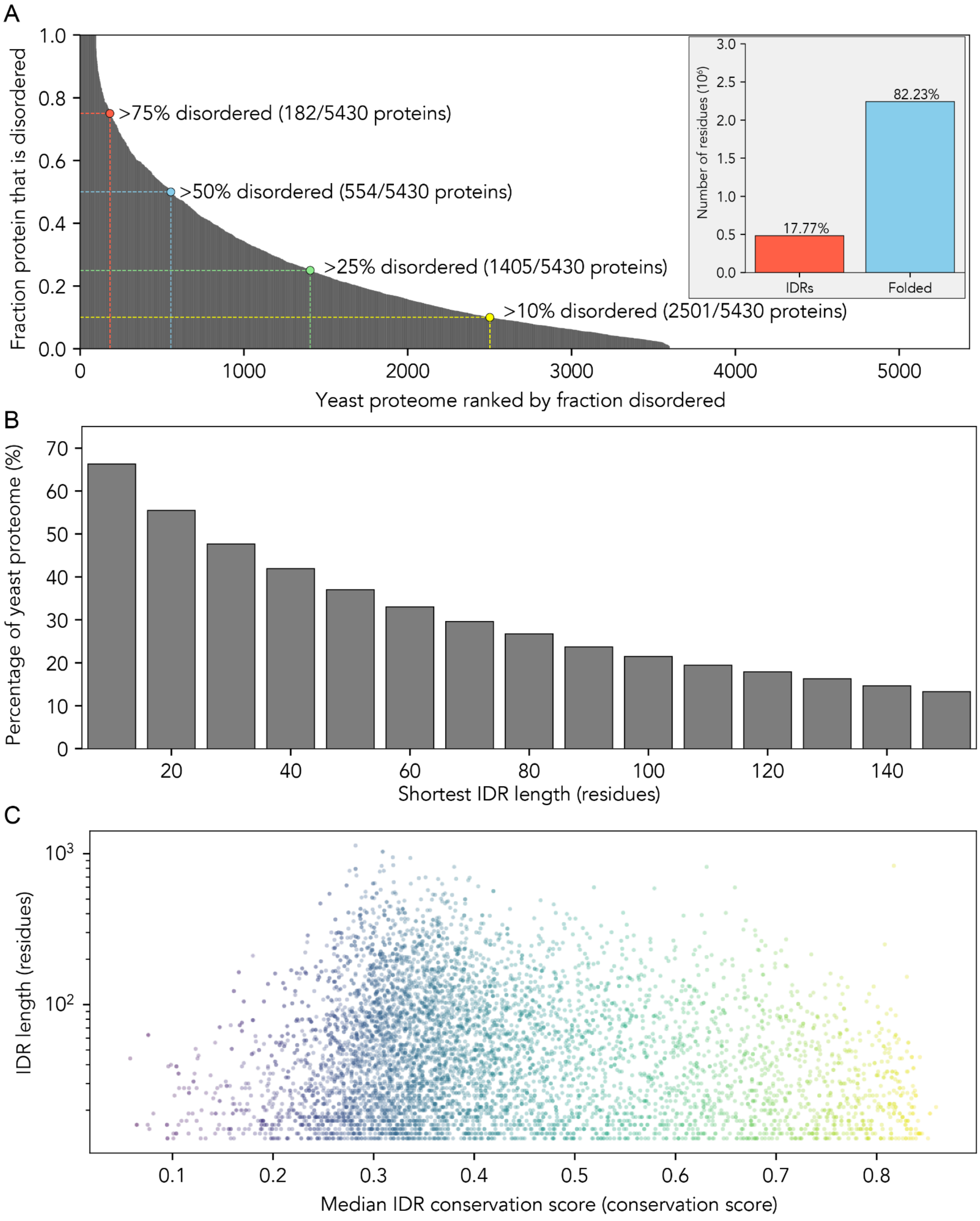


**Fig. S4 Proteome-wide analysis of disordered regions in *S. cerevisiae.* (A)** Rank-ordering every yeast protein by fraction disordered illustrates how prevalent disordered regions are predicted to be in yeast, where almost half of all proteins are 10% disordered or more, while about 18% of all residues reside in IDRs (inset). **(B)** To further convert from fractional disorder into the number of residues, we show the fraction of proteins in the yeast proteome that contains one (or more) IDRs of various lengths. Over 30% of yeast proteins contain one or more IDRs of sixty residues or longer. **(C)** Conservation vs. IDR length reveals that while many IDRs are poorly conserved, there exists a subpopulation of IDRs with a wide range of lengths that are relatively highly conserved. Colors track with median IDR conservation. Gene ontology (GO) analysis reveals that the top 200 most conserved IDR-containing proteins are enriched for nucleosomal proteins, ribosomal proteins, and DNA repair proteins (*35*).

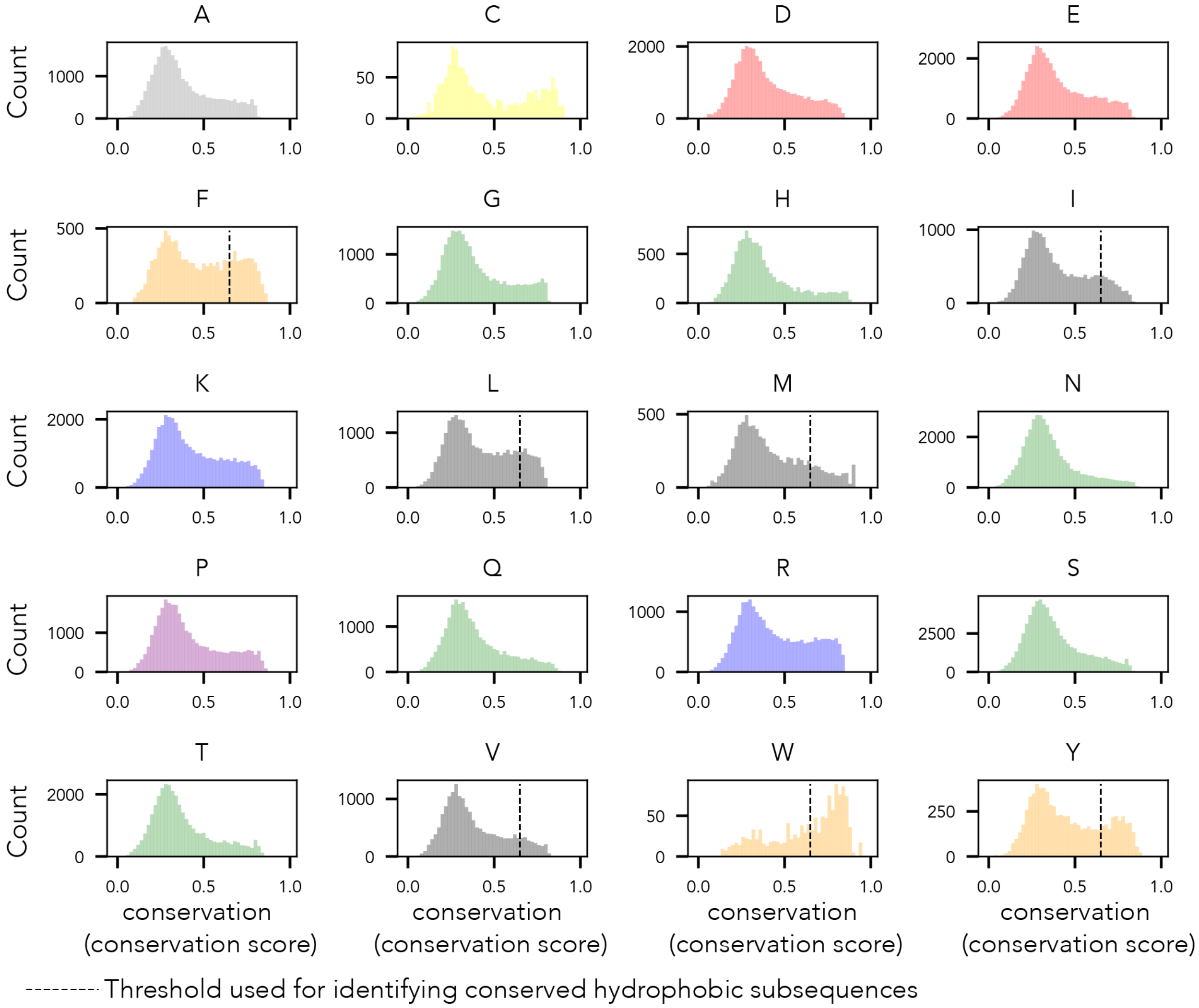


**Fig. S5 Histograms of per-residue conservation for each amino acid type within IDRs.** We calculated the per-residue conservation for each instance of each amino acid, separated out by amino acid type, and histogrammed the values. This analysis revealed that for hydrophobic and aromatic residues especially, while there exists a large population of poorly-conserved residues (conservation scores of 0.3 to 0.4) the second population of highly-conserved residues of these types exists. This is most noticeable for tryptophan (W), tyrosine (Y), and phenylalanine (F). Given short linear motifs are often driven by hydrophobic residues, we defined a threshold confidence score of 0.65 and defined any of the three aromatic (Y/W/F) and four bulky aliphatic (L/I/V/M) residues with a score above this as ‘highly conserved’ (dashed lines). Based on this delineation, we extracted sub-sequences centered on highly conserved hydrophobic residues as possible signatures for binding motifs.

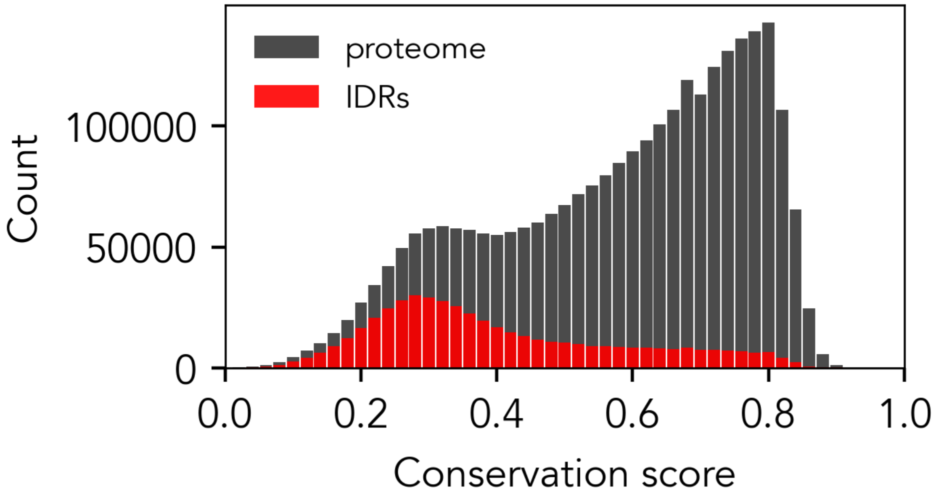


**Fig. S6. Histograms that show the distribution of per-residue conservation scores across the entire proteome and for all IDRs**. The per-residue conservation scores for every residue across the whole yeast proteome (black) or within disordered regions (red) were histogrammed. Disordered regions show a monomodal distribution with a peak around 0.3, while the whole proteome shows a bimodal distribution, with one peak in line with the disordered regions at around 0.3 and a second peak with substantial skew at around 0.75.

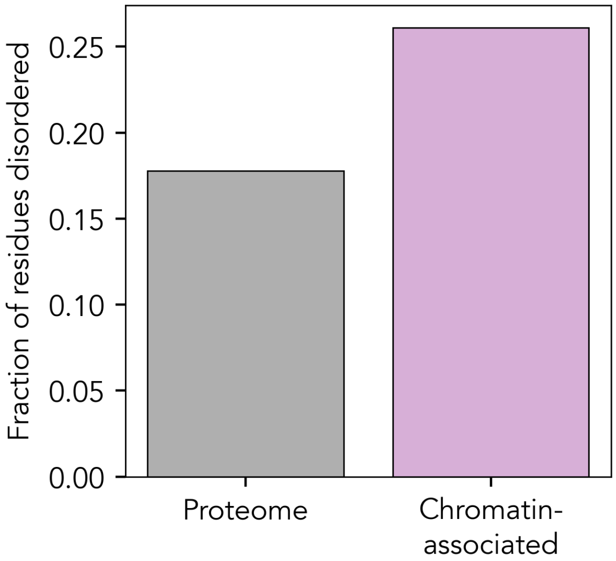


**Fig. S7 Comparison of disordered residues in chromatin-associated proteins compared to all proteins.** Fraction of disordered residues in the entire yeast proteome (n = 5430, left) compared with chromatin-associated proteins (n=190, right). Chromatin association is based on annotation with the GO term "chromatin" (GOA=785). Disordered regions are defined based on metapredict with standard thresholds (*30*).

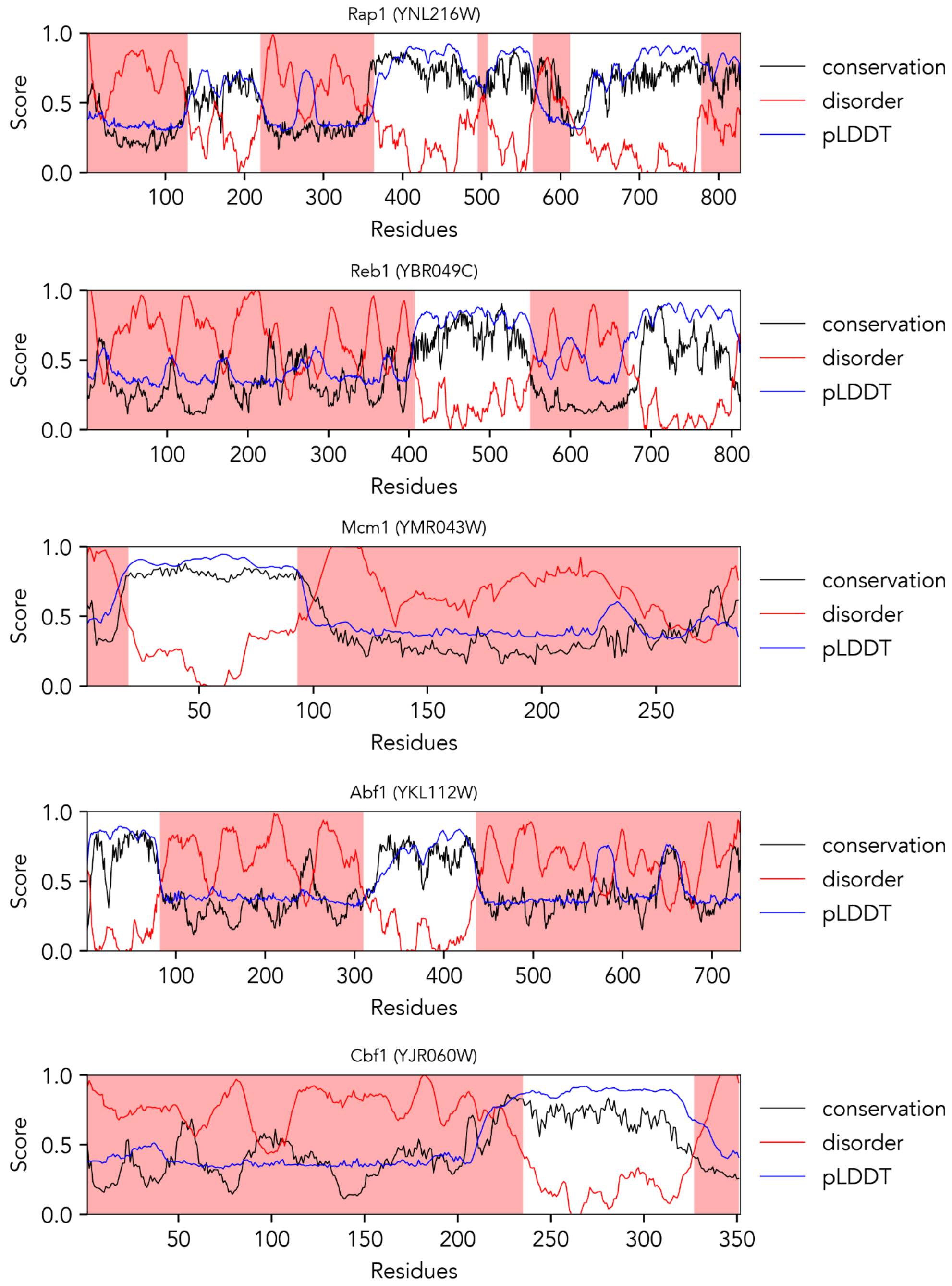


**Fig. S8. Disorder, conservation, and predicted structure profiles for representative general regulatory factor (GRF) proteins**. We analyzed linear sequence profiles for Rap1, Reb1, Mcm1, Abf1, and Cbf1. The shaded red regions are contiguous IDRs, the red line denotes the per-residue disorder score, the blue line per-residue predicted pLDDT score, and the black line per-residue conservation score. Predicted pLDDT score reports on the likelihood of structure (see *Methods*).

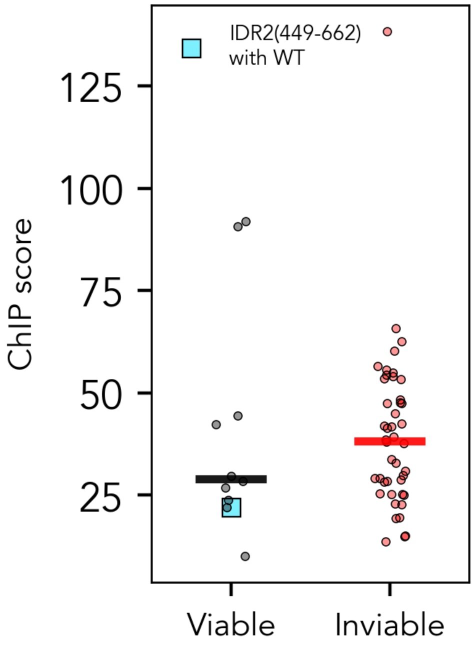


**Fig. S9. ChIP scores for viable vs. inviable constructs confirm the ability of variants to bind Abf1-specific genomic loci even in the event of inviability.** Distribution of ChIP scores as listed for individual viable and inviable constructs were plotted. Horizontal bars mark the median value. Boxed values of viable constructs were measured in the presence of untagged wildtype Abf1 (see *Methods*).

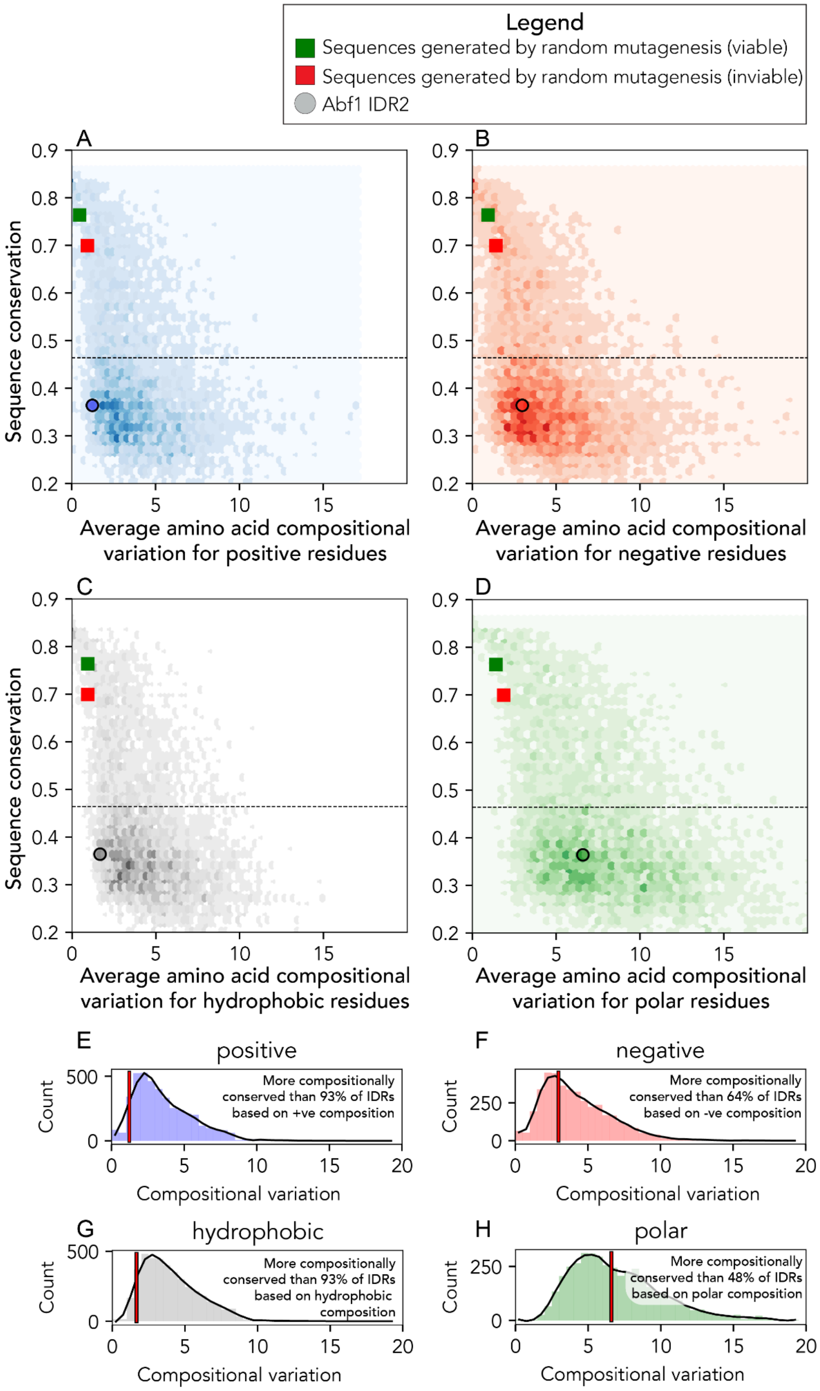


**Fig. S10. Sequence conservation vs. compositional conservation as compared to all other similarly conserved IDRs in the yeast proteome. (A-D)** Panels showing median sequence conservation as assessed by alignment (y-axis for all) compared to the per-residue compositional conservation (x-axis) for four different compositional groups: **(A)** negative (E,D), **(B)** positive (R,K), **(C)** hydrophobic (I,L,V,M,Y,F,W) and **(D)** polar (Q,S,N,H,G) residues. In each panel, we also show the compositional conservation and sequence conservation scores for the set of IDRs generated via the random mutagenesis that was either viable (green square) or inviable (red square). In addition, the position of IDR2 is noted (circles). **(E-H)** All IDRs that fall below the dashed line in panels A-D are histogrammed on compositional conservation, with IDR2 from Abf1 shown as a red horizontal line on the histogram. **(E)** Positive amino acid composition in IDR2 is more well-conserved than 93% of other IDRs. **(F)**Negative amino acid composition in IDR2 is more well-conserved than 64% of other IDRs. **(G)** Hydrophobic amino acid composition in IDR2 is more well-conserved than 93% of other IDRs. **(H)** Polar amino acid composition in IDR2 is more well-conserved than 48% of other IDRs. From this, we naively conclude that charge and hydrophobicity are more constrained in IDR2 than polar amino acid content.

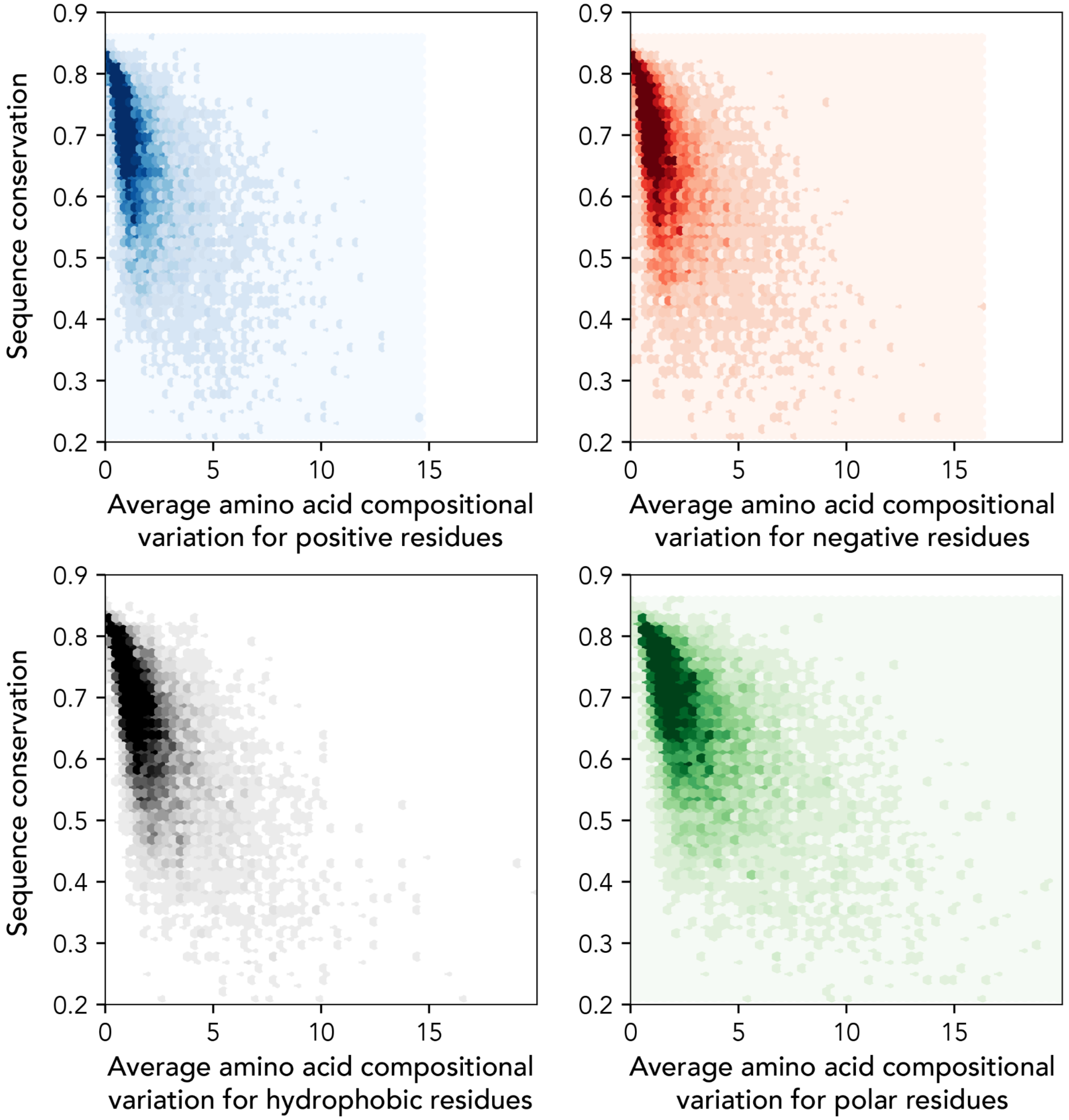


**Fig. S11. Sequence conservation vs. compositional conservation as compared to all other similarly conserved folded domains in the yeast proteome.** Folded domains were identified using predicted pLDDT-based sequence analysis in a manner analogous to identifying disordered regions. Compositional and linear conservation analysis confirms that folded domains tend to be substantially more conserved in both regards.

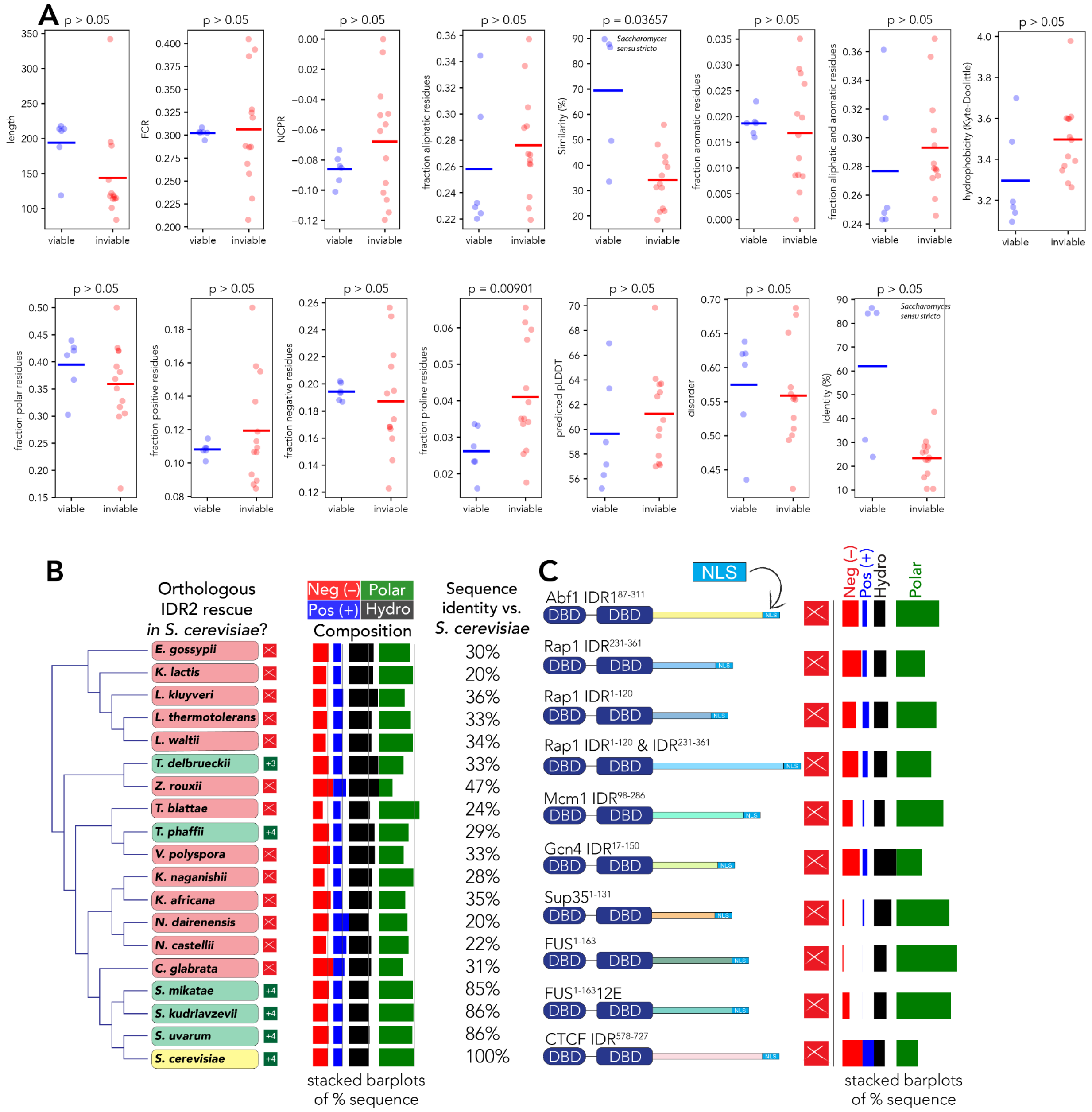


**Fig. S12. Composition analysis across IDR2 orthologs. (A)** Comparison of various sequence features for viable vs. inviable IDR2 orthologs. Across the 12 sequence analyses tested, only one (fraction of proline, bottom right) had a Welch's t-test p-value of below 0.05 when testing for a statistical difference in viable vs. inviable sequences. In addition to sequence features, we computed sequence similarity and identity to wildtype. Including species from *Saccharomyces sensu stricto*, an evolutionarily close species complex which includes *Saccharomyces cerevisiae, Saccharomyces uvarum, Saccharomyces mikatae,* and *Saccharomyces kudriavzevii*, sequence identity is not statistically different between viable and inviable orthologs. While sequence similarity has a p-value of less than 0.05 if the *Saccharomyces sensu stricto* species are included, the viable and inviable sequences are indistinguishable with respect to similarity without this species complex. Sequences were calculated using localCIDER or metapredict (see *Methods*)(*30*, *31*). **(B)** Evolutionary tree (as in Fig. 2D) showing amino acid



Version 2.1 (October 2024)

composition for each construct as a stacked bar chart. Grey lines reflect composition of *S. cerevisiae* IDR2. **(C)** Comparison of tested IDRs from other proteins with amino acid composition for each construct shown as a stacked bar chart.

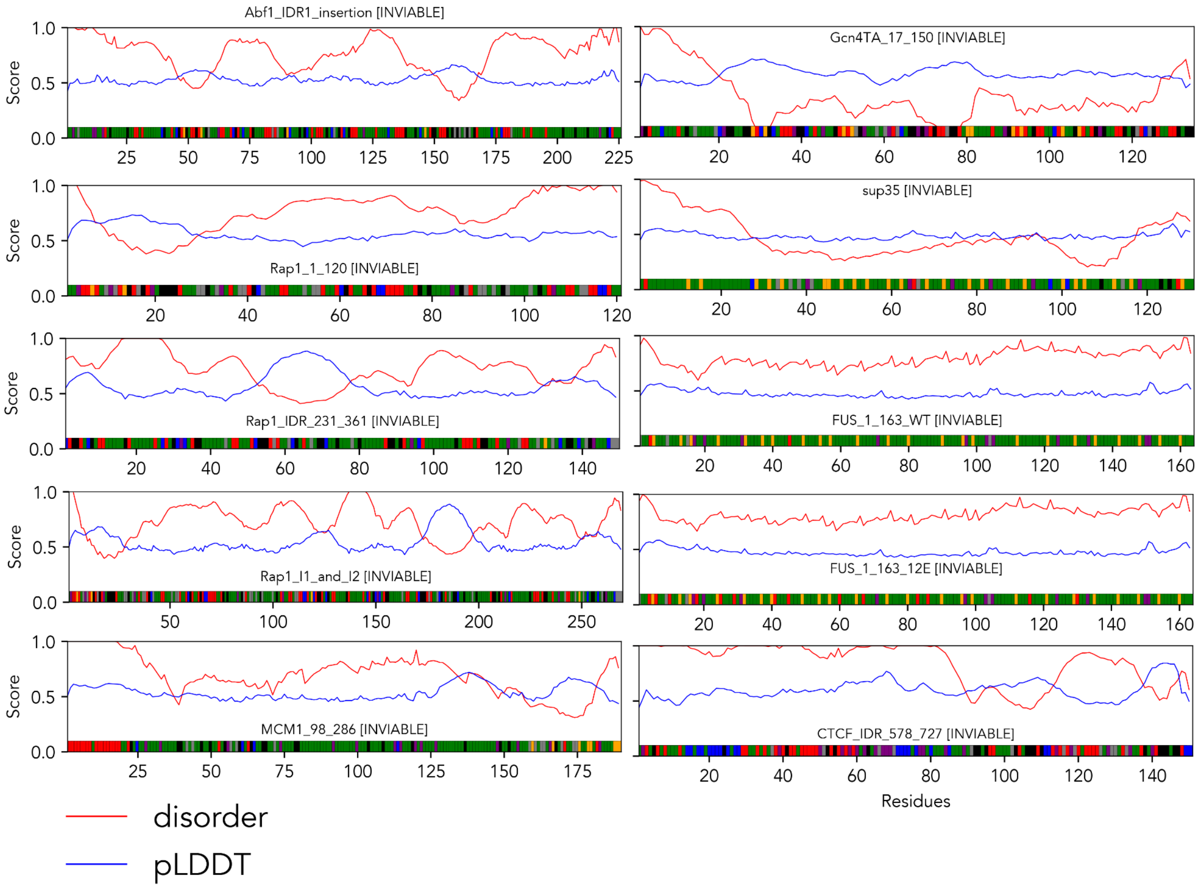


**Fig. S13. Linear sequence analysis of inviable IDRs.** We analyzed linear sequence profiles for IDRs taken from the yeast proteins Abf1, Gcn4, Rap1, Sup35, and Mcm1. In addition, we examined IDRs taken from the human proteins FUS and CTCF. The per-residue disorder score (red) and predicted pLDDT score (blue) provide linear sequence descriptions. The amino acid composition is shown based on amino acid chemistry along the bottom of each panel. Acidic residues are red, basic residues are blue, polar residues are green, aliphatic hydrophobic residues are black, a

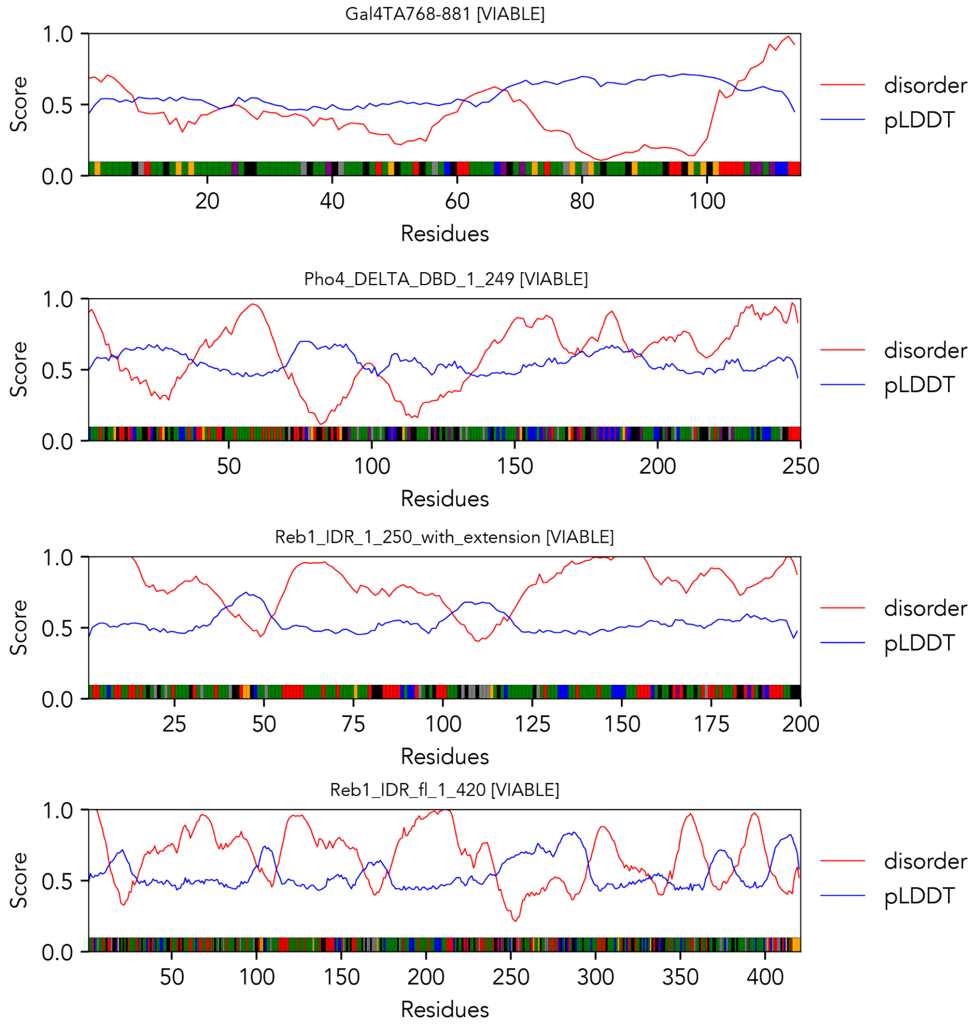


**Fig. S14 Sequence profiles for viable IDRs**. Linear sequence analysis for Gal4, Pho4, and Reb1. The analysis here is analogous to **fig. S13**. Viable sequences vary in length from 114 residues (top) to over 400 residues (bottom), are largely predicted to be disordered, and contain a similar fraction of acidic, polar, and hydrophobic residues compared to Abf1-IDR2.

```
Wt        NNNNNNNDGELSGTNLRSNSID------Y----AKHQEISSAGTSSNTTKNVNNNKNDSND
50 gal4ta -ANFNQSGNIADSSLSFTFTNSSNGPNLITTQTNSQALSQPIASSNVHDNFMNNEITASK
59          * *:.*:::.:.*  .  :            :: * :*.  :***. .*. **: :..

wt         DNNGNNNNDASNLMESVLDKTSSHRYQPKKMPSVNKWSKPDQITHSDVSMVGLDESNDGG
110 gal4ta IDDGN-----------------------NSKPLSPGWTDQTAYNAF-----GITTGM---
88          ::**                       :. *    *:.    .       *:  .

                 EM              Abf1G4
wt         NENVHPTLAEVDAQEARETAQLAIDKINSYKRSIDDKNGDGHNNSSRNVVDENLINDMDS
170 gal4ta ------------------FNTTTMDDVYNY--LFDDE---------------------DT
107                              ::*.: .*   :**:                     *:
                                 Gal4G4

wt         EDAHKSKRQHLSDITLEERNEDDKLPHEVAEQLRLLSSHLKEVE
214 gal4ta P-------------------------PNPKKE-------------
                                     *:   *
```

**Fig. S15 Alignment of Abf1-IDR2 (wt) and Gal4$^{768\text{-}881}$ (gal4ta).** The essential motif (EM), Abf1$^{G4}$ motif, and Gal4$^{G4}$ motifs are highlighted. While other regions in the alignment are also reasonable (i.e. the N-terminus of both proteins) our goal here was to identify if any regions in Gal4 aligned specifically with the essential motif. The alignment was performed using EMBOSS Needle (*52*).

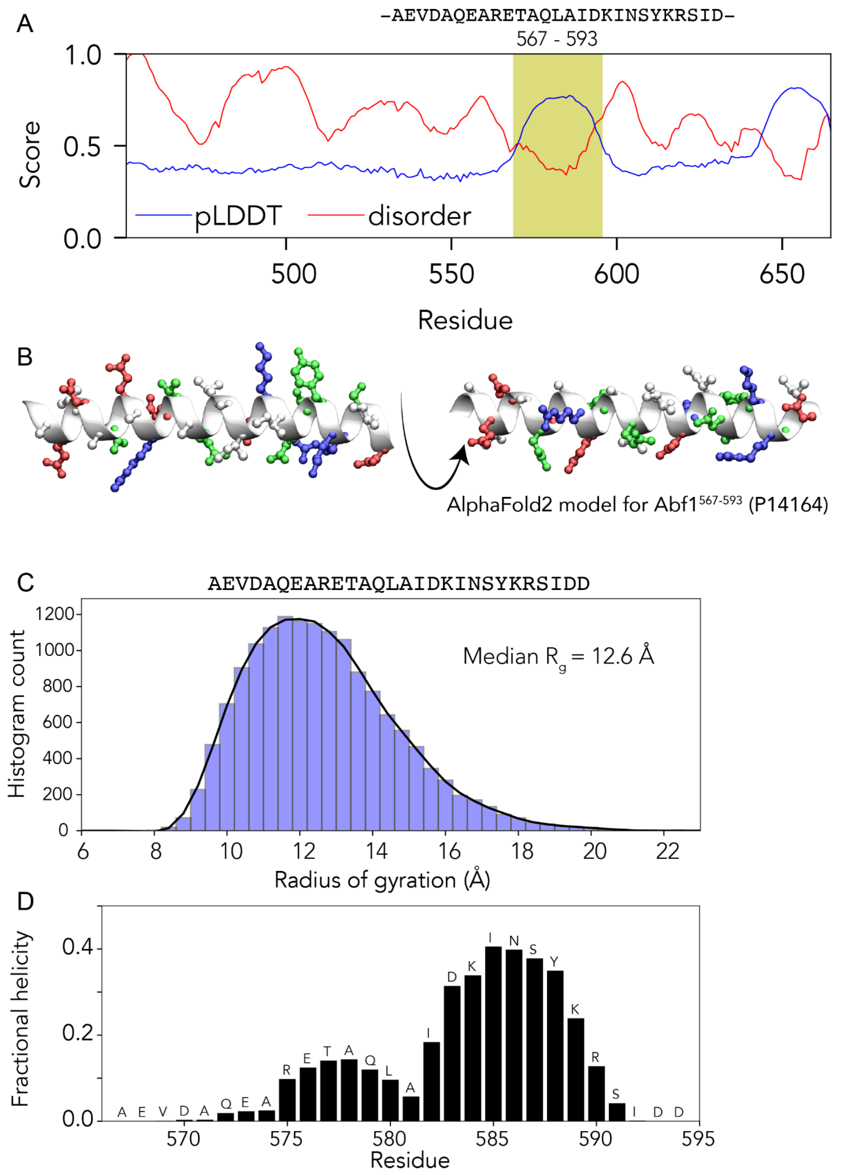


**Fig. S16. The essential motif is predicted to form a transient helix.** (**A**, **B**) AlphaFold2 predicts the essential motif overlaps with a helical region within IDR2. **(C)** All-atom simulations of the essential motif region in isolation predict a disordered region, as illustrated by the radius of gyration distribution which reports on a relatively expanded and broad conformational ensemble, consistent with IDRs studied *in vitro* (*53*, *54*). **(D)** Analysis of local secondary structure reveals that despite being disordered, residues 575-591 are predicted to exist in a heterogeneous ensemble in which a helical region flickers into and out of existence. These simulations provide a rational explanation for how this region can be both disordered and helical.

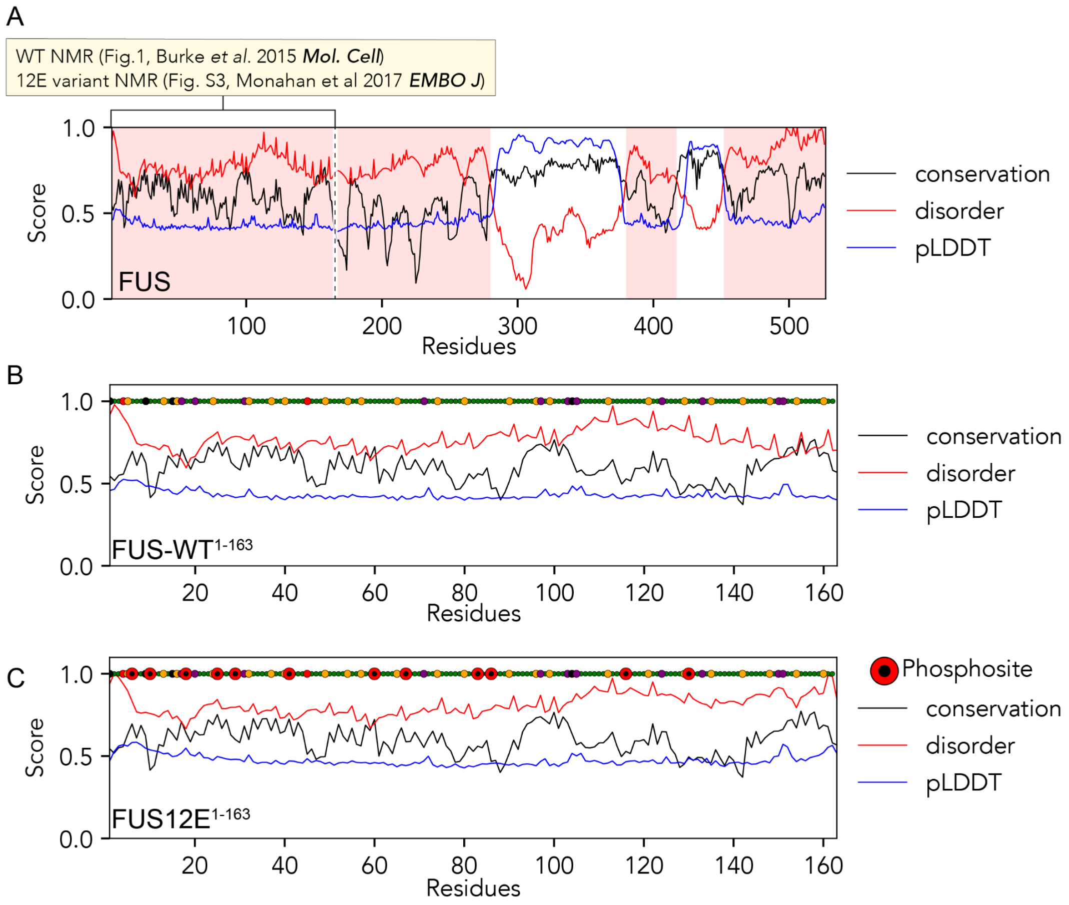


D

MASNDYEQQAEQSYGAYPEQPGQGYEQQSEQPYGQQSYSGYEQSTDTSGY
GQSSYSSYGQEQNTGYGEQSTPQGYGSTGGYGSEQSEQSSYGQQSSYPGY
GQQPAPSSTSGSYGSSEQSSSYGQPQSGSYEQQPSYGGQQQSYGQQQSYN
PPQGYGQQNQYNS

**Fig. S17 FUS$^{1\text{-}163}$ and FUS12E$^{1\text{-}163}$ are low complexity disordered regions, as characterized by both experimental and computational approaches. (A)** Full-length protein with IDRs highlighted in red (identified using both disorder scores and pLDDT scores). Residues 1-163 are relatively poorly conserved, and based on NMR data collected on both WT and the 12E phosphomimetic version are predicted to be fully disordered (references and associated figures shown in inset). **(B)** FUS$^{1\text{-}163}$ WT sequence and **(C)** FUS12E$^{1\text{-}163}$ regions zoomed in with residues from different physicochemical groups illustrated by color. Green = polar (S/T/G/Q/N/H/C), red is acidic (E/D), blue is basic (R/K), orange is aromatic (Y/W/F), black is aliphatic (A/I/L/V/M) and purple is proline. **(D)** Complete FUS12E$^{1\text{-}163}$ sequence.

A

| | | FCR | NCPR | Hydro.[a] | Aro |
|---|---|---|---|---|---|
| Gal4$^{G4}$ | FNTTTMDDVYNYLFDDE | 0.294 | -0.294 | 3.694 | 4 |
| Gal4$^{M2}$ | FMPRDALHGFDWSEEDD | 0.412 | -0.294 | 3.365 | 3 |
| p65$^{M1}$ | DEDFSSIADMDFSALLS | 0.294 | -0.294 | 4.647 | 2 |
| GR$^{M1}$ | VSLLEVIEPEVLYAGYD | 0.235 | -0.235 | 5.141 | 2 |
| VP16$^{M1}$ | GEDVAMAHADALDDFDL | 0.353 | -0.353 | 4.447 | 1 |
| GCN4$^{M1}$ | ESFFSSSTDSTPMFEY | 0.188 | -0.188 | 3.969 | 4 |

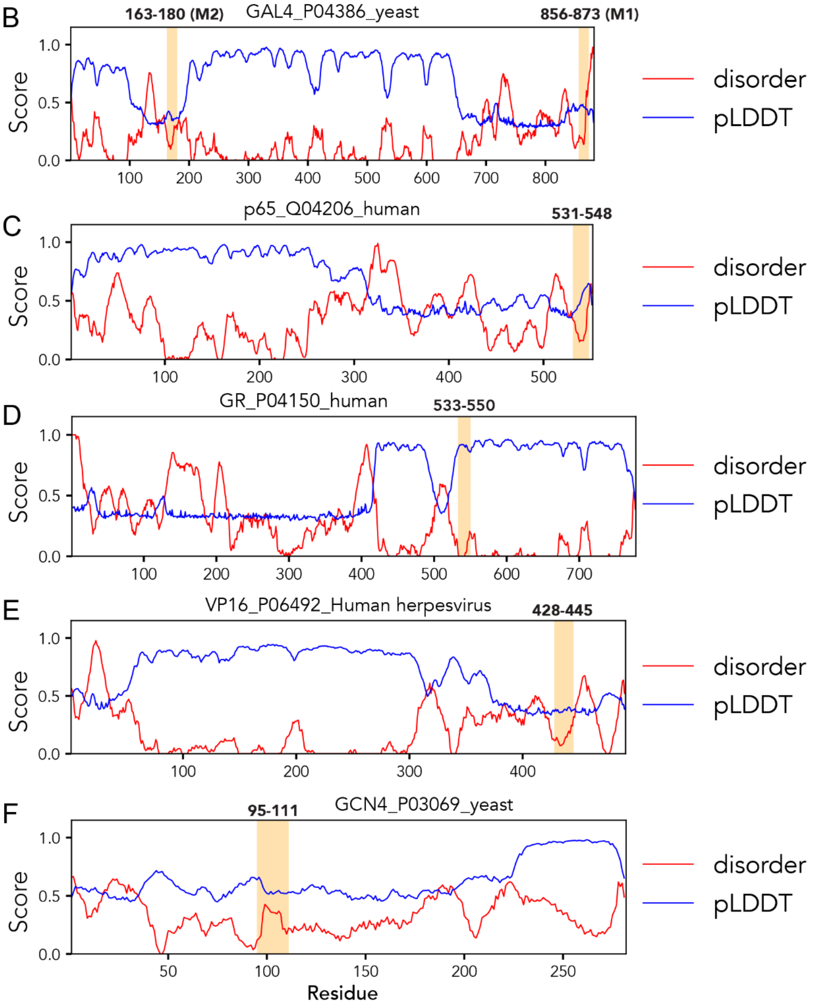


**Fig. S18 Compositionally-selected 16-17 residue subsequences from unrelated proteins**. Subsequences were selected from Gal4, p65, GR, VP16, and Gcn4. **(A)** Subsequences were selected solely based on composition and size. For each protein, the gene name, UniProt ID, and species of origin are provided. **(B-F)** Relative positions of subsequences within their associated full-length proteins, along with per-residue disorder scores and per-residue predicted pLDDT scores. Given selection was based solely on composition within a sliding window, the motif from GR falls within a folded region, which we retained as a convenient control given this region cannot possibly have evolved to function as a binding motif in a disordered protein region.

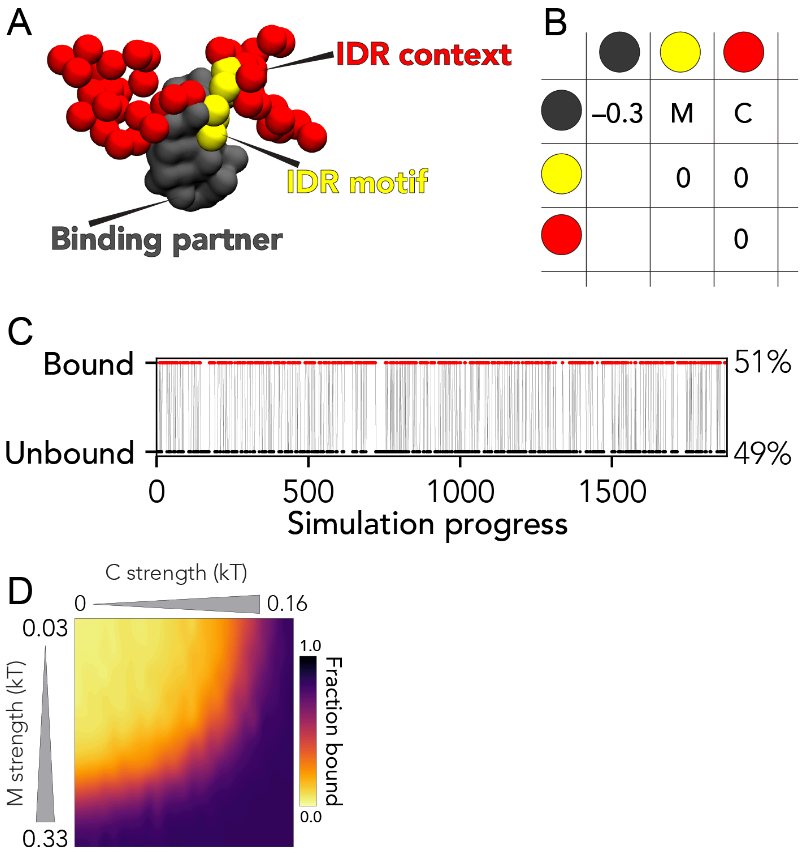


**Fig. S19 Summary of coarse-grained simulations for constructing the naive two-dimensional binding landscape**. **(A)** Coarse-grained simulations make use of a simplified representation in which an IDR consists of a central binding motif (yellow) with longer N- and C-terminal flanking residues (red). This species binds to a partner represented as a globular polypeptide, shown in gray. **(B)** Interaction strength for inter-bead interactions. "M" defines an interaction strength between the IDR motif and globular binding partner. "C" defines an interaction between the IDR context of the globular binding partner. Globular intra-bead interactions are strong, whereas motif:globular protein ("M") and context:globular protein ("C") binding strength are systematically varied. **(C)** Example of single simulation trajectory, revealing an almost 50:50 split of bound and unbound. Simulation progress here is measured in terms of Monte Carlo steps. **(D)** Reproduction of **Fig. 4B/D** - the two-dimension landscape is generated by systematically varying both C and M parameters as defined above.

```
>FUS12E_normal
MASNDYEQQAEQSYGAYPEQPGQGYEQQSEQPYGQQSYSGYEQSTDTSGYGQSSYSSYGQEQNTGYGEQSTPQGYGSTGGYG
SEQSEQSSYGQQFNTTTMDDVYNYLFDDEGSYGSSEQSSSYGQPQSGSYEQQPSYGGQQQSYGQQQSYNPPQGYGQQNQYNS
```

G4 motif

```
>FUS12E_patchy
MASNDYYYYEQQAEQSGAPEQPGQGEQQSEQPYYYYGQQSSGEQSTDTSGGQSSYYYYSSGQEQNTGGEQSTPQGGSTGGYYY
YGSEQSEQSSGQQFNTTTMDDVYNYLFDDEGSGSSEQSSSGQPQSGSYYYYEQQPSGGQQQSGQQQSNPPQGYYYYGQQNQNS
```

G4 motif

**Fig. S20. Aromatic clustered FUS12E.** The top sequence shows the “normal” FUS12E sequence with the G4 motif, whereas the bottom sequence shows the patchy variant. Compositionally these two sequences are identical, but the patterning of aromatic residues is very different. Specifically, using the inverse-weighted distance (IWD) to compute residue clustering, the aromatic clustering of the wildtype sequence has a value of 0.82, while the aromatic IWD clustering in the patchy sequence is 2.49 (*55*).

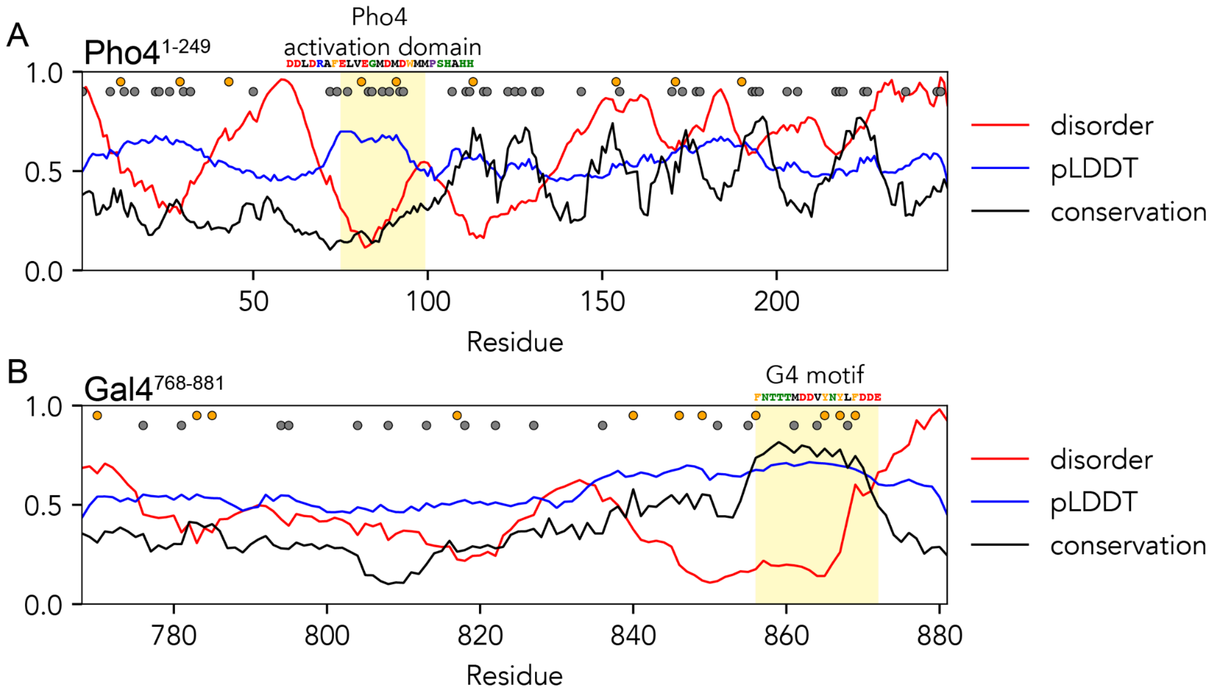


**Fig. S21 Outliers in Fig. 4J may have cryptic motif-like sequence features**. Sequence analysis for the two viable outlier sequences that lack the essential motif in **Fig. 4J**. In each analysis disorder (red), pLDDT (blue) and conservation (black) are shown as lines, while alphatic and aromatic positions are shown as black or orange circles in the associated location, respectively. **(A)** Analysis for Pho4$^{1\text{-}249}$ reveals a series of conserved short regions from residue 100 to 230. In addition, the predicted pLDDT score identifies a hydrophobic-enriched region that corresponds with a known transactivation domain at residues 70-100. Intriguingly, the per-residue conservation of the transactivation domain is relatively low, illustrating that functionally important subregions need not be conserved across orthologs, mirroring results obtained in Abf1. **(B)** Analysis of Gal4 reveals a local set of more conserved hydrophobic residues N-terminal of the conserved G4 motif. Intriguingly, while the G4 motif was originally identified based on homology to Abf1-IDR (**fig. S15**), this region is also the most well-conserved region in Gal4$^{768\text{-}881}$.

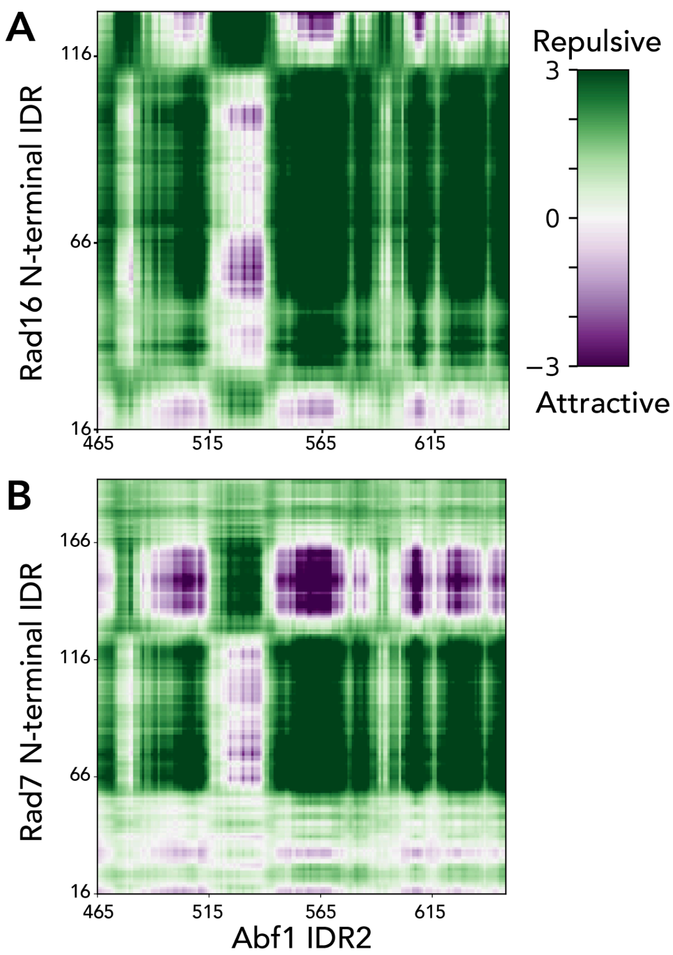


**Fig. S22 Intermaps between IDR2 and Rad16 and Rad7.** Predicted intermolecular interaction maps (intermaps) generated using FINCHES predict specific subregions in Rad7 (UniProt P06779, 1-207) and Rad16 (UniProt P31244, 1-143) that will interact with IDR2. Intermaps can be reproduced using these sequences at [http://finches-online.com/](http://finches-online.com/).

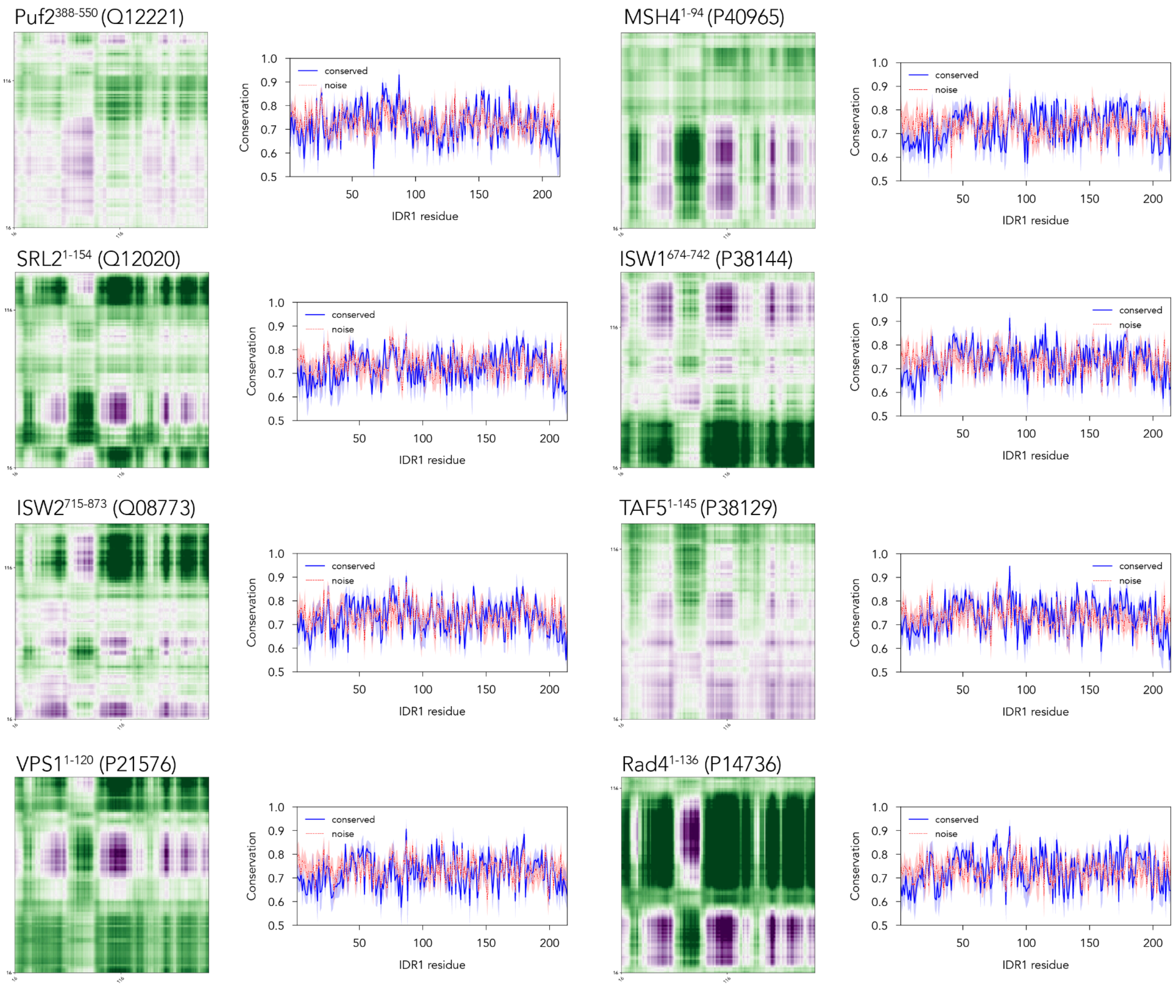


**Fig. S23. IDR2 conservation against a diverse array of possible binding partners yields per-residue conservation scores indistinguishable from noise despite robust conservational pressures.**

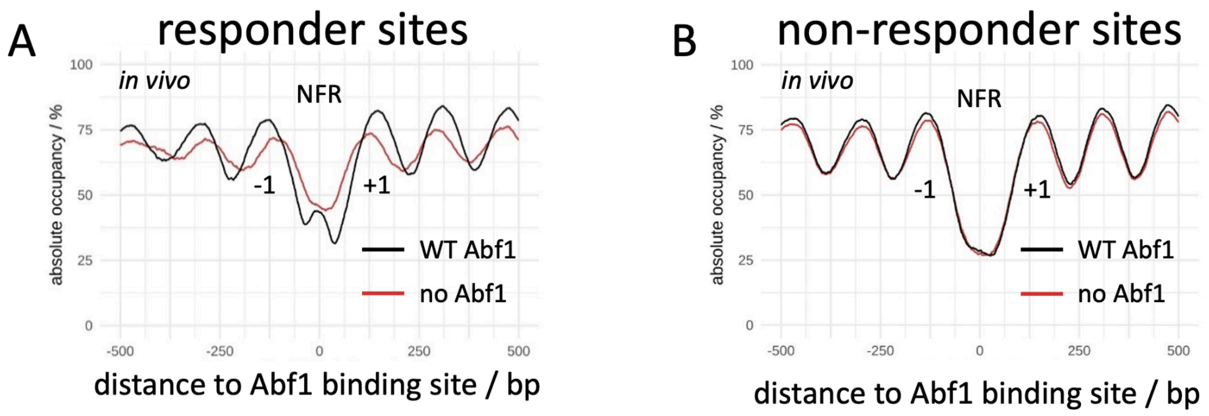


**Fig. S24 Nucleosomal changes as a function of Abf1 depletion via anchor away.**
In the absence of Abf1 from the nucleus, the nucleosome-free region (NFR) around Abf1 sites began to be filled up with nucleosomes while the flanking nucleosomes (-1, +1) and up-/downstream nucleosome arrays became disordered as seen by decreased peak-to-trough ratios and shifts of occupancy peaks towards the NFR. **(A)** As **Fig. 6D**, but data are aligned at responder Abf1 sites (class I and II, n = 397)) as identified by Kubik et al. (*17*) based on MNase-seq data in an analogous Abf1 anchor away strain and oriented such that a gene is to the right for unidirectional promoters. **(B)** As panel A but for non-responder sites (class III to V, n = 478) according to Kubik et al. (*17*).

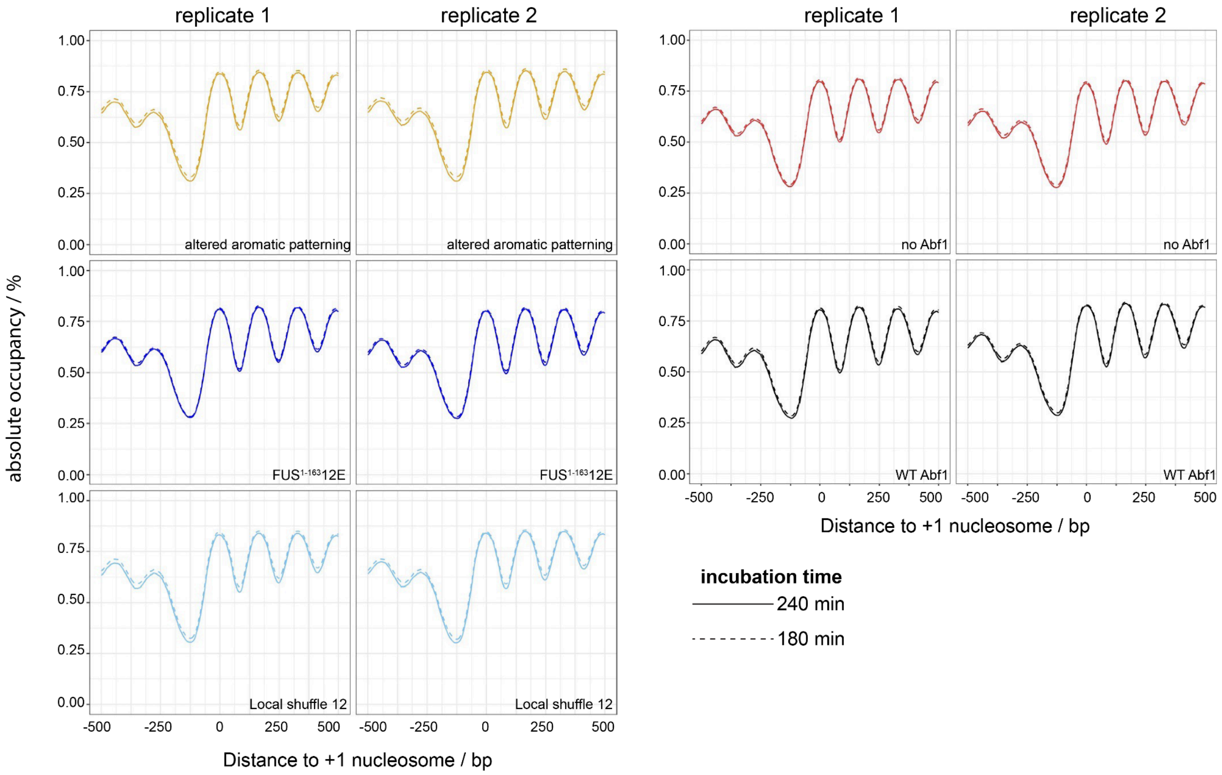


**Fig. S25. ODM-seq reaches saturated DNA methylation**. Incubation of each chromatin preparation with CpG-specific DNA methyltransferase M.SssI for 180 vs. 240 min results in the same occupancy patterns of in vivo +1 nucleosome-aligned composite plots. Importantly, active methyltransferase activity during the 60 additional minutes of incubation for the 240 min samples was demonstrated by de novo methylation of plasmid pFMP233 spiked-in at 180 min (**fig. S26**).

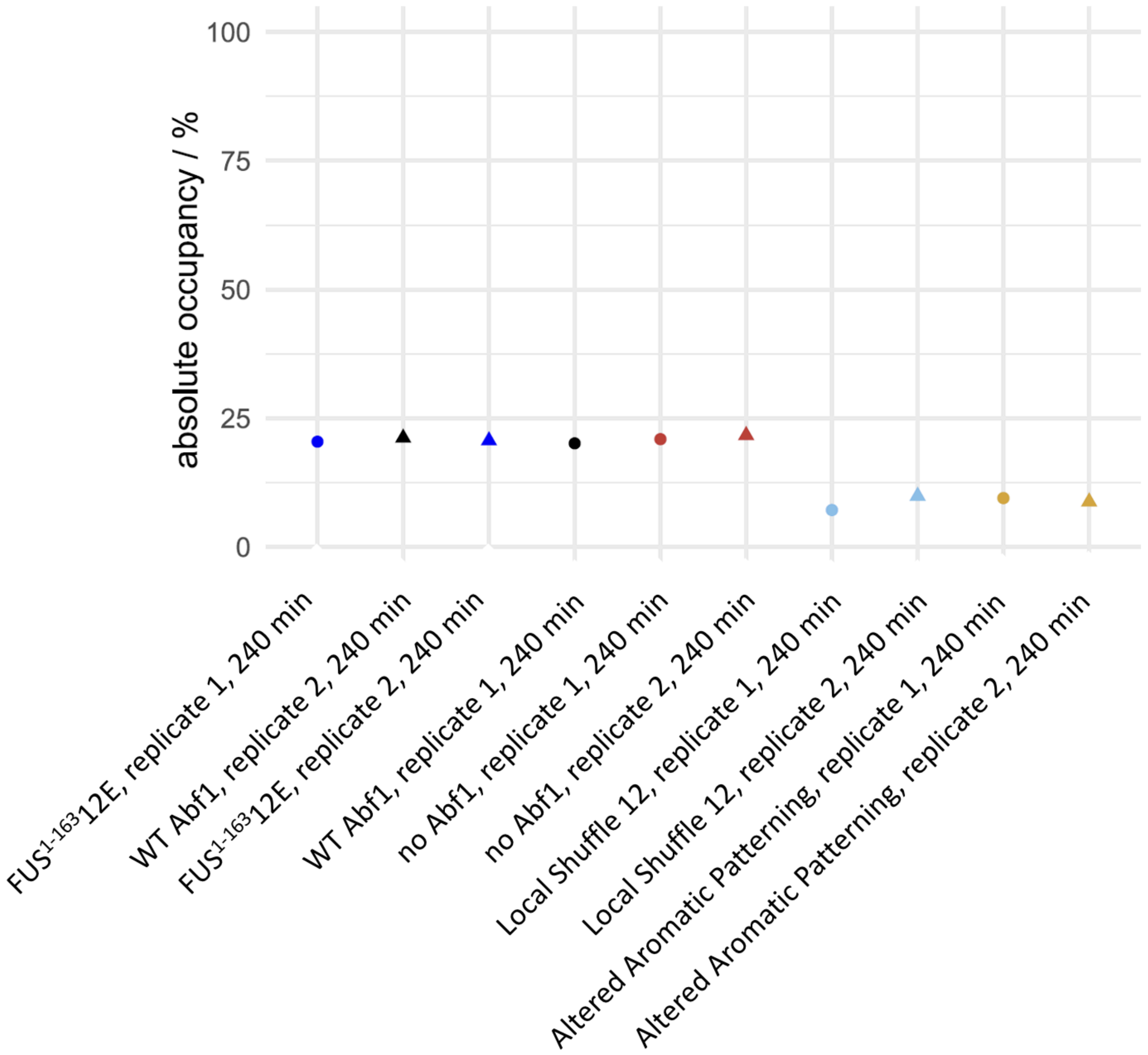


**Fig. 26. Average absolute occupancy** (ODM-seq; note that occupancy = 1 - methylation) data for plasmid pFMP233 that was spiked into the indicated samples at 180 min.

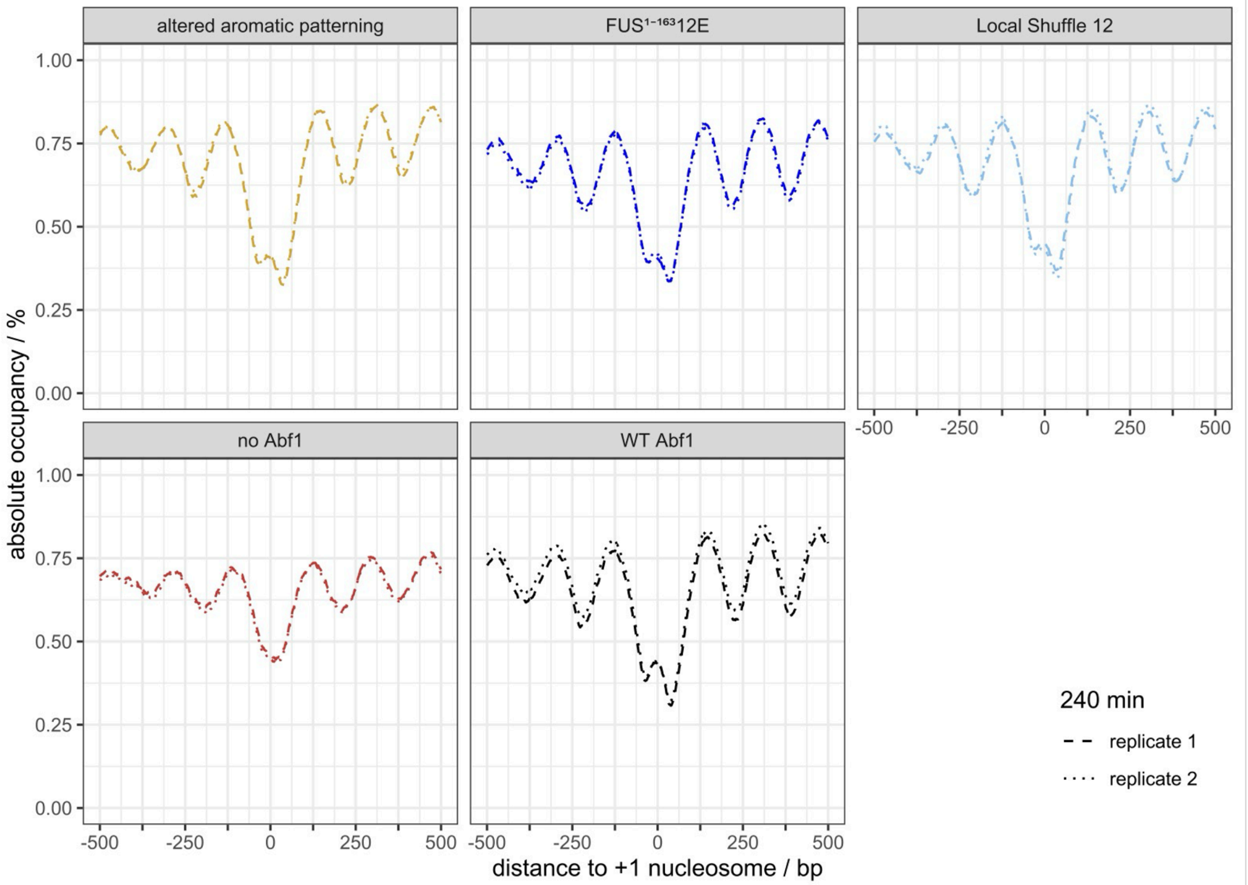


**Fig. S27. Independent biological replicates demonstrate near-perfect reproducibility in ODM-seq occupancy patterns**. ODM-seq data for the indicated samples after 240 min incubation with M.SssI are aligned at responder Abf1 sites according to Kubik et al. (*17*) (see also **fig. S24A**).

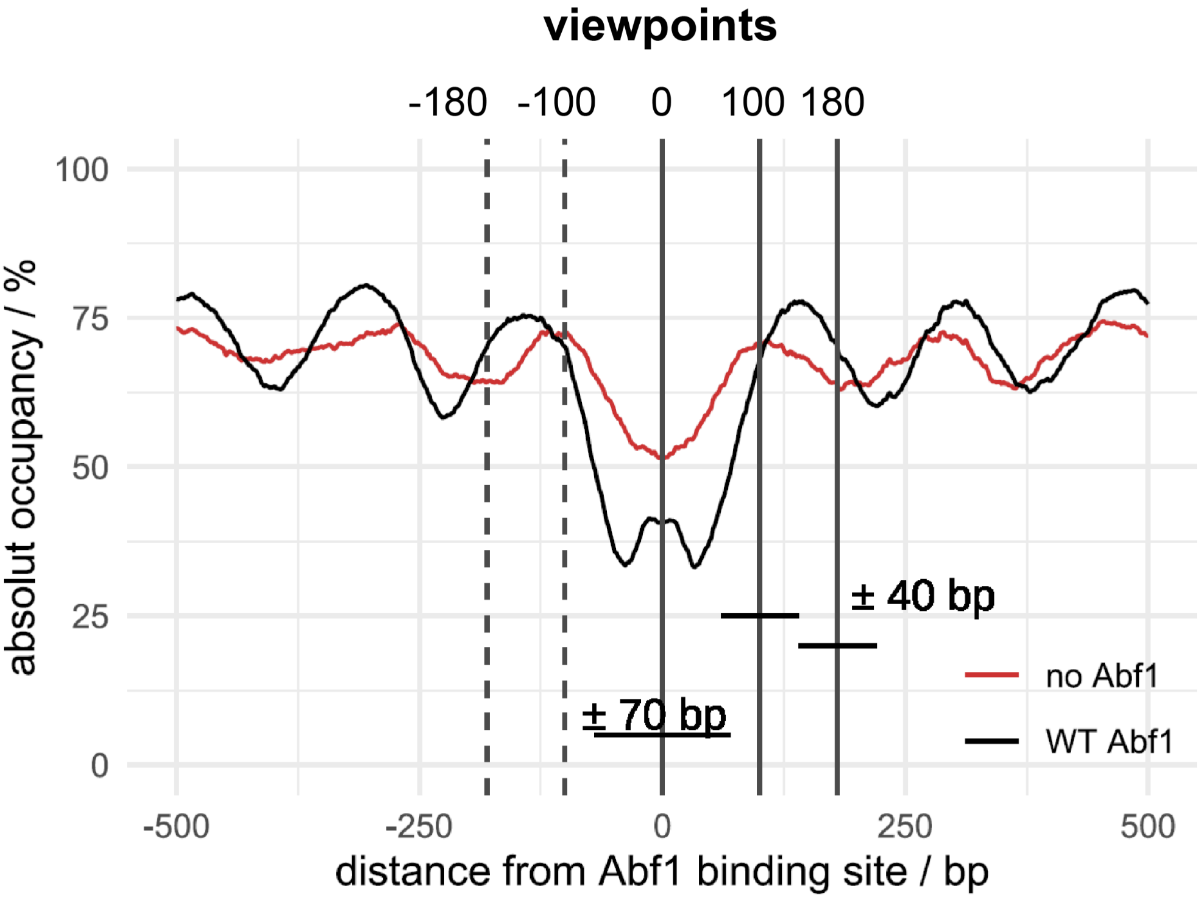


**Fig. S28. Schematics for visualizing different types of responders.** Schematics that visualize the different viewpoints and their windows where we searched for significant differences between ODM-seq occupancy patterns of WT Abf1 and no or variant Abf1 constructs with regard to the average absolute occupancy (1 - DNA methylation) across the window. Windows were centered either at the Abf1 sites (0 on the x-axis, viewpoint "0") or ± 100 bp or ± 180 bp (viewpoints "100" or "180") on both sides of the Abf1 sites and had a size of either 141 bp (± 70 bp from center) or 81 bp (± 40 bp from center). As an example, ODM-seq data is shown for WT and no Abf1 aligned at all promoter Abf1 sites (within 80-140 bp upstream of the transcription start sites, n = 188), where the no Abf1 sample showed significantly more occupancy in the 141 bp window centered at the Abf1 sites.

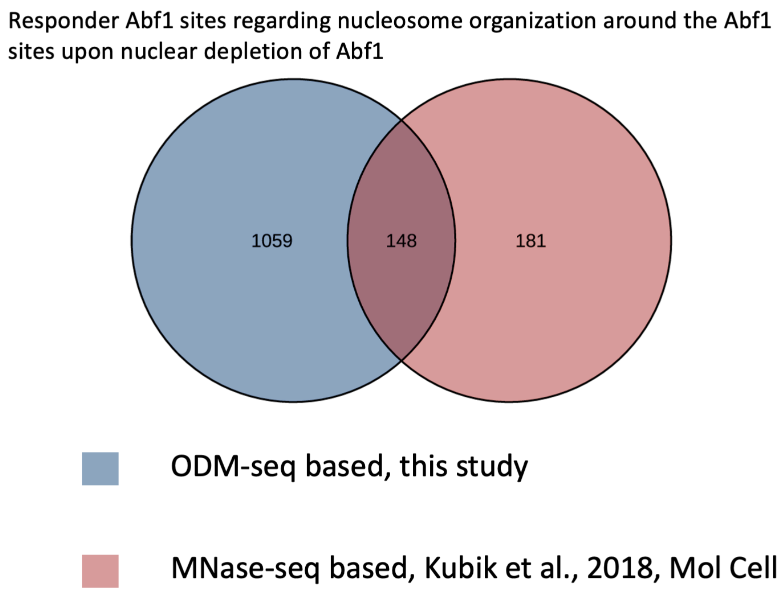


**Fig. S29 Repsonder sites identified by ODM-seq vs. responder sites identified by MNase-seq.** Venn diagram comparing responder sites after depletion of Abf1 from the nucleus in the Abf1 anchor away strain according to our own analysis of ODM-seq data and starting from just PWM-predicted Abf1 sites versus according to MNase-seq data and starting from only promoter Abf1 sites predicted by PWM and with positive ChIP signal (*17*). ODM-seq recovered many more responder sites than previously identified, and also some sites identified by Kubik *et al.* were not recovered in our analysis. As discussed in the methods, the larger number of responder sites may be explained because we did not limit our search to Abf1 sites in promoters that were confirmed by ChIP data, but included in an unbiased way all 2126 sites predicted by position weight matrices (PWMs; (*18*, *19*)). Conversely, the responder sites missed by us relative to the responder sites called by Kubik *et al.* may be due to our stringent significance threshold of $p \leqslant 0.01$.

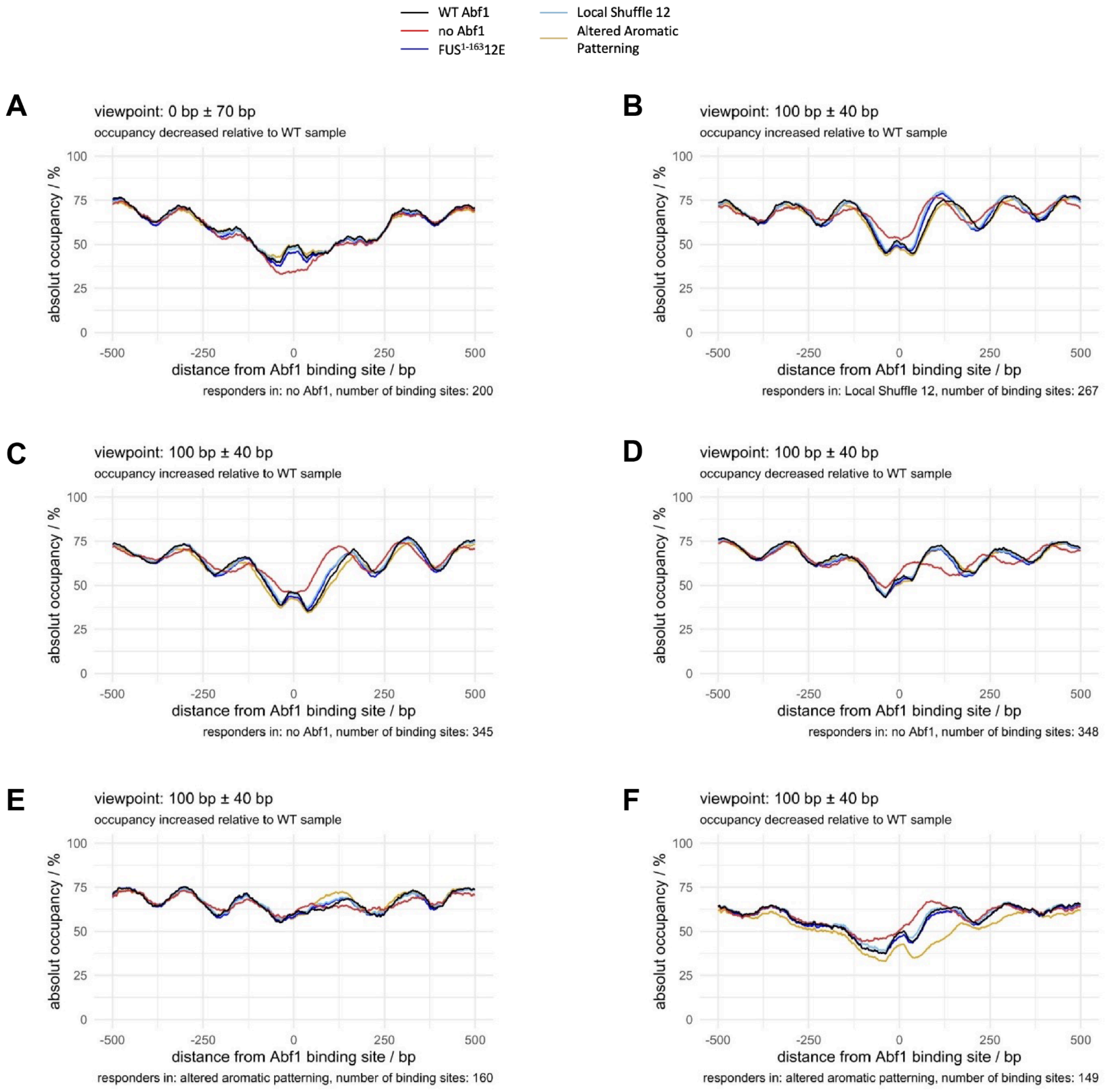


**Fig. S30 (Part 1/4; caption with part 4)**

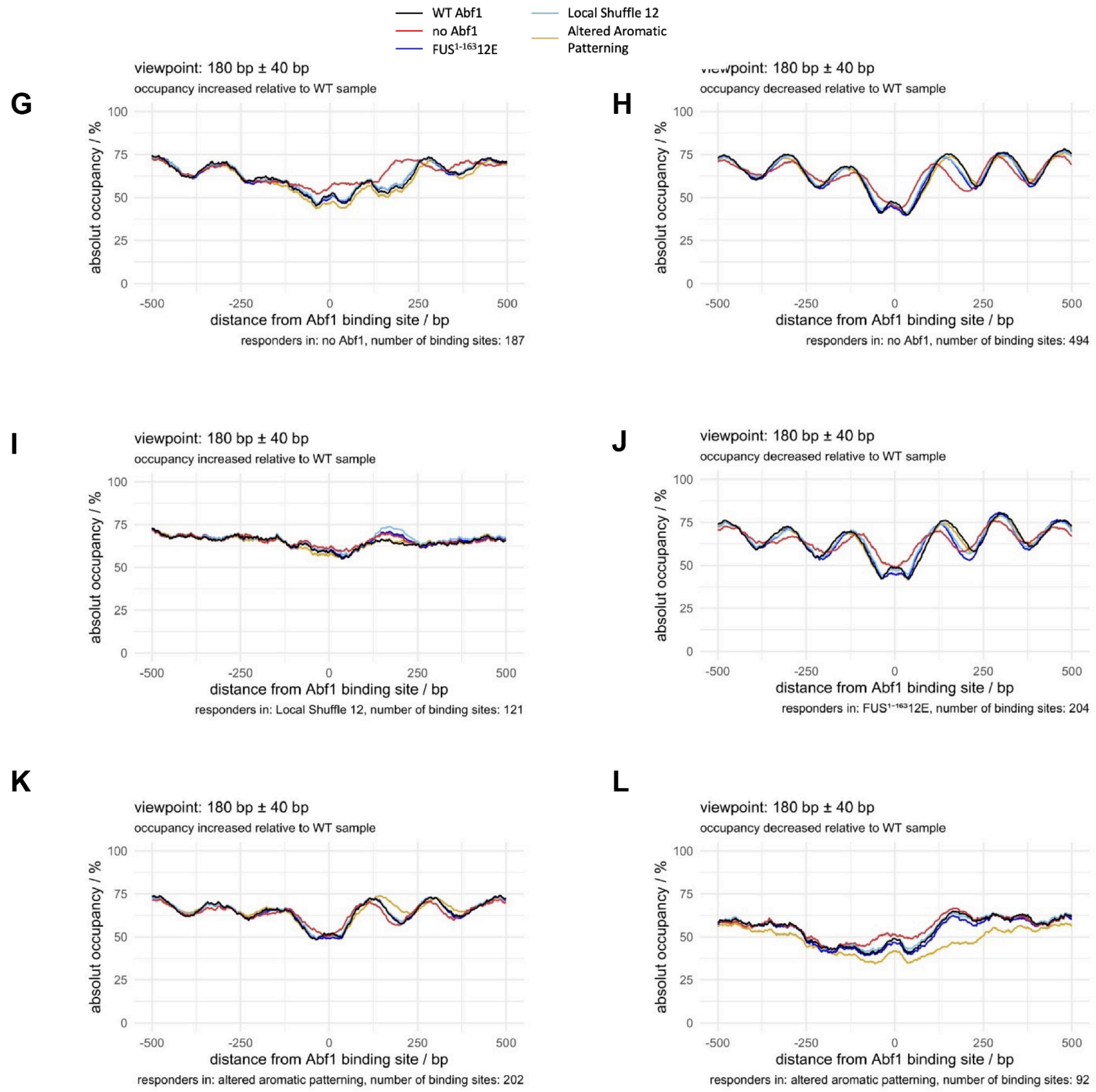


**Fig. S30 (Part 2/4; caption with part 4)**

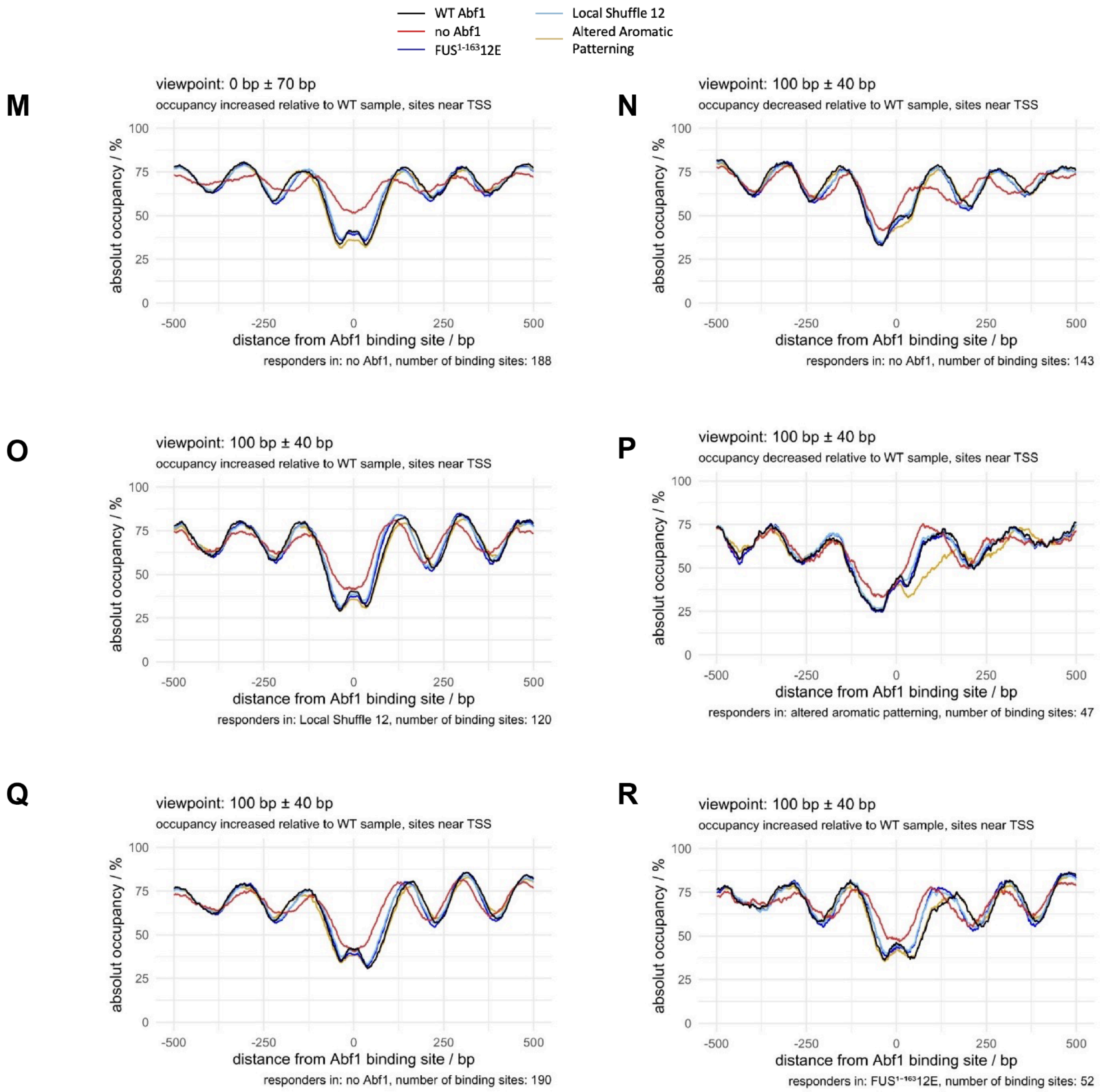


**Fig. S30 (Part 3/4; caption with part 4)**

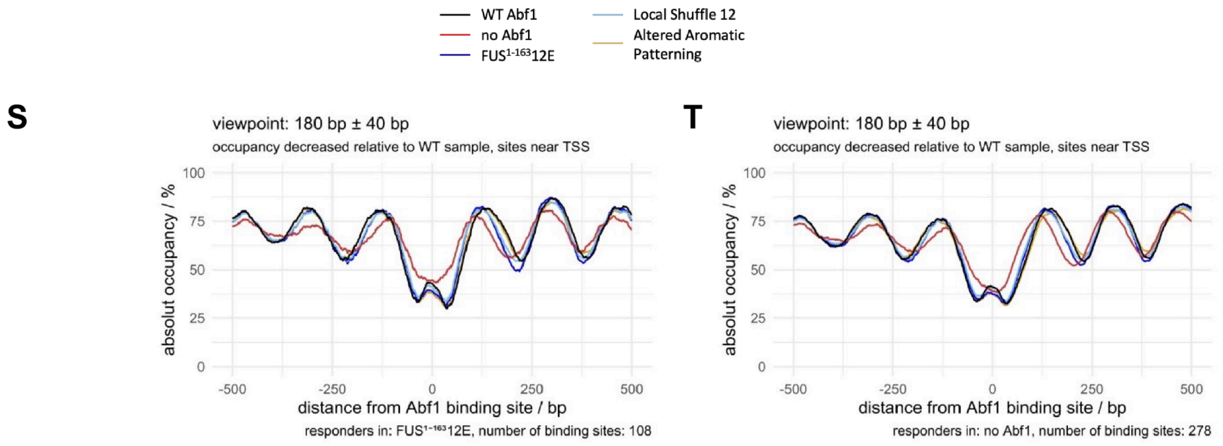


**Fig. S30 (Part 4/4).** Like **Fig. 6D,F,G**, but for the Abf1 responder sites, either all or 80-180 bp upstream of a transcription start site (TSS), as defined in **fig. 28** and Methods and counted sites are listed in **Table S8**.

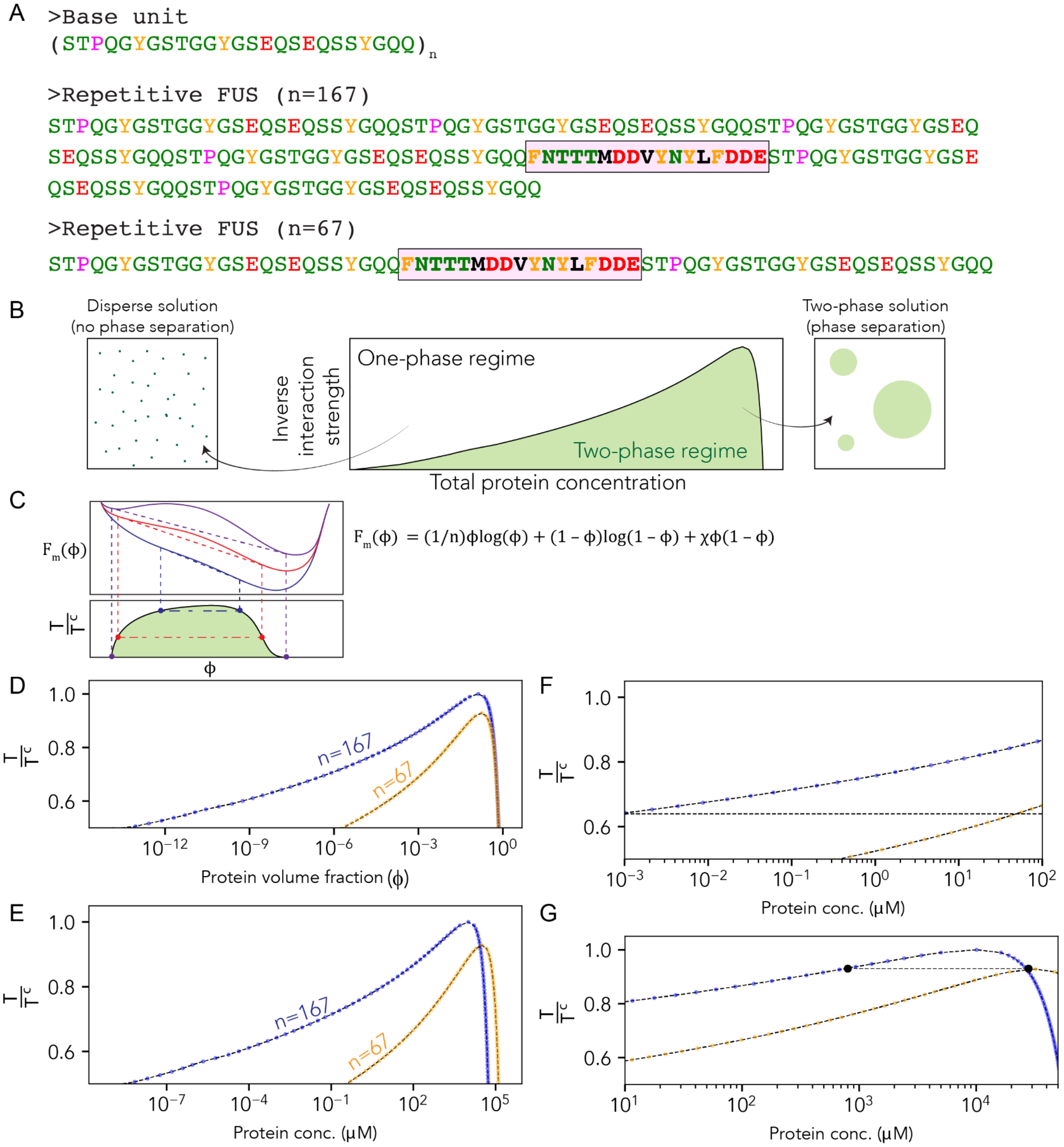


**Fig. S31 Flory-Huggins theory predicts that sequence variants that alter sequence length should change the saturation concentration by five orders of magnitude within the relevant regime.** To ask if a phase separation model might be reasonable we used Flory-Huggins theory to predict the expected shift in the saturation concentration ($c_{sat}$) as a function of IDR length. **(A)** We designed a basic repeating unit (top) and used this base unit to generate n=6 and n=2 units of synthetic polymers (middle and bottom). **(B)** Flory-Huggins theory allows us to predict an expected change in saturation concentration for two polymers with identical attractive interactions but of different lengths. As such, in a phase separation model, we can conceptually predict how changing the IDR length would shift the saturation concentration. **(C)** To construct phase diagrams requires reconstruction from the free energy of mixing to extract coexisting phases. The expression for the free energy of mixing is shown to the right (*56*). **(D)** Phase diagram for n=67 and n=167 polymers in reduced temperature (y-axis)

vs. volume fraction (x-axis) space. Reduced temperature provides an effective parameter that tunes intramolecular interactions up to the critical temperature (reduced temperature = 1.0). As such, the y-axis position reflects the normalized intermolecular interaction between polymers. With this in mind, we can convert from volume fraction into molar concentration and ask how, at any arbitrary intermolecular interaction strength, the saturation concentration (in µM) would change in response to a change in IDR length. **(E)** Full phase diagram showing a change in molar concentration of the phase diagram expected for the two IDRs of length 67 and 167 residues. **(F)** Under concentration regimes compatible with cellular concentrations, the change in length from 167 residues to 67 residues yields a predicted change in the saturation concentration of almost five orders of magnitude. **(G)** Even at the top of the phase diagram, where the difference is minimal, we find a difference in concentration of almost two orders of magnitude.

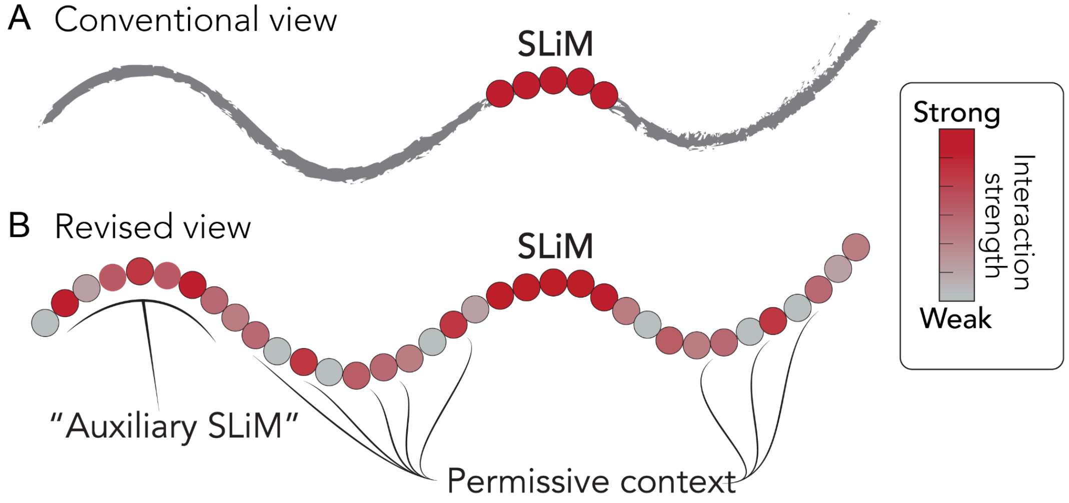


**Fig. S32. A revised model for IDR-dependent molecular recognition affords evolutionary change.** In a more conventional view of IDRs that possess SLiMs, a specific subregion is considered the key determinant of binding. We propose a revised version of this in which an apparent motif reflects the single strongest region, but many additional regions (that could be designated as auxiliary motifs) along the IDR have weak, transient contributions to binding affinity and, possibly, binding specificity. Under this model, new motifs can emerge and disappear through compensatory changes across an IDR.

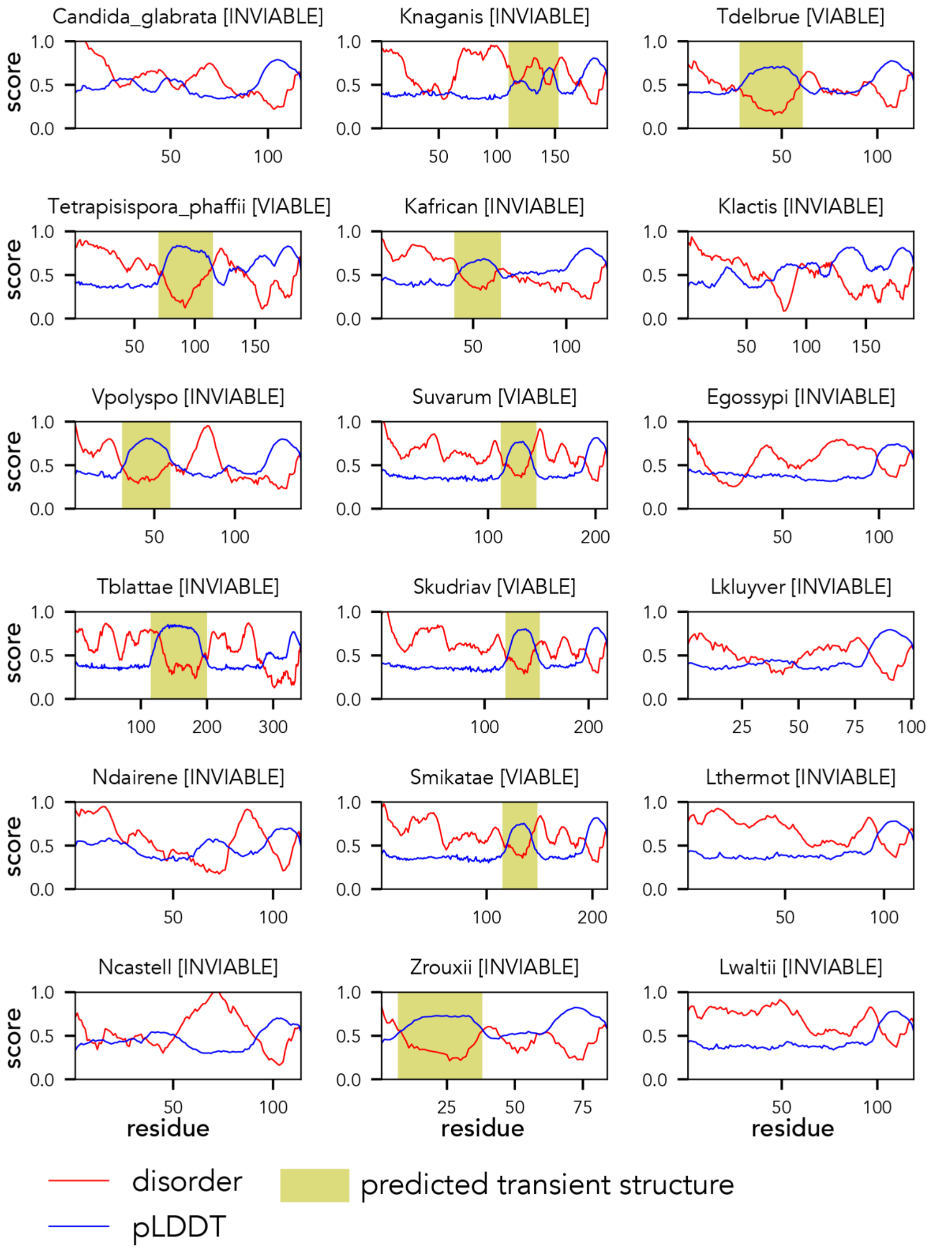


**Fig. S33 Abf1 orthologs IDR2 analysis**. All the viable orthologous sequences have a predicted structured region that we interpret to be a local helical region (highlighted in yellow). However, several inviable sequences also show a predicted structured subregion, suggesting this is not a sufficient feature for viability. The fact that a helical sub-region is present and absent across orthologs is consistent with a model in which rapid evolutionary changes in IDRs can drive the gain and loss of SLiMs within transient helical regions, although we emphasize we do not know if these predicted transient helices do harbor SLiMs.

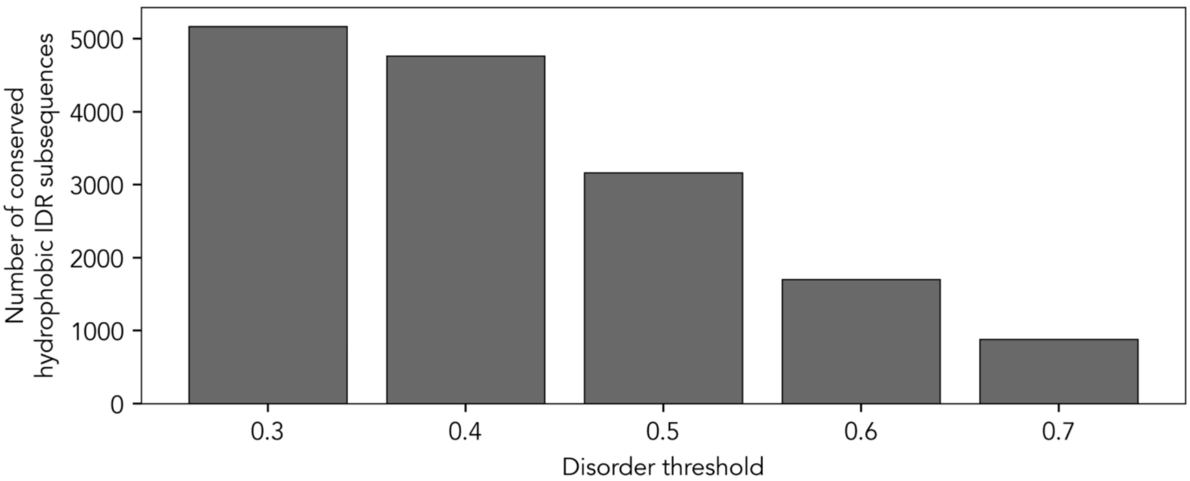


**Fig. S34 Number of conserved hydrophobic subsequences**. Using the conservation threshold of 0.65 (**fig. S4**) we identified subsequences based on the presence of conserved hydrophobic residues centered on a +/-6 residues window to identify conserved subregions. Hydrophobic residues here were defined as I, L, M, V, Y, F, W. Initiator methionines were discarded from the analysis. Depending on the stringency of disorder score for the resulting subsequence, between 5173 and 884 conserved subsequences are identified across IDRs in the yeast genome. All subsequences at all disorder thresholds are reported in **Table S10-S14**.

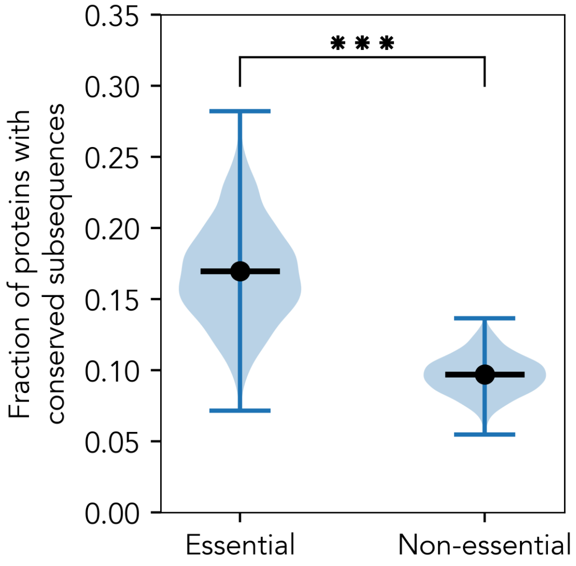


**Fig. S35 Number of conserved hydrophobic subsequences in essential vs. non-essential yeast proteins**. On average there are almost twice as many conserved subsequences in essential than in non-essential proteins. Distribution generated by bootstrapping (black line is overall mean) with a p-value below the detectable threshold by independent t-test with unequal variance. The analysis here uses the most stringent disorder threshold (=0.7, see **fig. S6**) but analogous results are obtained regardless of the used threshold.

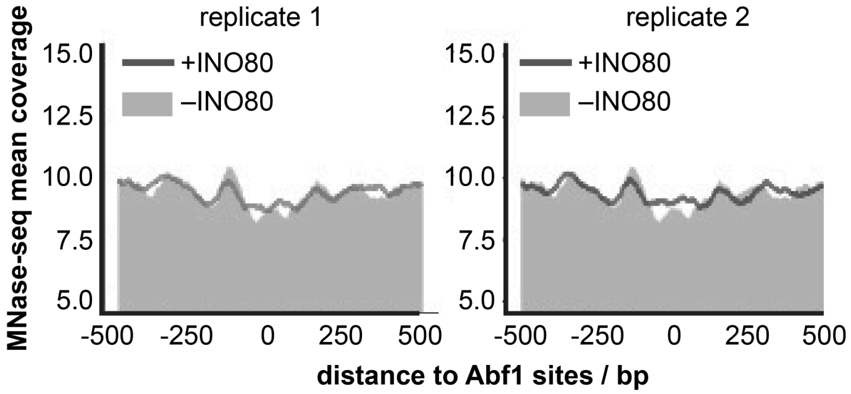


**Fig. S36.** MNase-seq data for yeast genome shows that INO80 cannot generate nucleosomal arrays phased to Abf1 sites without Abf1. As **Fig. 6E** but with n = 1071 and indicated conditions and two independent replicates.

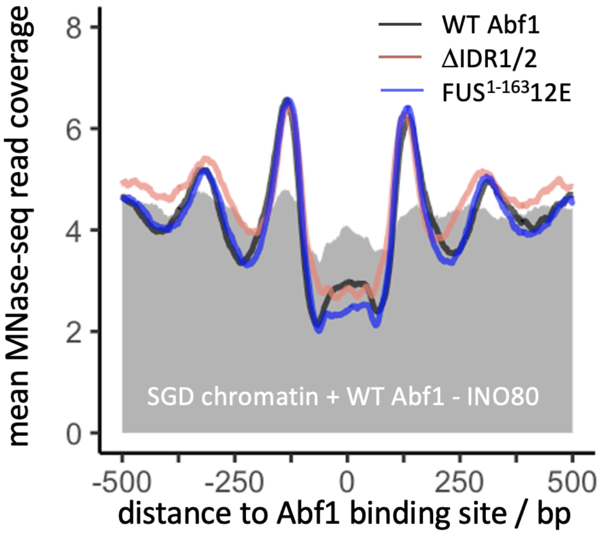


**Fig. S37.** Replicate in vitro reconstitution experiment as in **Fig. 6E** but with some altered conditions as detailed in *Methods*.

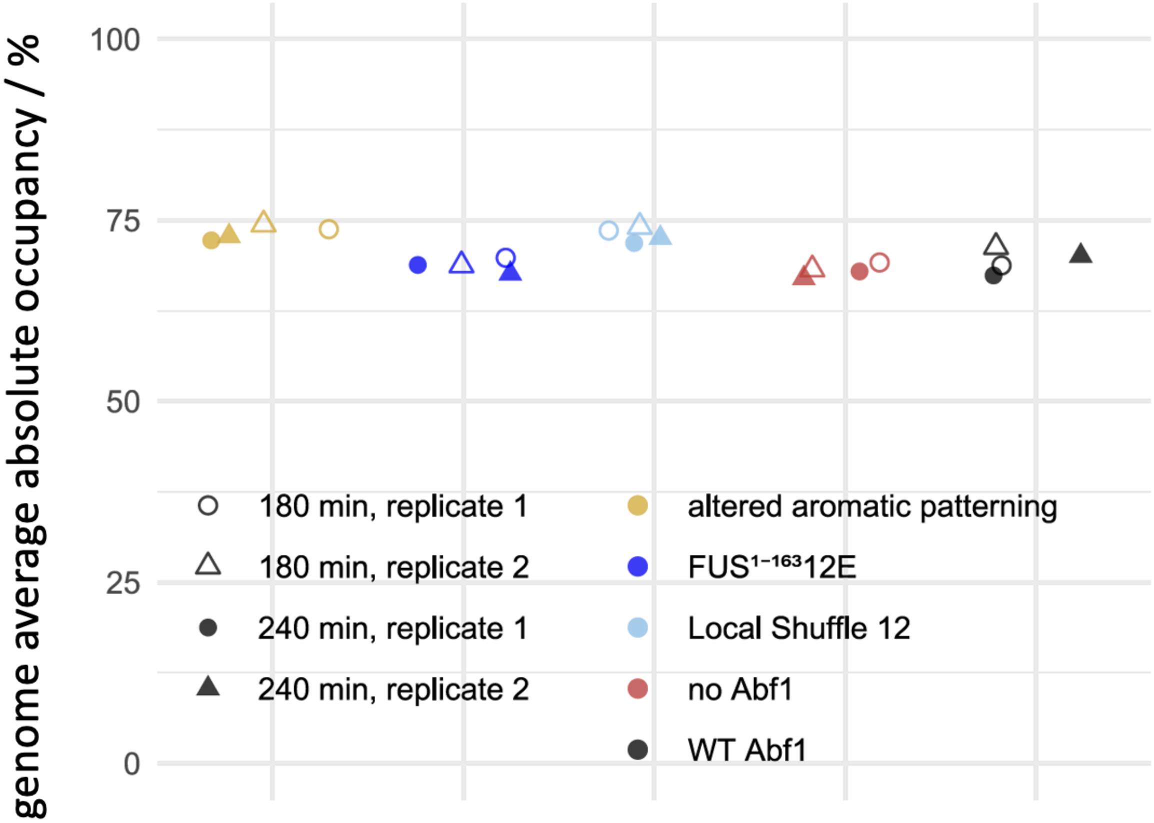


**Fig. S38. Genomic average methylation for each sample across all coordinates.**

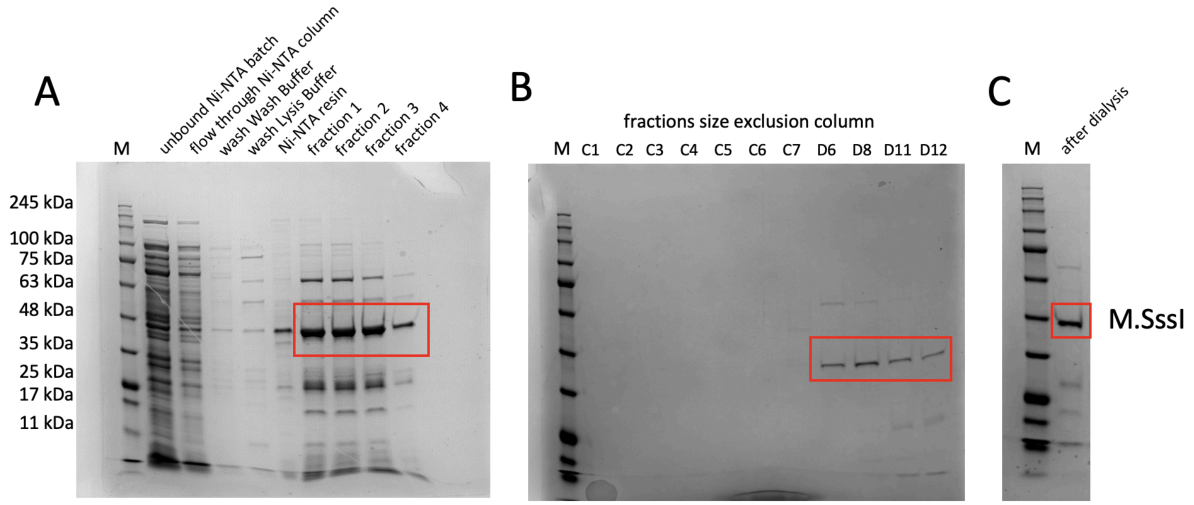


**Fig. S39**. **Purification of M.SssI. (A)**, **(B)**, **(C)** show 10% SDS-PAGE (Serva) analysis of the indicated steps and fractions during the purification of M.SssI (Methods). Red boxes highlight the bands corresponding to M.SssI (42 kDa). Marker lane (M): Triple Color Protein Standard III (39258, Serva).

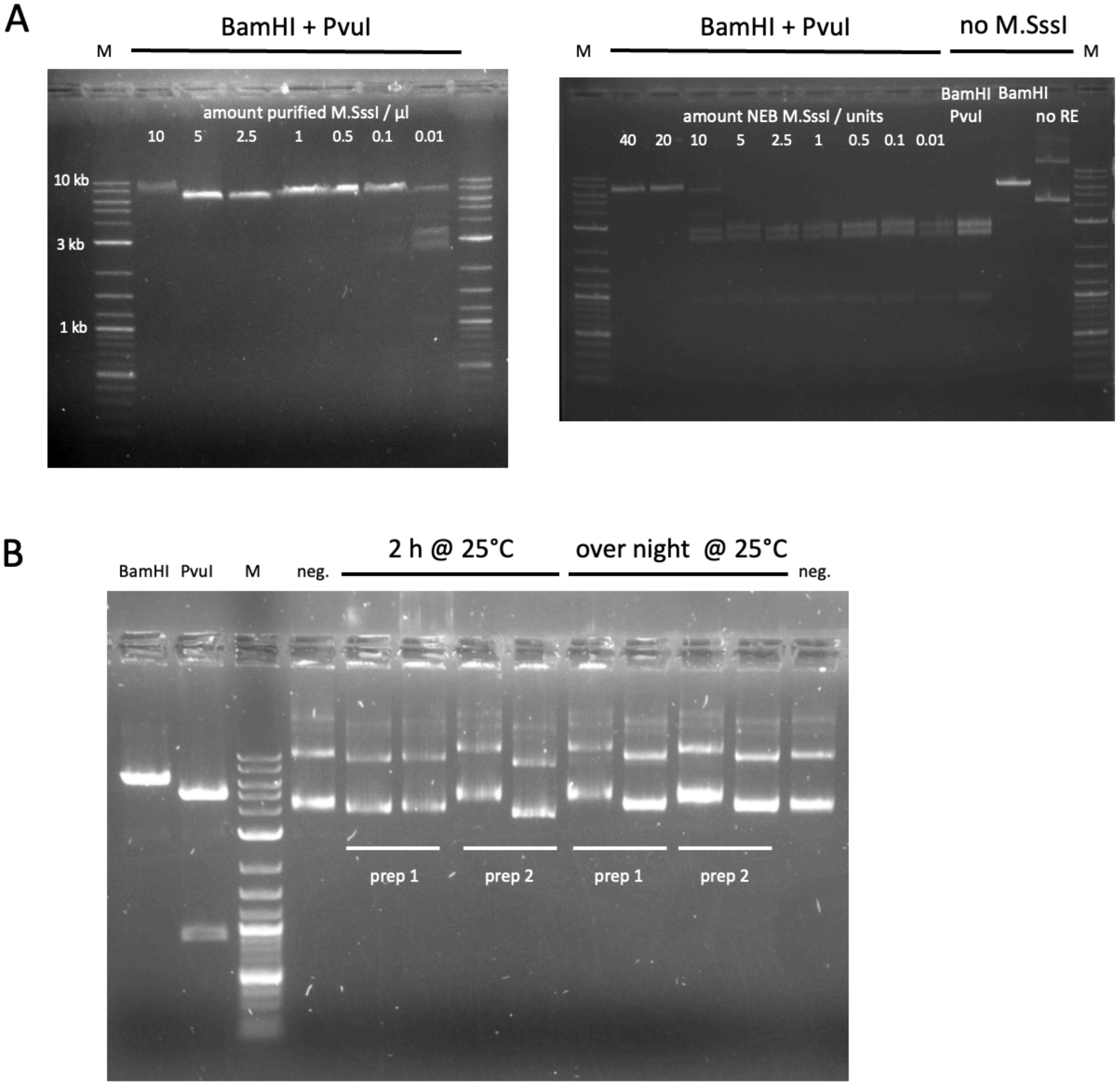


**Fig. S40. Characterization of purified M.SssI. (A)** Comparison of DNA methylation activity at CpG sites between our purified (left) and commercial (NEB, right) M.SssI. After plasmid pUC1-ftz (5769 bp) was incubated with indicated amounts of M.SssI in the presence of SAM and purified, it was digested with CpG methylation-non-sensitive BamHI (unique site: linearization) or BamHI + CpG methylation-sensitive PvuI (2544, 2329, 896 bp fragments if not blocked by CpG methylation). DNA fragments were electrophoresed in 1% agarose 1xTAE gels. 0.5µl of our here-shown M.SssI preparation had similar activity as 20 units of the commercial M.SssI, i.e. a concentration of 40 u/µl. Marker lane: 1 kb plus ladder (NEB). **(B)** Purified M.SssI preparations (prep 1 and prep 2) were free from nuclease contamination as they did not remove the supercoiled form of plasmid pUC19-ftz even after overnight incubation in two different buffers, NEB rCutSmart and NEB Buffer 2, in the left and right lane, respectively, for each test reactions of prep 1 and 2.

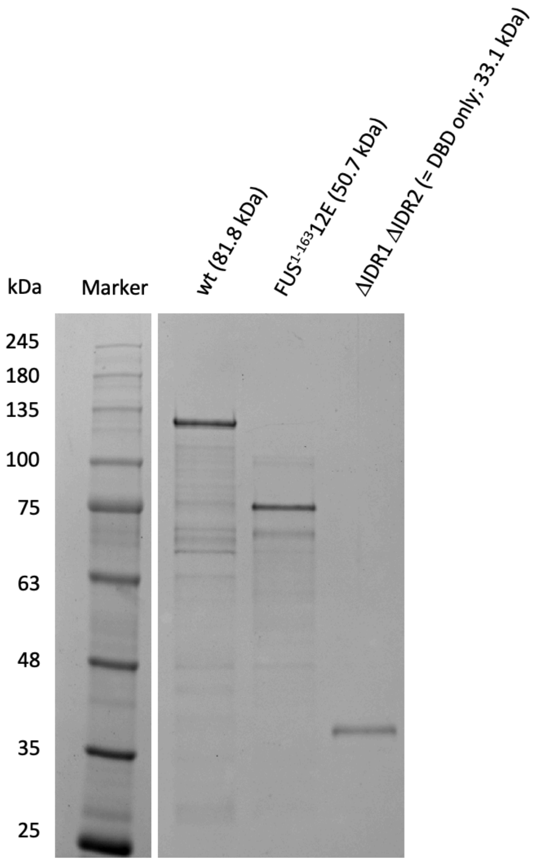


**Fig. S41.** 10% SDS-Page (Serva) analysis of indicated WT or variant Abf1 preparations. Marker: Triple Color Protein Standard III (39258, Serva)

**Table S1.**
Summary of results for Abf1 wildtype and mutant constructs. Found in Supplementary_tables.xlsx file.

**Table S2.**
PCR primers and templates for generation and resulting sequences of *ABF1* wildtype and truncation constructs. Found in Supplementary_tables.xlsx file.

**Table S3.**
PCR primers, templates and synthesized DNA sequence for generation of rationally designed constructs. Found in Supplementary_tables.xlsx file.

**Table S4.**
PCR primers and templates for generation and resulting sequences of random mutagenesis constructs. Found in Supplementary_tables.xlsx file.

**Table S5.**
Sanger sequencing primers. Found in Supplementary_tables.xlsx file.

**Table S6.**
qPCR primers for ChIP. Found in Supplementary_tables.xlsx file.

**Table S7**
Viablity scores for all viable constructs. Found in Supplementary_tables.xlsx file.

**Table S8**
Abf1 responder sites. Found in Supplementary_tables.xlsx file.

**Table S9**
IDRs that are chemically conserved and not conserved in terms of sequence

**Table S10-S14.**
Conserved hydrophobic subsequences were identified across the yeast proteome. Found in Supplementary_tables.xlsx file.

**Table S15**
Linear sequence conservation and compositional conservation for all IDRs in the yeast proteome. Found in Supplementary_tables.xlsx file.

**Table S16**
Compositional classification weights that were used in **Fig. 4J**. Found in Supplementary_tables.xlsx file.

**Table S17**
Subsequences from disordered regions with predicted transient structure. Found in Supplementary_tables.xlsx file.
