## Supplementary material for "Sequence- and chemical specificity define the functional landscape of intrinsically disordered regions": Fig S1

### Supplemental figures

#### figure S1: Viability of Abf1 constructs assessed by 5-FOA plasmid shuffling assay.

Strains harboring both pRS416-*ABF1* plasmid (*URA3* marker) and pRS315 plasmid bearing the *abf1* mutant construct gene and the *LEU2* marker in the strain background with deleted *abf1* chromosomal gene were re-streaked from patches from YNB w/o ura, leu plates onto 5-FOA w/o leu plates. Only constructs discussed in the paper are labeled in plate schemes, where names of viable strains are in blue, names of inviable ones in black. Viability on 5-FOA plates was visually scored by comparison to known viable or inviable strains on the same plate. Inviable strains were distinguished from viable strains because they showed a smeary appearance and at most tiny/sparse single colonies, whereas viable strains grew as a more or less dense patch.
For each construct, independent clones were tested with the clone number given in the plate schemes. On each slide, a table summarizes the result of the shown plate and in brackets the total result obtained for this construct if all tested clones on all plates were considered. If the same clone showed conflicting results in technical replicates, it was not counted as viable or inviable, but included in the total number of tested clones. The final categorization of each construct as viable or inviable followed the majority of tested clones excluding the clones with ambiguous results in technical replicates.

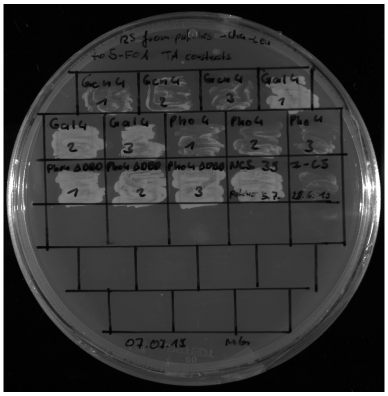

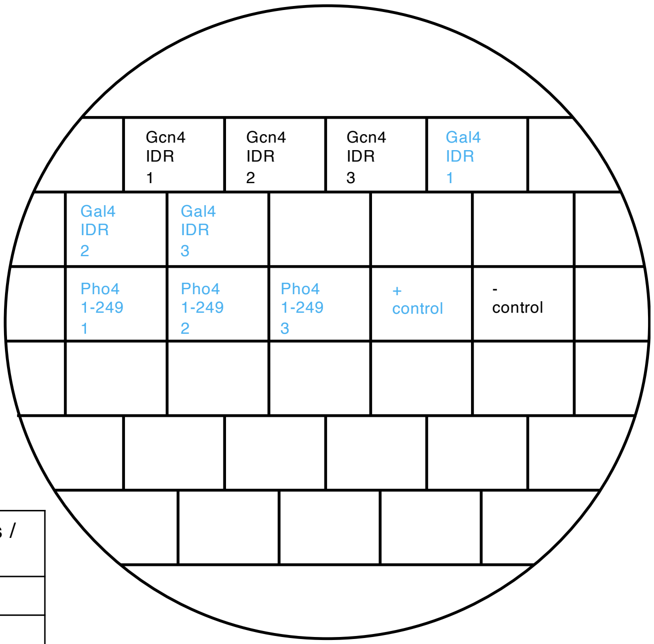


Restreak from YNB –ura –leu to 5-FOA –leu

| Construct | Result regarding viability on 5-FOA | Number of (in)viable clones / total number of clones |
|---|---|---|
| Gcn4 IDR 17-150 | inviable | 3 out of 3 |
| Gal4 IDR 768-881 | viable | 3 out of 3 |
| Pho4 1-249 | viable | 3 out of 3 |

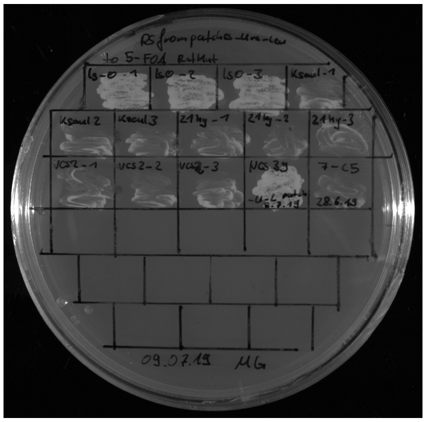

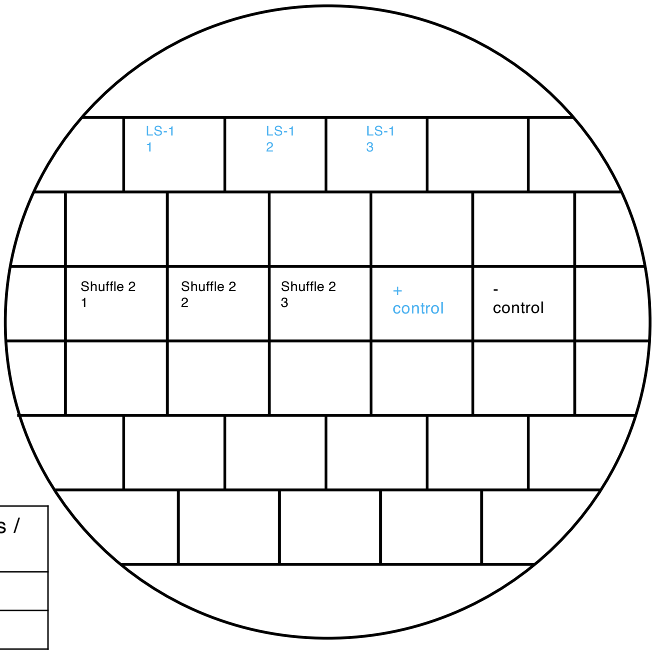


Restreak from YNB –ura –leu to 5-FOA –leu

| Construct | Result regarding viability on 5-FOA | Number of (in)viable clones / total number of clones |
|---|---|---|
| LS-1 | viable | 3 out of 3 |
| Shuffle 2 | inviable | 3 out of 3 |

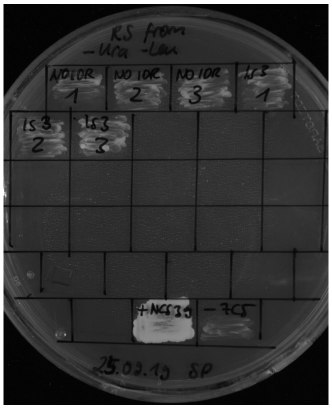

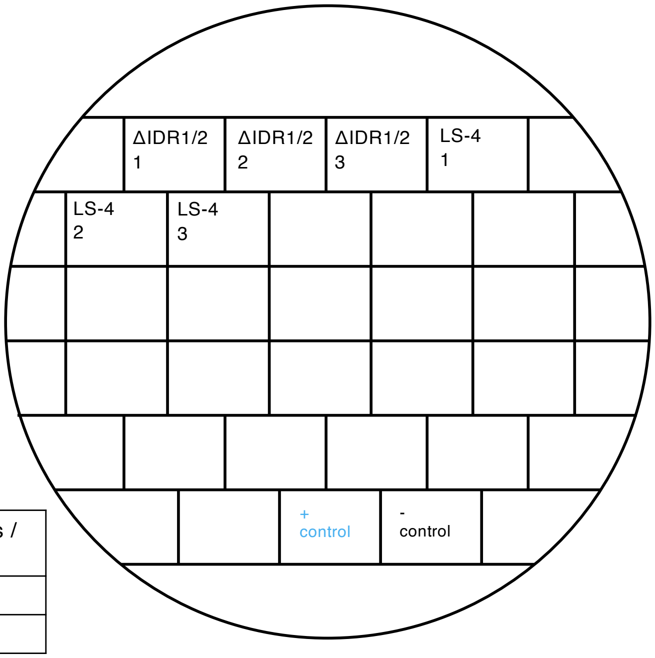


Restreak from YNB –ura –leu to 5-FOA –leu

| Construct | Result regarding viability on 5-FOA | Number of (in)viable clones / total number of clones |
| --- | --- | --- |
| ΔIDR1/2 (NLS-FLAG) | inviable | 3 out of 3 |
| LS-4 | inviable | 3 out of 3 |

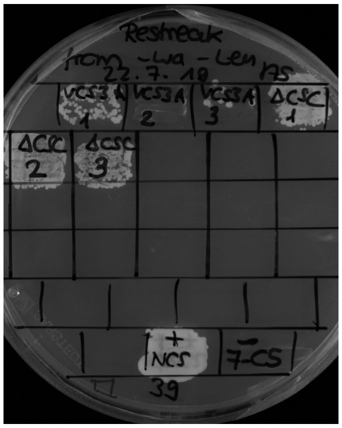

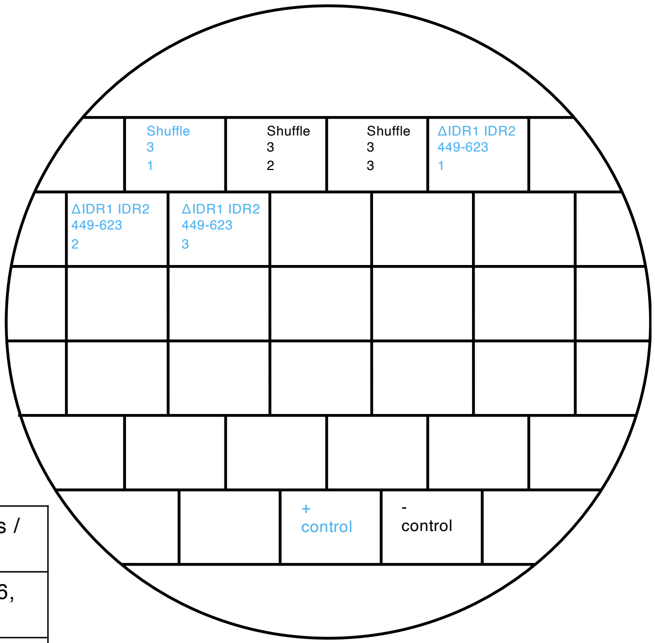


Restreak from YNB –ura –leu to 5-FOA –leu

| Construct | Result regarding viability on 5-FOA | Number of (in)viable clones / total number of clones |
| --- | --- | --- |
| Shuffle 3 | inviable | 2 out of 3 (in total 5 out ot 6, see also slide 4) |
| ΔIDR1 IDR2 449-623 | viable | 3 out of 3 |

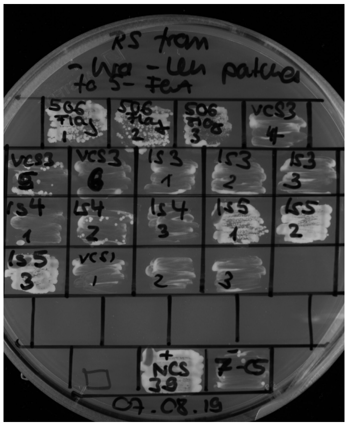

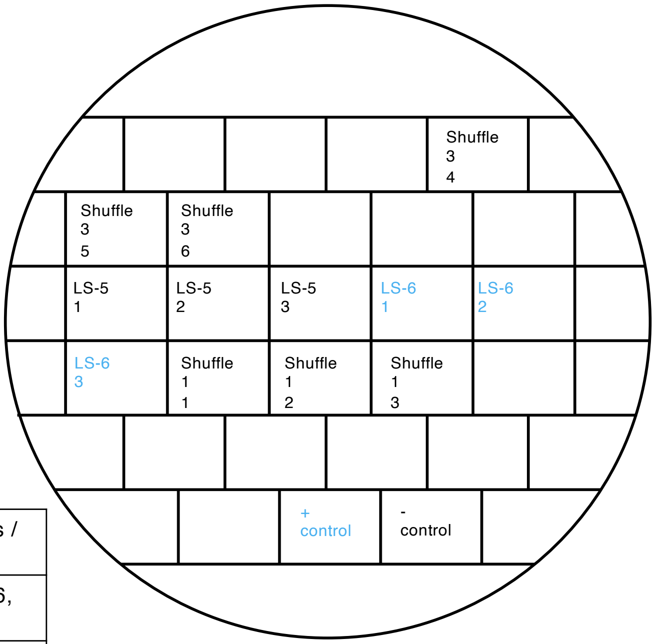


Restreak from YNB –ura –leu to 5-FOA –leu

| Construct | Result regarding viability on 5-FOA | Number of (in)viable clones / total number of clones |
|---|---|---|
| Shuffle 3 | inviable | 3 out of 3 (in total 5 out ot 6, see also slide 3) |
| LS-5 | inviable | 3 out of 3 |
| LS-6 | viable | 3 out of 3 |
| Shuffle 1 | inviable | 3 out of 3 |

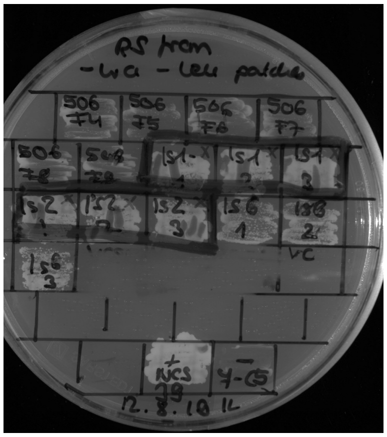

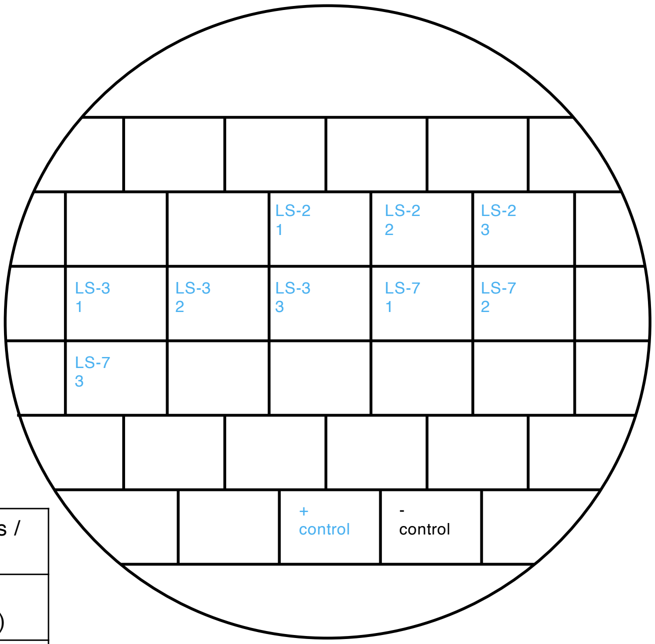


Restreak from YNB –ura –leu to 5-FOA –leu

| Construct | Result regarding viability on 5-FOA | Number of (in)viable clones / total number of clones |
|---|---|---|
| LS-2 | viable | 3 out of 3 (technical replicates, see also slide 6) |
| LS-3 | viable | 3 out of 3 (technical replicates, see also slide 6) |
| LS-7 | viable | 3 out of 3 (in total 8 out of 8, see also slide 5) |

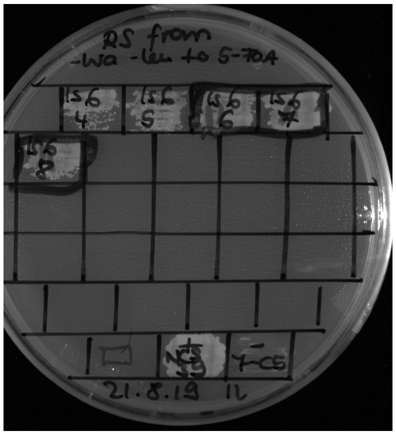


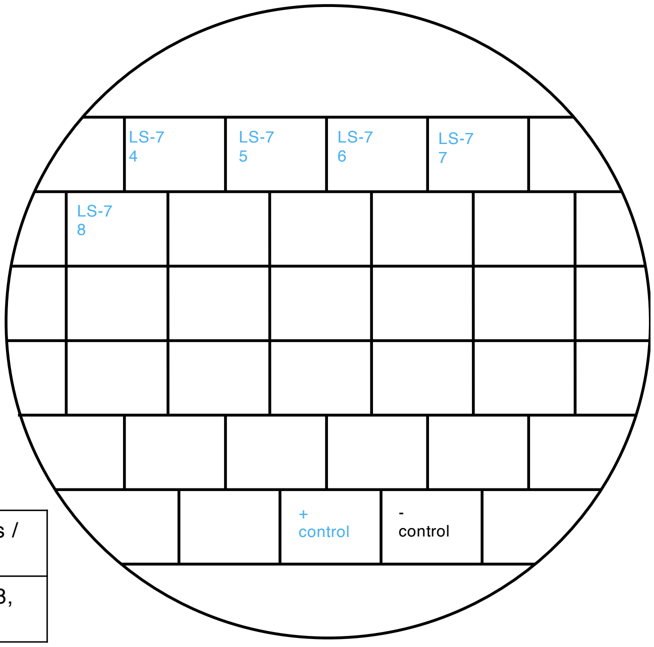


Restreak from YNB –ura –leu to 5-FOA –leu

| Construct | Result regarding viability on 5-FOA | Number of (in)viable clones / total number of clones |
|---|---|---|
| LS-7 | viable | 5 out of 5 (in total 8 out of 8, see also slide 4) |

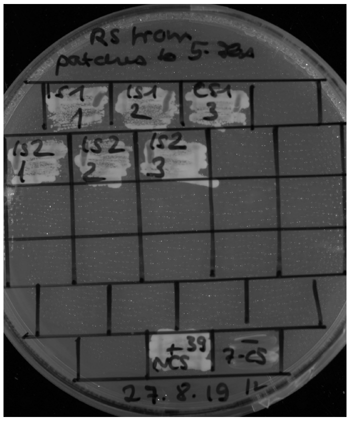

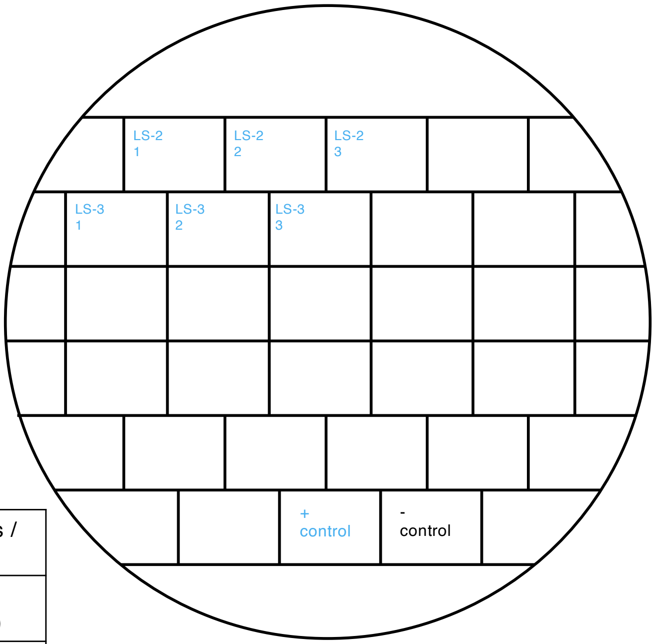


Restreak from YNB –ura –leu to 5-FOA –leu

| Construct | Result regarding viability on 5-FOA | Number of (in)viable clones / total number of clones |
|---|---|---|
| LS-2 | viable | 3 out of 3 (technical replicates, see also slide 4) |
| LS-3 | viable | 3 out of 3 (technical replicates, see also slide 4) |

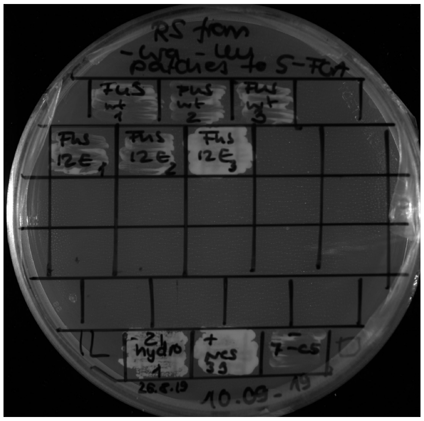

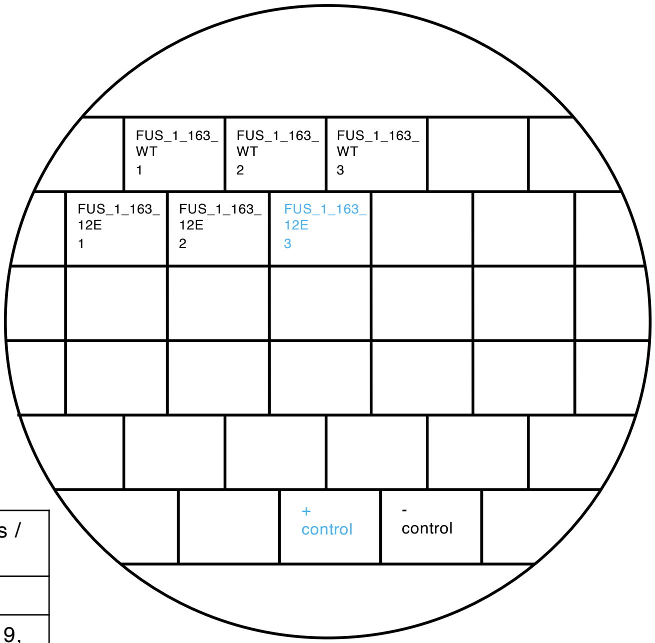


Restreak from YNB –ura –leu to 5-FOA –leu

| Construct | Construct | Number of (in)viable clones / total number of clones |
|---|---|---|
| FUS_1_163_WT | inviable | 3 out of 3 |
| FUS_1_163_12E | inviable | 2 out of 3 (in total: 8 out of 9, see also slides 7 and 14) |

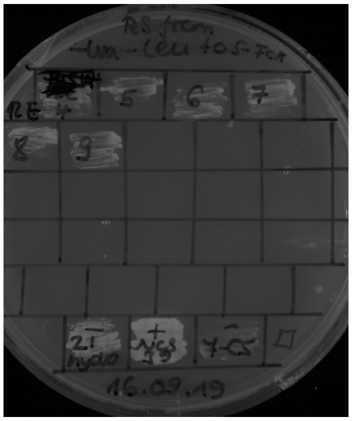

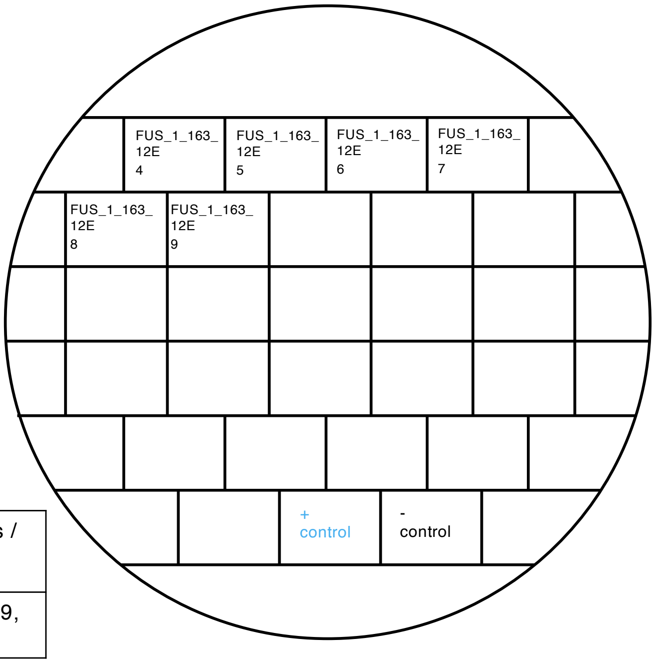


Restreak from YNB –ura –leu to 5-FOA –leu

| Construct | Result regarding viability on 5-FOA | Number of (in)viable clones / total number of clones |
|---|---|---|
| FUS_1_163_12E | Inviable | 6 out of 6 ((in total 8 out of 9, see also slides 6 and 14) |

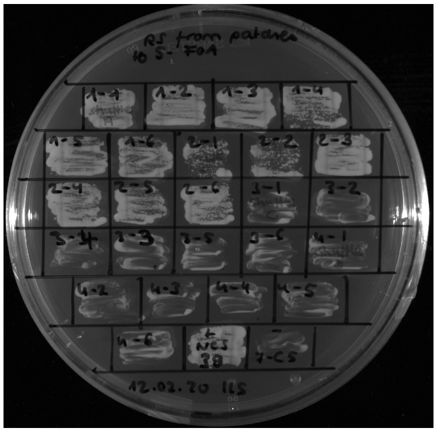

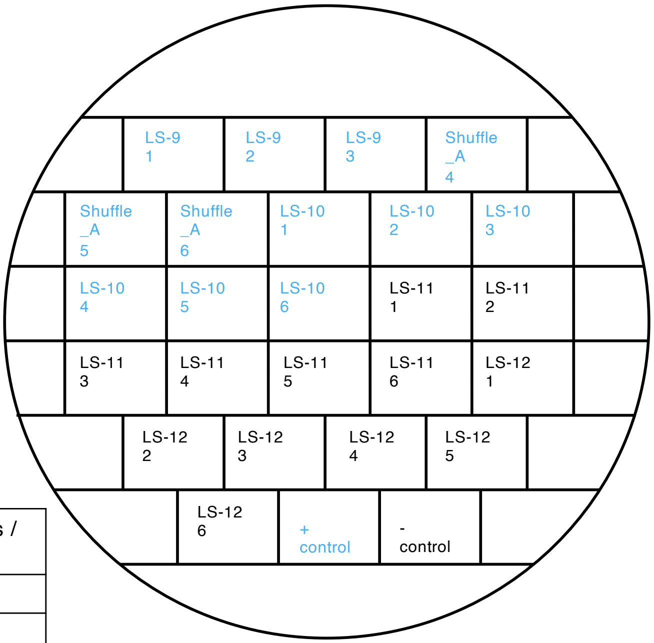


Restreak from YNB –ura –leu to 5-FOA –leu

| Construct | Result regarding viability on 5-FOA | Number of (in)viable clones / total number of clones |
|---|---|---|
| LS-9 | viable | 6 out of 6 |
| LS-10 | viable | 6 out of 6 |
| LS-11 | inviable | 6 out of 6 |
| LS-12 | inviable | 6 out of 6 |

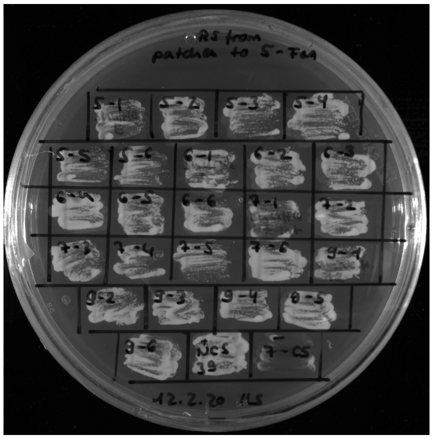

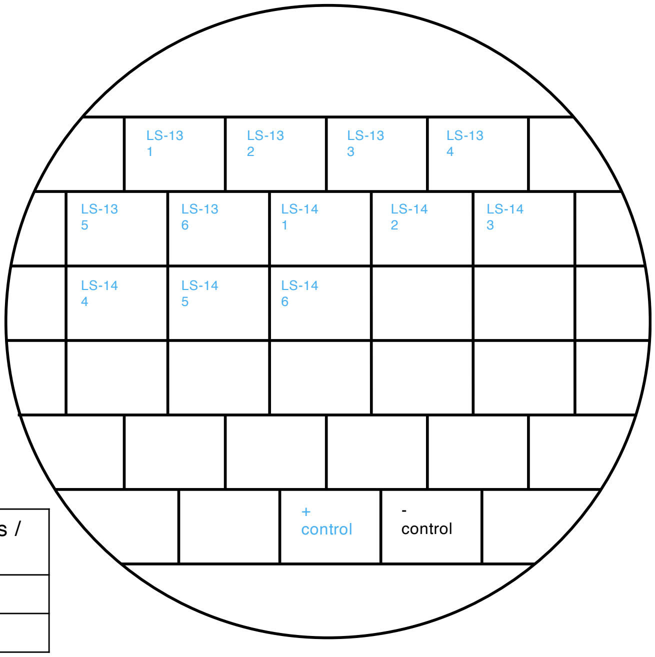


Restreak from YNB –ura –leu to 5-FOA –leu

| Construct | Construct | Number of (in)viable clones / total number of clones |
|---|---|---|
| LS-13 | viable | 6 out of 6 |
| LS-14 | viable | 6 out of 6 |

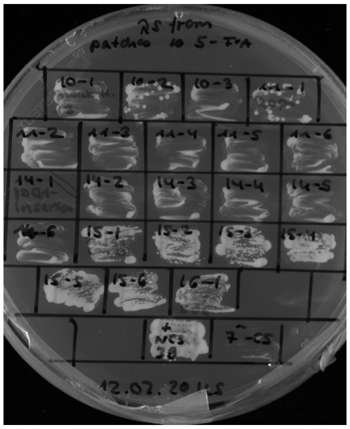

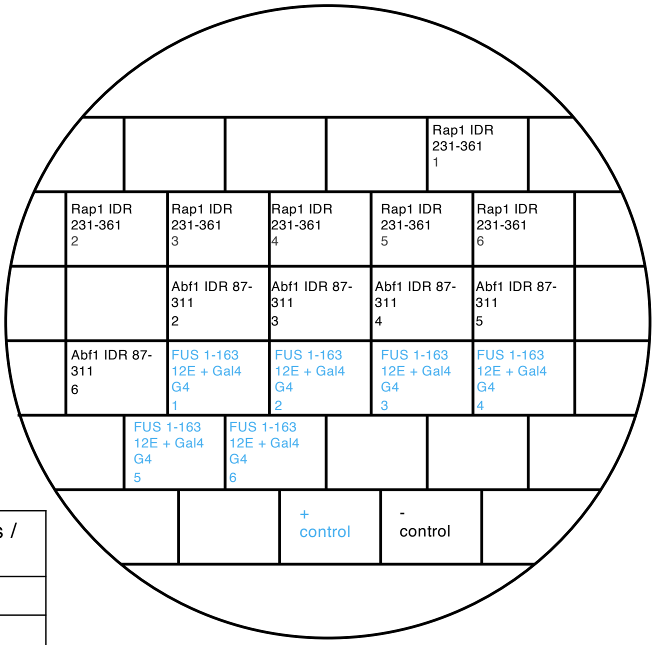


Restreak from YNB –ura –leu to 5-FOA –leu

| Construct | Result regarding viability on 5-FOA | Number of (in)viable clones / total number of clones |
|---|---|---|
| Rap1 IDR 231-361 | inviable | 6 out of 6 |
| Abf1 IDR 87-311 | inviable | 6 out of 6 |
| FUS 1-163 12E + Gal4 G4 | viable | 6 out of 6 |

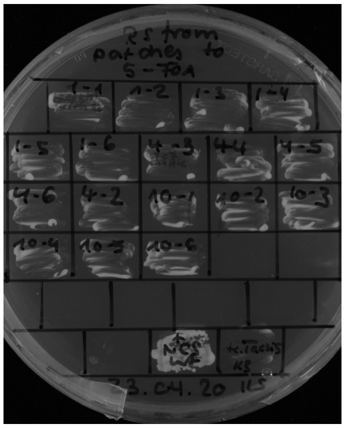

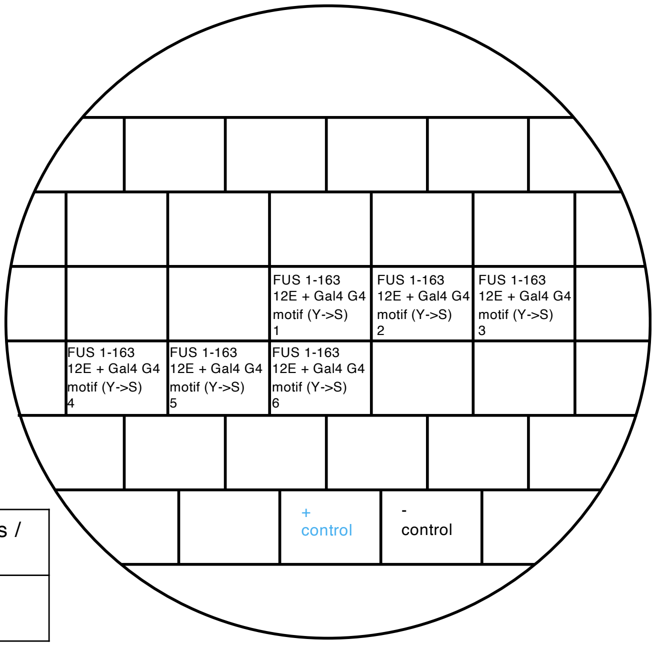


Restreak from YNB –ura –leu to 5-FOA –leu

| Construct | Construct | Number of (in)viable clones / total number of clones |
|---|---|---|
| FUS 1-163 12E + Gal4 G4 motif (Y->S) | inviable | 6 out of 6 |

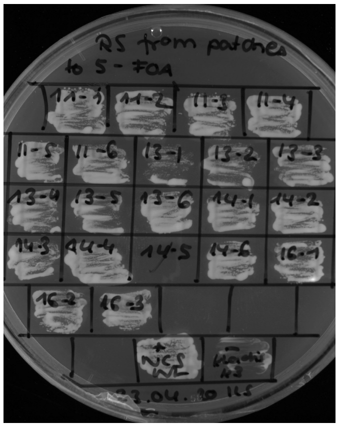

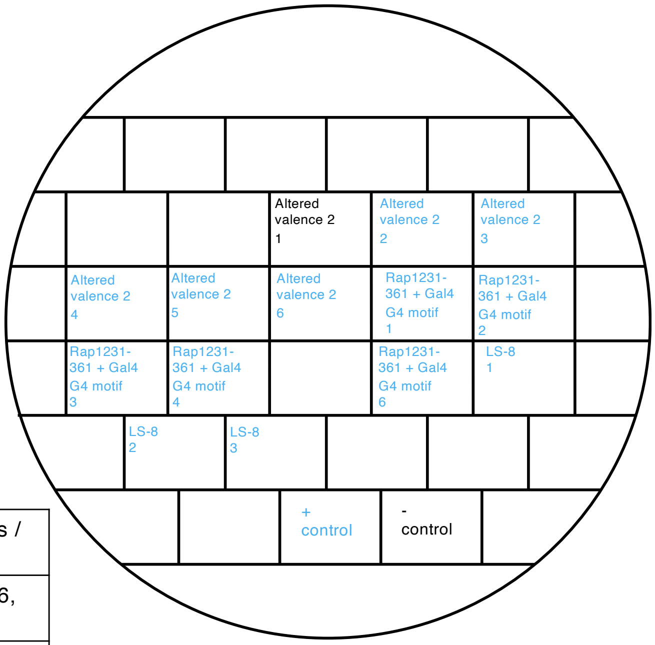


Restreak from YNB –ura –leu to 5-FOA –leu

| Construct | Construct | Number of (in)viable clones / total number of clones |
|---|---|---|
| Altered valence 2 | viable | 5 out of 6 (in total 5 out of 6, see also slide 22) |
| Rap1 231-361 + Gal4 G4 motif | viable | 5 out of 5 |
| LS-8 | viable | 3 out of 3 |

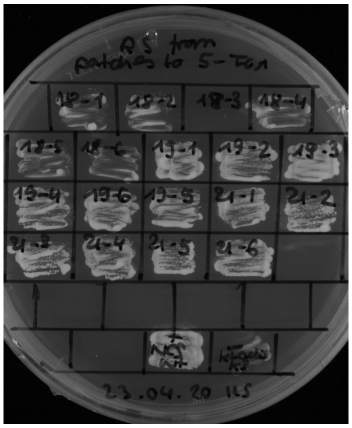

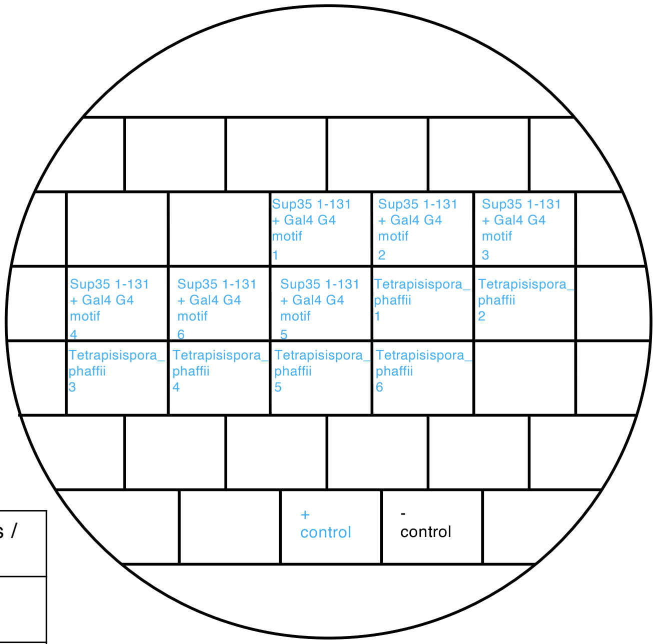


Restreak from YNB –ura –leu to 5-FOA –leu

| Construct | Result regarding viability on 5-FOA | Number of (in)viable clones / total number of clones |
|---|---|---|
| Sup35 1-131 + Gal4 G4 motif | viable | 6 out of 6 |
| Tetrapisispora_phaffii | viable | 6 out of 6 |

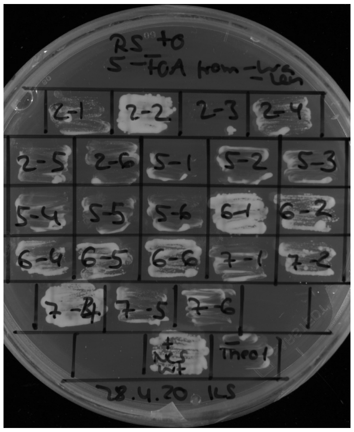

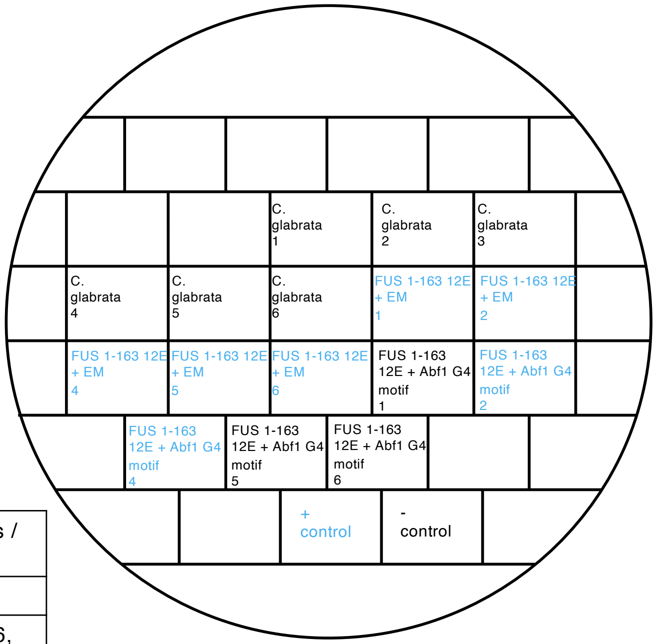


Restreak from YNB –ura –leu to 5-FOA –leu

| Construct | Result regarding viability on 5-FOA | Number of (in)viable clones / total number of clones |
|---|---|---|
| C. glabrata | inviable | 6 out of 6 |
| FUS 1-163 12E + EM | viable | 5 out of 5 (in total 6 out of 6, see also slides 11, 22, 23) |
| FUS 1-163 12E + Abf1 G4 motif | viable | 2 out of 5 (in total 2 out of 2, technical replicate, see also slide 11) |

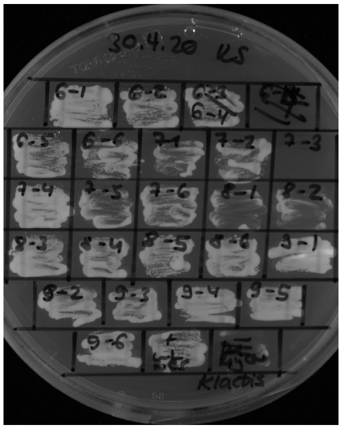


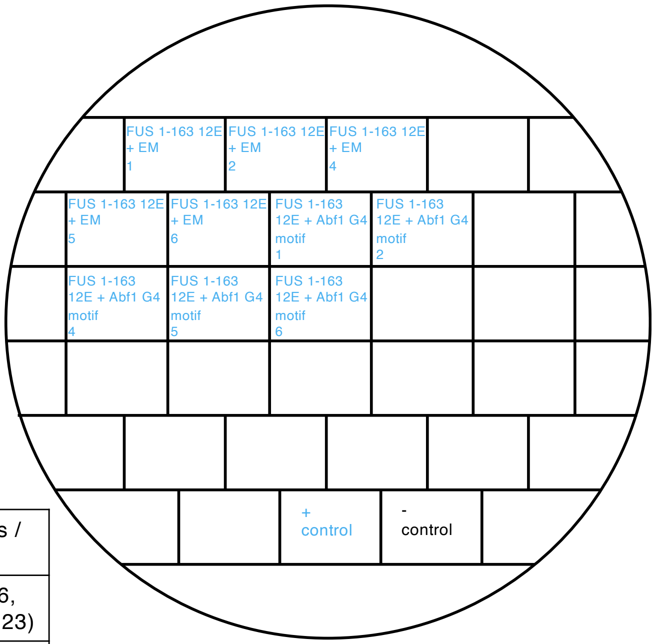


Restreak from YNB –ura –leu to 5-FOA –leu

| Construct | Construct | Number of (in)viable clones / total number of clones |
|---|---|---|
| FUS 1-163 12E + EM | viable | 5 out of 5 (in total 6 out of 6, see also slides 10, 22 and 23) |
| FUS 1-163 12E + Abf1 G4 motif | viable | 5 out of 5 (in total 2out of 2, technical replicate, see also slide 10) |

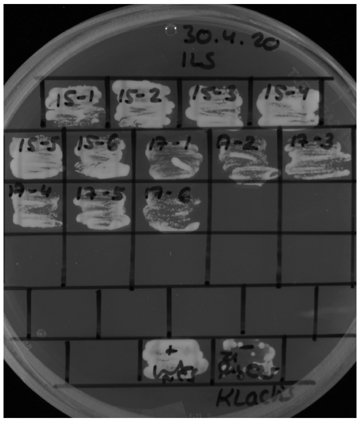


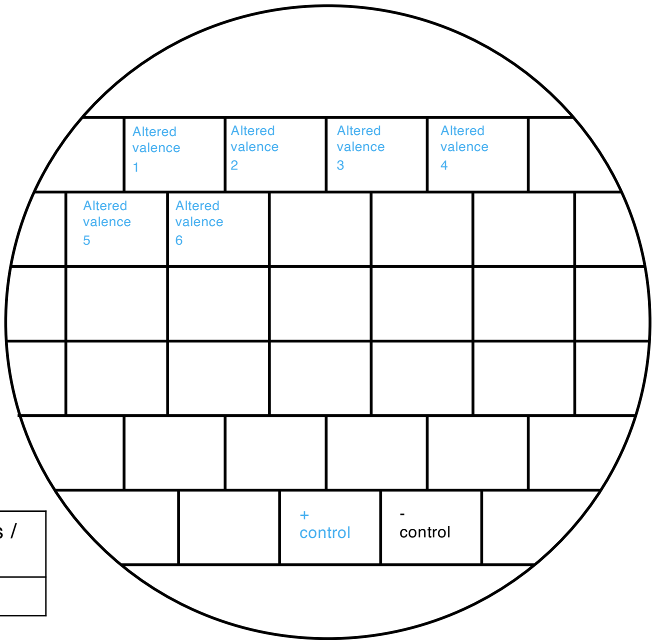


Restreak from YNB –ura –leu to 5-FOA –leu

| Construct | Result regarding viability on 5-FOA | Number of (in)viable clones / total number of clones |
|---|---|---|
| Altered valence 1 | viable | 6 out of 6 |

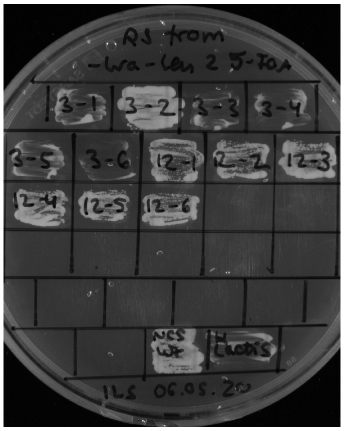


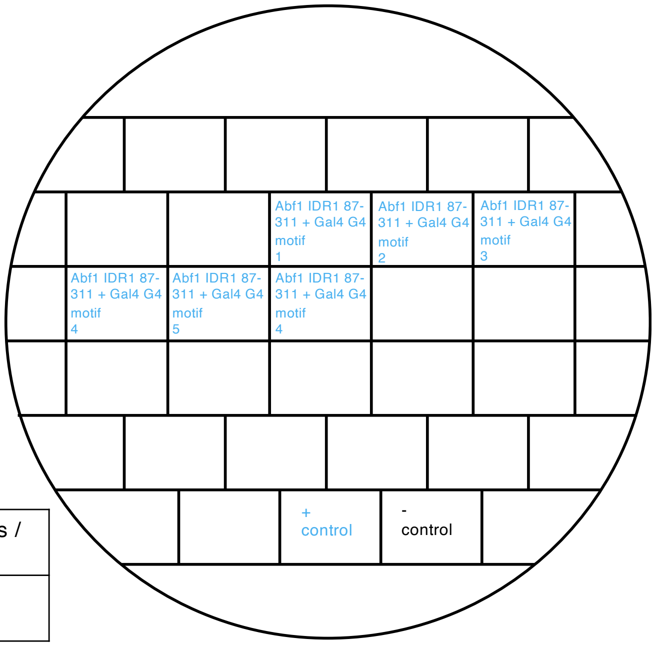


Restreak from YNB –ura –leu to 5-FOA –leu

| Construct | Construct | Number of (in)viable clones / total number of clones |
|---|---|---|
| Abf1 IDR1 87-311 + Gal4 G4 motif | viable | 6 out of 6 |

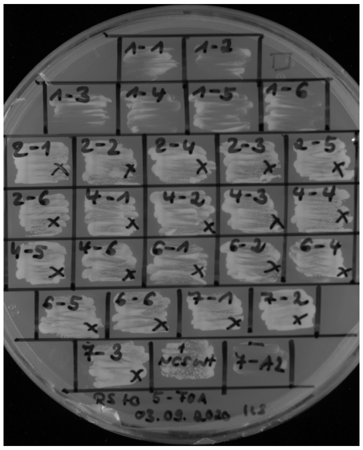


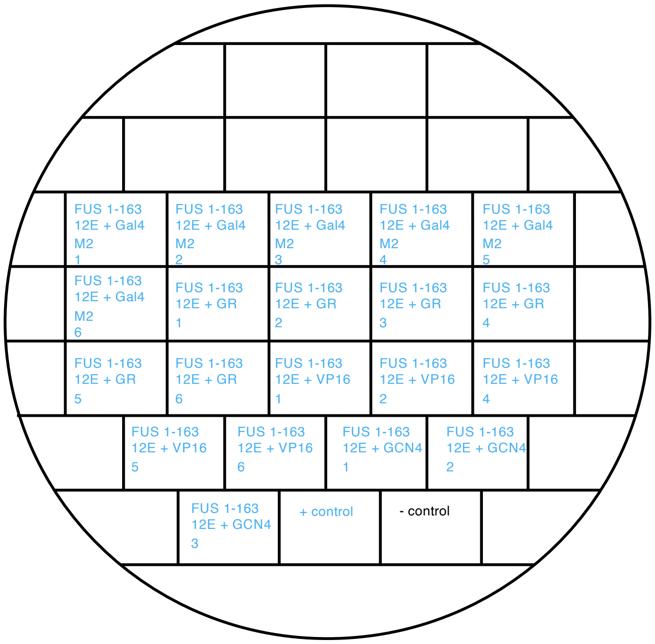


Restreak from YNB –ura –leu to 5-FOA –leu

| Construct | Result regarding viability on 5-FOA | Number of (in)viable clones / total number of clones |
|---|---|---|
| FUS 1-163 12E + Gal4 M2 | viable | 6 out of 6 |
| FUS 1-163 12E + GR | viable | 6 out of 6 |
| FUS 1-163 12E + VP16 | viable | 5 out of 5 |
| FUS 1-163 12E + GCN4 | viable | 3 out of 3 (in total 6 out of 6, see also slide 13) |

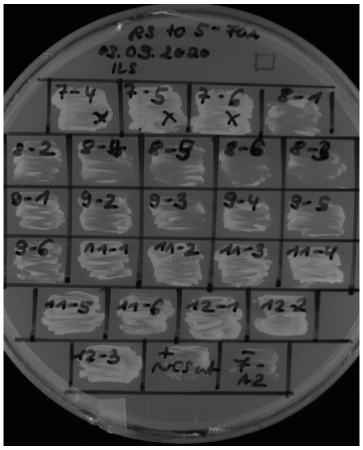


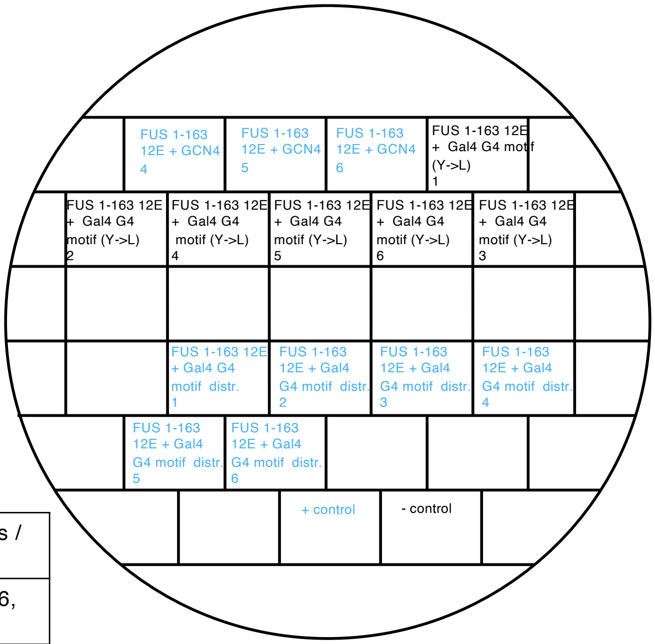


Restreak from YNB –ura –leu to 5-FOA –leu

| Construct | Result regarding viability on 5-FOA | Number of (in)viable clones / total number of clones |
|---|---|---|
| FUS 1-163 12E + GCN4 | viable | 3 out of 3 (in total 6 out of 6, see also slide 12) |
| FUS 1-163 12E + Gal4 G4 motif (Y->L) | inviable | 6 out of 6 |
| FUS 1-163 12E + Gal4 G4 motif distr. | viable | 6 out of 6 |

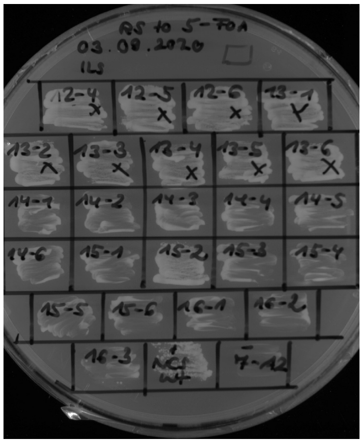


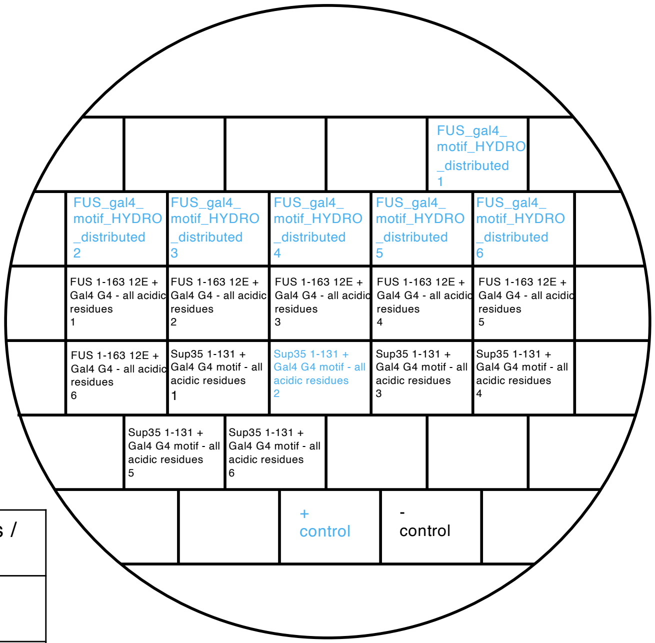


Restreak from YNB –ura –leu to 5-FOA –leu

| Construct | Result regarding viability on 5-FOA | Number of (in)viable clones / total number of clones |
|---|---|---|
| FUS_gal4_motif_HYDRO_distributed | viable | 6 out of 6 |
| FUS 1-163 12E + Gal4 G4 - all acidic residues | inviable | 6 out of 6 |
| Sup35 1-131 + Gal4 G4 motif - all acidic residues | inviable | 5 out of 6 |

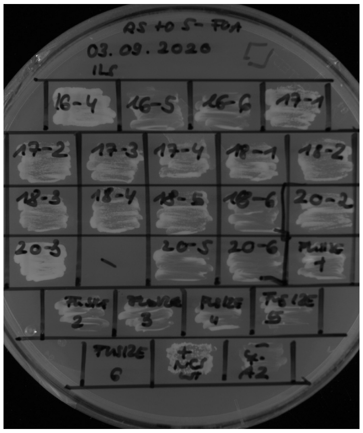

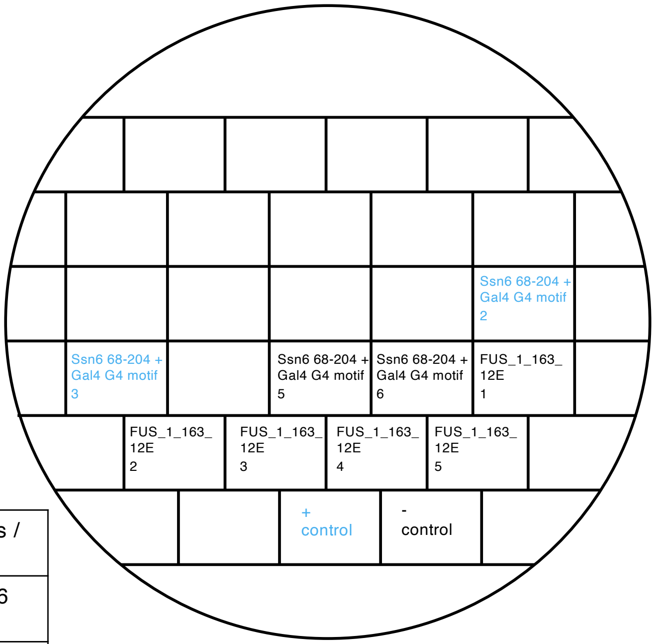


Restreak from YNB –ura –leu to 5-FOA –leu

| Construct | Result regarding viability on 5-FOA | Number of (in)viable clones / total number of clones |
|---|---|---|
| Ssn6 68-204 + Gal4 G4 motif | inviable | 2 out of 4 (in total 4 out of 6 see plate below) |
| FUS_1_163_12E | inviable | 5 out of 5 (in total 8 out of 9, see also slides 6 and 7) |

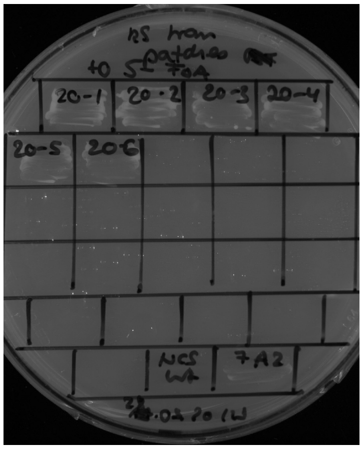

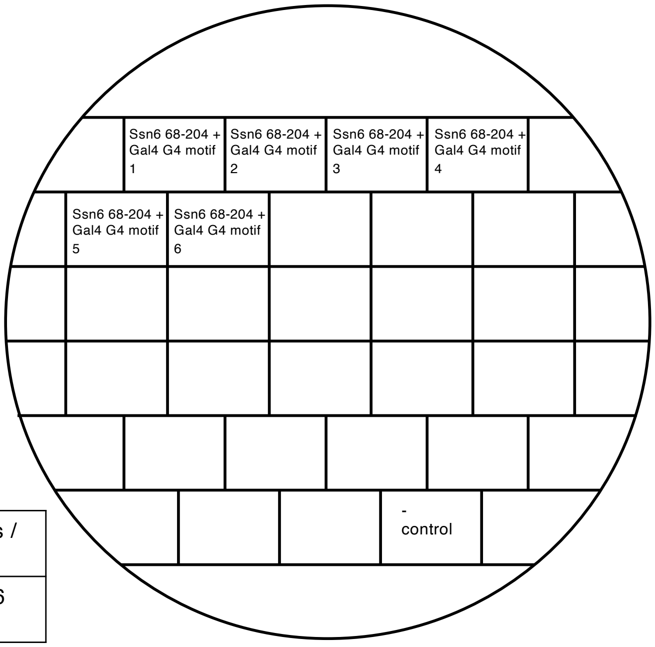


Restreak from YNB –ura –leu to 5-FOA –leu

| Construct | Result regarding viability on 5-FOA | Number of (in)viable clones / total number of clones |
|---|---|---|
| Ssn6 68-204 + Gal4 G4 motif | Inviable | 6 out of 6 (in total 4 out of 6 see plate above) |

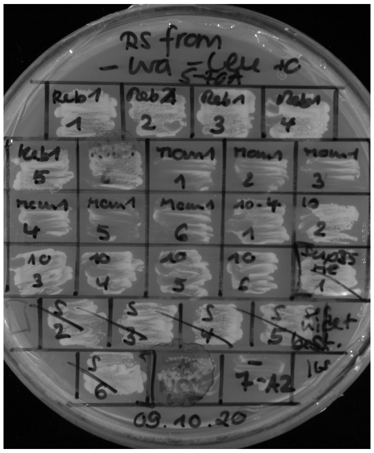

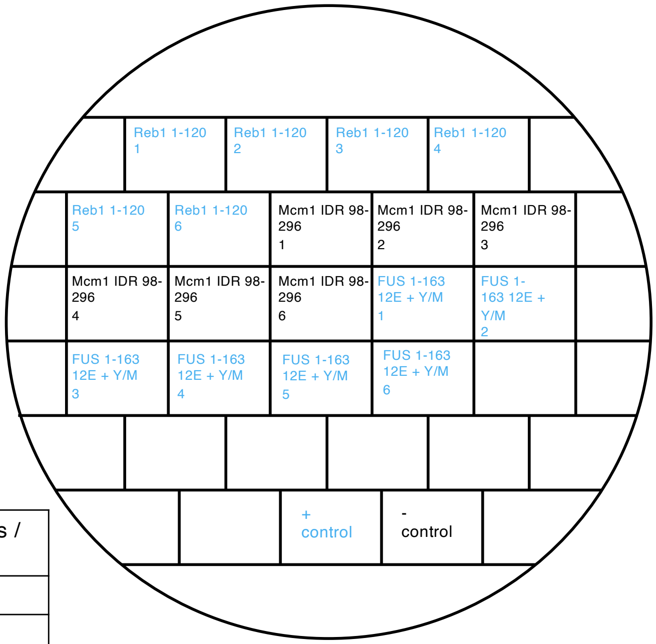


Restreak from YNB –ura –leu to 5-FOA –leu

| Construct | Result regarding viability on 5-FOA | Number of (in)viable clones / total number of clones |
|---|---|---|
| Reb1 1-120 (frameshift) | viable | 6 out of 6 |
| Mcm1 IDR 98-296 | inviable | 6 out of 6 |
| FUS 1-163 12E + Y/M | viable | 6 out of 6 |

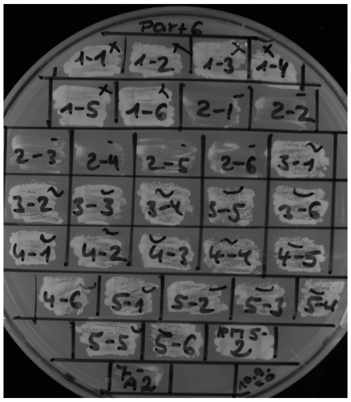

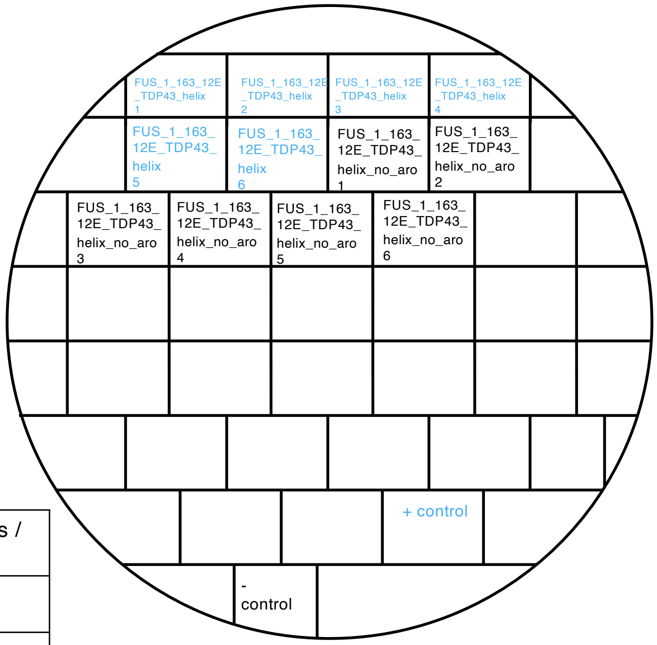


Restreak from YNB –ura –leu to 5-FOA –leu

| Construct | Result regarding viability on 5-FOA | Number of (in)viable clones / total number of clones |
|---|---|---|
| FUS_1_163_12E_TDP43_helix | viable | 6 out of 6 |
| FUS_1_163_12E_TDP43_helix_no_aro | inviable | 6 out of 6 |

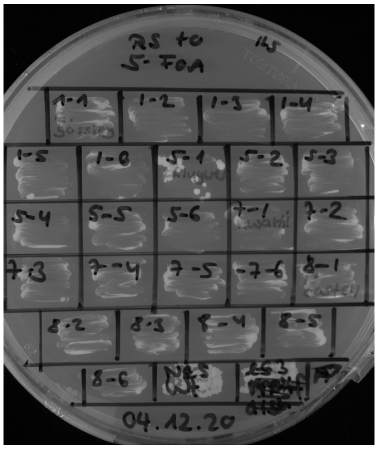


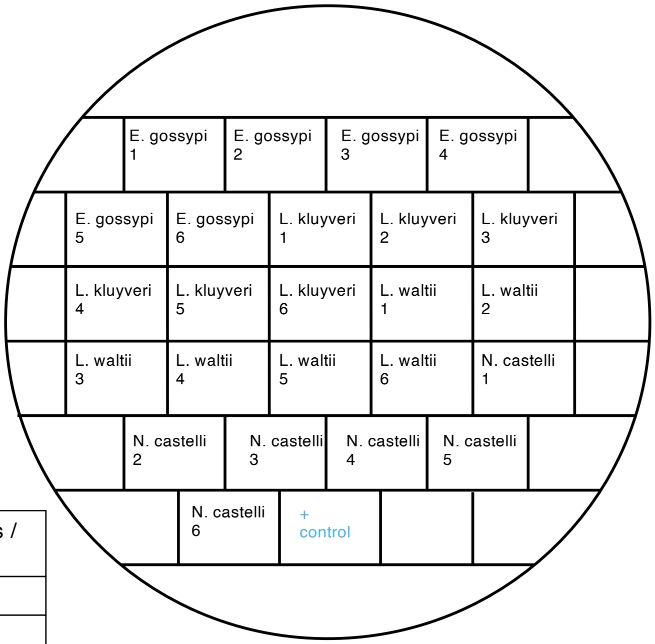


Restreak from YNB –ura –leu to 5-FOA –leu

| Construct | Result regarding viability on 5-FOA | Number of (in)viable clones / total number of clones |
|---|---|---|
| E. gossypi | inviable | 6 out of 6 |
| L. kluyveri | inviable | 6 out of 6 |
| L. waltii | inviable | 6 out of 6 |
| N. castelli | inviable | 6 out of 6 |

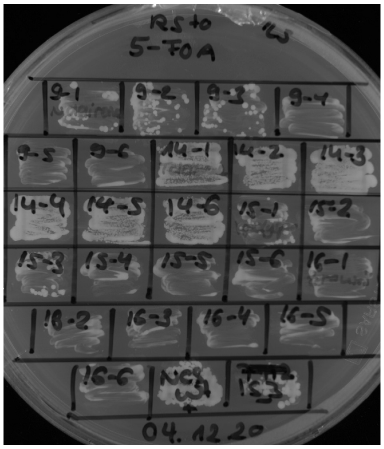


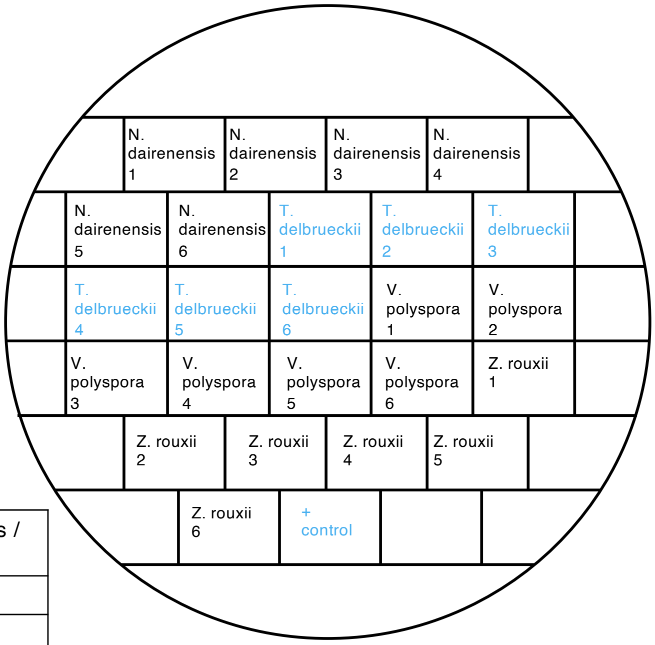


Restreak from YNB –ura –leu to 5-FOA –leu

| Construct | Result regarding viability on 5-FOA | Number of (in)viable clones / total number of clones |
|---|---|---|
| N. dairenensis | inviable | 6 out of 6 |
| T. delbrueckii | viable | 6 out of 6 |
| V. polyspora | inviable | 6 out of 6 |
| Z. rouxii | inviable | 6 out of 6 |

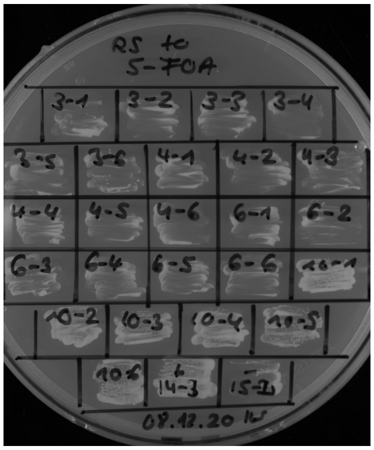


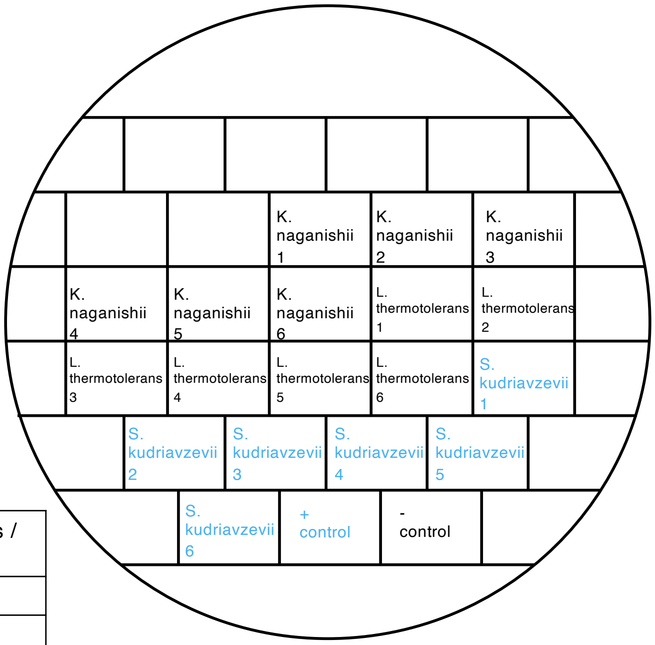


Restreak from YNB –ura –leu to 5-FOA –leu

| Construct | Result regarding viability on 5-FOA | Number of (in)viable clones / total number of clones |
|---|---|---|
| K. naganishii | inviable | 6 out of 6 |
| L. thermotolerans | inviable | 6 out of 6 |
| S. kudriavzevii | viable | 6 out of 6 |

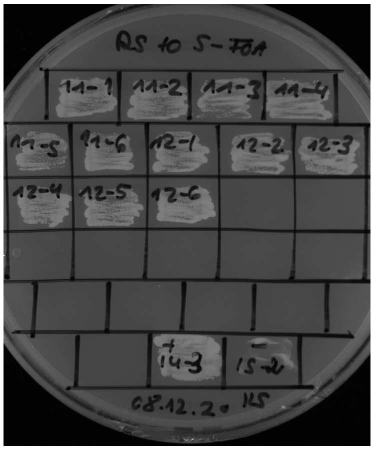


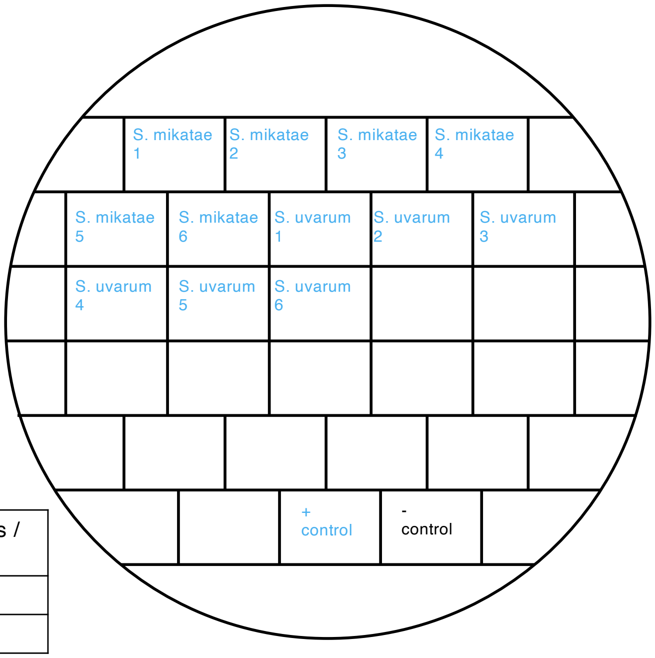


Restreak from YNB –ura –leu to 5-FOA –leu

| Construct | Result regarding viability on 5-FOA | Number of (in)viable clones / total number of clones |
|---|---|---|
| S. mikatae | viable | 6 out of 6 |
| S. uvarum | Viable | 6 out of 6 |

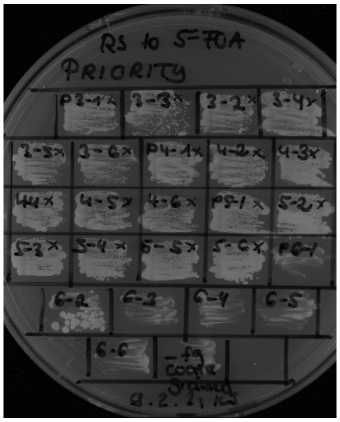


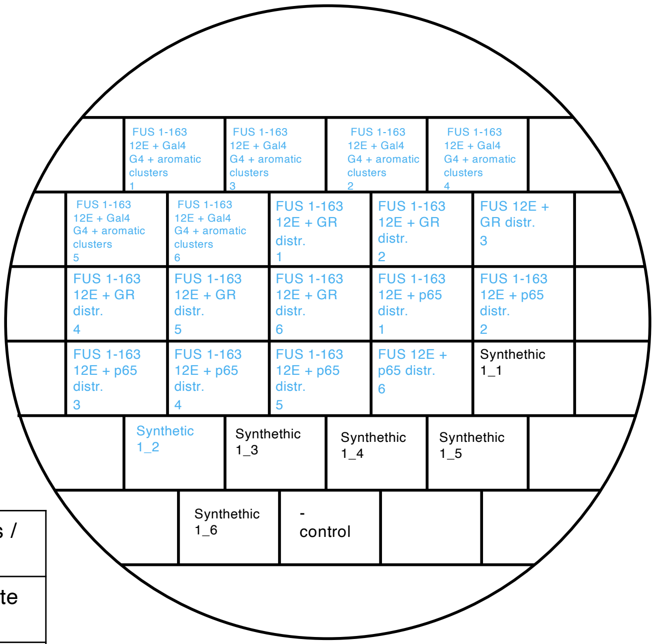


Restreak from YNB –ura –leu to 5-FOA –leu

| Construct | Result regarding viability on 5-FOA | Number of (in)viable clones / total number of clones |
|---|---|---|
| FUS 1-163 12E + Gal4 G4 + aromatic clusters | viable | 6 out of 6 (technical replicate see slide 22) |
| FUS 1-163 12E + GR distr. | viable | 6 out of 6 |
| FUS 1-163 12E + p65 distr. | viable | 6 out of 6 |
| Synthetic 1 | inviable | 5 out of 6 |

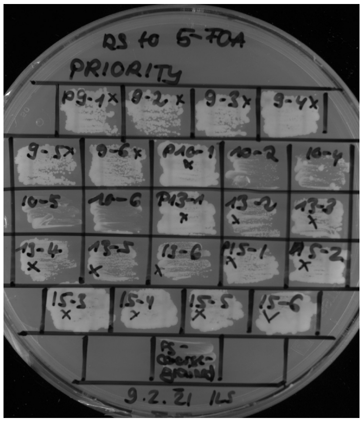


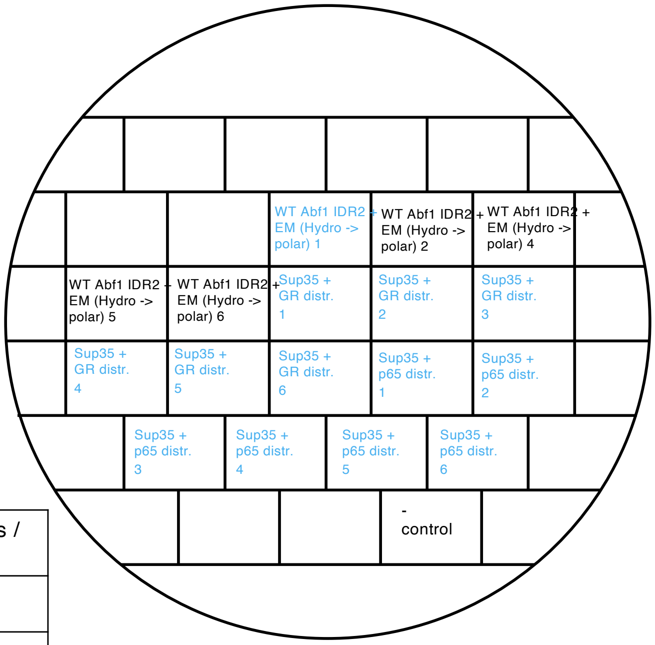


Restreak from YNB –ura –leu to 5-FOA –leu

| Construct | Result regarding viability on 5-FOA | Number of (in)viable clones / total number of clones |
|---|---|---|
| WT Abf1 IDR2 + EM (Hydro -> polar) | inviable | 4 out of 5 |
| Sup35 + GR distr. | viable | 6 out of 6 |
| Sup35 + p65 distr. | viable | 6 out of 6 |

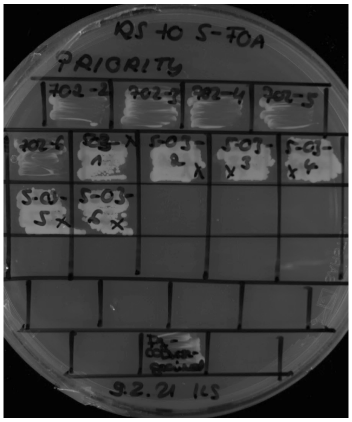

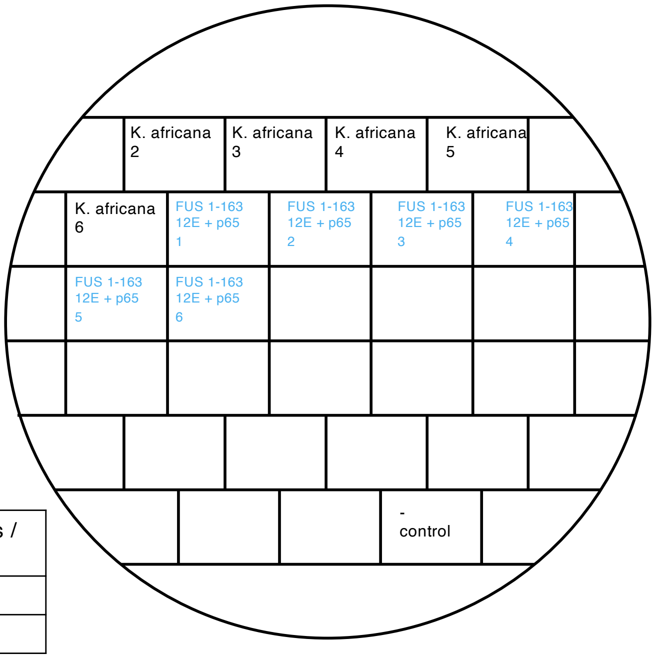


Restreak from YNB –ura –leu to 5-FOA –leu

| Construct | Result regarding viability on 5-FOA | Number of (in)viable clones / total number of clones |
|---|---|---|
| K. africana | inviable | 5 out of 5 |
| FUS 1-163 12E + p65 | viable | 6 out of 6 |

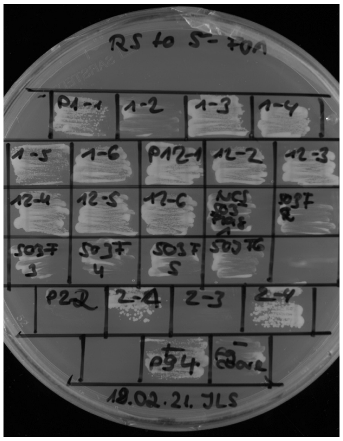

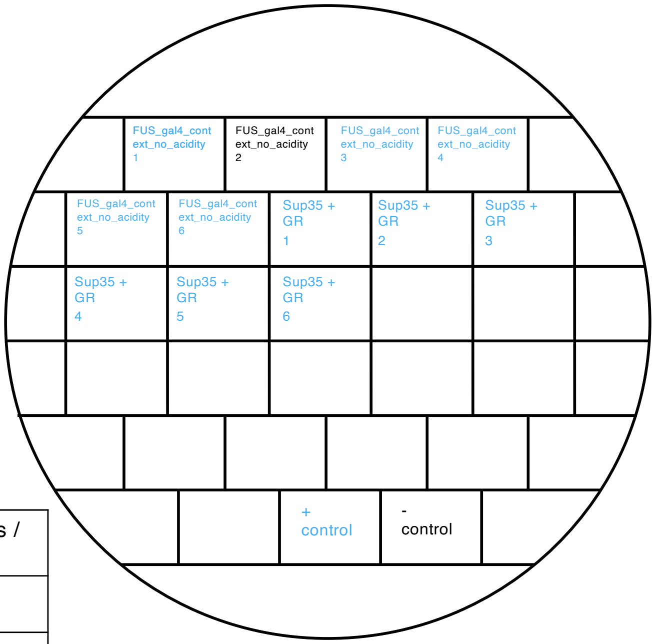


Restreak from YNB –ura –leu to 5-FOA –leu

| Construct | Result regarding viability on 5-FOA | Number of (in)viable clones / total number of clones |
|---|---|---|
| fus_gal4_context_no_acidity | viable | 5 out of 6 |
| Sup35 + GR | viable | 6 out of 6 |

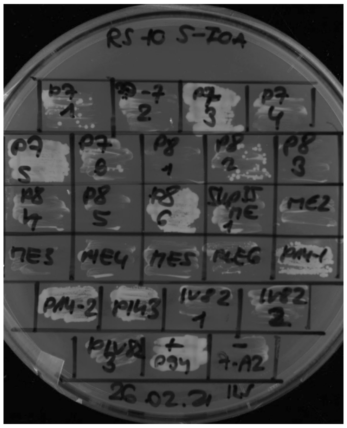

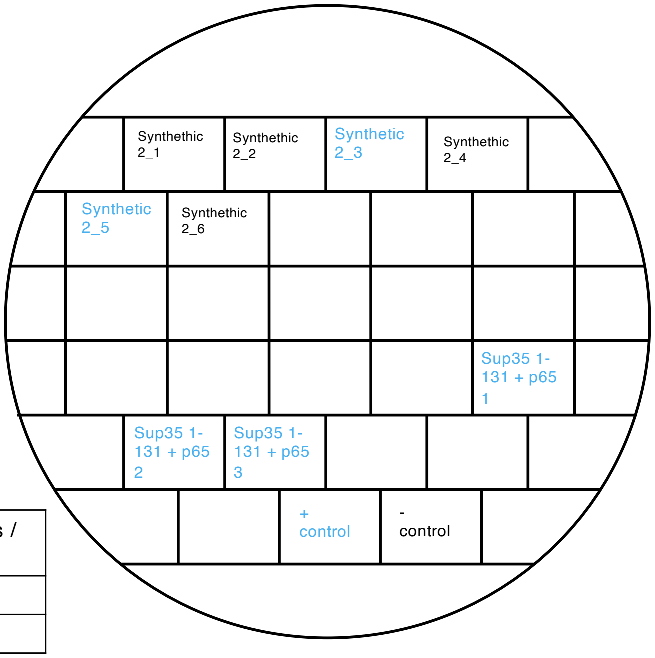


Restreak from YNB –ura –leu to 5-FOA –leu

| Construct | Result regarding viability on 5-FOA | Number of (in)viable clones / total number of clones |
|---|---|---|
| Synthetic 2 | inviable | 4 out of 6 |
| Sup35 1-131 + p65 | viable | 3 out of 3 |

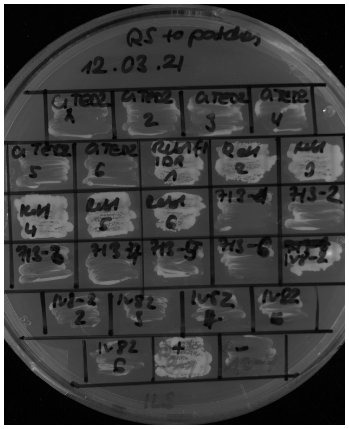

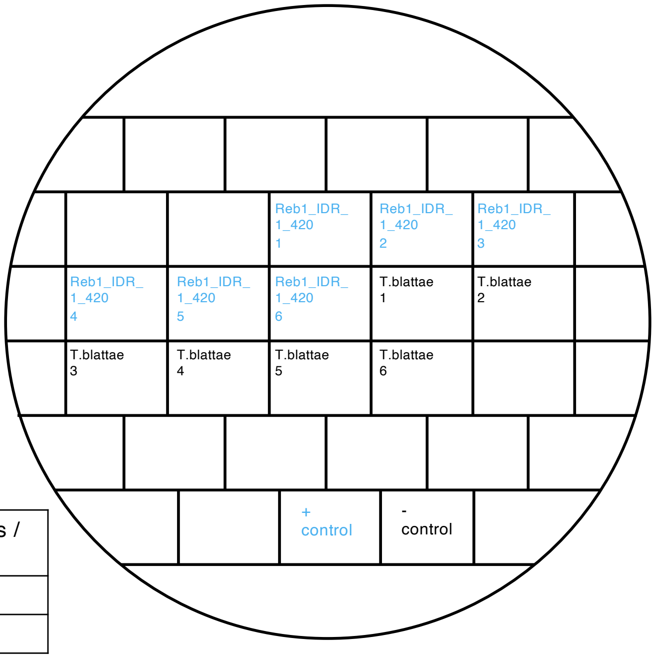


Restreak from YNB –ura –leu to 5-FOA –leu

| Construct | Result regarding viability on 5-FOA | Number of (in)viable clones / total number of clones |
|---|---|---|
| Reb1_IDR_1_420 | viable | 6 out of 6 |
| T.blattae | inviable | 6 out of 6 |

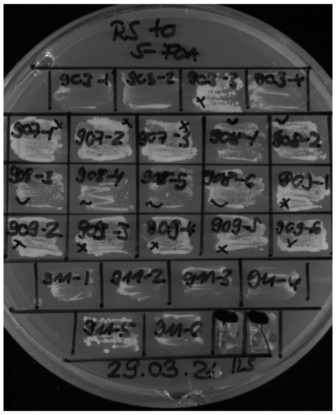


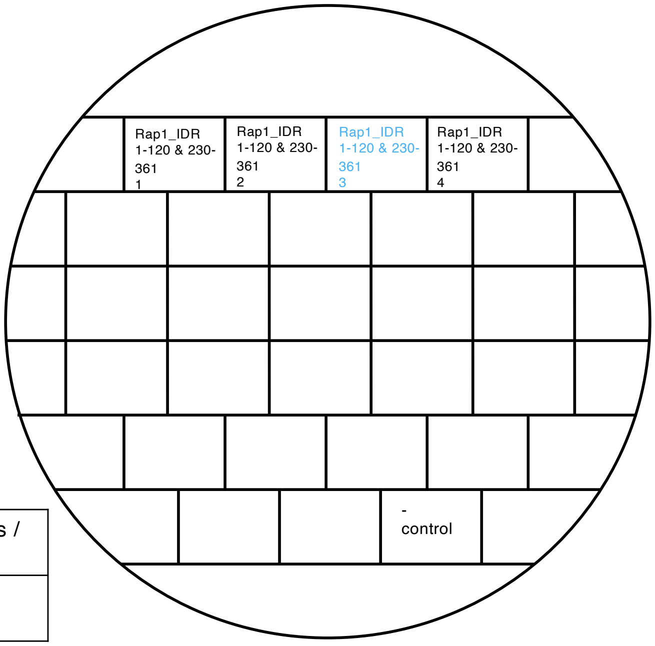


Restreak from YNB –ura –leu to 5-FOA –leu

| Construct | Result regarding viability on 5-FOA | Number of (in)viable clones / total number of clones |
|---|---|---|
| Rap1_IDR 1-120 & 230-361 | inviable | 3 out of 4 |

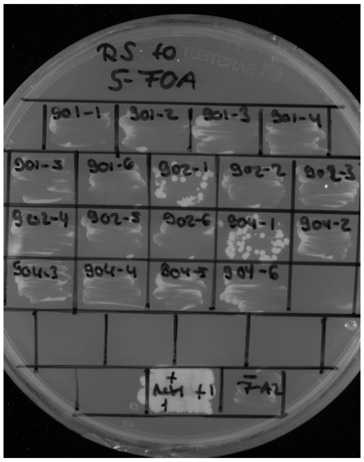


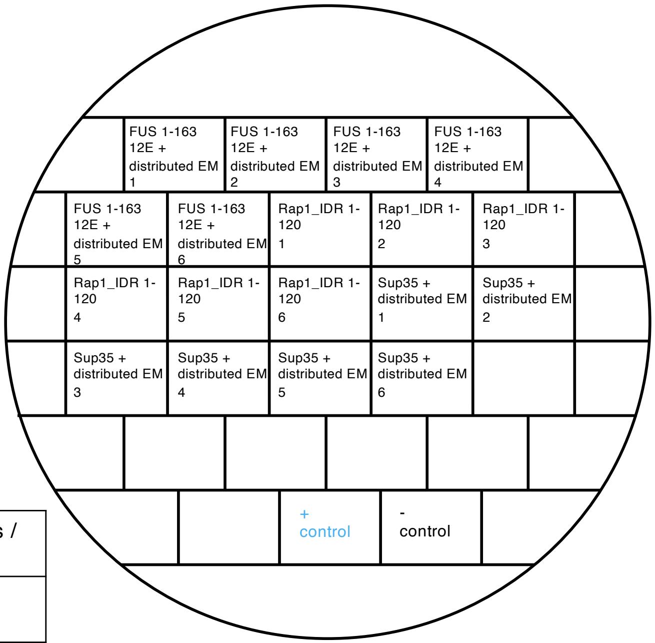


Restreak from YNB –leu to 5-FOA –leu

| Construct | Result regarding viability on 5-FOA | Number of (in)viable clones / total number of clones |
|---|---|---|
| FUS 1-163 12E + distributed EM | inviable | 6 out of 6 |
| Rap1_IDR1 1-120 | inviable | 6 out of 6 |
| Sup35 + distributed EM | inviable | 6 out of 6 |

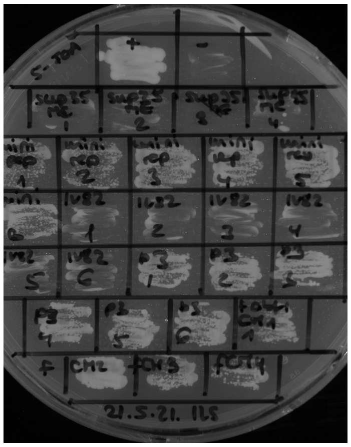

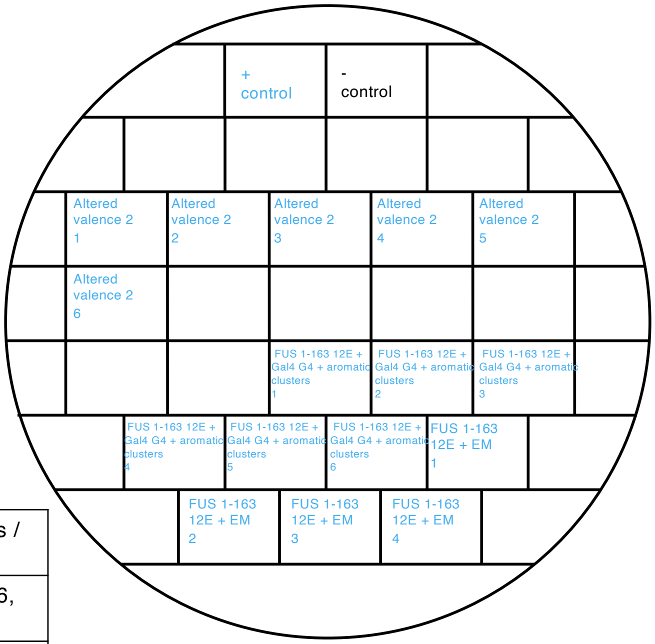


Restreak from YNB –ura –leu to 5-FOA –leu

| Construct | Result regarding viability on 5-FOA | Number of (in)viable clones / total number of clones |
|---|---|---|
| Altered valence 2 | viable | 6 out of 6 (in total 5 out of 6, see also slide 9) |
| FUS 1-163 12E + Gal4 G4 + aromatic clusters | viable | 6 out of 6 (technical replicate see slide 18) |
| FUS 1-163 12E + EM | viable | 4 out of 4 (in total 6 out of 6, see also slides 10, 11, 23) |

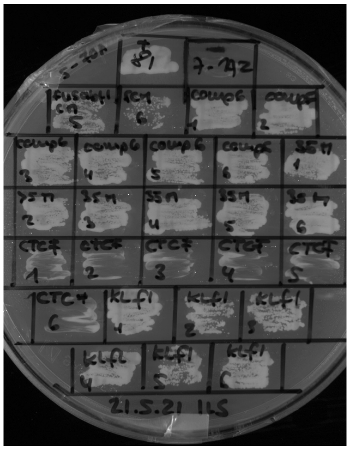

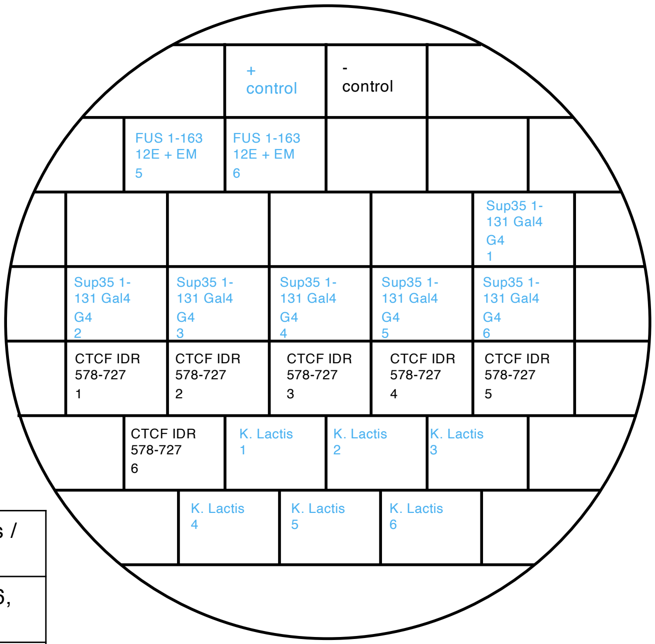


Restreak from YNB –ura –leu to 5-FOA –leu

| Construct | Result regarding viability on 5-FOA | Number of (in)viable clones / total number of clones |
|---|---|---|
| FUS 1-163 12E + EM | viable | 2 out of 2 (in total 6 out of 6, see also slides 10, 11, 22) |
| Sup35 1-131 + Gal4 G4 motif | viable | 6 out of 6 |
| CTCF IDR 578-727 | inviable | 6 out of 6 (technical repliate, see slide 24) |
| K. Lactis | viable | 6 out of 6 |

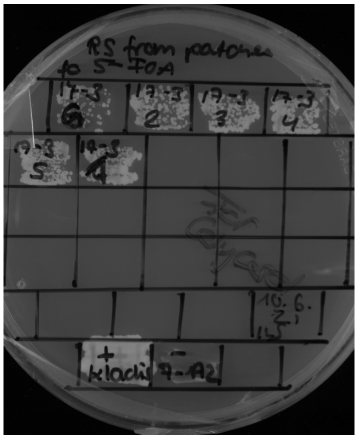

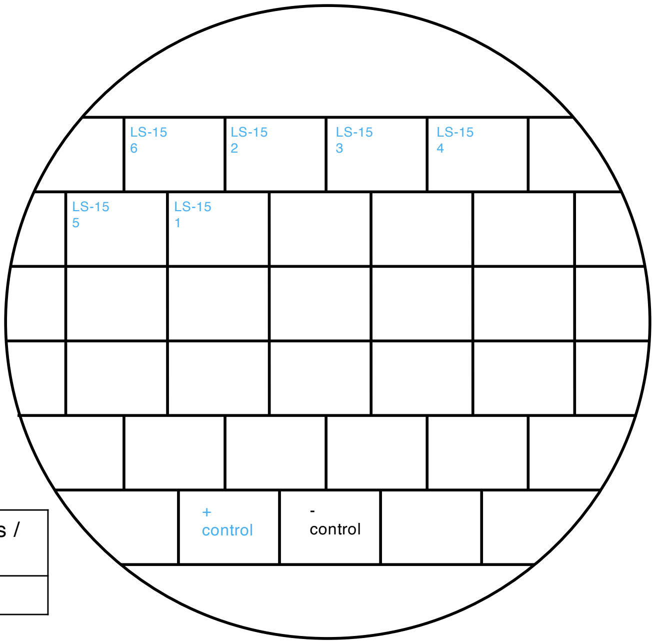


Restreak from YNB –ura –leu to 5-FOA –leu

| Construct | Result regarding viability on 5-FOA | Number of (in)viable clones / total number of clones |
|---|---|---|
| LS-15 | viable | 6 out of 6 |

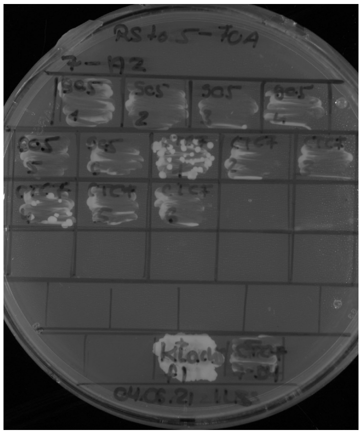

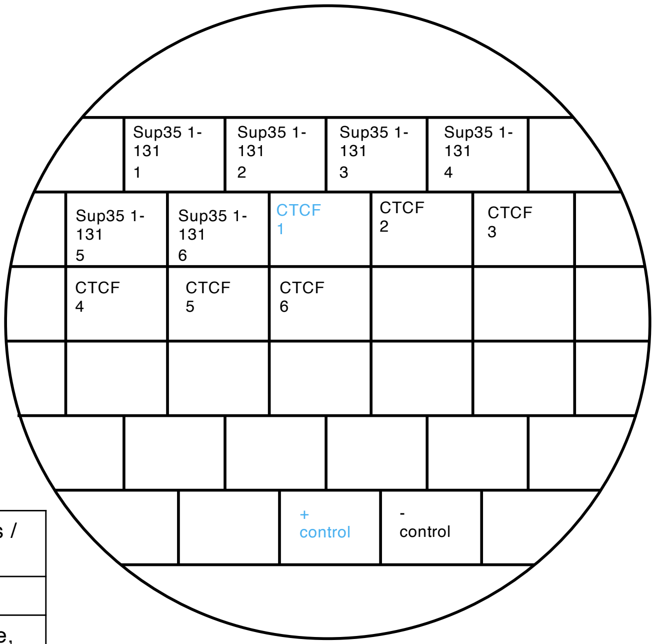


Restreak from YNB –ura –leu to 5-FOA –leu

| Construct | Result regarding viability on 5-FOA | Number of (in)viable clones / total number of clones |
|---|---|---|
| Sup35 1-131 | inviable | 6 out of 6 |
| CTCF IDR 578-727 | inviable | 5 out of 6 (technical repliate, see slide 23) |

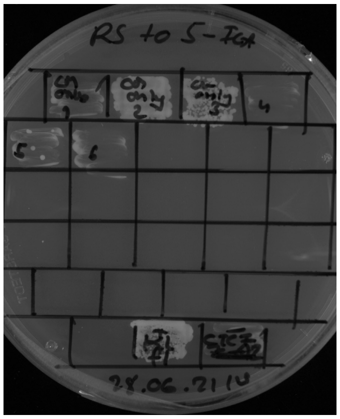

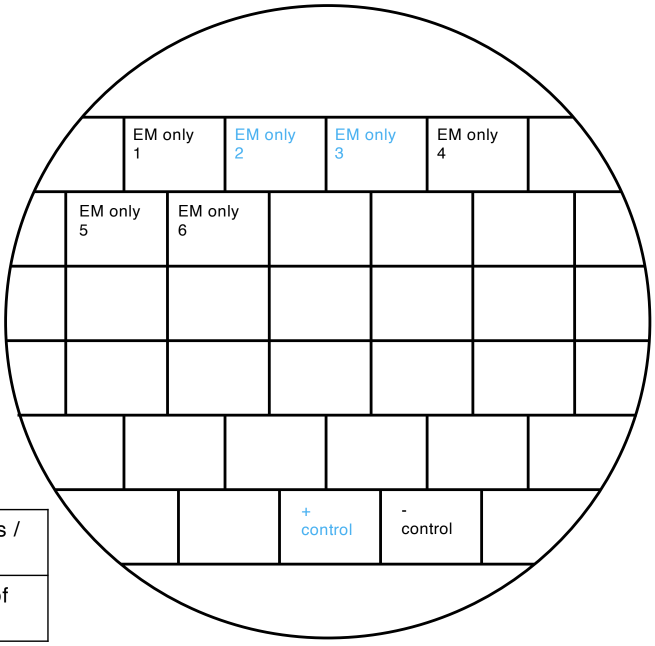


Restreak from YNB –ura –leu to 5-FOA –leu

| Construct | Result regarding viability on 5-FOA | Number of (in)viable clones / total number of clones |
|---|---|---|
| EM only | inviable | 4 out of 6 (in total: 10 out of 12, see also slide 25) |

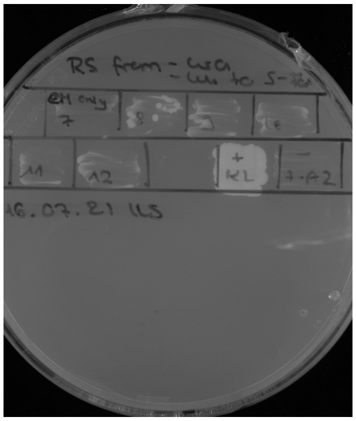

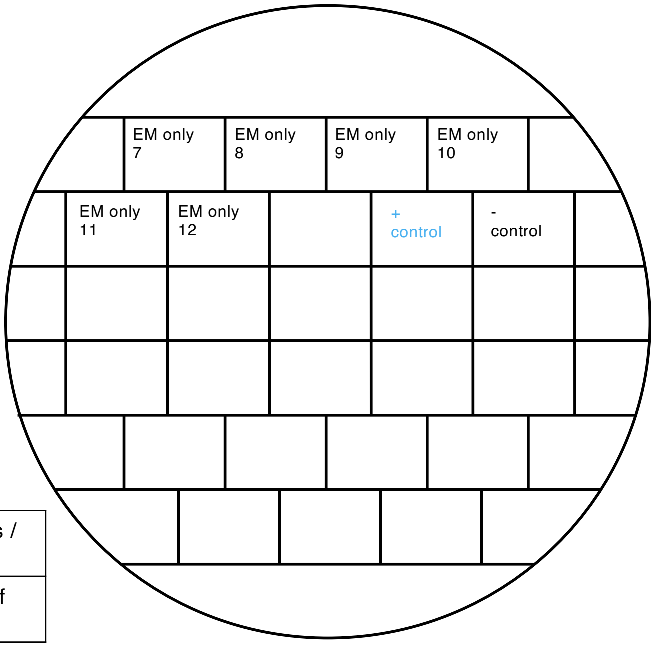


Restreak from YNB –ura –leu to 5-FOA –leu

| Construct | Result regarding viability on 5-FOA | Number of (in)viable clones / total number of clones |
|---|---|---|
| EM only | Inviable | 6 out of 6 (in total: 10 out of 12, see also slide 24) |

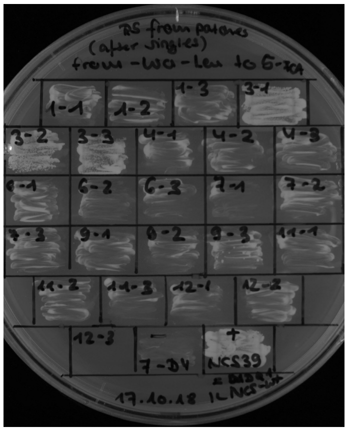

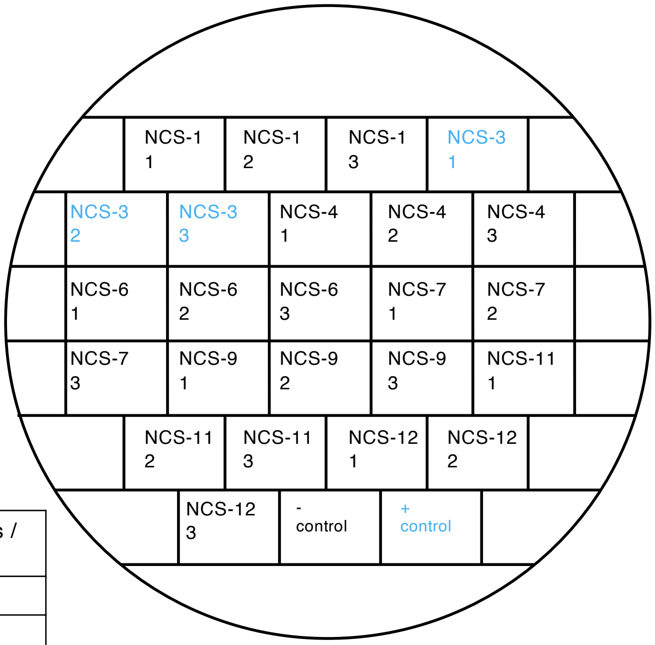


Restreak from YNB –ura –leu to 5-FOA –leu

| Construct | Result regarding viability on 5-FOA | Number of (in)viable clones / total number of clones |
|---|---|---|
| NCS-1 | inviable | 3 out of 3 |
| NCS-3 | viable | 3 out of 3 |
| NCS-4 | inviable | 3 out of 3 |
| NCS-6 | inviable | 3 out of 3 |
| NCS-7 | inviable | 3 out of 3 |
| NCS-9 | inviable | 3 out of 3 |
| NCS-11 | inviable | 3 out of 3 |
| NCS-12 | inviable | 3 out of 3 |

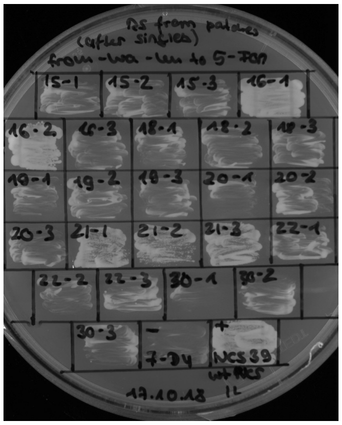

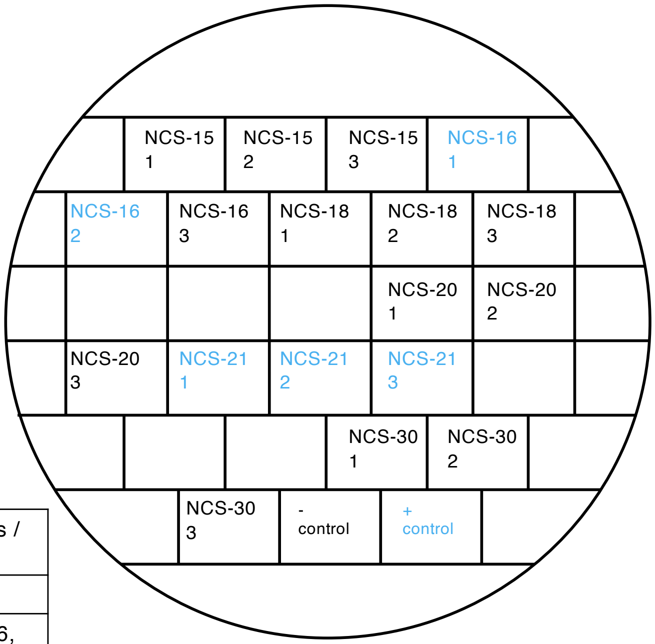


Restreak from YNB –ura –leu to 5-FOA –leu

| Construct | Result regarding viability on 5-FOA | Number of (in)viable clones / total number of clones |
|---|---|---|
| NCS-15 | inviable | 3 out of 3 |
| NCS-16 | inviable | 1 out of 3 (in total 4 out of 6, see slide 27) |
| NCS-18 | inviable | 3 out of 3 |
| NCS-20 | inviable | 3 out of 3 |
| NCS-21 | viable | 3 out of 3 |
| NCS-30 | inviable | 3 out of 3 |

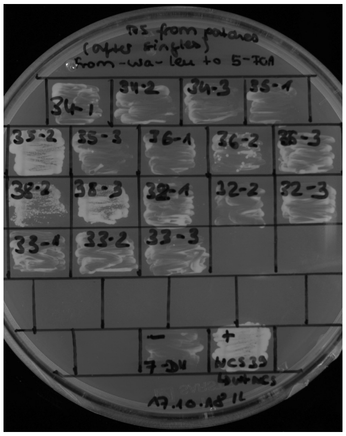

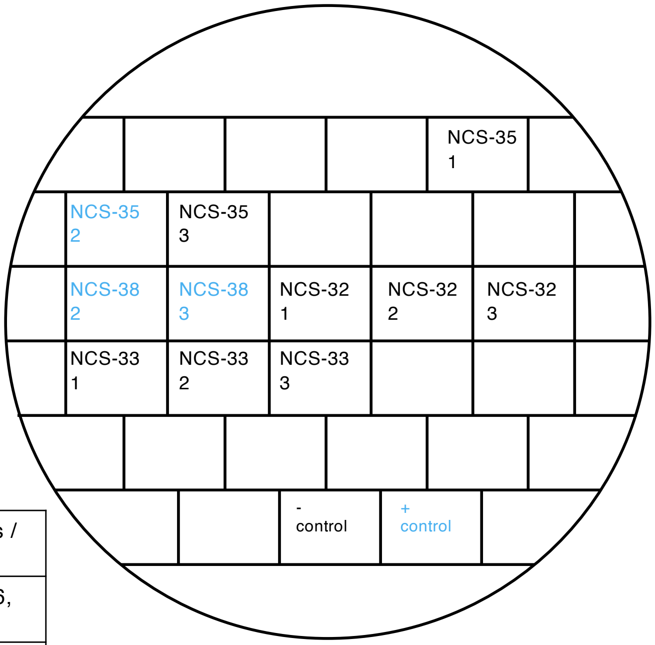


Restreak from YNB –ura –leu to 5-FOA –leu

| Construct | Result regarding viability on 5-FOA | Number of (in)viable clones / total number of clones |
|---|---|---|
| NCS-35 | inviable | 2 out of 3 (in total 5 out of 6, see also slide 26) |
| NCS-38 | viable | 2 out of 2 (in total 5 out of 6, see also slide 26) |
| NCS-32 | inviable | 3 out of 3 |
| NCS-33 | inviable | 3 out of 3 |

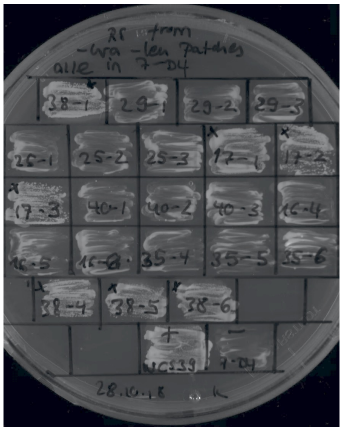

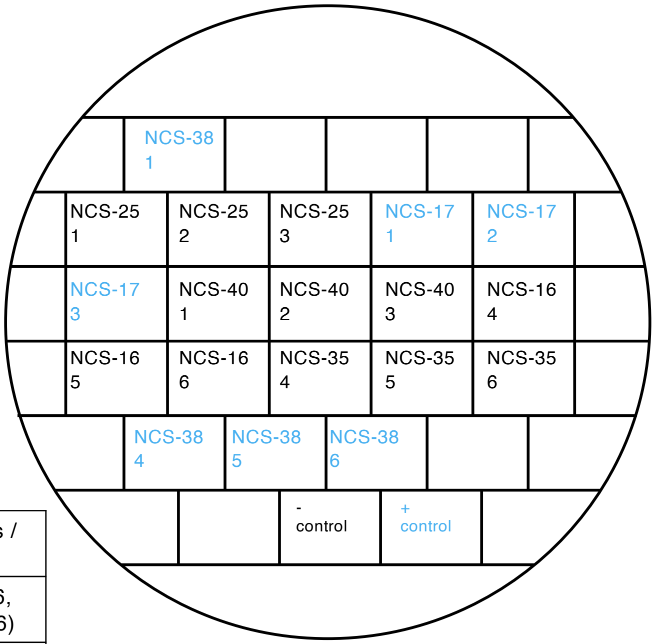


Restreak from YNB –ura –leu to 5-FOA –leu

| Construct | Result regarding viability on 5-FOA | Number of (in)viable clones / total number of clones |
|---|---|---|
| NCS-38 | viable | 4 out of 4 (in total 5 out of 6, see also below and slide 26) |
| NCS-25 | inviable | 3 out of 3 |
| NCS-17 | viable | 3 out of 3 |
| NCS-40 | inviable | 3 out of 3 |
| NCS-16 | inviable | 3 out of 3 (in total 4 out of 6, see below and slide 26) |
| NCS-35 | inviable | 3 out of 3 (in total 5 out of 6, see also below and slide 26) |

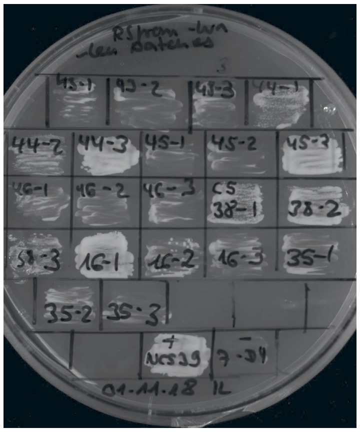

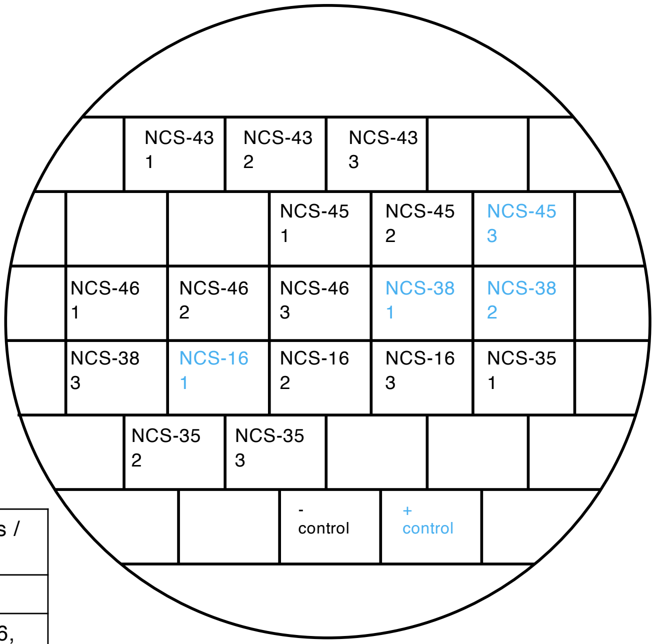


Restreak from YNB –ura –leu to 5-FOA –leu

| Construct | Result regarding viability on 5-FOA | Number of (in)viable clones / total number of clones |
|---|---|---|
| NCS-43 | inviable | 3 out of 3 |
| NCS-45 | inviable | 2 out of 3 (in total 5 out of 6, see also slide 29) |
| NCS-46 | inviable | 3 out of 3 |
| NCS-38 | viable | 2 out of 3 (in total 5 out of 6, see also above and slide 26) |
| NCS-16 | inviable | 2 out of 3 (in total 4 out of 6, see above and slide 26) |
| NCS-35 | inviable | 3 out of 3 (in total 5 out of 6, see also above and slide 26) |

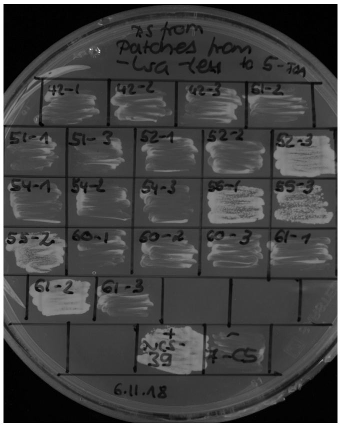

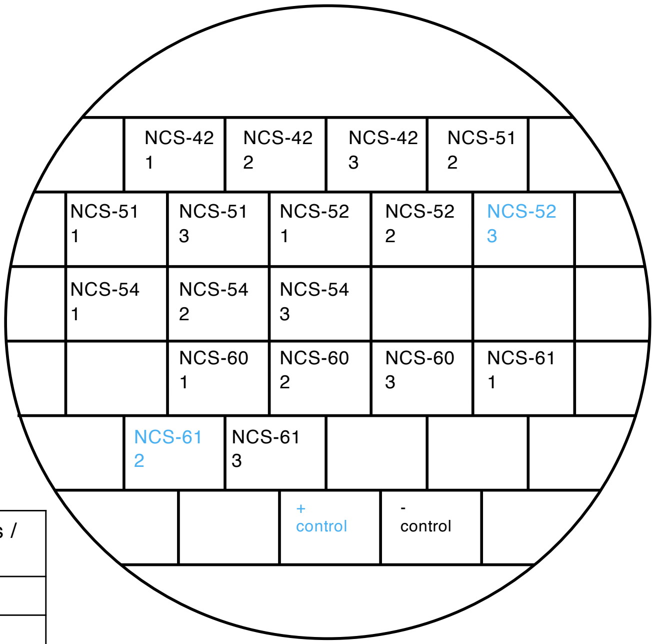


Restreak from YNB –ura –leu to 5-FOA –leu

| Construct | Result regarding viability on 5-FOA | Number of (in)viable clones / total number of clones |
|---|---|---|
| NCS-42 | inviable | 3 out of 3 |
| NCS-51 | inviable | 3 out of 3 |
| NCS-52 | inviable | 2 out of 3 (in total 5 out of 6 see also below) |
| NCS-54 | inviable | 3 out of 3 |
| NCS-60 | inviable | 3 out of 3 |
| NCS-61 | inviable | 2 out of 3 (in total 5 out of 6 see also below) |

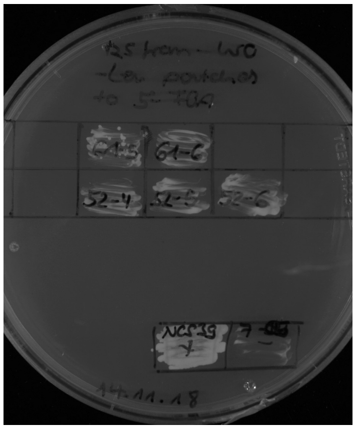

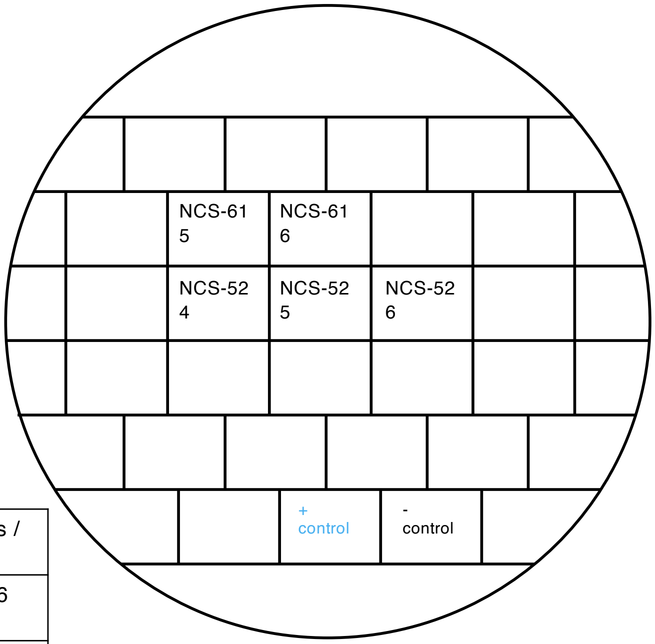


Restreak from YNB –ura –leu to 5-FOA –leu

| Construct | Result regarding viability on 5-FOA | Number of (in)viable clones / total number of clones |
|---|---|---|
| NCS-61 | inviable | 2 out of 2 (in total 5 out of 6 see also above) |
| NCS-52 | inviable | 3 out of 3 (in total 5 out of 6 see also above) |

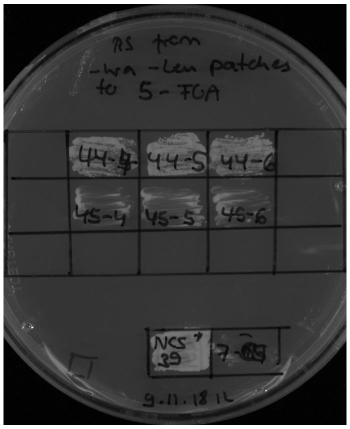


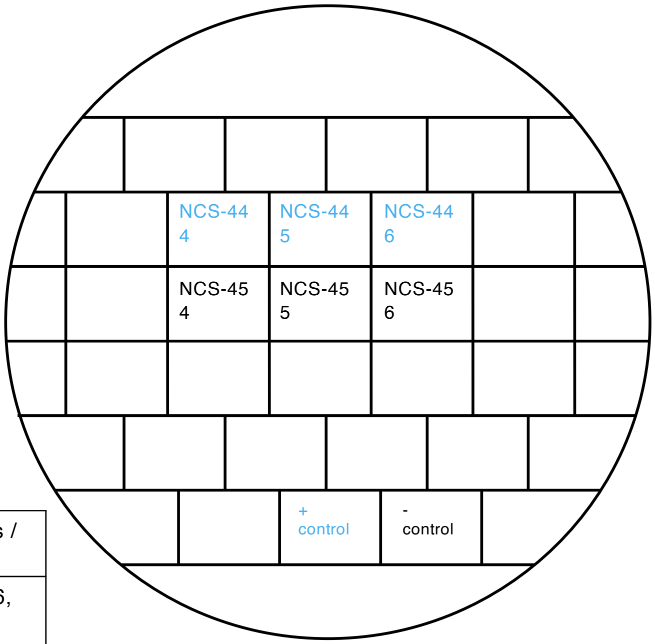


Restreak from YNB –ura –leu to 5-FOA –leu

| Construct | Result regarding viability on 5-FOA | Number of (in)viable clones / total number of clones |
|---|---|---|
| NCS-44 | viable | 3 out of 3 (in total 6 out of 6, technical replicate see also below) |
| NCS-45 | inviable | 3 out of 3 (in total 5 out of 6, see also slide 27) |

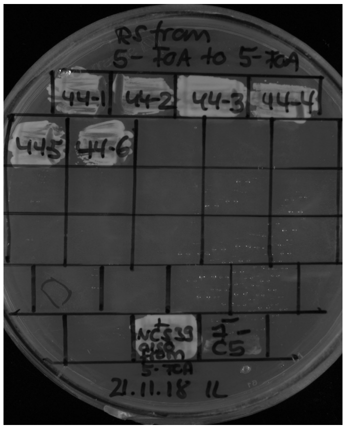


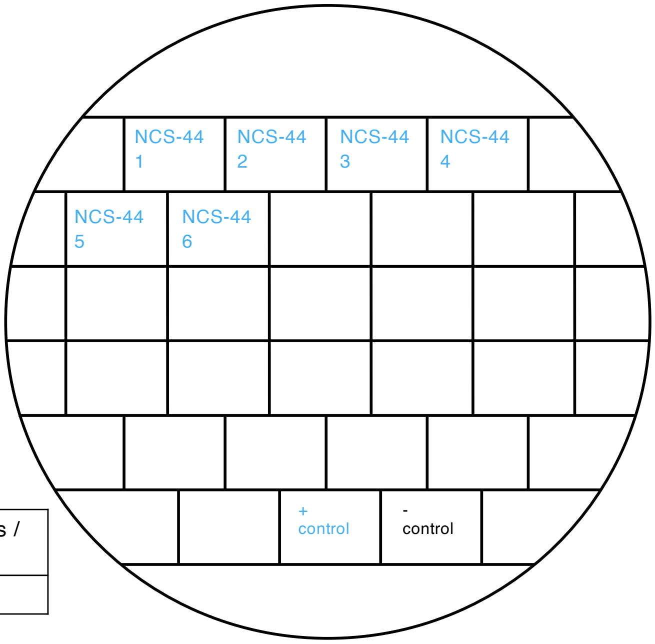


Restreak from 5-FOA – leu to 5-FOA –leu

| Construct | Result regarding viability on 5-FOA | Number of (in)viable clones / total number of clones |
|---|---|---|
| NCS-44 | Viable | 6 out of 6 |

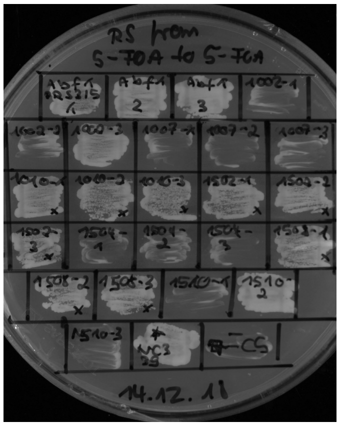


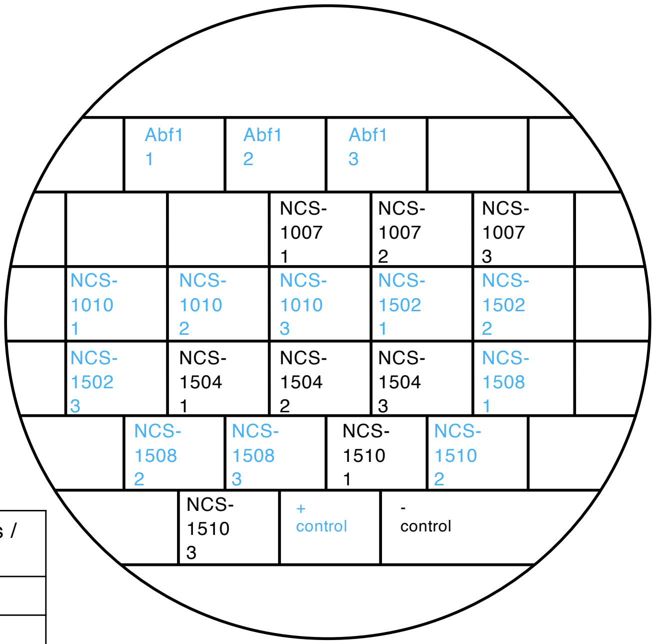


Restreak from YNB –ura –leu to 5-FOA –leu

| Construct | Result regarding viability on 5-FOA | Number of (in)viable clones / total number of clones |
|---|---|---|
| Abf1 | viable | 3 out of 3 |
| NCS-1007 | inviable | 3 out of 3 |
| NCS-1010 | viable | 3 out of 3 |
| NCS-1502 | viable | 3 out of 3 |
| NCS-1504 | inviable | 3 out of 3 |
| NCS-1508 | viable | 3 out of 3 |
| NCS-1510 | inviable (see also slides 31 and 32) | 2 out of 3 (in total 12 out of 15) |

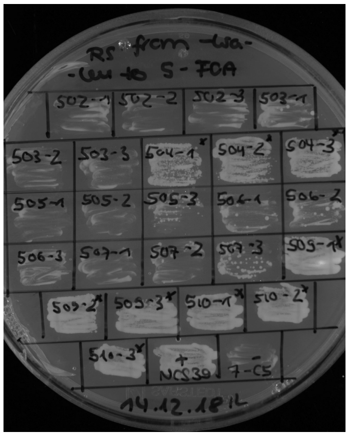


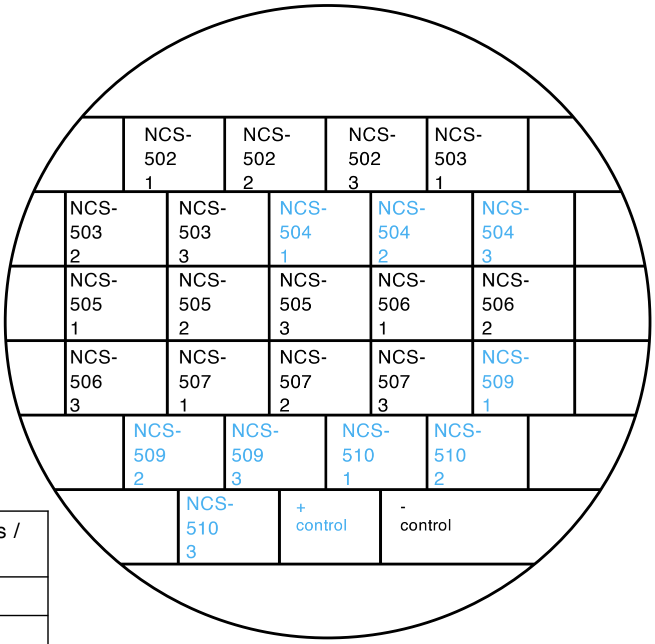


Restreak from YNB –ura –leu to 5-FOA –leu

| Construct | Result regarding viability on 5-FOA | Number of (in)viable clones / total number of clones |
|---|---|---|
| NCS-502 | inviable | 3 out of 3 |
| NCS-503 | inviable | 3 out of 3 |
| NCS-504 | viable | 3 out of 3 |
| NCS-505 | inviable | 3 out of 3 |
| NCS-506 | inviable | 3 out of 3 |
| NCS-507 | inviable | 3 out of 3 |
| NCS-509 | viable | 3 out of 3 |
| NCS-510 | viable | 3 out of 3 |

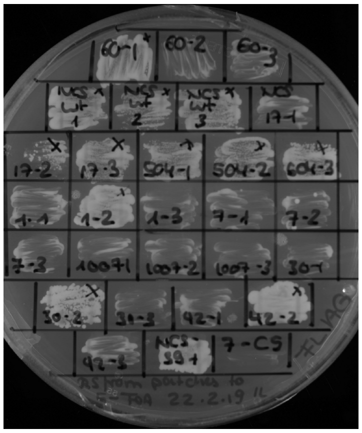

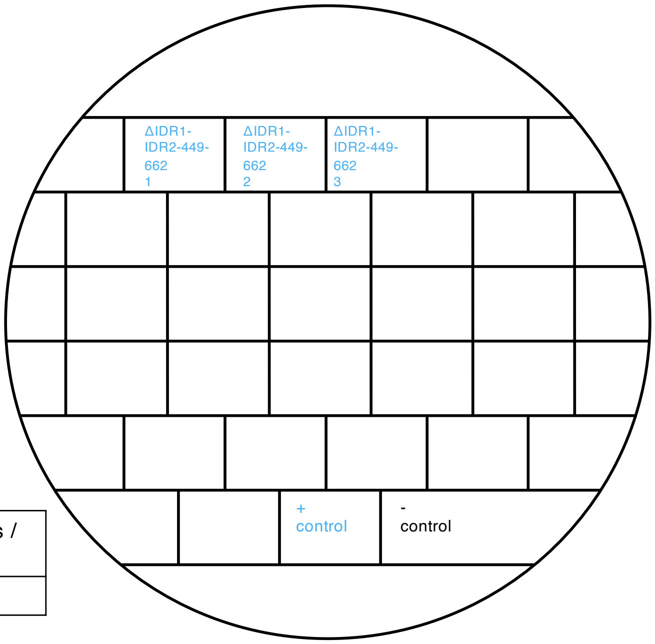


Restreak from YNB –ura –leu to 5-FOA –leu

| Construct | Result regarding viability on 5-FOA | Number of (in)viable clones / total number of clones |
|---|---|---|
| ΔIDR1-IDR2-449-662 | viable | 3 out of 3 |

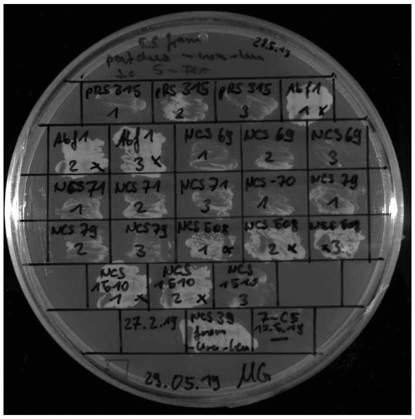

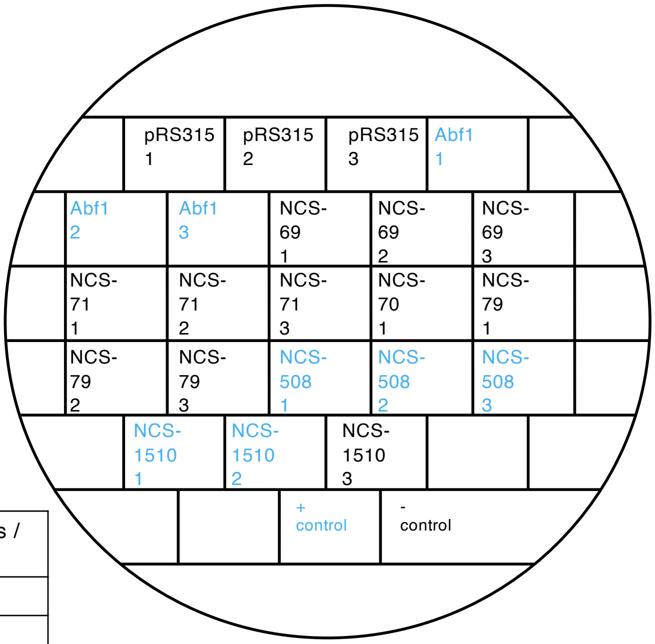


Restreak from YNB –ura –leu to 5-FOA –leu

| Construct | Result regarding viability on 5-FOA | Number of (in)viable clones / total number of clones |
|---|---|---|
| pRS315 | inviable | 3 out of 3 |
| Abf1-NLS-FLAG | viable | 3 out of 3 |
| NCS-69 | inviable | 3 out of 3 |
| NCS-71 | inviable | 3 out of 3 |
| NCS-70 | inviable | 1 out of 1 (in total 3 out of 3, see slide 32) |
| NCS-79 | inviable | 3 out of 3 |
| NCS-508 | viable | 3 out of 3 |
| NCS-1510 | inviable (see also slides 30 and 32) | 1 out of 3 (in total: 12 out of 15) |

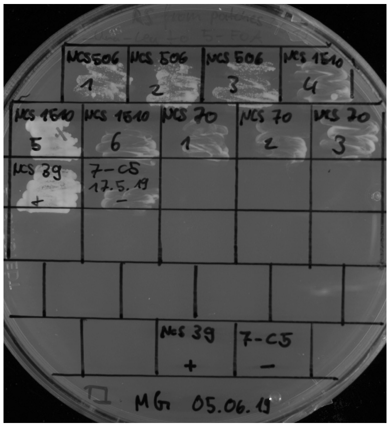

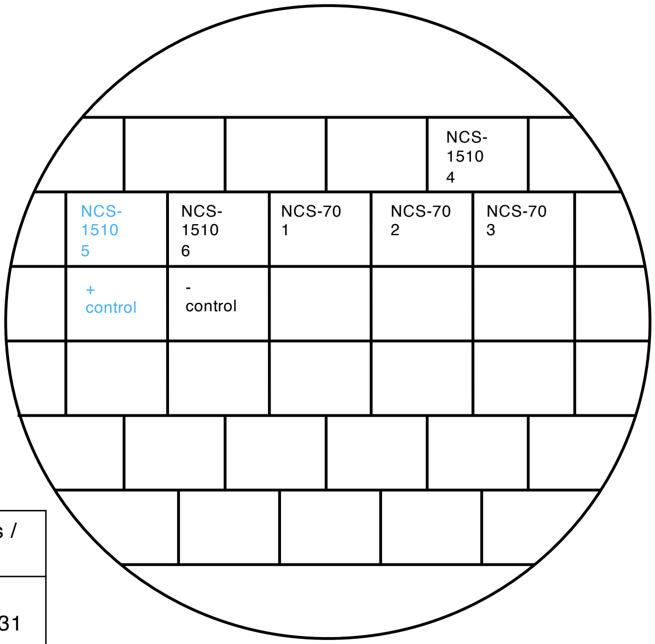


Restreak from YNB –ura –leu to 5-FOA –leu

| Construct | Result regarding viability on 5-FOA | Number of (in)viable clones / total number of clones |
|---|---|---|
| NCS-1510 | inviable | 2 out of 3 (in total 12 out of 15, see also slides 30 and 31 and below) |
| NCS-70 | inviable | 3 out of 3 (in total 3 out of 3, see slide 31) |

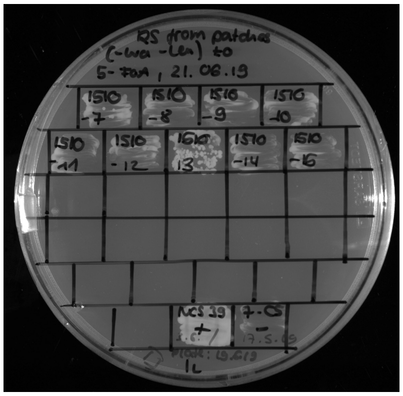

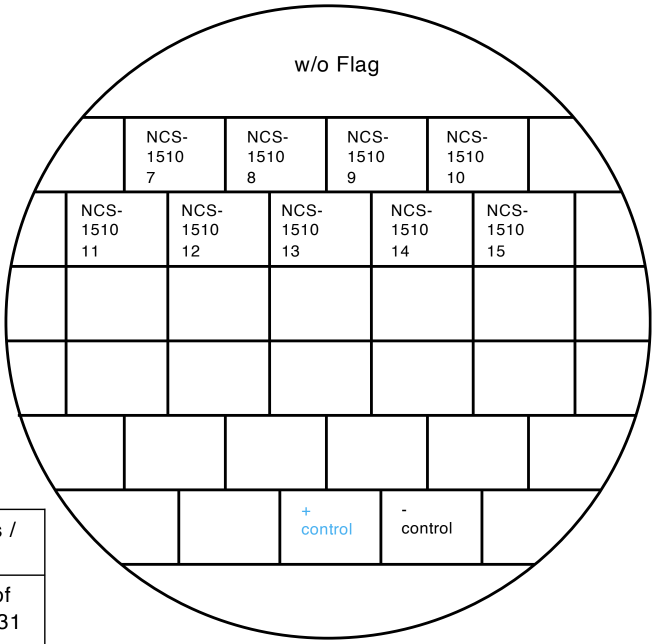


Restreak from YNB –ura –leu to 5-FOA –leu

| Construct | Result regarding viability on 5-FOA | Number of (in)viable clones / total number of clones |
|---|---|---|
| NCS-1510 | inviable | 9 out of 9 (in total: 12 out of 15, see also slides 30 and 31 and above) |

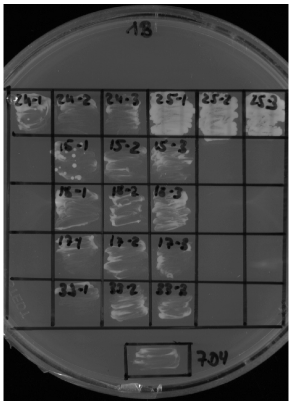

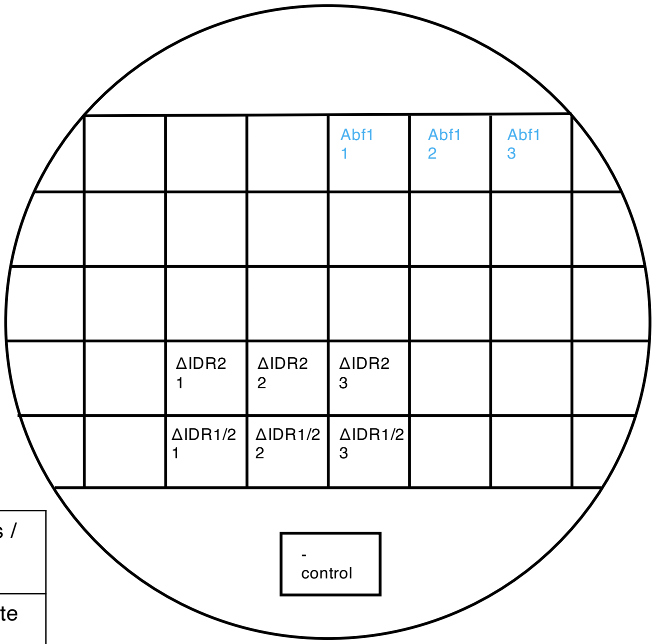


Restreak from YNB –ura –leu to 5-FOA –leu

| Construct | Result regarding viability on 5-FOA | Number of (in)viable clones / total number of clones |
|---|---|---|
| Abf1 (NLS) | viable | 3 out of 3 (technical replicate see below) |
| ΔIDR2 | inviable | 3 out of 3 |
| ΔIDR1/2 | inviable | 3 out of 3 |

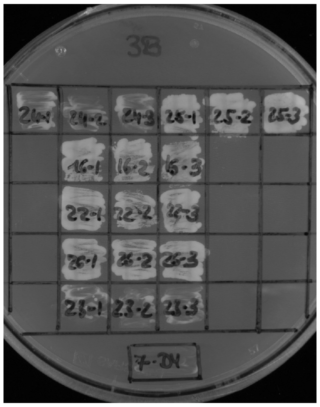

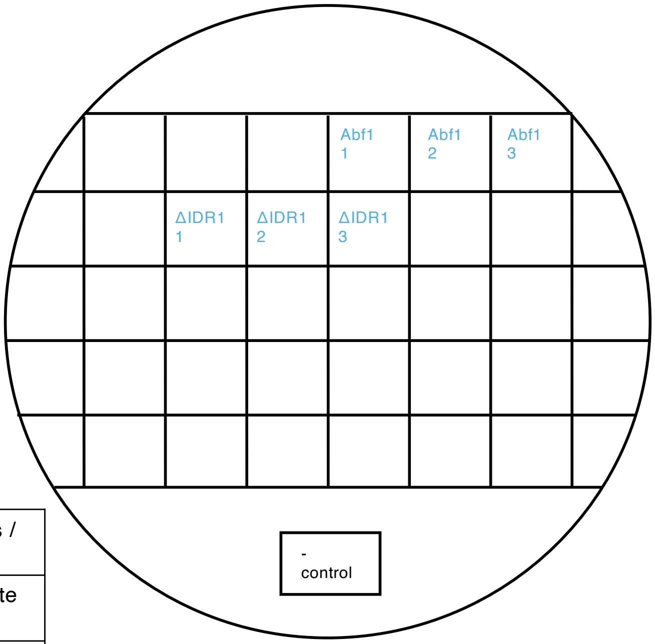


Restreak from YNB –ura –leu to 5-FOA –leu

| Construct | Result regarding viability on 5-FOA | Number of (in)viable clones / total number of clones |
|---|---|---|
| Abf1 (NLS) | viable | 3 out of 3 (technical replicate see above) |
| ΔIDR1 | viable | 3 out of 3 |

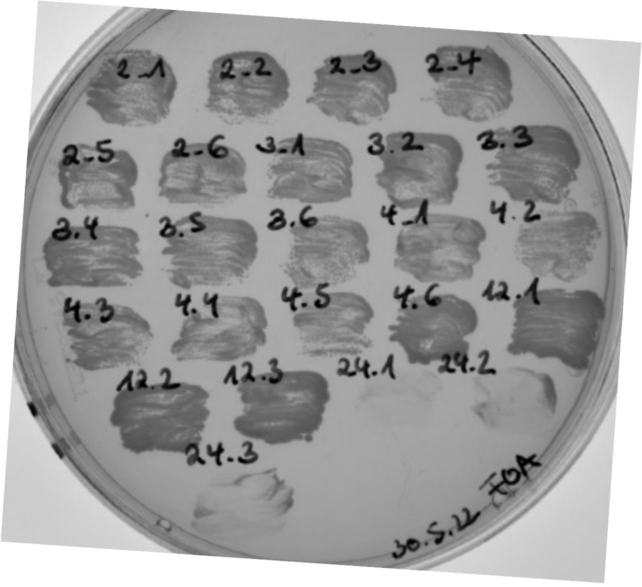


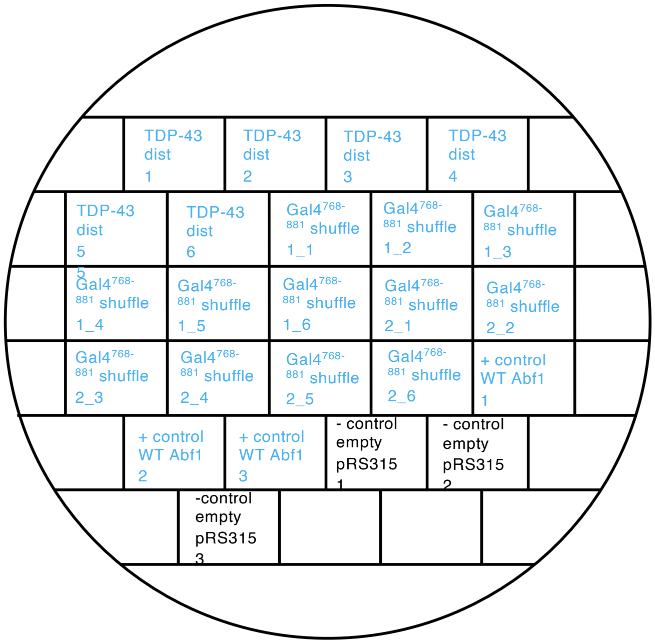


Restreak from YNB –ura –leu to 5-FOA –leu

| Construct | Result regarding viability on 5-FOA | Number of (in)viable clones / total number of clones |
|---|---|---|
| TDP-43 dist | viable | 6 out of 6 |
| Gal4$^{768-881}$ shuffle 1 | viable | 6 out of 6 |
| Gal4$^{768-881}$ shuffle 2 | viable | 6 out of 6 |

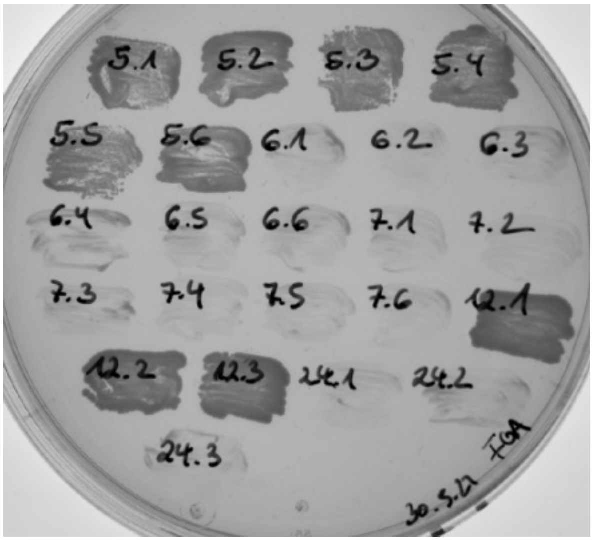


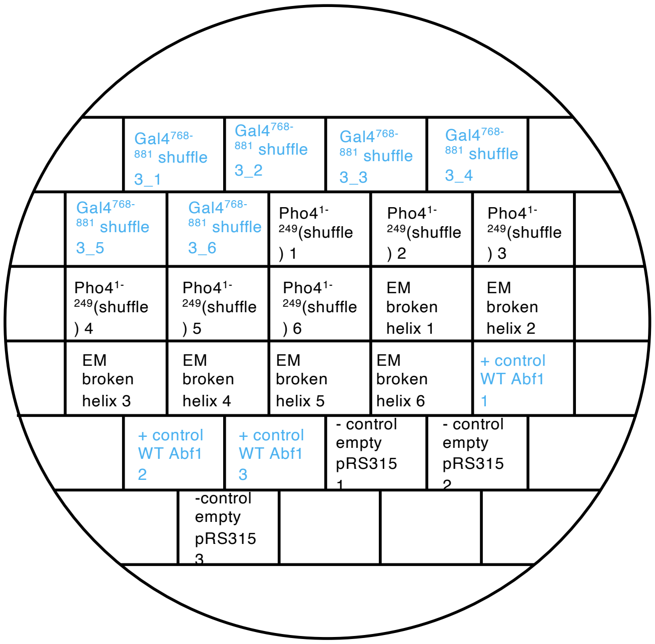


Restreak from YNB –ura –leu to 5-FOA –leu

| Construct | Result regarding viability on 5-FOA | Number of (in)viable clones / total number of clones |
|---|---|---|
| Gal4$^{768-881}$ shuffle 3 | viable | 6 out of 6 |
| Pho4$^{1-249}$(shuffle) | inviable | 6 out of 6 |
| EM broken helix | inviable | 6 out of 6 |

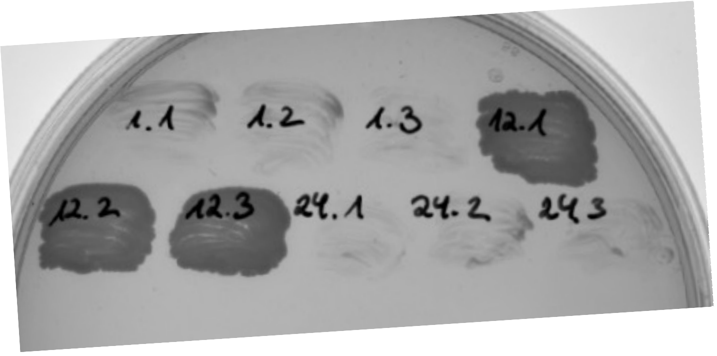


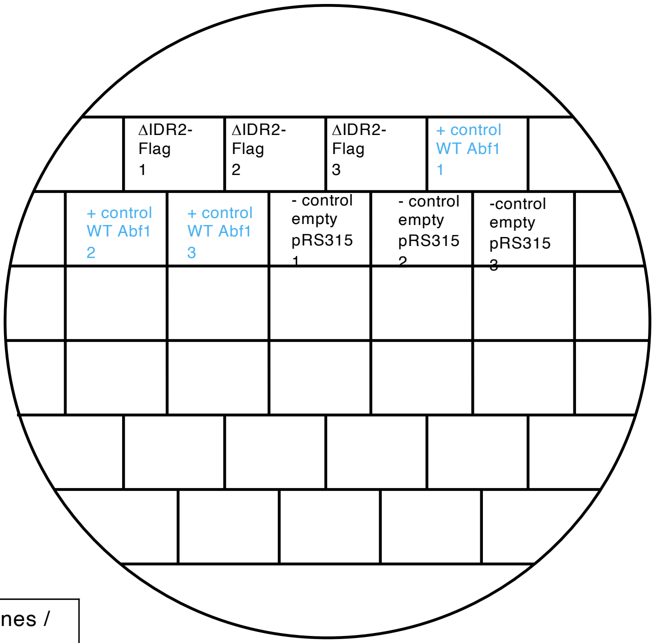


Restreak from YNB –ura –leu to 5-FOA –leu

| Construct | Result regarding viability on 5-FOA | Number of (in)viable clones / total number of clones |
|---|---|---|
| ΔIDR2-Flag | inviable | 3 out of 3 |
